# 40 new specimens of *Ichthyornis* provide unprecedented insight into the postcranial morphology of crownward stem group birds

**DOI:** 10.1101/2022.01.11.475364

**Authors:** Juan Benito, Albert Chen, Laura E. Wilson, Bhart-Anjan S. Bhullar, David Burnham, Daniel J. Field

**Affiliations:** Department of Earth Sciences, University of Cambridge, Cambridge, UK; Department of Biology & Biochemistry, Milner Centre for Evolution, University of Bath, Bath, UK; Sternberg Museum of Natural History and Department of Geosciences, Fort Hays State University, Hays, KS, USA; Yale Peabody Museum of Natural History, New Haven, CT, USA; Department of Earth & Planetary Sciences, Yale University, New Haven, CT, USA; Biodiversity Institute and Natural History Museum, University of Kansas, Lawrence, KS, USA; University Museum of Zoology, Cambridge, UK

## Abstract

*Ichthyornis* has long been recognized as a pivotally important fossil taxon for understanding the latest stages of the dinosaur–bird transition, but little significant new postcranial material has been brought to light since initial descriptions of partial skeletons in the 19^th^ Century. Here, we present new information on the postcranial morphology of *Ichthyornis* from 40 previously undescribed specimens, providing the most detailed morphological assessment of *Ichthyornis* to date. The new material includes four partially complete skeletons and numerous well-preserved isolated elements, enabling new anatomical observations such as muscle attachments previously undescribed for Mesozoic euornitheans. Among the elements that were previously unknown or poorly represented for *Ichthyornis*, the new specimens include an almost-complete axial series, a hypocleideum-bearing furcula, radial carpal bones, fibulae, a complete tarsometatarsus bearing a rudimentary hypotarsus, and one of the first-known nearly complete three-dimensional sterna from a Mesozoic avialan. Several pedal phalanges are preserved, revealing a remarkably enlarged pes presumably related to foot-propelled swimming. Although diagnosable as *Ichthyornis*, the new specimens exhibit a substantial degree of morphological variation, some of which may relate to ontogenetic changes. Phylogenetic analyses incorporating our new data and employing alternative morphological datasets recover *Ichthyornis* stemward of Hesperornithes and *Iaceornis*, in line with some recent hypotheses regarding the topology of the crownward-most portion of the avian stem group, and we establish phylogenetically-defined clade names for relevant avialan subclades to help facilitate consistent discourse in future work. The new information provided by these specimens improves our understanding of morphological evolution among the crownward-most non-neornithine avialans immediately preceding the origin of crown group birds.

## INTRODUCTION

*Ichthyornis* is a key taxon in the history of avian palaeontology. First described by Marsh in 1872, early specimens of *Ichthyornis* provided some of the first known fossil evidence documenting the evolutionary origins of birds, thereby bolstering evolutionary theory in the late 19^th^ Century. Indeed, “Darwin’s Bulldog”, Thomas Henry Huxley, declared—presumably in partial reference to *Ichthyornis* and *Hesperornis*—that:

*“There is nothing in any way comparable … for their scientific importance, to the series of fossils which Professor Marsh has brought together.”* (Rieppel, 2019).

And Darwin himself complimented Marsh by writing:

*“Your work on these old birds, & on the many fossil animals of N. America has afforded the best support to the theory of evolution, which has appeared within the last 20 years.*” (Burkhardt, 2020).

Now, almost 150 years later, *Ichthyornis* remains a key taxon for understanding the morphological transitions that gave rise to crown bird anatomy—or, as O.C. Marsh put it, “to break down the old distinction between Birds and Reptiles” (Marsh, 1873), and it has been consistently recovered in a phylogenetic position close to the origin of the avian crown group (e.g., Clarke, 2004; Field et al., 2018a).

### Previous research on Ichthyornis

Fossil remains attributable to *Ichthyornis* have been commonly recovered from Late Cretaceous deposits of the Western Interior Seaway of North America, usually from the middle to late Santonian rocks of the Niobrara Formation in Kansas, USA (Marsh, 1972a, 1880; Clarke, 2004; Field et al., 2018a) and from the early Campanian deposits of the Mooreville Chalk in Alabama, USA (Wetmore, 1962; Olson, 1975, Field et al., 2018a). Additional material has been recovered from the Cenomanian of Saskatchewan, Canada (Tokaryk et al.,1997, Sanchez, 2010) and Kansas (Shimada & Wilson, 2016), the Turonian of Alberta, Canada (Fox, 1984), Kansas (Shimada & Fernandes, 2006), and New Mexico, USA (Lucas & Sullivan, 1982), the Campanian of Texas, USA (Parris & Echols, 1992) and the Coniacian–Campanian of Coahuila, Mexico (Porras-Muzquiz et al., 2014). Fragmentary fossil material showing similarities to *Ichthyornis* has additionally been recovered from the Cenomanian of Russia (Zelenkov et al., 2017) and Egypt (Mohesn et al., 2020), and the Maastrichtian of Belgium (Dyke et al., 2002), but the precise phylogenetic relationships of these remains have yet to be assessed in detail.

Eight different species of *Ichthyornis* have been erected in the past on the basis of specimens from the Niobrara and Mooreville Formations (Marsh, 1872a, 1873, 1876, 1880; Wetmore, 1962). However, despite the substantial geographic and temporal distribution of *Ichthyornis* (ranging in age from 95MYA to 83.5MYA), Clarke (2004) found no discrete morphological differences among the YPM *Ichthyornis* specimens to substantiate their designation as distinct species, and therefore synonymized 5 of the 8 previously named species into *Ichthyornis dispar* in addition to providing a detailed definition and diagnosis of this taxon. One of the remaining species was separated into its own genus, *Guildavis,* while the specimens belonging to *I. celer*, already assigned to their own genus, *Apatornis*, by Marsh (1880), were separated into two distinct taxa, *A. celer* and *Iaceornis marshi*.

The comprehensive study by Clarke (2004) reevaluated all of the 19^th^ Century specimens of *Ichthyornis* housed at the Yale Peabody Museum (YPM), greatly updating our understanding of the taxon and clarifying its morphological differences with respect to more stemward Mesozoic avialans and the avian crown group. However, the fragmentary nature of many YPM specimens seriously hinders our understanding of *Ichthyornis* postcranial anatomy, particularly in key skeletal regions such as the sternum, pelvis, and hindlimbs. Unfortunately, only a small fraction of the YPM material was clearly figured in Clarke (2004) in order to avoid duplicating many illustrations of specimens that were initially figured by Marsh (1880). However, Clarke (2004) noted several inaccuracies in these 19^th^ Century illustrations; thus, the relative dearth of unambiguous images of *Ichthyornis* postcranial morphology limits the availability of anatomical data on this key taxon for comparative morphological studies.

A substantial amount of new cranial material belonging to *Ichthyornis* has recently been described (Field et al., 2018a; Torres et al., 2021), with recent work on *Ichthyornis* predominantly targeting questions related to jaw morphology, the origin of the neornithine beak, and brain architecture (Gingerich, 1972; Martin & Stewart 1977; Dumont et al., 2016; Field et al., 2018; Brocklehurst & Field, 2021; Torres et al., 2021). By contrast, postcranial material reported since the original descriptions of *Ichthyornis* in the 19^th^ Century is limited, consisting mostly of several isolated humeri and other highly fragmentary elements (e.g., Olson, 1975; Fox, 1984; Porras-Muzquiz, 2014; Shimada & Wilson, 2016). As a result, our knowledge of the postcranial osteology of *Ichthyornis* still primarily rests on the material originally described by Marsh (1872a, 1880) and redescribed by Clarke (2004), housed in the collections of the YPM.

Importantly, the postcranial anatomy of *Ichthyornis* has yet to be reinvestigated in light of a surge of crownward euornithean discoveries over the last two decades. Since the redescription of *Ichthyornis* by Clarke (2004), a wealth of recently described taxa, mainly from the Early Cretaceous of China has shed much needed light on the diversity and morphology of Mesozoic avialans, and particularly on Euornithes, the avialan subclade including crown birds (Neornithes) and their closest relatives (Pittman et al., 2020a,b). These discoveries have documented the gradual evolutionary acquisition of crown-bird-like postcranial anatomy, revealing an increasingly complex picture of euornithean evolution, and affirming the phylogenetic position of *Ichthyornis* as among the most crownward Mesozoic avialans yet known. Newly-recognized euornitheans that have been described since the last substantial work on *Ichthyornis* postcranial morphology include clades such as Hongshanornithidae (Zhou & Zhang, 2005; O’Connor et al., 2010; Chiappe et al., 2014; Wang et al., 2016a), Songlingornithidae (or Yanornithidae; Zhou & Zhang, 2001; Clarke et al., 2006; Zheng et al., 2014; Wang et al., 2013, 2019, 2020, 2021), and Schizoouridae (Zhou et al., 2012; Wang et al., 2020). Of particular relevance are taxa such as *Gansus* (You et al., 2006; Wang et al., 2016b) and *Iteravis* (Liu et al., 2014; Zhou et al., 2014; O’Connor et al., 2015; Wang et al., 2018), which have been recovered in phylogenetic positions close to *Ichthyornis* along the most crownward portion of the avialan stem lineage. In light of these recent discoveries, renewed investigations into the morphology of *Ichthyornis* may provide important insights into key morphological transitions immediately preceding the origin and diversification of the avian crown group, as well as the refinement of powered flight capacity among Mesozoic avialans (Pittman et al. 2020c).

Recently, the description of a substantial amount of new skull material from *Ichthyornis*, aided by high-resolution µCT imaging, revealed a striking mosaic of crown bird-like and plesiomorphic avialan features (Field et al., 2018a; Torres et al., 2021). This transitional cranial architecture departs in important ways from earlier, largely hypothetical reconstructions of the *Ichthyornis* skull on the basis of poorly preserved, incomplete remains (e.g., Marsh, 1880). Despite this recent advance in our understanding of its skull and jaws, much about the postcranial morphology of *Ichthyornis* remains unknown. Here, we investigate the postcranial morphology of numerous new *Ichthyornis* specimens, including those whose cranial material was previously described by Field et al., (2018a). The new information revealed here has enabled us to reconstruct the postcranial skeletal morphology of *Ichthyornis* in unprecedented detail, leaving only a small number of minor skeletal components unknown for this taxon.

### Phylogenetic interrelationships of Cretaceous euornitheans

The exact phylogenetic position of *Ichthyornis* with respect to Neornithes and other Ornithurae (the most exclusive clade uniting Ichthyornithes, Hesperornithes, and Neornithes; see below for full phylogenetic definitions) remains controversial (Pittman et al., 2020b), but *Ichthyornis* has been consistently recovered in a phylogenetic position close to the origin of crown group birds (Clarke et al., 2006; O’Connor et al., 2011, 2016; Huang et al., 2016; Wang et al., 2016b, 2017, 2020a,b,c; Atterholt et al., 2018; Field et al., 2018a; Zheng et al., 2018; Torres et al., 2021). A few additional taxa, such as *Apsaravis* (Clarke & Norell, 2002), *Ambiortus* (Kurochkin, 1985; O’Connor & Zelenkov, 2013), *Hollanda* (Bell et al., 2010) and *Patagopteryx* (Chiappe, 1996, 2002) have occasionally been recovered within Ornithurae, close to *Ichthyornis* and Hesperornithes, but these results have not been consistently recovered in most studies (O’Connor & Zelenkov, 2013; Field et al., 2018a; Pittman et al., 2020b; Wang et al., 2020a,b). Recent analyses have recovered alternative phylogenetic positions for *Ichthyornis* with respect to the diving Hesperornithes, which have been recovered in a position either slightly crownward of (O’Connor et al., 2011, 2020; Wang et al., 2017, 2019; Atterholt et al., 2018; Field et al., 2018a), slightly stemward (Chiappe, 2002; Clarke, 2004; You et al., 2006; Huang et al., 2016; Wang et al., 2020b; Torres et al., 2021) or in an unresolved polytomy (Wang et al., 2021) with *Ichthyornis* (Fig. 1). The relative position of both groups is highly sensitive to both the dataset and the methods used in phylogenetic analyses (Wang et al., 2017; Field et al., 2018a, Pittman et al., 2020b). Moreover, numerous postcranial character states have been impossible or very difficult to score with accuracy for *Ichthyornis*, due to a lack of suitably complete and well-preserved material, highlighting the need for additional data on *Ichthyornis* in order to recover a more consistent phylogenetic topology for the most crownward portion of the avian stem lineage. Other than Hesperornithes, very few Mesozoic euornithean stem-birds have been recovered in a phylogenetic position crownward of *Ichthyornis*. Such taxa, such as *Guildavis, Apatornis*, and *Iaceornis* (Clarke, 2004), as well as *Limenavis* (Clarke & Chiappe, 2001), are generally based on highly fragmentary material, which limits their informativeness and complicates the assessment of phylogenetic interrelationships among the crownward-most stem birds.

**Figure 1.**
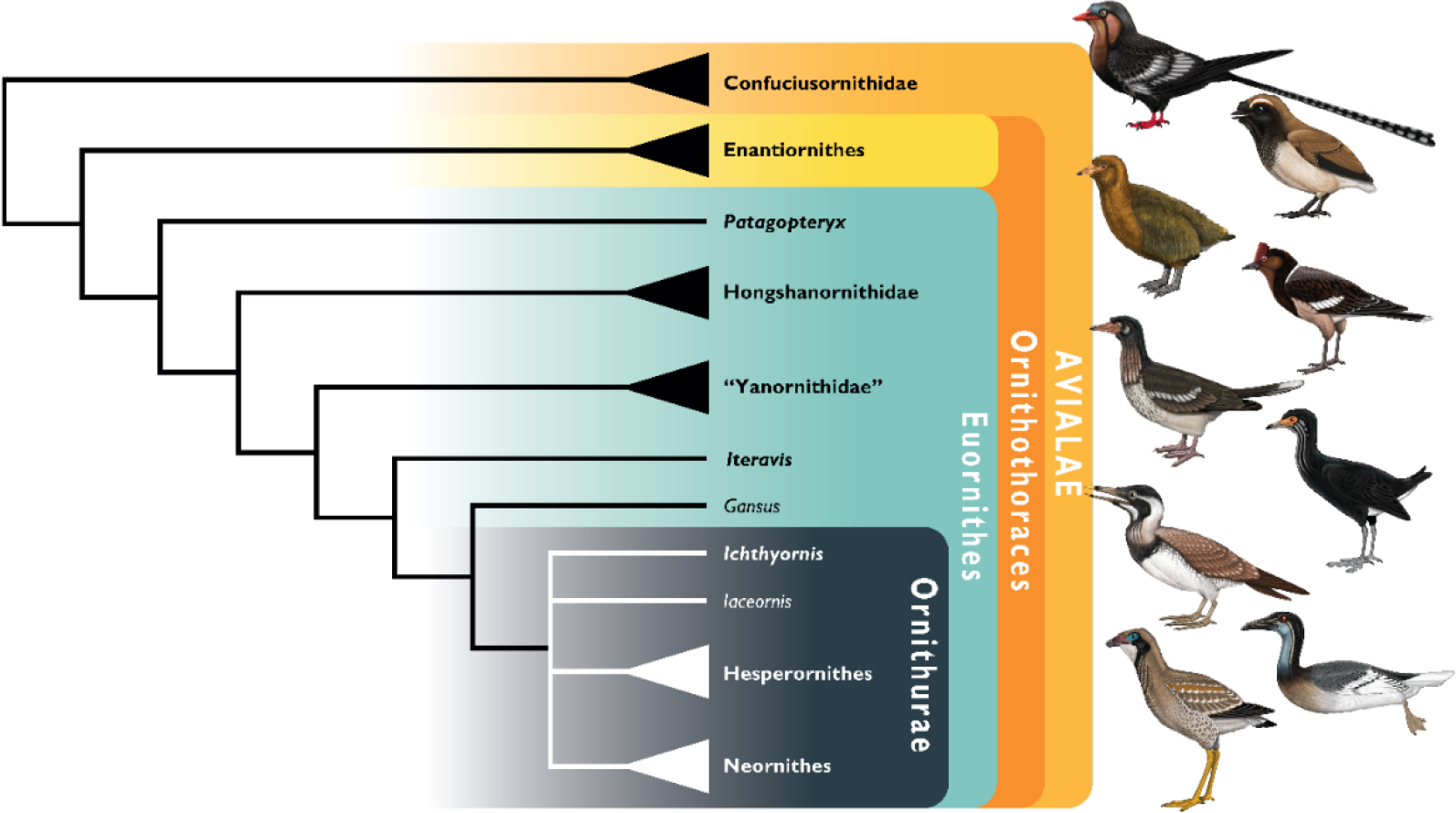
Simplified cladogram showing the most commonly recovered phylogenetic positions of *Ichthyornis* and relevant Mesozoic avialans. White branches highlight phylogenetic uncertainty within the major clade Ornithurae, which includes Ichthyornis, Hesperornithes, and the bird crown group (Neornithes).. Taxa in bold are figured, superscript numbers indicate the corresponding illustration. For Confuciusornithidae, the illustration corresponds to *Eoconfuciusornis zhengi*; for Enantiornithes, the illustration corresponds to *Cathayornis yandica*; for Hongshanornithidae the illustration corresponds to *Hongshanornis longicresta*; for “Yanornithidae” the illustration corresponds to *Abitusavis lii*; for Hesperornithes the illustration corresponds to *Brodavis varneri*; for Neornithes the illustration corresponds to *Asteriornis maastrichtensis.* Illustrations courtesy of R. Olive, used with permission.

### Reconstructing the morphology of the earliest crown birds

The origin of the bird crown group is well-established to have occurred during the Cretaceous Period (Jarvis et al., 2014; Prum et al., 2015; Berv & Field, 2018), but the scarcity of Late Cretaceous crown bird material complicates our understanding of the early morphology and evolutionary history of the group (Chatterjee, 1989, 2000; Clarke et al., 2005, 2016; Longrich et al. 2011; Field et al., 2020a,b). Given this significant gap in the crown bird fossil record, work attempting to understand aspects of the ecology, biology, and morphology of the earliest crown birds must rely on inferences based on extant birds and the most crownward-known stem birds (Zheng et al., 2014, 2018; Berv & Field, 2018; Field et al., 2018b; Wang et al., 2018; O’Connor, 2019; O’Connor & Zhou, 2015, 2020; Torres et al., 2021). Thus, an improved understanding of the morphology of the closest relatives of crown birds from the Late Cretaceous is pivotal for reconstructing the nature of the earliest Neornithes. Unfortunately, despite their abundance and their crownward position among Mesozoic Ornithurae, Hesperornithes were secondarily flightless, exhibiting highly specialized, autapomorphic postcranial features including greatly reduced wings, strongly modified hindlimbs for foot-propelled diving, and osteosclerotic skeletons. These specialized features preclude the use of many aspects of hesperornithine postcranial osteology as a reliable source for reconstructions of the plesiomorphic condition of the avian crown group (Bell & Chiappe, 2016). By contrast, *Ichthyornis* was obviously less ecologically specialized than hesperornithines, and easily falls within the size range of extant volant marine birds.

These features suggest that the morphology of *Ichthyornis* provides a more useful approximation of the ancestral condition of the crown bird postcranium (Clarke, 2004; Field et al., 2018a), with some aspects of *Ichthyornis* postcranial morphology hypothesized to fall within the range of variation of extant bird diversity (Mayr, 2017). Given the scarcity of Mesozoic fossil material recovered crownward of *Ichthyornis* and Hesperornithes, the postcranial morphology of *Ichthyornis* may be more representative of the ancestral condition of crown birds than that of any other known Mesozoic avialan; thus, its study has crucial implications for understanding morphological evolution immediately preceding the great radiation of the avian crown group.

### Focus of the present study

Despite the substantial number of well-preserved Mesozoic euornitheans that have recently been described, its crownward phylogenetic position continues to render *Ichthyornis* a key taxon in our understanding of avian evolution, and the abundance of its remains makes it almost unique in its potential for revealing important insights into avialan intraspecific variation (Clarke, 2004). Here, we substantially advance our understanding of the morphology of this pivotal taxon by describing three-dimensional µCT scans of the postcranial morphology of 40 new specimens of *Ichthyornis,* including four substantially complete partial skeletons with recently-described cranial remains (Field et al., 2018a). The new material includes several skeletal elements that have not previously been described for *Ichthyornis*, including the radial carpal and the fibula, as well as significantly better-preserved examples of elements previously known from highly fragmentary remains, such as the sternum, the furcula, the pelvis, the tibiotarsus, and the foot (Fig. 2). Many of the new specimens are exceptionally well-preserved in three dimensions, exceeding the completeness and degree of preservation of much of the classic YPM material. Together, the new material offers a nearly complete view of *Ichthyornis* postcranial osteology (Fig. 3), facilitating a detailed reinvestigation of the phylogenetic position of *Ichthyornis* among Mesozoic Avialae.

**Figure 2.**
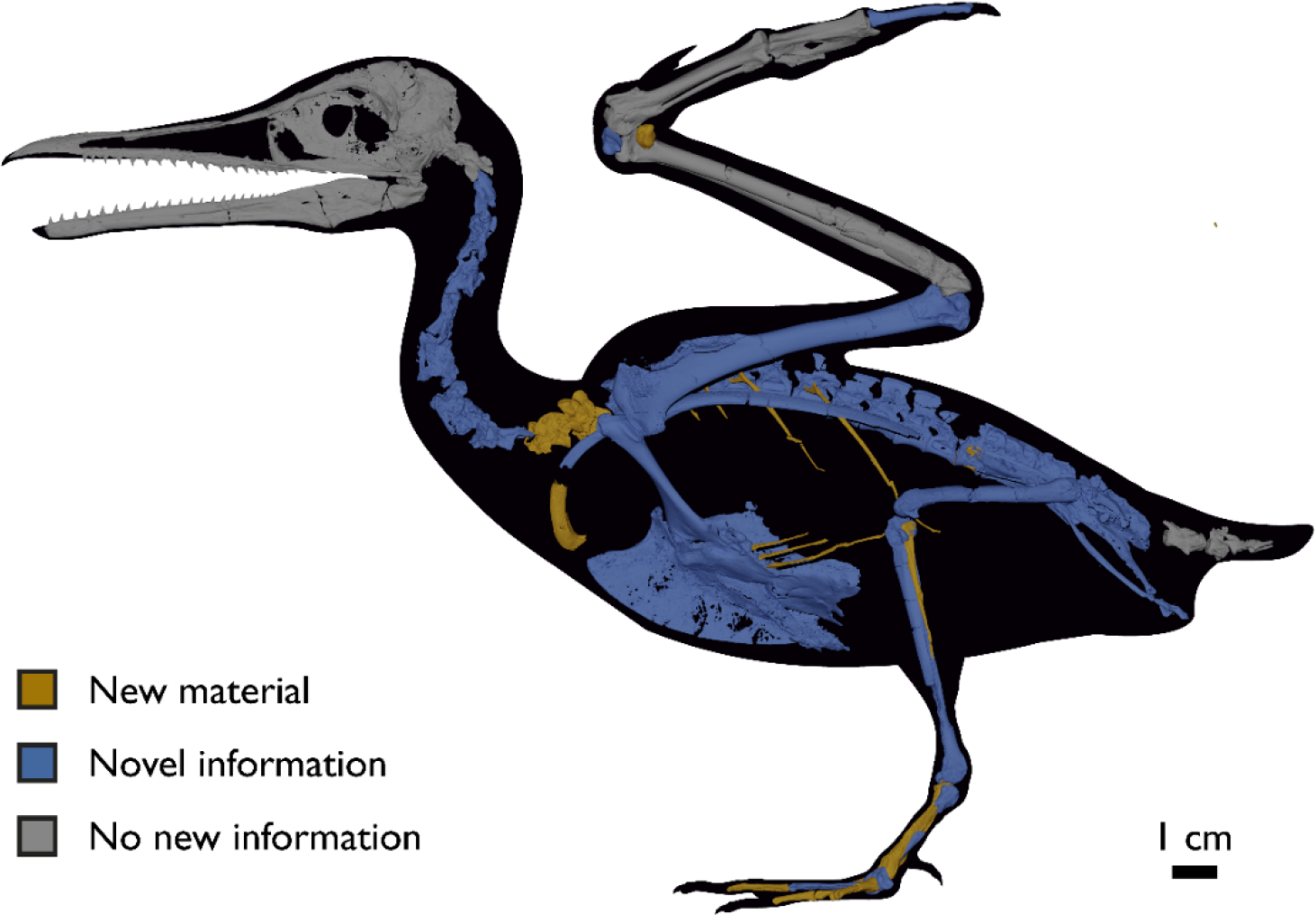
Reconstruction of the skeleton of *Ichthyornis dispar*, showing elements described in the present study that exhibit novel morphological information for *Ichthyornis*. The reconstructed skeleton is a composite incorporating multiple specimens described in this study (see Table 1). All specimens are scaled to the dimensions of the FHSM VP-18702 specimen.

**Figure 3.**
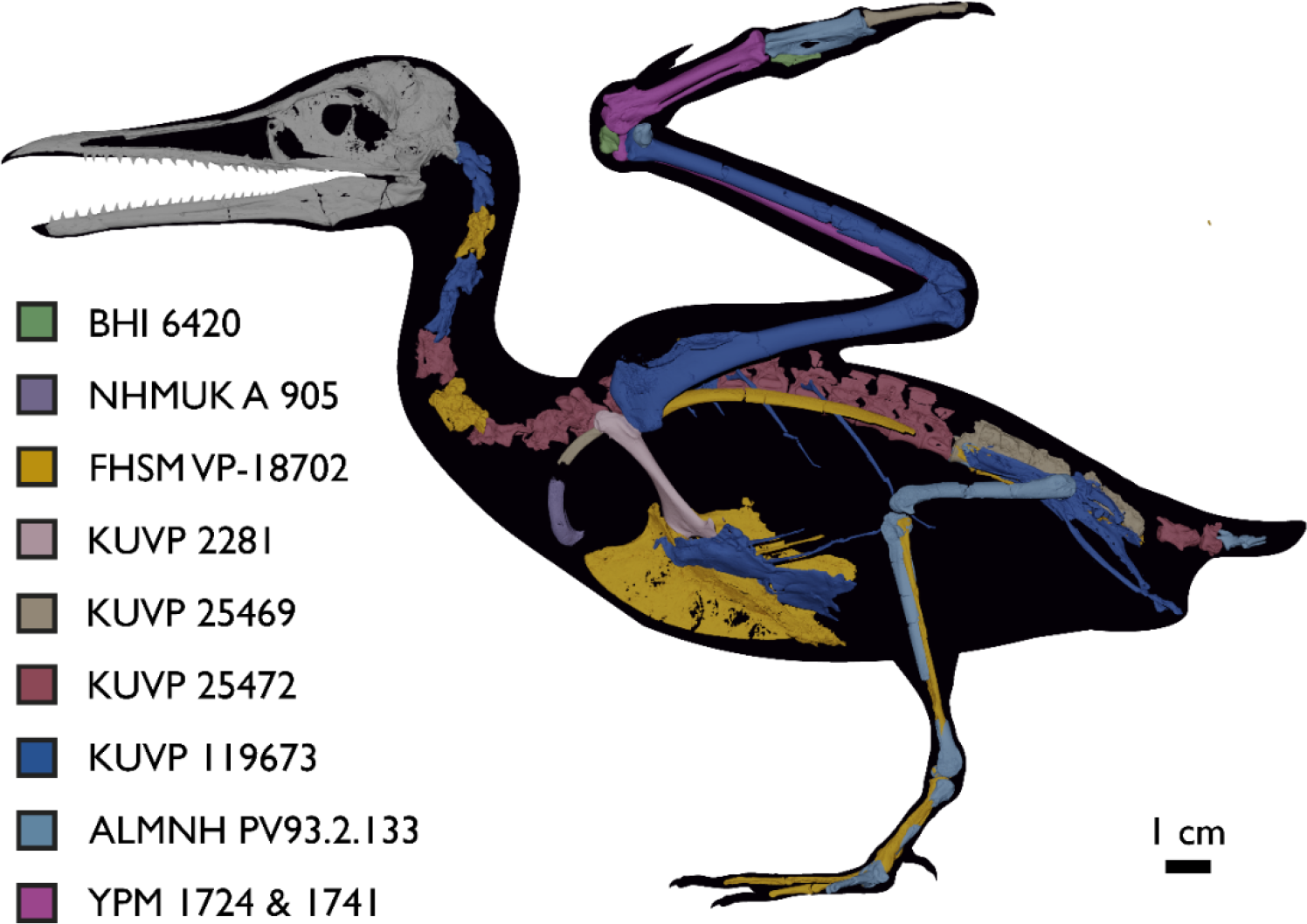
Reconstruction of the skeleton of *Ichthyornis dispar*, showing elements that are described here for the first time. The reconstructed skeleton is a composite incorporating numerous new specimens described in this study (see colour-coded legend and Table 1). All specimens are scaled to the dimensions of FHSM VP-18702.

**Table 1.** List of specimens included in this study. All specimens, with the exception of those from the YPM collections and the skull of FHSM VP 18702, are newly described in the present study. The elements preserved in each specimen are listed; see description and Supplementary Information for additional information. Synapomorphies diagnosing *Ichthyornis* from Clarke (2004) that are recognizable in each specimen are indicated. Where none of these apomorphies are preserved, specimens were identified based on morphological similarity with other diagnosed *Ichthyornis* specimens. Synapomorphies: (**1**) a single large pneumatic foramen situated on the anteromedial surface of the quadrate, (**2**) amphicoelous cervical vertebrae, (**3**) free caudal vertebrae exhibiting well-developed and elongated prezygapophyses, (**4**) scapula exhibiting an extremely diminutive acromion process, (**5**) pit-shaped fossa on the distal end of the bicipital crest of the humerus, (**6**) ulnar trochlear surface equal in length across its caudal and distal surfaces, (**7**) oval scar located on the caudoventral surface of the distal radius, (**8**) large tubercle developed close to the articular surface of phalanx II:1 in the carpometacarpus and (**9**) presence of an internal index process on the distal end of manual phalanx II:1.

| Specimen Number | Elements represented | Origin | Age | Discrete synapomorphies supporting referral to *Ichthyornis* |
| --- | --- | --- | --- | --- |
| ALMNH PV 1988.20.180 | Distal tarsometatarsus | Mooreville Fm, Alabama | Early Campanian | - |
| ALMNH PV 1988.20.181 | Distal tarsometatarsus | Mooreville Fm, Alabama | Early Campanian | - |
| ALMNH PV 1988.20.183.1 | Distal femur | Mooreville Fm, Alabama | Early Campanian | - |
| ALMNH PV 1988.20.186 | Proximal femur | Mooreville Fm, Alabama | Early Campanian | - |
| ALMNH PV 1988.20.375.3 | Proximal tarsometatarsus | Mooreville Fm, Alabama | Early Campanian | - |
| ALMNH PV 1988.20.427.1 | Proximal humerus | Mooreville Fm, Alabama | Early Campanian | 5 |
| ALMNH PV 1988.20.5 | Proximal humerus & bone fragment | Mooreville Fm, Alabama | Early Campanian | 5 |
| ALMNH PV 1988.26.1 | Possible pedal phalanx | Mooreville Fm, Alabama | Early Campanian | - |
| ALMNH PV 1993.2.191.1 | Distal tibiotarsus | Mooreville Fm, Alabama | Early Campanian | - |
| ALMNH PV 1993.2.133 | Partial skeleton. Premaxilla, maxillae, mandible, 7 vertebrae, rib, coracoid, omal furcula fragment, distal ulna, ulnar carpals, carpometacarpus, manual phalanges, femur, tibiotarsus, tarsometatarsus & pedal phalanx | Mooreville Fm, Alabama | Early Campanian | 6,8,9 |
| BHI 6420 | Partial skeleton. Scapulae, coracoids, omal furcula fragment, humeri, radius, proximal ulna, radial carpal, carpometacarpus, manual phalanges | Niobrara Fm, Kansas | Middle Santonian | 4,5,8,9 |
| BHI 6421 | Partial skeleton. Mandibles, quadrate, 3 thoracic vertebrae, distal humerus, distal ulna, manual phalanx & distal tibiotarsus | Niobrara Fm, Kansas | Middle Santonian | 5,9 |
| FHSM VP 18702 | Largely complete skeleton. Partial skull, lower jaws, 2 cervical vertebrae, synsacrum, ribs, sternum, scapula, coracoids, partial humeri, ulna, radii, radial carpals, ulnar carpal, carpometacarpus, manual phalanx, femur, distal tibiotarsus, fibula, tarsometatarsus & pedal phalanges | Niobrara Fm, Kansas | Middle Santonian | 1,2,4,5,7,8,9 |
| KUVP 2281 | Scapula & coracoid | Niobrara Fm, Kansas | Middle Santonian | 4 |
| KUVP 2300 | Humerus | Niobrara Fm, Kansas | Middle Santonian | 5 |
| KUVP 5969 | Coracoid, humerus, ulna, radius, radial carpal, manual phalanx | Niobrara Fm, Kansas | Middle Santonian | Lacks 9 |
| KUVP 25469 | Synsacrum, omal furcula fragment, distal humerus, distal radius, carpometacarpus & manual phalanx | Niobrara Fm, Kansas | Middle Santonian | 7 |
| KUVP 25471 | Fragmentary vertebra & humerus | Niobrara Fm, Kansas | Middle Santonian | 5 |
| KUVP 25472 | 17 vertebrae (cervical, thoracic & caudal), radius & proximal ulna | Niobrara Fm, Kansas | Middle Santonian | 2,3 |
| KUVP 119673 | Partial skeleton. Quadrate, jugal, mandible, 13 vertebrae (cervical, thoracic & caudal), synsacrum, sternum, ribs, coracoid, scapula, humeri, radii, ulnae, pelves, femur, distal tibiotarsus & fibula | Niobrara Fm, Kansas | Middle Santonian | 1,2,3,4,5,6 |
| KUVP 123459 | Partial carpometacarpus & distal ulna | Niobrara Fm, Kansas | Middle Santonian | 8 |
| MSC 13214 | Partial tarsometatarsus | Mooreville Fm, Alabama | Early Campanian | - |
| MSC 13868 | Fragmentary pedal phalanx | Mooreville Fm, Alabama | Early Campanian | - |
| MSC 34426 | Distal carpometacarpus | Mooreville Fm, Alabama | Early Campanian | 8 |
| MSC 34427 | Partial coracoid, distal ulna & partial carpometacarpus | Mooreville Fm, Alabama | Early Campanian | 6,8 |
| NHMUK A 905 | Sternum, medial portion of furcula, coracoids, scapulae, humeri & proximal radius | Niobrara Fm, Kansas | Middle Santonian | 4,5 |
| RMM 2841 | Distal carpometacarpus | Mooreville Fm, Alabama | Early Campanian | 8 |
| RMM 3394 | Distal carpometacarpus | Mooreville Fm, Alabama | Early Campanian | 8 |
| RMM 5794 | Proximal humerus | Mooreville Fm, Alabama | Early Campanian | - |
| RMM 5895 | Distal humerus, proximal radius & manual phalanx | Mooreville Fm, Alabama | Early Campanian | 9 |
| RMM 5916 | Proximal ulna | Mooreville Fm, Alabama | Early Campanian | - |
| RMM 5937 | Partial coracoid | Mooreville Fm, Alabama | Early Campanian | - |
| RMM 6200 | Proximal radius & distal ulna | Mooreville Fm, Alabama | Early Campanian | 6 |
| RMM 6201 | Manual phalanx | Mooreville Fm, Alabama | Early Campanian | 9 |
| RMM 6202 | Carpometacarpus | Mooreville Fm, Alabama | Early Campanian | 8 |
| RMM 7841 | Omal coracoid, humeri & distal ulna | Mooreville Fm, Alabama | Early Campanian | 5,6 |
| RMM 7842 | Partial humerus | Mooreville Fm, Alabama | Early Campanian | - |
| RMM 7844 | Distal humerus | Mooreville Fm, Alabama | Early Campanian | - |
| YPM 1450 | Partial skeleton. Partial skull, mandibles, 4 vertebrae, synsacrum, partial sternum, ribs, coracoid, humerus, ulna, radius, carpometacarpus, femur & tibiotarsus. This study includes only scans of its sternum & ulna | Niobrara Fm, Kansas | Middle Santonian | Holotype. 2,5,6,7,8 |
| YPM 1461 | Sternum, ribs, coracoid & humerus. This study includes only scans of its sternum. | Niobrara Fm, Kansas | Middle Santonian | - |
| YPM 1724 | Carpometacarpus | Niobrara Fm, Kansas | Middle Santonian | 8 |
| YPM 1733 | Partial skeleton. 10 vertebrae, synsacrum, coracoid, scapula, humerus, radius & partial ilium. This study includes only scans of its coracoid. | Niobrara Fm, Kansas | Middle Santonian | 2,4 |
| YPM 1740 | Ulna | Niobrara Fm, Kansas | Middle Santonian | 6 |
| YPM 1741 | Coracoid, humerus & radius. This study includes only scans of its radius | Niobrara Fm, Kansas | Middle Santonian | 7 |

## MATERIAL AND METHODS

### SPECIMENS STUDIED

The present work is based on the study of a series of previously undescribed postcranial specimens referred to *Ichthyornis*, with the cranial material of several of these specimens previously described in Field et al. (2018a). The most complete specimens included in the study are FHSM VP-18702 (a partial skeleton from a single fossil block, including cranial material, the pectoral girdle and forelimbs, and the synsacrum and hindlimbs); KUVP 119673 (a partial skeleton from a single block filled with radiopaque inclusions, preserving cranial material, most cervical and anterior thoracic vertebrae, the pectoral girdle, a partial forelimb, the pelvic girdle and partial hindlimb); ALMNH PV93.2.133 (a partial skeleton including cranial material, thoracic, sacral, and caudal vertebrae, a partial forelimb and complete hindlimb); and BHI 6420 (preserving the complete pectoral girdle and forelimb). Thirty-six additional undescribed specimens that are less complete are also described here for the first time—see Table 1 for the complete list of material. In addition to this substantial amount of new material, several skeletal elements from YPM specimens previously described by Marsh (1880) and Clarke (2004) have been incorporated into this study: the *Ichthyornis dispar* holotype YPM 1450 (ulna), 1724 (carpometacarpus), 1733 (coracoid), 1740 (ulna) and 1741 (radius).

All specimens studied come from middle to late Santonian rocks of the Niobrara Formation in Kansas (US), and from the early Campanian deposits of the Mooreville Chalk in Alabama (US), the same localities that produced the majority of the classic *Ichthyornis* material (Marsh, 1880; Clarke, 2004; Field et al., 2018a). See Table 1 and the Supplemental Information for available information on the provenance of each specimen.

The newly described specimens were referred to *Ichthyornis dispar* based on the presence of multiple autapomorphies, previously described by Clarke (2004). Where no diagnostic features were preserved, specimens were referred to *Ichthyornis* based on their morphological similarity to specimens preserving autapomorphies. See Table 1 for a full list of the diagnostic features preserved in each specimen.

## METHODS

### CT-Scanning

The specimens were scanned at the Cambridge Biotomography Centre, the University of Texas High-Resolution CT Facility (UTCT), and the Center for Nanoscale Systems at Harvard. Scan parameters and details for each specimen are provided in the Supplemental Information. The scans were assembled and digitally segmented using VG Studio Max 3.3 (Volume Graphics, Heidelberg, Germany), from which 3D surface meshes of individual elements were extracted and exported. Skeletal models of each specimen in anatomical connection were built in Autodesk Maya 2020.

### Anatomical Comparisons

The main references for comparative morphological information on *Ichthyornis* were descriptions of the classic *Ichthyornis* specimens from the YPM collections by Marsh (1880) and Clarke (2004). Comparisons with other fossil euornithean taxa were based on available literature (e.g., Zhou et al., 2014; Wang et al., 2016, 2020b). Comparisons with extant taxa were based on specimens from the University of Cambridge Museum of Zoology (UMZC). Osteological and myological nomenclature follows that of Baumel & Witmer (1993) and Baumel & Raikow (1993), with additional nomenclature from Livezey & Zusi (2007) and Mayr (2014, 2016). We acknowledge the complicated developmental identities of the free carpal bones (Botelho et al., 2014), and how these may be at odds with the traditional usage of the terms ulnare and radiale. For clarity, in light of Botelho et al. (2014), we have decided to refer to these elements as the ulnar carpal and radial carpal, respectively. We use standard terminology for morphological orientation (medial/lateral, dorsal/ventral, etc.), and preferentially apply the terms cranial/caudal instead of anterior/posterior, except for cases where disambiguation is required (e.g., anterior caudal vertebrae).

### Phylogenetic Analyses

We tested the phylogenetic position of *Ichthyornis* by re-scoring it in updated versions of the morphological matrices from Huang et al. (2016) and Wang et al. (2020b). Taxa were also added and re-scored from Wang & Zhou (2020), Hu et al. (2020), Wang et al. (2020c) and O’Connor et al. (2020). With the exception of O’Connor et al. (2020), these studies did not incorporate the additional 5 characters and character re-scorings from Field et al. (2018a). We produced a new dataset by combining the matrices from Wang et al. (2020b), including updates from Field et al. (2018a), with the matrix from O’Connor et al. (2020). The Huang et al. (2016) matrix was found to contain multiple scoring errors, affecting at least 6 characters, including several characters scored for more states than described. Several of these problems were already present in a previous version of that dataset from Liu et al. (2014), but not in Clarke (2004) or Clarke et al. (2006). These issues were corrected for the present study, but an exhaustive overhaul of that dataset would be beyond the scope of this study. Therefore, we recommend caution in future investigations employing the Huang et al. (2016) morphological matrix. Some taxa were re-scored based on published literature, such as *Gansus* (Wang et al., 2016b) and *Iteravis* (Zhou et al., 2014). A complete list of scoring changes and corrections to published matrices is provided in the Supplemental Information.

Given the presence of several morphological differences among the specimens described here (see morphological descriptions), well-represented specimens were initially included as distinct operational taxonomic units (OTUs) in our phylogenetic analyses. In all cases, these were recovered either within an exclusive clade including the *Ichthyornis* holotype, or in a polytomy comprising the holotype of *Ichthyornis* and the clade formed by Hesperornithes + crown birds (see supplementary trees in the Supplemental Information).

Based on these results, all specimens were treated as a single, combined OTU for *Ichthyornis dispar* in subsequent analyses.

We performed phylogenetic analyses under both parsimony and Bayesian analytical frameworks in order to account for differences introduced by alternative optimality criteria. As found by Field et al. (2018a), *Apsaravis* was identified as a wildcard taxon, and alternative analyses were performed including and excluding it. Its removal yielded better-resolved relationships and higher node support values within Euornithes. Parsimony analyses were conducted using TNT 1.5 (Goloboff & Catalano, 2016, made available with the sponsorship of the Willi Hennig Society). An unconstrained heuristic search with equally weighted characters was performed, with 1,000 replicates of random stepwise addition using the tree bisection reconnection (TBR) algorithm. 10 trees were saved per replicate, and all most parsimonious trees (MPTs) were used to calculate a strict consensus. Bremer support values were calculated in TNT using TBR from existing trees. Bootstrap analyses were performed using a traditional search and 1,000 replicates, with outputs saved as absolute frequencies. Our Bayesian analyses followed the same protocol as Field et al. (2020a). We conducted Bayesian analyses with MrBayes (Ronquist et al., 2012) using the CIPRES Science Gateway (Miller et al. 2010), and data were analysed under the Mkv model (Lewis, 2001). Gamma-distributed rate variation was assumed in order to allow for variation in evolutionary rates across different characters. Analyses were conducted using four chains and two independent runs, with a tree sampled every 4,000 generations and a burn-in of 25%. Analyses were run for 30,000,000 generations, and analytical convergence was assessed using standard diagnostics provided in MrBayes (average standard deviation of split frequencies < 0.02, potential scale reduction factors = 1, effective sample sizes > 200). Results obtained from independent runs of the same analyses were summarized using the sump and sumt commands in MrBayes. Morphological synapomorphies of recovered tree topologies were optimized under parsimony, by exporting the recovered trees into TNT.

## INSTITUTIONAL ABBREVIATIONS

The following abbreviations denote the museum collections where the specimens mentioned in this article are accessioned. **ALMNH**, Vertebrate Paleontology Collection, Alabama Museum of Natural History, University of Alabama, Tuscaloosa, AL, USA; **BHI**, Black Hills Institute of Geological Research, Hill City, SD, USA; **FHSM**, Sternberg Museum of Natural History, Fort Hays State University, Hays, KS, USA; **KUVP**, Vertebrate Paleontology Division, University of Kansas Biodiversity Institute & Natural History

Museum, Lawrence, KS, USA; **NHMUK**, National History Museum, London, UK; **MSC & RMM**, McWane Science Center (formerly Red Mountain Museum), Birmingham, AL, USA; **UMZC**, University of Cambridge Museum of Zoology, Cambridge, UK; **YPM**, Yale Peabody Museum, Yale University, New Haven, NY, USA.

## CLADE DEFINITIONS

To facilitate consistent phylogenetic nomenclature in the present and in future work on crownward stem birds, we establish definitions for the following clade names in accordance with rules outlined by the *International Code of Phylogenetic Nomenclature* (*PhyloCode*) (de Queiroz and Cantino, 2020). All names and definitions have been registered in the online database RegNum (Cellinese and Dell, 2020).

[NOMENCLATURAL ACTS REDACTED]

## SYSTEMATIC PALAEONTOLOGY

Avialae Gauthier, 1986

Ornithurae, Haeckel, 1866

Ichthyornithes Marsh, 1873b *sensu* Clarke, 2004

*Ichthyornis dispar* Marsh, 1872b

### Holotype

YPM 1450, a partial skeleton consisting of portions of the skull, mandible, most of the axial elements, pectoral girdle, wings and hindlimbs. The specimen was illustrated and described most recently by Clarke (2004), with additional elements identified and described by Field et al. (2018a).

### Locality and horizon

YPM 1450 was collected from sediments of the Smoky Hill Chalk Member, Niobrara Formation, near the Solomon River in Section 1, Township 6, Range 19, in Rooks County (Marsh, 1880; Brodkorb, 1967; Clarke, 2004).

### Referred specimens in this study

ALMNH PV 1988.20.180, 1988.20.181, 1988.20.183.1, 1988.20.186, 1988.20.375.3, 1988.20.427.1, 1988.20.5, 1988.26.1, 1993.2.191.1, 1993.2.133; BHI 6420, 6421; FHSM VP-18702; KUVP 2281, 2300, 5969, 25469, 25471, 25472, 119673, 123459, MSC 13214, 13868, 34426, 34427; NHMUK A 905; RMM 2841, 3394, 5794, 5895, 5916, 5937, 6200, 6201, 6002, 7841, 7842, 7844; YPM 1461, 1724, 1733, 1740, 1741. See Table 1 for summaries of the material associated with each specimen.

### Diagnosis

Following Clarke (2004), *Ichthyornis dispar* shows the following autapomorphies: a single large pneumatic foramen located on the anteromedial surface of the corpus of the quadrate, amphicoelous or biconcave cervical vertebrae, anterior free caudal vertebrae with well-developed prezygapophyses clasping the dorsal surface of the preceding vertebra, an extremely diminutive acromion process of the scapula, a pit-shaped fossa for muscle attachment at the distal end of the humeral bicipital crest, the length of the trochlear surface along the posterior surface of the distal ulna approximately equal to the width of the trochlear surface, an oval scar on the posteroventral surface of the distal radius, a large turbercle close to the articular surface for the phalanx II:1 in the carpometacarpus, and the presence of an internal index process in the phalanx II:1 (Clarke, 2004).

## ANATOMICAL DESCRIPTION

### VERTEBRAL COLUMN

Seven of the newly described specimens preserve some vertebral material, with three of them—KUVP 15472, KUVP 119673, and ALMNH PV93.2.133—preserving a significant portion of the axial series (Fig. 4,5 & 6), although no complete vertebral column is yet known for *Ichthyornis*. BHI 6421 preserves three isolated thoracic vertebrae, with two of them being extremely fragmentary. FHSM VP-18702 preserves two complete but severely distorted cervical vertebrae, as well as a complete but poorly preserved synsacrum. KUVP 25469 preserves a complete synsacrum, and KUVP 2471 preserves only a single very fragmentary thoracic vertebra.

**Figure 4.**
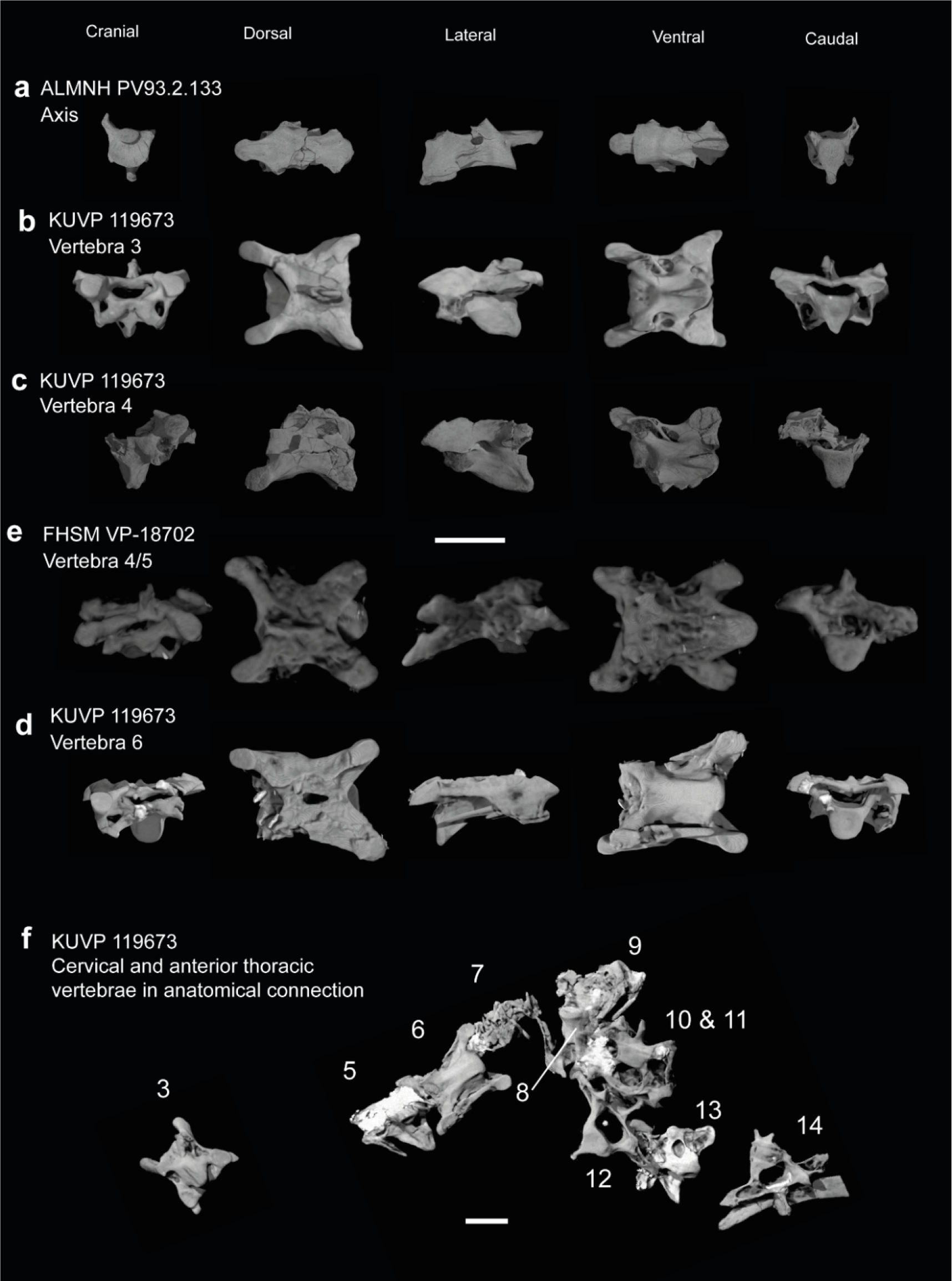
Anterior cervical vertebrae of *Ichthyornis*. (a) ALMNH PV93.2.133 axis, (b) KUVP 119673 possible 3^rd^ cervical, (c) KUVP 119673 possible 4^th^ cervical, (d) KUVP 119673 possible 5^th^ cervical, (e) FHSM VP-18702 possible 4^th^ or 5^th^ cervical; in cranial, dorsal, lateral, ventral, and caudal views. (f) KUVP 119673, cervical and anterior thoracic vertebrae in anatomical connection as preserved in the specimen, with probable vertebral numbers indicated. Scale bar equals 5 mm.

**Figure 5.**
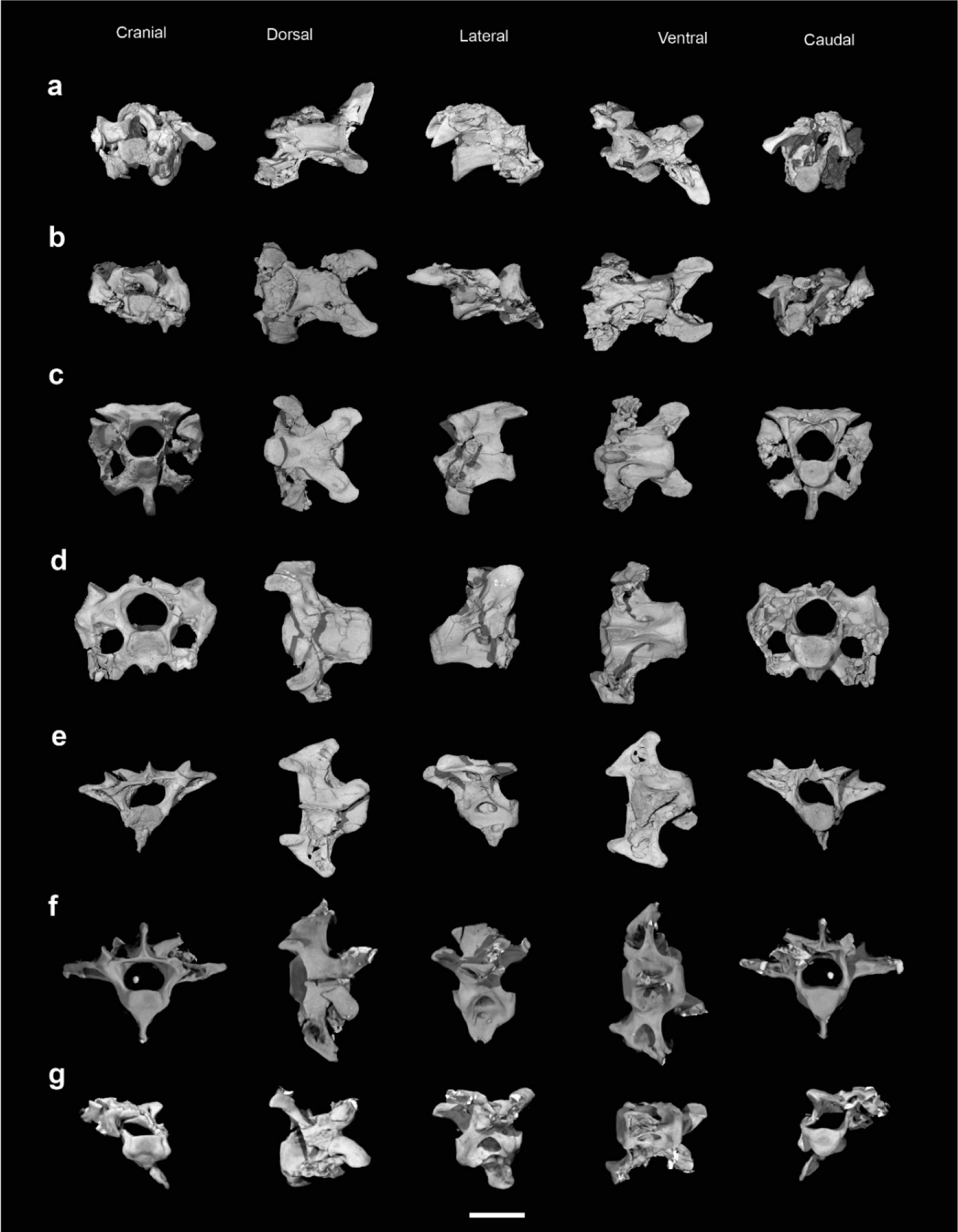
Mid and posterior cervical and anterior thoracic vertebrae of *Ichthyornis* specimen KUVP 25472. (a) 6^th^ or 7^th^ cervical vertebra, (b) 7^th^ or 8^th^ cervical vertebra, (c) 9^th^ or 10^th^ cervical vertebra, (d) 10^th^ or 11^th^ cervical vertebra, (e) 1^st^ thoracic vertebra, (f) 2^nd^ thoracic vertebra, (g) 3^rd^ thoracic vertebra; in cranial, dorsal, lateral, ventral, and caudal views. Scale bar equals 5 mm.

KUVP 25472 is the specimen that preserves the highest presacral vertebral count, with five cervical (Fig. 5 & 6) and nine thoracic vertebrae (Fig. 6), most of them in an exceptional state of preservation, as well as three caudal vertebrae (Fig.8). Of these, only two posterior thoracic and two of the caudal vertebrae are articulated. KUVP 119673 preserves the highest total vertebral count of any *Ichthyornis* specimen known to date, with eight cervical vertebrae, three anterior thoracic vertebrae, a complete synsacrum, and three caudal vertebrae. KUVP 119673 is remarkable as well for being the only specimen known to preserve a significant portion of its vertebral column in articulation, with at least seven of the cervical vertebrae and the anterior thoracic vertebrae found in anatomical connection, although five of the posterior cervical vertebrae are badly distorted and crushed against one-another (Fig. 4). ALMNH PV93.2.133 preserves the axis, two fragmentary posterior cervical vertebrae (Fig. 4), two anterior and two posterior thoracic vertebrae and a partial synsacrum.

**Figure 6.**
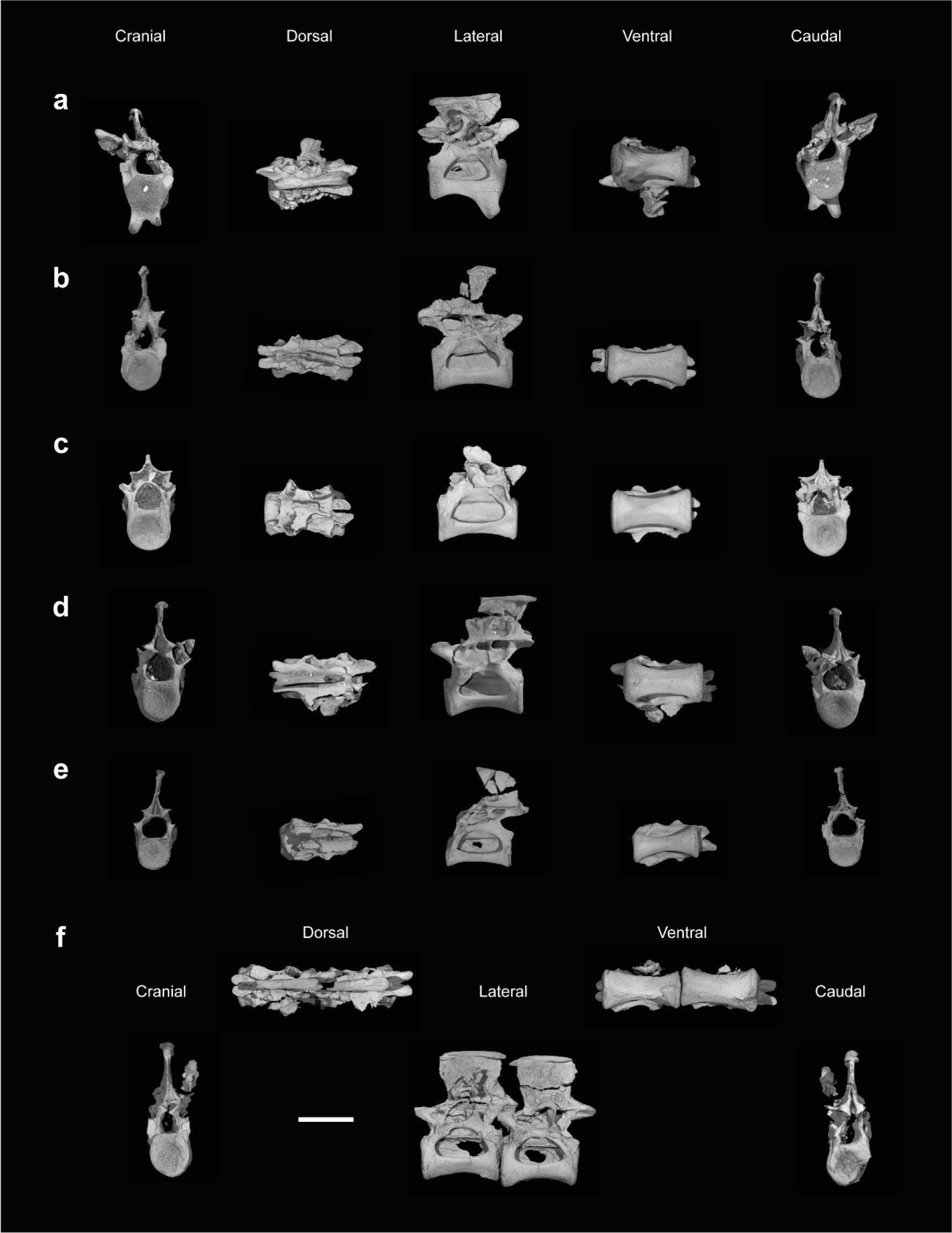
Anterior and mid-thoracic thoracic vertebrae of *Ichthyornis* specimen KUVP 25472. (a) 3^rd^ thoracic vertebra, (b) indeterminate mid thoracic vertebra 1, (c) indeterminate mid thoracic vertebra 2, (d) indeterminate mid thoracic vertebra 3, (e) indeterminate posterior thoracic vertebra 1, (f) 2 mid-thoracic vertebrae in anatomical connection; in cranial, dorsal, lateral, ventral, and caudal views. Scale bar equals 5 mm.

The wealth of axial material preserved among the new specimens included in this study contrasts with the limited number of vertebrae previously described for *Ichthyornis*, although both Marsh (1880) and Clarke (2004) extensively described and illustrated the limited YPM axial material. Of the previously described specimens, only four, including the holotype (YPM 1450), preserve axial material (Marsh 1880, Clarke, 2004). YPM 1733 exhibits the highest vertebral count, with four cervical vertebrae and six thoracic vertebrae, as well as a complete synsacrum. In total, the YPM material includes 19 presacral and possibly 6 caudal vertebrae, while the new specimens preserve 38 presacral vertebrae, four partial or complete synsacra and six caudal vertebrae.

In spite of the considerable amount of newly preserved vertebral material, it is currently impossible to establish a precise total vertebral count for *Ichthyornis*, although the new specimens offer a much better opportunity for estimating this figure and drawing comparisons with other Mesozoic avialans. The vertebrae preserved in KUVP 25472 and 119673 indicate a minimum number of 21 presacral vertebrae, with at least eleven cervical vertebrae (including the axis and atlas) and ten thoracic vertebrae. This estimate is the same as that of Marsh (1880), who based his estimate on the axial skeleton of the extant *Sterna maxima,* although he considered the incompletely fused first sacral vertebra of *Ichthyornis* (YPM 1732) the last thoracic vertebra (Clarke, 2004). Although it is not possible to verify this vertebral count without additional, more complete specimens, similar presacral counts have been described for the few Mesozoic euornitheans preserving sufficiently complete vertebral columns, such as *Yixianornis* with 22 presacral vertebrae (12 cervical and 10 thoracic; Clarke et al., 2006), and the hesperornithines *Hesperornis* and *Parahesperornis,* both with 23 presacral vertebrae, although in Hesperornithes the relative count of cervical vertebrae is notably higher (17 cervical and 6 thoracic, Marsh, 1880; Bell & Chiappe, 2020).

### SACRAL VERTEBRAE

The synsacrum is represented in four of the studied specimens, of which three preserve virtually the entire element: FHSM VP-18702, KUVP 119673, and KUVP 25469. The synsacrum of ALMNH PV93.2.133 is divided into two matching fragments but appears to be missing the cranialmost sacral vertebrae. The preservation of FHSM VP-18702 is fairly poor, preventing the discernment of several salient morphological details, such as the morphology of most transverse processes and the clear delineation of several individual vertebrae. Despite the breakage of the transverse processes and several missing parts, both KUVP 25469 and ALMNH PV93.2.133 show very little distortion, and the sacral vertebrae are easily distinguishable. KUVP 119673 is dorsoventrally flattened, and many of its morphological features are obscured by radiopaque inclusions. Nonetheless, this specimen shows minimal breakage and preserves most of the vertebral transverse processes. Synsacral measurements are provided in Table 2.

**Table 2.** Measurements of the synsacrum of *Ichthyornis* specimens. Maximum length corresponds to the total craniocaudal length of the ankylosed or fused vertebrae, not including disassociated sacral vertebrae. Acetabulum width is measured from the total width of the complete acetabulum transverse processes. Maximum width and height are provided for the cranialmost and caudalmost fused sacral vertebrae. Asterisks (*) denote measurements that might be unreliable due to breakage or distortion, but are included for completeness. All measurements are in mm. - = not measurable.

| Specimen | Max. length | Acetabulum width | Cranial centrum width | Cranial centrum height | Caudal centrum width | Caudal centrum height |
| --- | --- | --- | --- | --- | --- | --- |
| ALMNH PV 1993.2.133 | - | - | - | - | 2.78 | 1.42 |
| FHSM VP 18702 | 44.30 | - | 4.08 | 1.94* | 2.70 | 1.50* |
| KUVP 25469 | 43.01 | - | 4.03 | 4.70* | 2.59 | 2.43* |
| KUVP 119673 | 31.64 | 17.12 | 4.06 | 2.61 | 2.32 | - |

The number of fused sacral vertebrae varies among the studied specimens; the same was reported by Clarke (2004) in reference to the two complete synsacra in the YPM collection. YPM 1450 preserves ten vertebrae, but YPM 1732, a larger specimen, preserves twelve. They also differ in terms of which vertebra bears the perpendicular costal processes attaching to the acetabular area: this was reported to be the seventh vertebra in YPM 1450, and the ninth in YPM 1732 (Clarke, 2004). Both FHSM VP-18702 and KUVP 25469, the largest specimens investigated here, include twelve ankylosed vertebrae (Fig. 7a,c), although the poor preservation of FHSM VP-18702 makes it difficult to differentiate individual vertebrae. The first sacral of FHSM VP-18702, much larger than the second, is poorly ankylosed, connected only by ossified tendons dorsally, with a noticeable gap between its centrum and that of the second sacral. The same vertebra is fully fused in KUVP 25469, and no suture is visible (Fig. 8c). An intermediate condition seems to be present in YPM 1732, in which the first sacral is fully ankylosed to the second but a suture is clearly visible (Clarke, 2004). The acetabular bar is present in both specimens on the ninth vertebra. KUVP 119673, the smallest synsacrum, preserves only ten fused sacral vertebrae, with the acetabular bar on the eighth vertebra (Fig. 7b). Interestingly, a large, isolated vertebra with a dorsoventrally compressed centrum is preserved for KUVP 119673, which, given the size of its caudal articular surface, probably sits just cranial to the first fused sacral. The morphology and proportions of this vertebra match those of the first sacral of FHSM VP-18702 and KUVP 25469. Thus, this element probably represents the first sacral of KUVP 119673, even though it is unfused. From this point onwards, this element will be referred as the first sacral of this specimen. KUVP 119673 also differs from all other specimens in the number of postacetabular fused sacrals, exhibiting two distinguishable vertebrae instead of three. ALMNH PV93.2.133 is incomplete, preserving ten vertebrae; however, given its size (which is comparable to that of KUVP 25469) and the presence of three postacetabular vertebrae, it is probably missing the two cranialmost sacrals (Fig. 7d).

**Figure 7.**
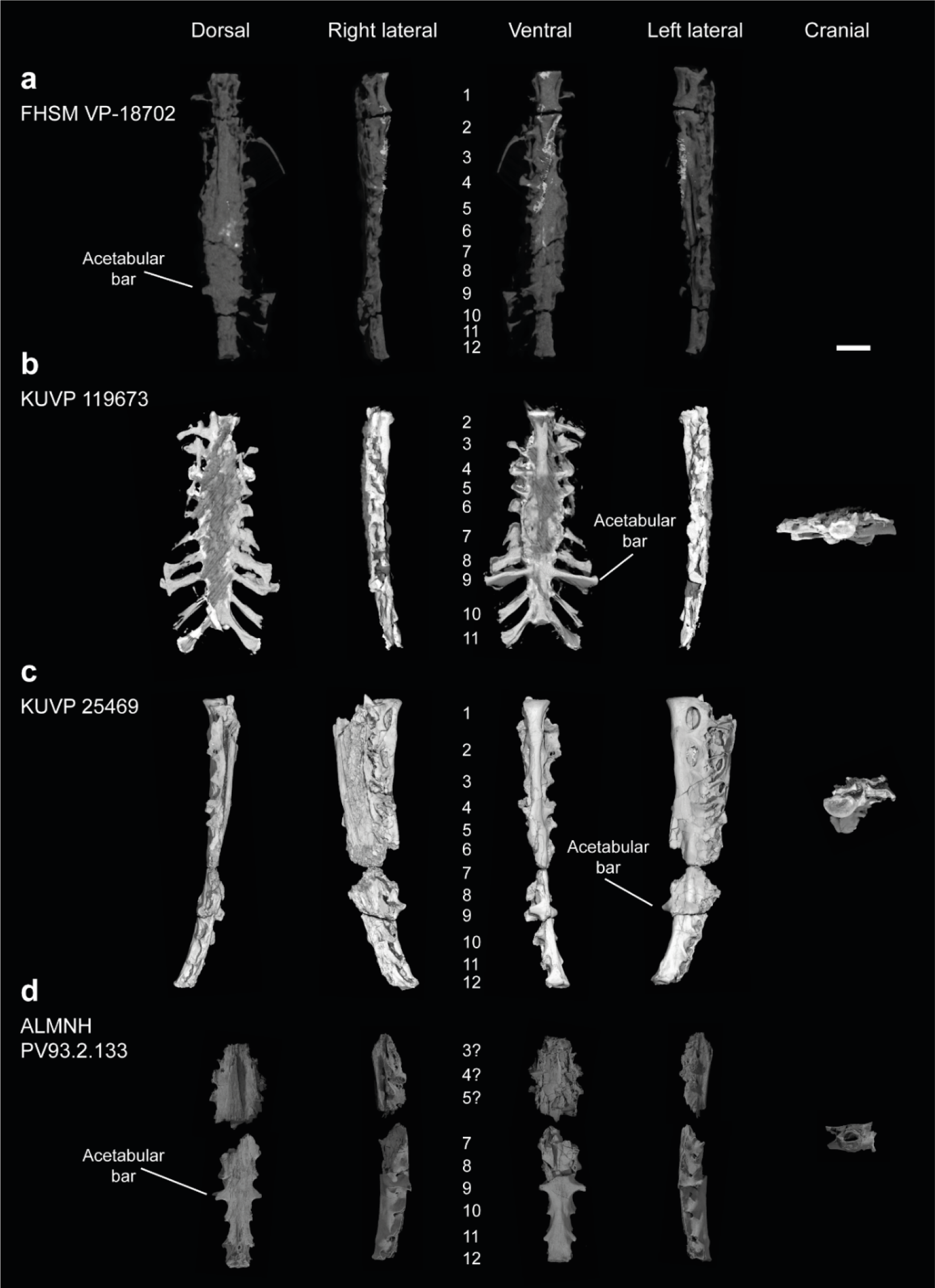
Synsacra of *Ichthyornis*. (a) FHSM VP-18702, (b) KUVP 119673 (c) KUVP 25469 and (d) ALMNH PV93.2.133 in dorsal, right lateral, ventral, left lateral and cranial views. Numbers indicate sacral vertebral order. The 1^st^ sacral vertebra in FHSM VP-18702 is incompletely fused, and the 1^st^ and 11^th^ sacral vertebrae in KUVP 119673 are unfused and not pictured here. Scale bar equals 1 cm.

**Figure 8.**
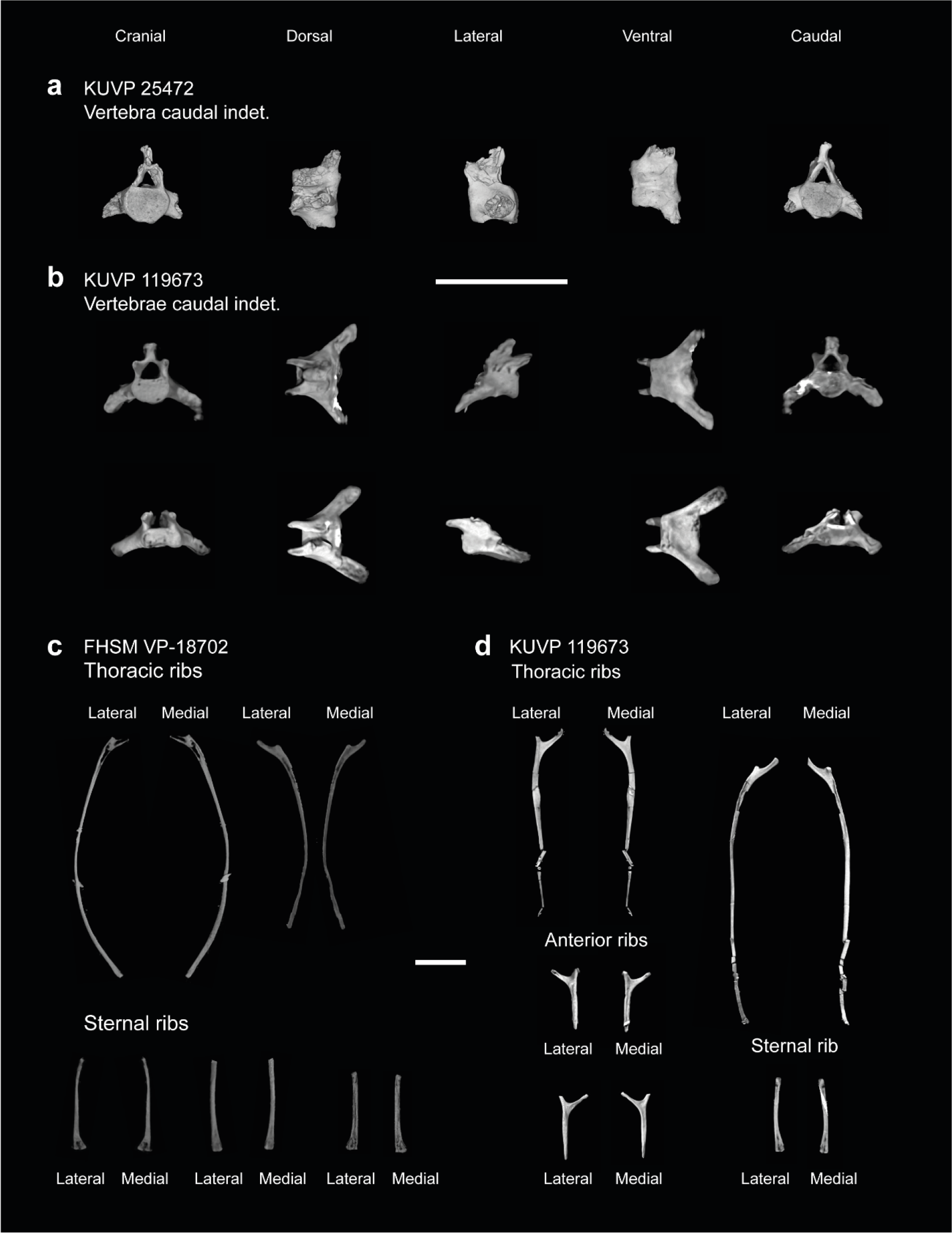
Caudal vertebrae and ribs of *Ichthyornis*. (a) KUVP 25472, caudal vertebra indeterminate, (b) KUVP 119673, caudal vertebrae indeterminate; in cranial, dorsal, lateral, ventral and caudal views; (c) FHSM-VP 18702, thoracic and sternal ribs and (d) KUVP 119673 anterior, thoracic, and sternal ribs, in lateral and medial views. Scale bar equals 1 cm.

The new specimens, therefore, bridge a gap between the YPM synsacra, revealing a pattern of vertebral fusion in which an additional sacral is added at both the cranial and caudal ends of the synsacrum. Given that the larger specimens are those with twelve fused vertebrae (Table 2), as well as the absence of a suture between the first and the second sacrals in KUVP 25469, we consider it likely that the differences among these specimens and the YPM synsacra are ontogenetic in nature. The largest specimens, KUVP 25469 and FHSM VP-18702, are respectively only 16.2 and 16.7% longer than KUVP 119673 (including the unfused first sacral and excluding the presumably missing last unfused postacetabular sacral), suggesting a late ontogenetic acquisition of complete bone fusion, as all specimens probably represent adults or subadults. A reduced number of fused sacral vertebrae in hatchlings and juveniles has been reported in Enantiornithes, with several immature specimens preserving variably five, six, or eight fused sacrals, with eight sacrals being the normal condition amongst adults (Chiappe et al., 2007; Knoll et al., 2018). Developmental variability in the number of fused sacrals is less well-known in stemward euornitheans, and probable ontogenetic variation in sacral fusion has only been described in *Archaeorhynchus*, where early-stage juveniles exhibit completely unfused sacrals (Foth et al., 2021), with at least four of the seven sacrals fused in subadults (Zhou et al., 2013) and a maximum of seven completely fused in fully-grown adults (Zhou & Zhang, 2006). The total number of fused sacrals varies among Mesozoic euornitheans, with seven in *Archaeorhynchus*, eight in *Zhongjianornis* (Zhou et al., 2010) and *Abitusavis* (Wang et al., 2020), nine in *Schizooura* (Zhou et al., 2012), *Mengciusornis* (Wang et al., 2020a), *Patagopteryx* (Chiappe, 2002), *Similiyanornis*, *Yanornis* and *Yixianornis* (Zhong & Zhou, 2001; Clarke et al., 2006; Wang et al., 2020), ten in *Apsaravis* (Clarke & Norell, 2002) and *Gansus* (You et al., 2006; Wang et al., 2016b), eleven in *Chaoyangia* (O’Connor & Zhou, 2013) and *Changmaornis* (Wang et al., 2013), twelve in *Juehuaornis* (Wang et al., 2015), and, although unclear, between ten and fourteen in Hesperornithes (Bell & Chiappe, 2016, 2020). This evidence suggests the possibility of a general crownward trend towards an increase in the number of fused sacrals across euornithean phylogeny. Although variation in sacral number and degree of sacral fusion is poorly characterized among crown birds, a greater degree of fusion in the posterior sacral vertebrae in more mature or older specimens has been reported in Anatidae (Woolfenden, 1960). The total number of fused sacral vertebrae shows a moderate degree of intraspecific variation among Anatidae (Verheyen, 1955; Woolfenden, 1960), Cuculiformes (Berger, 1956), and Gruiformes (Hiraga, 2013), with variation in total fused sacral count of up to two vertebrae documented. Although variation in the total number of documented sacral vertebrae has been suggested to be related differing counting approaches, this variation is considered to be genuine (Berger, 1956).

However, the relationship between total number of fused vertebrae in the synsacrum and ontogenetic stage is poorly studied and deserves research attention.

The cranial articular surfaces of the first sacral in KUVP 25469 and FHSM VP-18702 are taphonomically compressed dorsoventrally (Fig. 7a,c). The unfused first sacral of KUVP 119673 reveals that the articular surface is dorsoventrally shortened with respect to the thoracic vertebrae, but is similarly moderately concave. The caudal articular surface is broken and obscured by radiopaque inclusions in KUVP 119673 and barely appreciable in FHSM VP-18702, but seems to be smaller than the cranial articular surface, matching with the substantial size and dorsoventral height reduction between the first and second sacrals in KUVP 25469. The vertebra bears small but elongate prezygapophyses, which are triangular in dorsal view. A small gap is present between the first two sacrals in FHSM VP-18702, and a suture is visible between them in YPM 1732, but no equivalent suture is visible in KUVP 25469, in which only a moderate ventrolateral tubercle or ossification is present. Only the first two sacral vertebrae in KUVP 25469 and FHSM VP-18702 show large and deeply excavated lateral pleurocoels, which are probably non-pneumatic (O’Connor, 2006; Mayr, 2021), similar to those of the thoracic vertebrae; in KUVP 119673 these are only observable in the first isolated sacral, as they are obscured by radiopaque inclusions caudally. Similar lateral excavations or pleurocoels in the anterior sacral vertebrae have only been rarely reported for Mesozoic euornitheans, although they occur in a possible ornithuran synsacrum from the Maastrichtian of Madagascar (O’Connor & Forster, 2010). The first fused sacral of KUVP 119673 bears minute prezygapophyses that barely extend beyond the cranial articular surface of the vertebra. These are presumably fused or ankylosed with the postzygapophyses of the first sacral in the other specimens. The third sacrals of KUVP 2546 and FHSM VP-18702 show lateral concave depressions, but these are not fenestrated nor deeply excavated, and therefore seem apneumatic. A similar condition is present in the first preserved sacral of ALMNH PV93.2.133, here assumed to represent the third sacral (Fig. 7c). Only the first sacral of each specimen preserves clear parapophyses or articular surfaces for the sacral ribs similar to those of the thoracic vertebrae; the equivalent positions in the second, third and fourth sacrals show a shallow concavity pierced by numerous minute foramina. The transverse processes of the first three vertebrae are thin and craniocaudally compressed in FHSM VP-18702 and KUVP 25469—in which they are fused to a portion of the iliac preacetabular wing, but in contrast, they are proportionally more robust in KUVP 119673, in which they extend into wide, caudolaterally directed wing-like structures similar to those preserved in several thoracic vertebrae. The fourth vertebra shows no lateral excavations, and its transverse processes are much more robust than those of the preceding vertebrae in all specimens. The transverse processes are dorsolaterally oriented and extend along the entire dorsoventral extent of the centrum. The ventral portion of each transverse process is short and wide, and meets the subtriangular, flange-like and laterally oriented dorsal portion, which is only preserved in KUVP 119673, forming a marked cranial cavity (Fig. 7b). The centra of these first four vertebrae are short and circular in cross-section, and, following the pronounced reduction in size between the first and second sacrals, become progressively narrower caudally. A moderate ventrolateral expansion marks the suture between the centra of each vertebra.

The following three sacral vertebrae are dorsoventrally flattened, with laterally expanded centra forming a plate-like surface in ventral view, from which it is difficult to distinguish individual vertebrae. Their transverse processes are only well preserved in KUVP 119673. The transverse processes of the fifth sacral are similar in morphology to those of the 4th sacral, but with a thinner and craniocaudally compressed ventral portion and a shorter dorsal flange. The transverse processes of the sixth and seventh sacrals show minimal dorsoventral extension, and flat dorsal and slightly concave ventral surfaces. These processes are short, subquadrangular and laterally oriented on the sixth sacral vertebra, but longer and caudolaterally oriented on the seventh, with a caudal subtriangular expansion. The eighth sacral shows massively laterally expanded and elongate transverse processes, which lack any kind of ventral expansion. These are flat on both their dorsal and ventral surfaces, and are caudolaterally oriented and rectangular in shape, with a moderate caudolateral expansion at their lateral end, which contacts the processes of the ninth sacral caudally. Both the seventh and eighth sacral vertebrae show minute but deep lateral perforations just caudal to their transverse processes in ALMNH PV93.2.133, distinct from the large pleurocoels present in the anterior sacral vertebrae of KUVP 25469 and FHSM VP-18702 and those from the thoracic vertebrae, but these are either not preserved or not distinguishable in any of the other specimens. The ninth sacral shows the largest and most laterally expanded transverse processes, which attach in the acetabular area and form robust acetabular bars. These are only completely preserved in KUVP 119673 (Fig. 7b), with the rest of the specimens preserving only the base of the processes. They show a dorsoventral expansion similar to those of the fourth and fifth sacrals, with the ventral portion of each process extending along the entire height of the centrum. The ventral portion is laterally oriented and perpendicular to the main axis of the synsacrum; it is craniocaudally compressed and widens slightly on its ventral margin. Laterally, it expands into a flat and rounded articular facet for the ilium. The dorsal portions of the processes are similar in shape to those of the eighth sacral vertebra, and, in contrast to the ventral portions, are caudolaterally directed. A thin continuous sheet of bone extends dorsoventrally between both portions of the process in KUVP 119673, but this sheet is fenestrated in ALMNH PV93.2.133. The extent of the fenestration varies between the right and left processes, with the fenestra constituting a small round hole in the middle of the sheet on the left process, and a significantly larger opening on the right process, to the extent that it appears to that the bone sheet is almost entirely absent. Similar to the previous two sacral vertebrae, ALMNH PV93.2.133 shows tiny but deep perforations in the ninth sacral, absent in the other specimens.

Caudal to the ninth sacral, the number of fused vertebrae varies among the specimens investigated here, with two vertebrae in KUVP 119673 and three in the rest (Fig. 7). The tenth sacral is craniocaudally elongated, similar to the ninth, but the eleventh and twelfth are craniocaudally compressed. Their centra are narrow, with a low ridge running craniocaudally along their ventral surface. The contact between the eleventh and twelfth sacral vertebrae shows two moderately developed tubercles on its ventral surface, situated on either side of the aforementioned ridge. The caudal articular surface of the twelfth sacral is dorsoventrally short and almost quadrangular in shape, with a slightly concave caudal surface. The bases of the postzygapophyses seem to be present in ALMNH PV93.2.133, although these are broken. As in the previous sacral vertebrae, the eleventh and the twelfth sacrals of ALMNH PV93.2.133 exhibit minute perforations on their lateral surfaces. Only KUVP 119673 preserves the complete transverse processes for the tenth and the eleventh vertebrae, although their bases are well-preserved in ALMNH PV93.2.133. The shapes of the transverse processes of the tenth and eleventh sacrals are similar to those of the ninth sacral; this is particularly true for the eleventh, which shows a great dorsoventral extension, in contrast to the tenth, which is dorsoventrally short. The processes are more caudally oriented than those of the preceding sacral vertebrae. The dorsal portions of the processes are flat and craniocaudally narrow; the distal end is broken in the tenth sacral vertebra, but a large, rounded expansion is visible on the distal end of the processes of the eleventh. Transverse processes are only preserved on the unfused twelfth sacral of KUVP 119673; in all of the other specimens the lateral surfaces are eroded.

The internal morphology of the *Ichthyornis* synsacrum is obscured by imperfect preservation of the specimens, which, as mentioned, show variable degrees of flattening, distortion, and/or presence of radiopaque inclusions. Despite significant dorsoventral crushing cranially, the fragmentary synsacrum of ALMNH PV93.2.133 shows an enlarged neural canal towards the middle region of the synsacrum, congruent with an enlarged spinal cord and similar to the condition in the probable Maastrichthian euornithean synsacrum reported by O’Connor & Forster (2010). Although the presence of circumferential lumbosacral canals related to the balance-maintenance system (Necker, 2006) is difficult to verify in any of the specimens, circumferential indentations on the dorsal interior surface of ALMNH PV93.2.133 are congruent with the presence of such canals, constituting the second apparent occurrence of a lumbosacral sensory system in stem euornitheans (O’Connor & Forster, 2011).

*Guildavis* and *Apatornis*, represented exclusively by sacral remains, were differentially diagnosed from *Ichthyornis* by Clarke (2004) on the basis of only a few sacral characters. Clarke (2004) differentiated *Guildavis* from *Ichthyornis* based on the presence of a parapophysis on the left side of the first sacral vertebra, which was missing in YPM 1450 and 1732. These are well preserved in both KUVP 25469 and KUVP 119673, and are apparent in FHSM VP-18702; therefore, their absence in the YPM specimens may be taphonomic.

*Apatornis* was diagnosed as distinct from *Ichthyornis* based on the different number of fused sacral vertebrae (at least eleven), the number of mid-sacrals bearing dorsally directed transverse processes (four), and the lack of ossified tendons on the dorsal surface of the synsacrum. All of these characters are exhibited by one or more of the new specimens. As discussed above, the total number of fused sacral vertebrae varies among the studied specimens, presumably as a result of ontogenetic change. Both FHSM VP-18702 and KUVP 15469 exhibit four mid-sacrals with dorsally directed transverse processes, while KUVP 119673 exhibits only three such vertebrae. Both KUVP 119673 and ALMNH PV93.2.133 lack clear ossified tendons on their dorsal surfaces, while these are difficult to verify in FHSM VP-19702.

All the studied synsacra derive from specimens diagnosable as *Ichthyornis* on the basis of multiple apomorphic features (Table 1), and therefore it appears that the supposed diagnostic features of both *Guildavis* and *Apatornis* may fall within the range of variation of *Ichthyornis*. A more detailed reassessment of the preserved material of *Guildavis* and *Apatornis* will be necessary to confidently assess their taxonomic validity.

### RIBS

FHSM VP-18702 and KUVP 119673 both preserve several isolated vertebral and sternal ribs spread around other skeletal elements (Fig. 8). NHMUK A 905 preserves a single rib in association with the sternum, but comparison with the sternal ribs preserved in the other two specimens, and the shape of the sternal costal facets, reveal that it does not represent an articulated sternal rib.

No vertebral rib seems to be entirely complete, though most of the identifiable ribs preserve the proximal region. The ribs exhibit a flat and broad lateral surface but become progressively cylindrical in cross-section distally. KUVP 119673 preserves two floating ribs; these are short and lack the flat lateral surface seen in the other vertebral ribs (Fig. 8). The sternal ribs are short and wide, with a flattened caudal surface and a broad sternal articulation facet. They vary in length, with the shorter ones exhibiting wider sternal ends, matching the anterior sternal costal facets. The broad sternal end is caudally depressed and perforated by many small foramina. An element from FHSM VP-18702 described by Field et al. (2018a) as the palatine is reinterpreted here as a rib fragment. Although flattened and distorted, the size, proportions, and general morphology of this element agree with those of the proximal region of the preserved ribs (Fig. 8). The palatine of *Ichthyornis* was illustrated by Torres et al. (2021), and is notably different from that reported by Field et al. (2018a).

No uncinate processes, fused or unfused, are found in any of the studied specimens, and they seem to be absent in all of the YPM specimens (Clarke, 2004). These elements are found in both *Hesperornis* and *Parahesperornis* (Marsh, 1880; Bell & Chiappe 2020) and in several closely related euornitheans, like *Gansus* (Wang et al. 2016) and *Iteravis* (Wang et al. 2018). While the absence of these elements could be preservational, the fact that several of the known *Ichthyornis* specimens preserve numerous costal remains without any recognizable uncinate processes perhaps points to their genuine absence, a lack of (or incomplete) ossification of these processes, or at least a lack of fusion between the ribs and uncinate processes even in mature individuals. Whereas uncinate processes are present in most extant birds, they are absent in screamers (Anseriformes: Anhimidae; Codd, 2010).

### STERNUM

Three of the studied specimens preserve sterna in varying levels of completeness (Fig. 9). FHSM VP-18702 preserves a nearly complete, three-dimensional sternum, with the left side virtually undistorted from its original shape. This is similar to the skull associated with that specimen, in which the left side, preserved downward in the sediment, appears virtually undistorted, whereas the right side is mostly crushed flat (Field et al. 2018a). KUVP 119673 preserves a partial, dorsoventrally flattened sternum missing most of its right side but preserving the complete caudal portion of the element. Despite the excellent general preservation of KUVP 119673, much of the cranial portion of the sternum is infilled with radiopaque inclusions, which complicates the observation of several morphological features. NHMUK A 905, previously figured by Clarke (2004), exhibits a partial sternum, including the left side of the element and much of the sternal keel. Additionally, we CT-scanned and studied the two previously reported *Ichthyornis* sterna, those of the YPM 1450 and YPM 1461, which are considerably more fragmentary, but preserve clear pneumatic foramina. The specimens described here reveal the complete morphology of the sternum for the first time, including the first information on the caudal portion of this element. Measurements of the sternum from the most complete specimens are provided in Table 3.

**Figure 9.**
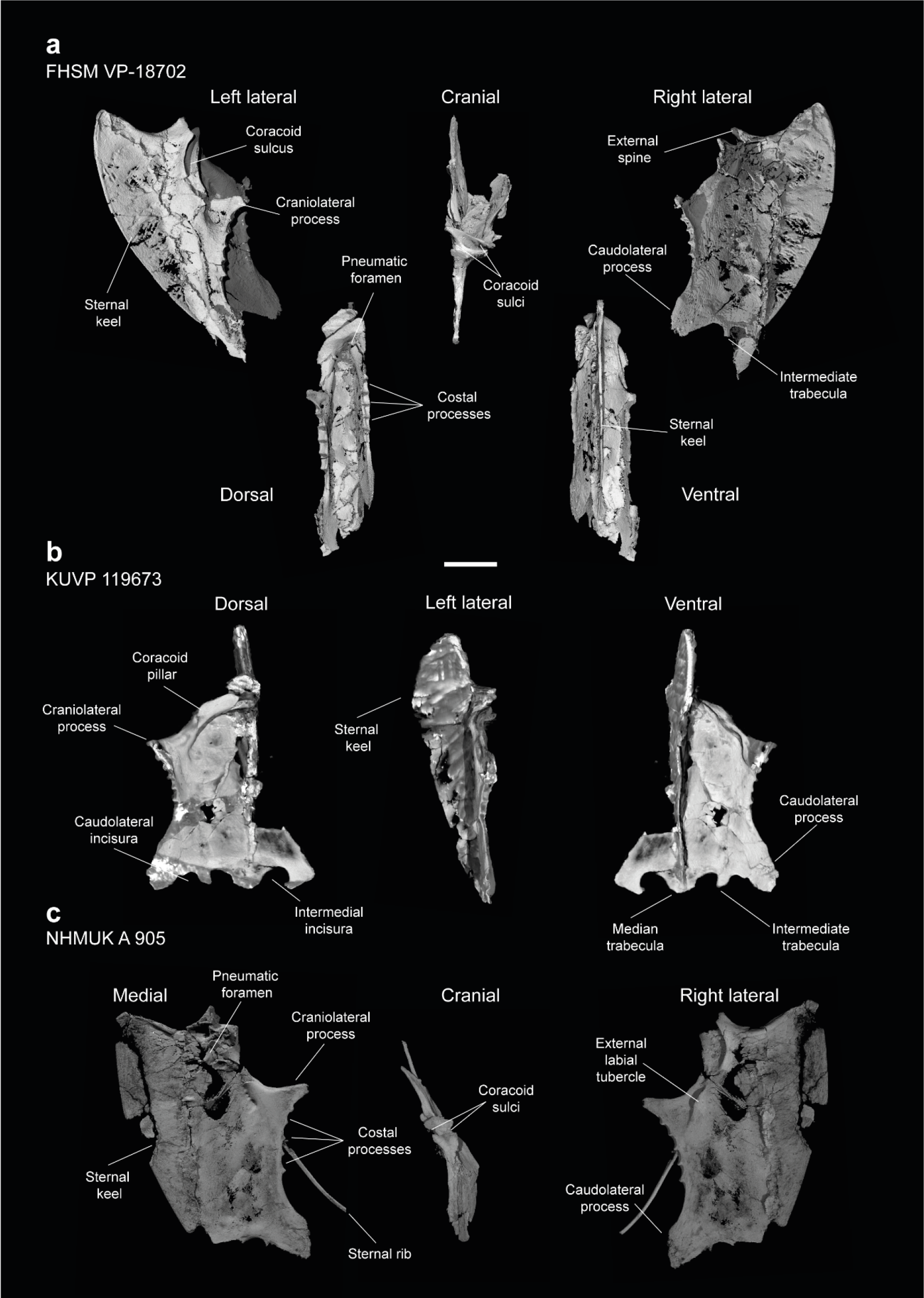
Sterna of *Ichthyornis*. (a) FHSM VP-18702 in left lateral, dorsal, cranial, ventral and right lateral views; (b) KUVP 119673 in dorsal, left lateral and ventral views and (c) NHMUK A 905 in medial, cranial and right lateral views. Scale bar equals 1 cm.

**Table 3.** Measurements of the sternum of *Ichthyornis* specimens. Total length measurements correspond to the maximum craniocaudal length of the sternum measured from the external spine to the caudal end of the median trabecula. Rostral width, craniolateral process width and caudal width are measured for one sternal side only (from the maximum extension of these structures to the sternal midline) since no specimen preserves both complete and undistorted sides of the sternum. Rostral width corresponds to the mediolateral extension of the sternal rostrum, measured from the maximum lateral extension of the coracoid pillar. Craniolateral process width corresponds to the maximum lateral extension of the craniolateral process. Caudal width is measured from the maximum lateral extension of the external trabeculae. Maximum keel depth is measured from the cranioproximal edge of the sternal keel to its maximum ventral extension. Asterisks (*) denote measurements that might be unreliable due to breakage or distortion, but are included for completeness. All measurements are in mm. - = not measurable.

| Specimen | Total length | Rostral width | Cranio-lat. processes width | Caudal width | Max. keel depth |
| --- | --- | --- | --- | --- | --- |
| FHSM VP 18702 | 45.38 | - | - | - | 13.90 |
| KUVP 119673 | 39.09* | 11.11 | 20.91 | 19.60 | 12.10* |
| NHMUK A 905 | 45.87* | - | 23.31 | 22.23 | 13.68 |
| YPM 1461 | - | - | - | - | 13.85 |

The rostral region of the sternum is well preserved in FHSM VP-18702 and NHMUK A 905, and as both Marsh (1880) and Clarke (2004) previously described, exhibits asymmetrical crossed coracoid sulci, in which the right coracoid sulcus lies ventral to the left, crossing at the midline (Fig. 9a). This condition is widespread among Neornithes, and is exhibited by numerous taxa such as Musophagidae, certain Gruidae, Phoenicopteridae, Ciconiidae, Threskiornithidae, Ardeidae, Scopidae, Aramidae, Eurypygidae, Phaethontidae, Procellariidae, Phalacrocoracidae, Balaenicipitidae, Accipitridae, Strigiformes, Falconinae and Psittaciformes, as well as the extinct total-clade anseriforms Presbyornithidae, the extinct total-clade palaeognaths Lithornithidae, and the extinct coliiform *Sandcoleus* (Houde, 1988; Houde & Olson, 1992; Ericson, 1997; Mayr & Clarke, 2003; Nesbitt & Clarke, 2016).

Amongst Mesozoic euornitheans, this condition is also present in *Iaceornis* (Clarke, 2004), and has been suggested to be present in *Yixianornis* (Clarke et al., 2006)*, Gansus*, and *Ambiortus,* though the condition is difficult to confirm in these taxa due to preservational issues (O’Connor & Zelenkov, 2013). This morphology constitutes an unusual example of midline asymmetry in a tetrapod, and the significance of this morphology and its functional implications are poorly understood. However, its wide phylogenetic distribution across crown birds as well as Mesozoic euornitheans suggests that the sternum of the last common ancestor of crown birds may have exhibited crossed coracoid sulci.

The coracoid sulci are deep and wide at their midlines, narrowing progressively towards both their medial and lateral margins, though they exhibit a limited dorsoventral flare on their caudal end in lateral view (Fig. 9a). The sulci extend caudolaterally from the midline in ventral view, turning parallel to the midline close to the base of the craniolateral processes. The dorsal edges of the coracoid sulci are flat along their anterior surfaces, and no internal spine of the sternum is present. A short but robust cranially-pointed external spine (spina externa; Baumel & Witmer, 1993) extends from the midline of the ventral edge of the sulci, which then extend laterally into two short processes pointing laterally and protruding cranially well beyond the dorsal edges of the sulci, giving this region a rhomboidal shape in ventral view (Fig. 9a). This observation departs from the morphological description of Clarke (2004), in which the cranial extent of these processes was assumed to be short. These processes, extending from the ventral lips of the coracoid sulci, coincide with the medial ends of both coracoid facets. Similar lateral processes of the ventral edges of the coracoid sulci are found in *Eurypyga helias,* which also shows crossed coracoid sulci, though the processes are more rounded and less marked than the condition in *Ichthyornis*. The ventral edges of the sulci turn caudolaterally just caudal to these processes, and then become completely caudally directed close to the base of the craniolateral processes, where they form a conspicuous and rounded external labial tubercle (tuberculum labri ext.; Baumel & Witmer, 1993). A low ridge extends ventrally from the external spine along the cranial surface of the sternal keel, which is wide and flattened.

Robust and wide coracoid pillars (pila coracoidea; Baumel & Witmer, 1993) extend caudolaterally from the dorsal region of the coracoid sulci as seen in KUVP 119673, forming an angle of roughly 100° between them, and delimiting a rounded internal depression (Fig. 9b). The dorsal region of the sternum (pars cardiaca; Baumel & Witmer, 1993) is well preserved in KUVP 119673, forming a broad and flat surface. The craniolateral processes (proc. craniolateralis; Baumel & Witmer, 1993) extend posteriorly from the coracoid pillars, pointing dorsolaterally and forming an angle of ∼ 120° with the cranial edge of the sternum. The craniolateral processes are subtriangular in shape, elongate, and gracile, exhibiting a shallow lateral depression extending from the coracoid sulci. Two rounded shallow depressions are visible along the length of the medial surface of the craniolateral processes.

A large pneumatic foramen is present on the dorsal region of the sternum, just caudal to the coracoid pillars. This foramen was previously reported by Clarke (2004) in YPM 1450 and YPM 1461, but our CT-scans reveal its complete morphology for the first time (Fig. 10). The foramen is subquadrangular, longer mediolaterally than craniocaudally and clearly delimited by thickened edges, which form almost straight angles on the vertices of the foramen. Both YPM 1450 and YPM 1461 are flattened, obscuring the internal structure of this pneumatic foramen. In contrast, in FHSM VP-18702 the external opening of the foramen is obscured by the flattening of the right side of the sternum, but it is observable in cross-section, where it opens into a large internal cavity. The right caudolateral edge of the foramen is preserved in NHMUK A 905, and the breakage of the rostral portion of the sternum allows the observation of the internal pneumatic cavity. KUVP 119673 preserves this region, but radiopaque inclusions prevent confirmation of the presence of the pneumatic foramen. A pneumatic foramen situated in the dorsal portion of the sternum is widespread and present in most major crown bird groups (Musser & Clarke, 2020), although it is absent in certain taxa, particularly those with reduced postcranial pneumatization, such as *Phalacrocorax carbo* or *Alca torda*. This foramen, when present, is generally large and circular, as in *Anas platyrhynchos* and *Gallus gallus*, ovoid as in *Ardea alba* or *Rynchops flavirostris*, or tear-shaped like in *Chroicocephalus novaehollandiae*, although certain taxa exhibit a condition where the foramen is divided by a medial septum, as in *Phaethon lepturus*. In certain taxa no single foramen is present, but this region of the sternum is instead perforated by a large number of smaller foramina, as in *Fregata minor*, *Puffinus lherminieri*, and, to a more extreme degree, *Chauna chavaria*. A foramen exhibiting a quadrangular shape similar to that of *Ichthyornis* was only observed in certain strisoreans, such as *Caprimulgus macrurus* and *Podargus strigoides* amongst the surveyed crown-group birds. This pneumatic foramen is absent in Hesperornithes, which are highly osteosclerotic and exhibit reduced skeletal pneumaticity (Bell & Chiappe, 2016, 2020), and has not been previously reported in euornitheans stemward of *Ichthyornis*.

**Figure 10.**
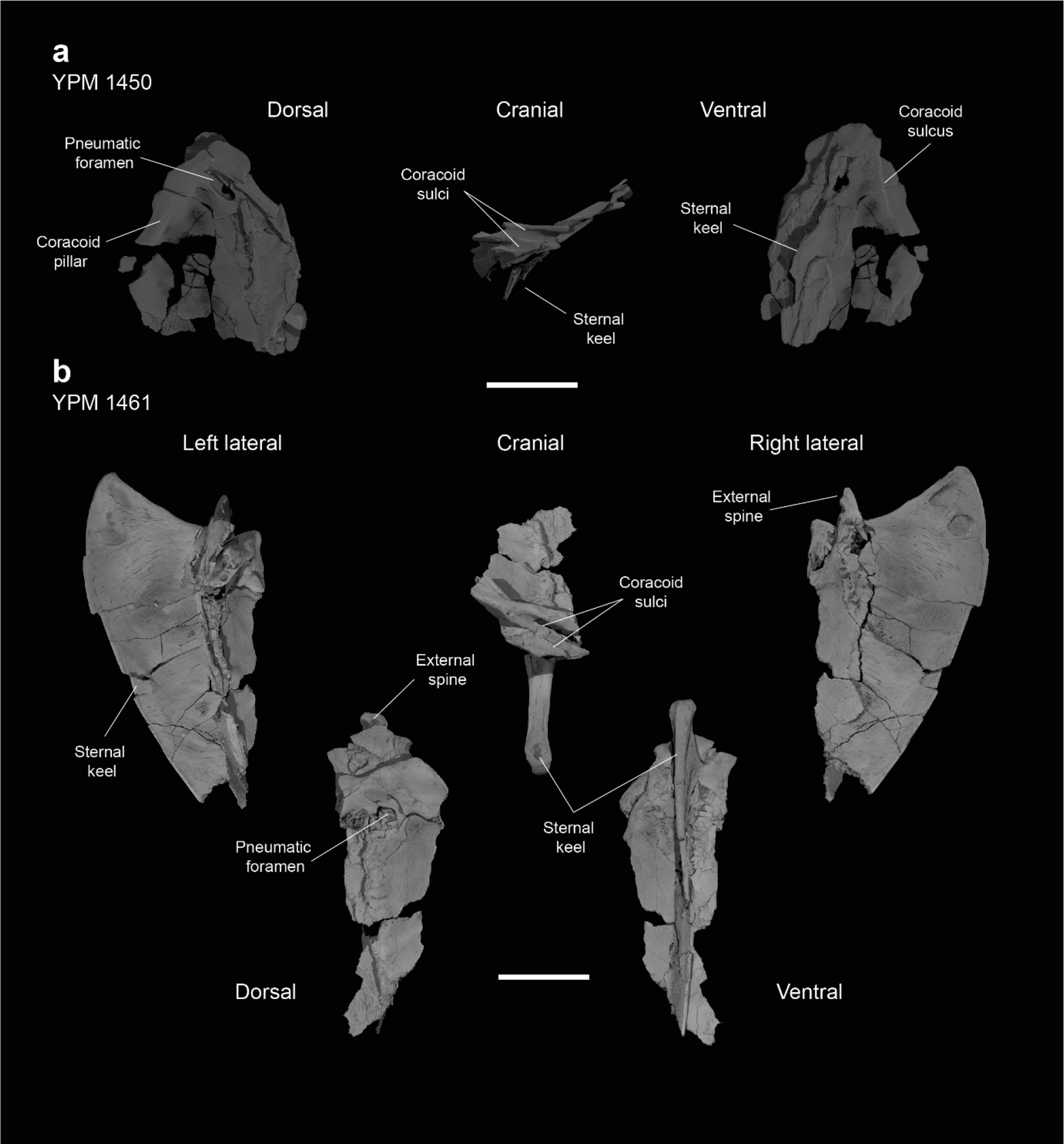
Sterna of *Ichthyornis*. (a) YPM 1450 in dorsal, cranial and ventral views and (b) YPM 1461 in left lateral, dorsal, cranial, ventral and right lateral views. Scale bar equals 1 cm.

Five costal processes are found posterior to the caudal edges of the craniolateral processes. Clarke (2004) describes the cranial- and caudalmost costal processes as only indicated by a narrowing in the edge of the sternum, but this does not seem to be the case in the new specimens described here, in which only the cranialmost process lacks a clear anterior edge. Minute foramina are visible in cross-section between the costal processes, found on the caudal edge of each process, just under the dorsal ridge at the apex of each process, in a position similar to the costal pneumatic foramina of some extant birds such as *Anser albifrons* (Anatidae) and *Fregata aquila* (Fregatidae). These foramina are present in FHSM VP-18702 and NHMUK A 905, and connect to an infilled internal cavity, though they are obscured by radiopaque inclusions in KUVP 119673. The presence of pneumatic foramina between the costal processes has previously been considered a synapomorphy of the avian crown group (Clarke, 2004). The foramina documented here are smaller than the pneumatic foramina observed in most extant birds, so their pneumatic nature remains to be confirmed, since small pneumatic foramina have been previously described as externally indistinguishable from nutrient and neurovascular foramina (O’Connor, 2006).

No lateral or ziphoid processes are developed caudal to the costal processes, contrasting with the condition in *Archaeorhynchus* (Zhou et al., 2013), *Yixianornis*, and *Gansus*. Clarke (2004) reported that no lateral trabeculae or caudolateral processes (proc. caudolateralis; Baumel & Witmer, 1993) were present in *Ichthyornis*, but the bases of these structures seem apparent in NHMUK A 905, which was described in that publication.

Caudolateral trabeculae are well preserved in both FHSM VP-18702 and KUVP 119673 as well. The caudolateral processes are robust and craniocaudally wide, being twice as long as the craniolateral processes. The dorsocranial edge of each caudolateral process exhibits a very shallow, rounded, cranially-directed extension halfway along the length of the process, but no distal flare is present, unlike in *Yanornis* (Zhou & Zhang, 2001; Wang et al., 2020) or *Yixianornis* (Clarke et al., 2006). The caudal end of the process is rounded and wide, showing an unfinished bone texture both in FHSM VP-18702 and NHMUK A 905.

The caudal end of the sternum of *Ichthyornis* has only been briefly reported in AMNH FARB 32773 (Torres et al., 2021), in which it is poorly preserved. Among the specimens described here, it is well preserved in FHSM VP-18702 and KUVP 119673, and we therefore describe this portion of the sternum in detail for the first time. This region clearly shows open lateral and medial sternal incisures medial to the caudolateral processes (Fig. 9b). These incisures are wide and rounded, and they do not penetrate deeply into the edge of the bone.

Torres et al. (2021) reported the presence of medial incisures and possible lateral incisures, although they could not verify whether the lateral incisure was a taphonomic artifact. The lateral incisure present in FHSM VP-18702 and KUVP 119673, congruent with the condition in AMNH FARB 32773, has a flat lateral margin and slightly concave medial margin, while the medial incisure shows a concave lateral margin and a slightly convex medial margin. The medial incisure extends further cranially than the lateral incisure, in a manner similar to that in the tern *Sterna hirundo* and the gull *Chroicocephalus novaehollandiae*. This morphology lacks clear analogues amongst other Mesozoic euornitheans, in which the lateral incisure tends to extend more cranially and the medial incisure is usually enclosed, forming a fenestra (O’Connor & Zelenkov, 2013; Wang et al., 2013, 2016; Zhou et al., 2014).

Both rounded incisures delimit a short, robust and flat intermediate trabecula (trabecula intermedia; Baumel & Witmer, 1993), preserved in all three studied specimens. The trabecula extends slightly less far caudally than the caudolateral processes, and it is straight and quadrangular in FHSM VP-18702, pointing caudally and widening slightly on its caudal end before ending abruptly, which might indicate breakage (Fig. 9a). The shape of the trabeculae differs in KUVP 119673, where they are short, rounded and caudomedially directed, with the medial edge extending further caudally than the lateral edge. This difference might be taphonomic since the morphology varies somewhat between the two sides of KUVP 119673 (Fig. 9b). The median trabecula is short, wide, and rounded, extending as far caudally as the intermediate trabeculae.

The sternal keel originates from a broad and flat ridge extending ventrally from the external spine, and the ventral apex (apex carinae; Baumel & Witmer, 1993) extends cranially slightly beyond the tip of the external spine (Fig. 9a; 10b). The keel is deep and well developed, extending for the entire craniocaudal length of the sternum. The preservation of the bone surface on the sternal keel is patchy in FHSM VP-18702 and NHMUK A 905, and no muscular attachment lines are clearly discernible. The cranial surface of the keel is flat and wide, becoming progressively narrower ventrally but widening slightly at the ventral apex.

The ventral margin of the keel is thick and wide, forming a ridge which progressively narrows towards the posterior region of the keel.

### FURCULA

Four specimens described here include fragments of the furcula: BHI 6420, ALMNH PV93.2.133 and KUVP 25469 preserve the omal end, while NHMUK A9 905 preserves the clavicular symphysis (Fig. 11). Previously, only two incomplete furcula fragments had been reported for *Ichthyornis*: YPM 1755, corresponding to the fused region between both clavicles, and SMM 2503, a fragment including the right omal tip (Clarke, 2004). The morphology of the newly described furcular fragments differs significantly from that of the previously referred specimens, which, given the scarcity of furcular material, may be attributable to taphonomic distortion of the previously described material.

**Figure 11.**
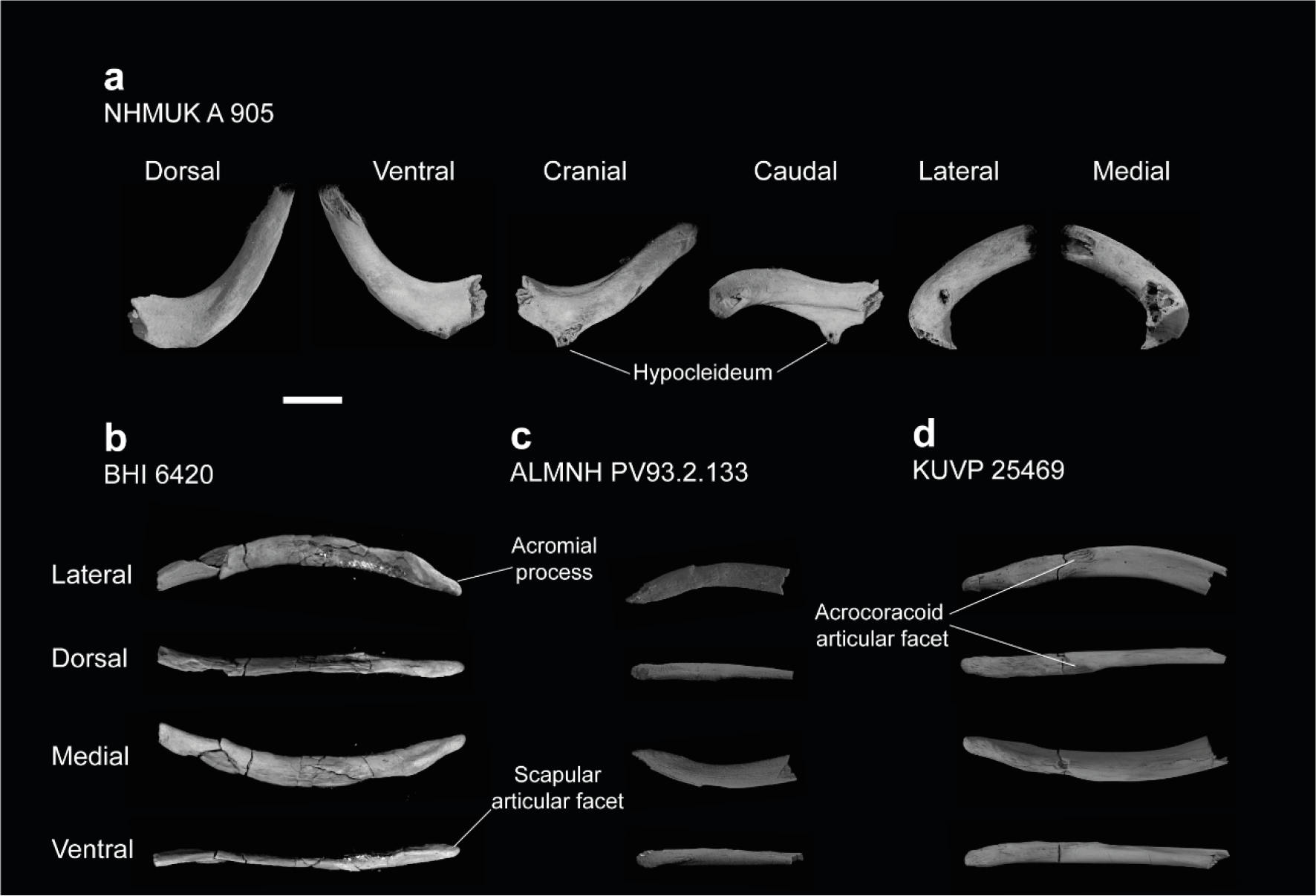
Furculae of *Ichthyornis*. (a) NHMUK A 905 medial furcula fragment in dorsal, ventral, cranial, caudal, lateral and medial views; (b) BHI 6420 left omal furcula fragment, (c) ALMNH PV93.2.133 right omal furcula fragment and (d) KUVP 25469 right omal furcula fragment, in lateral, dorsal, medial, and ventral views. Scale bar equals 5 mm.

The clavicular symphyseal region is preserved in NHMUK A 905 together with part of the left clavicular ramus (Fig. 11a). The furcula is roughly U-shaped, and while the broken clavicular rami prevent a precise measurement of the interclavicular angle, the morphology appears very similar to that of *Yixianornis* (Clarke et al., 2006) and *Iteravis* (Zhou et al., 2014), in which the clavicles meet at an angle of approximately 60°. Both clavicles are completely fused at the midline, and no suture is visible. *Contra* Clarke (2004), a clear hypocleideum (apophysis furculae; Baumel & Witmer, 1993) is preserved extending from the ventral surface of the symphysis, but only the base of the hypocleideum is preserved. A hypocleideum is not present in any other crownward euornitheans (O’Connor & Zelenkov, 2013; Zhou et al., 2014; Wang et al., 2016b) but it is widespread amongst Enantiornithes and extant crown birds (Nesbitt et al., 2009; Mayr, 2017). The hypocleideum is elongate and subtriangular in ventral view, and its caudal end extends caudally and ventrally to the clavicles. Despite being incomplete, the shape of the preserved hypocleideum points towards an elongated and enlarged lamina, similar to the condition in Laridae (e.g., *Sterna hirundo*), and Galliformes (e.g., *Gallus gallus*), instead of a reduced tubercle as in Procellariiformes like *Puffinus lherminieri* and Anseriformes like *Anas platyrhynchos*. It is unclear whether the hypocleideum contacted the apex of the sternal keel, as in some marine birds such as *Phaethon lepturus, Fregata minor*, *Morus bassanus*, and *Phalacrocorax carbo*.

Ridges extend on both craniolateral ends of the hypocleideum along the entire preserved cranial surface of the left clavicular ramus. These ridges delimit a subtriangular flat-to-convex surface on the cranial side of the symphyseal region. Similar ridges extend from the caudolateral edges of the hypocleideum onto the caudal or sternal surface of the clavicle, delimiting a concave depression on the sternal surface of the symphyseal region. The preserved left clavicle is subcircular in cross-section close to the symphyseal region, but becomes progressively mediolaterally compressed caudally, as the ridges extending from both the cranial and caudal ends of the hypocleideum become more marked.

The omal end of the furcula is preserved in BHI 6420 (left clavicle), KUVP 25469 (right clavicle) and ALMNH PV93.2.133 (right clavicle). The omal tip of the furcular ramus is tapered and very elongate, terminating in a pointed end instead of a blunt end as previously described (Clarke, 2004). Although most of the clavicular ramus is not preserved in either specimen, the cranial end of KUVP 25469 is slightly bowed ventrally and shows a shallow lateral excavation, bounded by marked ridges on both the dorsal and ventral edges of the lateral surface (Fig. 11d).

The articular facet for the acrocoracoid process sits on the lateral side of the clavicular ramus and is very poorly developed. It does not extend laterally from the main body of the clavicle (Fig. 11d). The facet is delimited cranially by a short ridge or tubercle, and forms a shallow circular concavity. The morphology of this facet is sparsely described in existing Cretaceous euornithean literature; it does not seem to extend beyond the ramus of the clavicle in any crownward euornithean taxa (Zhou et al., 2014; Wang et al., 2016b). Among Neornithes, the poorly developed acrocoracoid facet is seen in many taxa, including the shearwater *Puffinus lherminieri* and the plover *Charadrius rubricollis*. Just omal to the acrocoracoid facet there is a shallow, elongate facet that seems to correspond with the elongate and flattened medial surface of the procoracoid process. The omal end of the clavicle extends into an elongate and pointed acromial process, though it is not as well developed or pointed as it is in *Iaceornis* (Clarke, 2004). The process is robust, and it is not mediolaterally compressed as in the extant. *P. lherminieri* or *C. rubricollis*, instead remaining subcircular in cross-section for the whole of its length. The acromial process bears a short and flattened facet on the ventral surface of its omal tip. The dorsal portion of this facet seems to contact the shortened acromion process of the scapula near its base.

### CORACOID

Nine of the newly described specimens preserve coracoids: BHI 6420 (both), FHSM VP-18702 (both), NHMUK A 905 (both), ALMNH PV93.2.133 (right side), KUVP 119673 (right side), KUVP 2281 (left side), KUVP 5969 (right side), and RMM 7841 (right side). Although coracoids are among the most commonly preserved elements from *Ichthyornis,* with 24 coracoids in the YPM collections (Clarke, 2004), the coracoids of several of the newly described specimens (BHI 6420, FHSM VP-18702, KUVP 119673 and KUVP 2281), are among the best-preserved coracoids known for *Ichthyornis* and together reveal new information about the element’s morphology (Fig 12). Measurement of the coracoid of *Ichthyornis* specimens are provided in Table 4.

**Figure 12.**
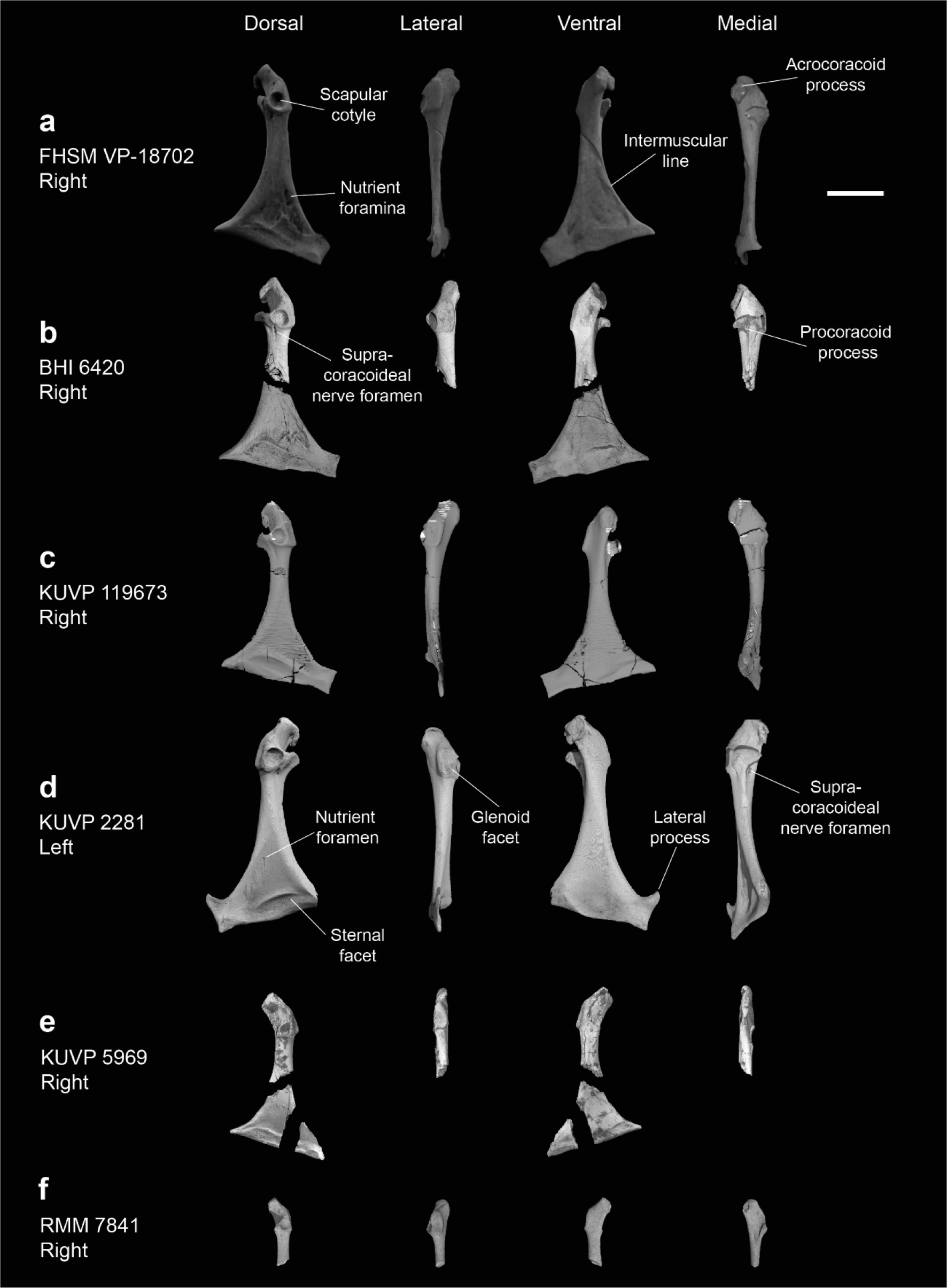
Coracoids of *Ichthyornis*. (a) FHSM VP-18702 right coracoid, (b) BHI 6420 right coracoid omal and sternal fragments, (c) KUVP 119673 right coracoid, (d) KUVP 2281 left coracoid, (e) KUVP 5969 right coracoid and (f) RMM 7841 right coracoid, in dorsal, lateral, ventral and medial views. Scale bar equals 1 cm.

**Table 4.**
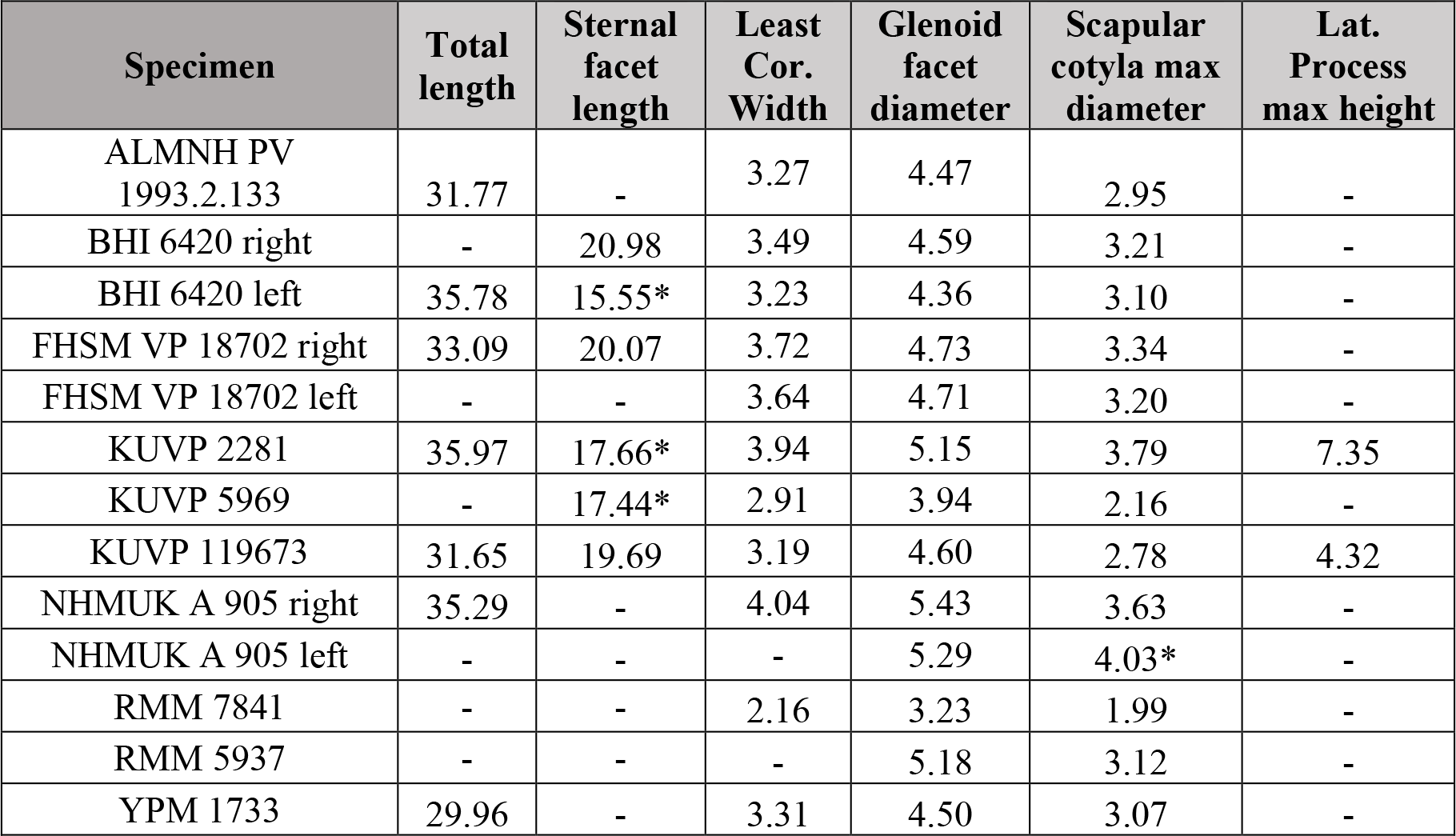
Measurements of the coracoid of *Ichthyornis* specimens. Standard coracoid measurements are taken from Field et al., (2013). Total length corresponds to the maximum coracoid length measured from its omal to sternal end. Sternal facet length corresponds to the maximum mediolateral expansion of the coracoid sternal facet. Least coracoid width corresponds to the minimum mediolateral width of the coracoid shaft. Glenoid facet diameter corresponds to the maximum diameter of the glenoid or humeral articular facet. Asterisks (*) denote measurements that might be unreliable due to breakage or distortion, but are included for completeness. All measurements are in mm. - = not measurable.

The studied specimens exhibit a broad size distribution (Table 4), with the largest specimen, KUVP 2281 (4.00 cm in length), being 38% longer than the smallest complete coracoid, KUVP 5969 (2.91 cm in length). Several features of the coracoid, such as total coracoid total length, shaft width, and particularly the maximum diameter of its glenoid facet, show strong correlations with body mass in extant volant birds (Field et al., 2013), thus revealing a wide size distribution for *Ichthyornis* (Fig. 34; see Body size estimates below). Despite these differences in size, the proportions of the omal and sternal regions of the studied coracoids remain essentially constant, though the effects of allometry are evident in the proportional length of the shaft, which is longer in the largest specimens (Table 4). For example, the sternal edge of the coracoid is 78% as long as the shaft in KUVP 5969, 65% in FHSM VP-18702 (total length 3.723 cm) and KUVP 119673 (total length 3.577 cm), and only 59% in KUVP 2281. Although generally similar in their overall morphology, the coracoids included in this study show some minor variation. While impossible to definitively resolve, we suspect these differences correspond to intraspecific variation, though some previous research has asserted that coracoid morphology may be relatively strongly conserved within bird species (Longrich, 2009; Longrich et al., 2011).

The morphology of the omal region is very similar to that of previously referred *Ichthyornis* specimens (Clarke, 2004). The shape of the scapular cotyle is subcircular and very deeply excavated in all specimens, though the shape of the surrounding rim varies somewhat, ranging from essentially circular (KUVP 5969) to ovoid (KUVP 119673, 2281) to subtriangular (BHI 6420, ALMNH PV93.2.133) (Fig. 12). The dorsocaudal opening of the supracoracoideal nerve foramen is positioned just caudal to the scapular cotyle. The opening lies in a short groove extending from the edge of the scapular cotyle on most specimens, as described by Clarke (2004), but this structure is not visible in either of the FHSM VP-18702 coracoids, similar to YPM 1446, which also lacks this groove (Clarke, 2004).

The glenoid facet is large and slightly concave, with very marked ventral and caudal margins forming a labrum, which separates the facet from the shaft. It is situated just lateral to the scapular cotyle, extending from its caudal end to the base of the acrocoracoid ligament scar (impressio lig. acrocoracohum.; Baumel & Witmer, 1993) in most specimens. In KUVP 5969 the glenoid originates more cranially, with its caudal margin situated in a position roughly equivalent to the centre of the scapular cotyle, instead of its caudal edge. The shape of the glenoid facet in lateral view is ovoid in most specimens but becomes almost quadrangular in KUVP 2281 and RMM 7841 (Fig. 12a, d). The position and shape of the glenoid contrasts with that of other Cretaceous ornithurines such as *Iaceornis* or *Cimolopteryx-*like taxa known mostly from coracoids (Hope, 2002; Agnolin, 2010; Longrich et al., 2011; Mohr et al., 2020), in which the glenoid is situated cranial to the scapular cotyle, a condition shared with most crown-group birds.

Although Clarke (2004, pg. 111) described the acrocoracoid process of *Ichthyornis* as “slightly hooked in posterior view” this structure is considerably more recurved and hook-shaped in BHI 6420, FHSM VP-18702, KUVP 119673, and KUVP 2281 than it is in any of the described and figured YPM specimens (Fig. 12). In these specimens, the acrocoracoid extends dorsocaudally for roughly 25% of the surface of the triosseal canal, curving slightly along its length. It does not taper, and exhibits a large and quadrangular furcular articular surface (Fig. 13a, b). The lack of this very recurved acrocoracoid morphology in NHMUK A 905, ALMNH PV93.2.133, and KUVP 5969 is apparently due to preservational factors, in that this process appears to have been broken or eroded, which is probably true for the numerous coracoids in the YPM collections as well. The cranial surface of the acrocoracoid process is flat to slightly concave, extending parallel to the cranial end of the triosseal canal, except in KUVP 5969, in which it is situated more perpendicularly—though this may be the result of taphonomic distortion. The acrocoracoid ligament scar is shallow and dorsally elongated, and it is clearly delimited by the caudal rim of the acrocoracoid and the cranial edge of the glenoid.

**Figure 13.**
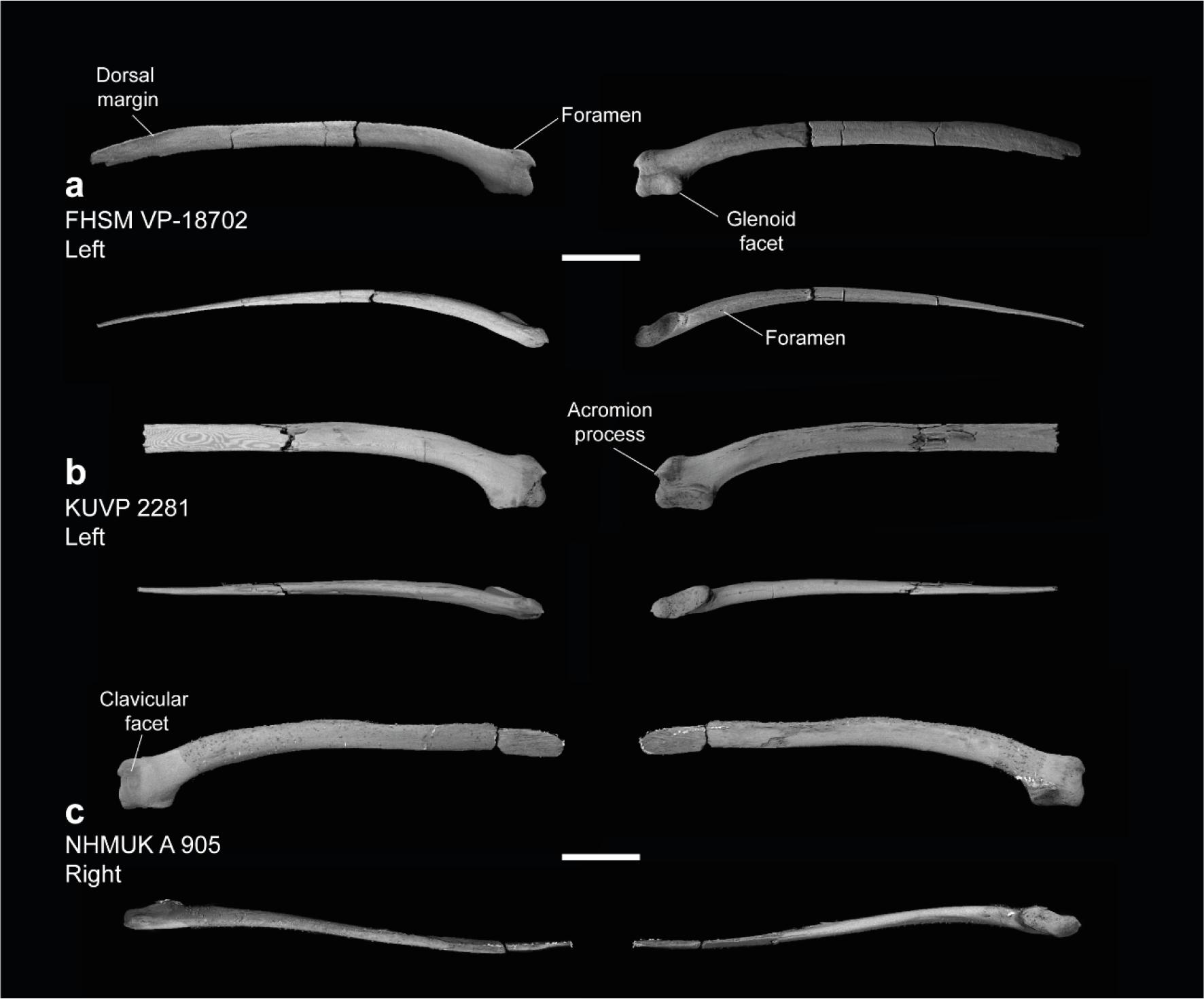
Scapulae of *Ichthyornis*. Views for each in clockwise order: medial, lateral, ventral, dorsal. (a) FHSM VP-18702 left scapula, (b) KUVP 2281 left scapula, and (c) NHMUK A 905 right scapula. Scale bar equals 1 cm.

Similar to the acrocoracoid, the procoracoid process in several studied specimens is much more developed than previously reported; in the specimens lacking a well-developed procoracoid, a clear breakage surface is visible just medial to the scapular cotyle. The procoracoid extends as a narrow subtriangular flange pointing ventromedially, reaching the ventral edge of the supracoracoideus sulcus, which gives the medial surface of the triosseal canal a claw-like shape (Fig. 12b, c, d).

The morphology of the triosseal canal region of the coracoid in *Ichthyornis* is similar to the condition in *Gansus*, *Yixianornis* and *Ambiortus*, which also possess elongated acrocoracoid and procoracoid processes (Clarke et al., 2006; O’Connor & Zelenkov, 2013; Wang et al., 2016b). However, the procoracoid in *Ichthyornis* is substantially more tapered than in the aforementioned taxa, in which it shows a more quadrangular shape with a blunt end. The proportions and shape of the acrocoracoid process of *Ichthyornis* show similarities with those of Lithornithidae (Houde, 1988), and Procellariiformes like the shearwater *Puffinus lherminieri*.

The shaft of the coracoid is long and robust, lateromedially broad, and becomes progressively dorsoventrally compressed sternally, particularly on its dorsal surface. The shaft is slightly more cylindrical and less mediolaterally expanded in KUVP 119673 and RMM 7841, which does not seem to be attributable to preservational factors since KUVP 2281 is exceptionally well-preserved and shows a more dorsoventrally compressed and mediolaterally expanded shaft that does not seem to be taphonomically flattened. The more cylindrical shaft is reminiscent of several isolated ornithurine coracoids from the Late Cretaceous of North America which have been suggested to represent *Ichthyornis*-like forms, such as ‘Ornithurine A’ from Longrich (2009) and Ornithurine D from Longrich et al. (2011), though these differ in other morphological features. Both lateral and medial surfaces of the shaft are strongly concave. A marked ridge extends from the caudal end of the procoracoid process along the medial surface of the bone, reaching the sternal end of the coracoid. The angle formed between the sternal margin of the coracoid and the main axis of the shaft varies among the studied specimens and does not seem to correlate with size, varying from 68° in KUVP 2281, to 72–74° in KUVP 5969, FHSM VP-18702, and BHI 6420, to 81° in KUVP 119673. The medial end of the sternal facet (angulus medialis coracoideum; Baumel & Witmer, 1993) is pointed, with a flattened medial surface. The lateral margin of the coracoid extends into a lateral process, which is broad and quadrangular. This process has only been previously reported in the specimen AMNH FARB 32773, in which it is badly crushed (Torres et al., 2021), and it is only completely preserved in KUVP 2281 (Fig. 12d), which shows a markedly tapered, subtriangular projection that is omally-directed, similar in shape and size to that of *Gansus* and some extant taxa such as *Puffinus lherminieri*.

As in other previously described *Ichthyornis* specimens, the ventral surface of the coracoid shows a shallow depression towards the sternal facet, probably for the implantation of m. supracoracoideus. This depression is variably excavated and delimited in the newly described specimens, showing a clear medial edge and a marked intermuscular line on its lateral edge running from the omal to the sternal end of the shaft in most specimens. Only KUVP 2281 shows a well-marked ridge on its sternal edge, as in YPM 1450. The sternal facet is marked and continuous between the dorsal and ventral surfaces of the coracoid, and, as described by Clarke (2004), it shows a clear projecting ridge delimiting the cranial edge of the facet on its dorsal surface. However, unlike the morphology described by Clarke (2004), it does not show a “vaguely sigmoidal” shape, and is instead somewhat quadrangular, with its proximal edge running parallel to the sternal edge of the coracoid (Fig. 12).

The dorsal surface of the coracoid is described by Clarke (2004) as preserving a clearly marked foramen in the impression of m. sternocoracoidei in all YPM specimens. The newly described specimens show the presence of either one (BHI 6420, KUVP 2281) or two foramina (FHSM VP-18702) in this region.

### SCAPULA

Five of the studied specimens preserve scapulae: BHI 6420 (both), FHSM VP-18702 (left side), KUVP 2281 (left side), KUVP 119673 (right side), and NHMUK A 905 (both). The preservation of this element in the new specimens is generally excellent, and these specimens all show complete or nearly complete scapulae, at worst missing only the acromion or the caudal terminus of the bone. The scapulae of FHSM VP-18702 and NHMUK A 905 may represent the best-preserved scapulae known for *Ichthyornis*. Measurements of the scapulae of the studied specimens are provided in Table 5.

**Table 5.** Measurements of the scapula of *Ichthyornis* specimens. Total length corresponds to the maximum scapular craniocaudal length. Glenoid facet length corresponds to the maximum craniocaudal extension of the glenoid or humeral articular facet. Omal height corresponds to the maximum dorsoventral extension of the omal end of the scapula, measured from the dorsalmost point of the omal end to the ventralmost point of the glenoid facet. Asterisks (*) denote measurements that might be unreliable due to breakage or distortion, but are included for completeness. All measurements are in mm. -= not measurable.

| Specimen | Total length | Glenoid facet length | Omal height |
| --- | --- | --- | --- |
| FHSM VP 18702 | 58.27 | 6.52 | 6.26 |
| KUVP 2281 | 52.46* | 7.19 | 6.95 |
| KUVP 119673 | 56.83 | 5.84 | 4.97* |
| NHMUK A 905 Right | 52.76* | 6.27 | 7.17 |
| NHMUK A 905 Left | 57.54 | 6.39 | 6.57 |

The omal end of the scapula is short, robust, and quadrangular in shape. The dorsal surface of the omal region is straight in all specimens with the exception of KUVP 2281, in which the region just distal to the acromion is raised (Fig. 13b), similar to the condition in YPM 1718. The base of the acromion is robust and inflated, with a marked depression on its ventral end that seems to accommodate the elongated terminus of the acromial process of the furcula.

The acromion is missing or distorted in most specimens, but it is well preserved in FHSM VP-18702 and KUVP 2281. In both specimens the acromion process extends slightly beyond the coracoid tubercle (tuberculum coracoideum; Baumel & Witmer, 1993), a condition differing from that seen in the YPM *Ichthyornis* scapulae, in which this process was described as being markedly shorter (Clarke, 2004). The shape of the acromion process differs among the new specimens. In FHSM VP-18702 the acromion is hook-shaped, curving slightly ventromedially towards the coracoid tubercle (Fig. 13a). The tip of the acromion is missing in BHI 6420, but the shape of the preserved base suggests a similar condition to that of FHSM VP-18702. The base of the acromion is similarly shaped in NHMUK A 905 as well, but in this case the acromion is short, relatively undeveloped, and ends in a rounded, unbroken tip (Fig. 13c). By contrast, the acromion in KUVP 2281 is conical and points cranially (Fig, 13b), similar to the shape seen in YPM 1452, the holotype of *Ichthyornis victor* (Marsh 1880, Clarke, 2004).

This variation in morphology is particularly notable, as an extremely diminutive acromion has been considered one of the autapomorphies of *Ichthyornis dispar* (Clarke, 2004), but no variation in the shape of the acromion or its orientation has been previously reported. It should be noted that, despite the presence of several scapulae among the YPM specimens, only a few of them preserve the proximal portion of the scapula or the acromion process, and only four specimens preserving this morphology have previously been mentioned or figured (YPM 1452, 1718,1763; Clarke, 2004; and AMNH FARB 32773, Torres et al., 2021). The disparity in the shape and length of the acromion noted here suggests either intraspecific or interspecific variation, which may have previously gone unappreciated due to the relative ease with which this element may break or erode, given its absence from many otherwise well-preserved specimens.

Despite this variation in shape, when preserved the acromion is very short in all *Ichthyornis* scapulae, which contrasts with the greatly elongated condition seen in other Late Cretaceous euornithean taxa like *Gansus*, *Apsaravis*, and *Iaceornis* (Clarke, 2004; Clarke & Norell, 2002; Wang et al., 2016b). The proportions and general shape of the acromion in *Ichthyornis* are most similar to those of *Yixianornis* (Clarke et al., 2006) and *Hongshanornis* (Chiappe et al., 2014), in which it is short, pointed, and sharply tapered, though still more elongate than in any *Ichthyornis* specimens. Amongst extant taxa, the acromion is greatly reduced in Gaviidae (e.g., *Gavia arctica*) and Anhimidae (e.g., *Chauna torquata*), but neither show the pointed or recurved morphology seen in *Ichthyornis*.

The coracoid tubercle is large and ball-shaped in all of the new specimens described here, but it is slightly more proximally extended in FHSM VP-18702 (Fig. 13). The glenoid facet is large, ovoid and mostly flat, though its distal end is medially extended. It shows very little variation among the different specimens (Fig. 13), with a morphology congruent with that described by Clarke (2004) for the YPM specimens.

No obvious pneumatic foramina are present, but FHSM VP-18702 preserves a very clear, small foramen situated on its medial surface, just caudal to the base of the acromion, which connects with a large internal hollow chamber in this region (Fig. 13a). Such a hollow chamber is also present in all of the other specimens investigated here, though the foramen is not. It is difficult to establish whether this represents a genuine pneumatic structure, as such foramina are rarely unambiguously pneumatic in fossil and extant taxa, though the hollow internal cavity and thin bone walls of the element would be consistent with it being pneumatized (O’Connor, 2006). The presence of a pneumatic foramen on the lateral surface of the proximal end of the scapula is widespread amongst Neornithes, but it remains to be seen whether it constitutes a true synapomorphy of the group (Clarke, 2004).

The scapular blade is preserved in its entirety in BHI 6420, FHSM VP-18702, NHMUK A 905 and KUVP 119673. The blade is elongate and slightly curved at its proximal- and distalmost ends, but is essentially straight for most of its length. Both BHI 6420 scapulae show greater curvature along the length of the blade with no straight central region—similar to the condition in YPM 1773 (Clarke, 2004). This elongated morphology is widespread amongst crownward euornitheans, such as *Yixianornis, Gansus* and *Apsaravis* (Wang et al., 2016b; Clarke & Norell, 2002; Clarke et al., 2006). The scapular blade is thick and ovoid in cross-section just distal to the omal region, but becomes more mediolaterally compressed along its length, thinning significantly at its distal end. The blade is dorsoventrally broad, and it maintains a similar width for most of its length in all specimens, tapering progressively towards its distal end. A small, thin flange is developed distally along the dorsal edge of the blade (margo dorsalis; Baumel & Witmer, 1993) (Fig. 13a). The surface of the blade is mostly smooth across both the lateral and medial surfaces of the bone, but a shallow groove is present on the distal lateral surface, just ventral to the dorsal flange. This groove has also been described in *Gansus*, *Hongshanornis* and *Ambiortus* (O’Connor & Zelenkov, 2013; Chiappe et al., 2014; Wang et al., 2016b), but is absent in *Yixianornis* (Clarke et al., 2006).

A shallow ridge, starting just distal to the glenoid facet, extends along the cranial region of the blade in all studied specimens. Clarke (2004) describes this ridge as beginning just distal to a small foramen, but it appears to originate well cranial to the foramen in all of the new specimens investigated here. This foramen is situated just next to the medial side of the ridge (Fig. 13a). The shallow scar suggested to be situated just dorsal to the cranial end of the ridge by Clarke (2004) is only visible in the NHMUK A 905 scapulae. Both the ridge and the scar were interpreted as being related to the implantation of m. scapulohumeralis cranialis and caudalis by Clarke (2004).

### HUMERUS

Complete and partial humeri are preserved in 12 of the studied specimens (Fig. 14): BHI 6420 (both, fragmentary), BHI 6421 (right, distal), FHSM VP-18702 (right proximal and left distal), KUVP 119673 (both, complete), KUVP 2300 (left, complete), KUVP 5969 (right, fragmentary), KUVP 25469 (right, distal portion), KUVP 25471 (left, complete), NHMUK A 905 (two left humeri), ALMNH PV93.20.5 (left proximal), ALMNH PV98.20.427.1 (left proximal) and RMM 7841 (both, fragmentary). Among these, KUVP 119673 and KUVP 2300 include the best-preserved humeri for *Ichthyornis* known to date, preserving the entire element and its three-dimensional morphology. KUVP 119673 is particularly remarkable in this regard, as both humeri are preserved, with the left element taphonomically flattened, allowing the effects of taphonomy on humeral morphology to be assessed. Measurements of the humeri of the most complete specimens included in this study are provided in Table 6.

**Figure 14.**
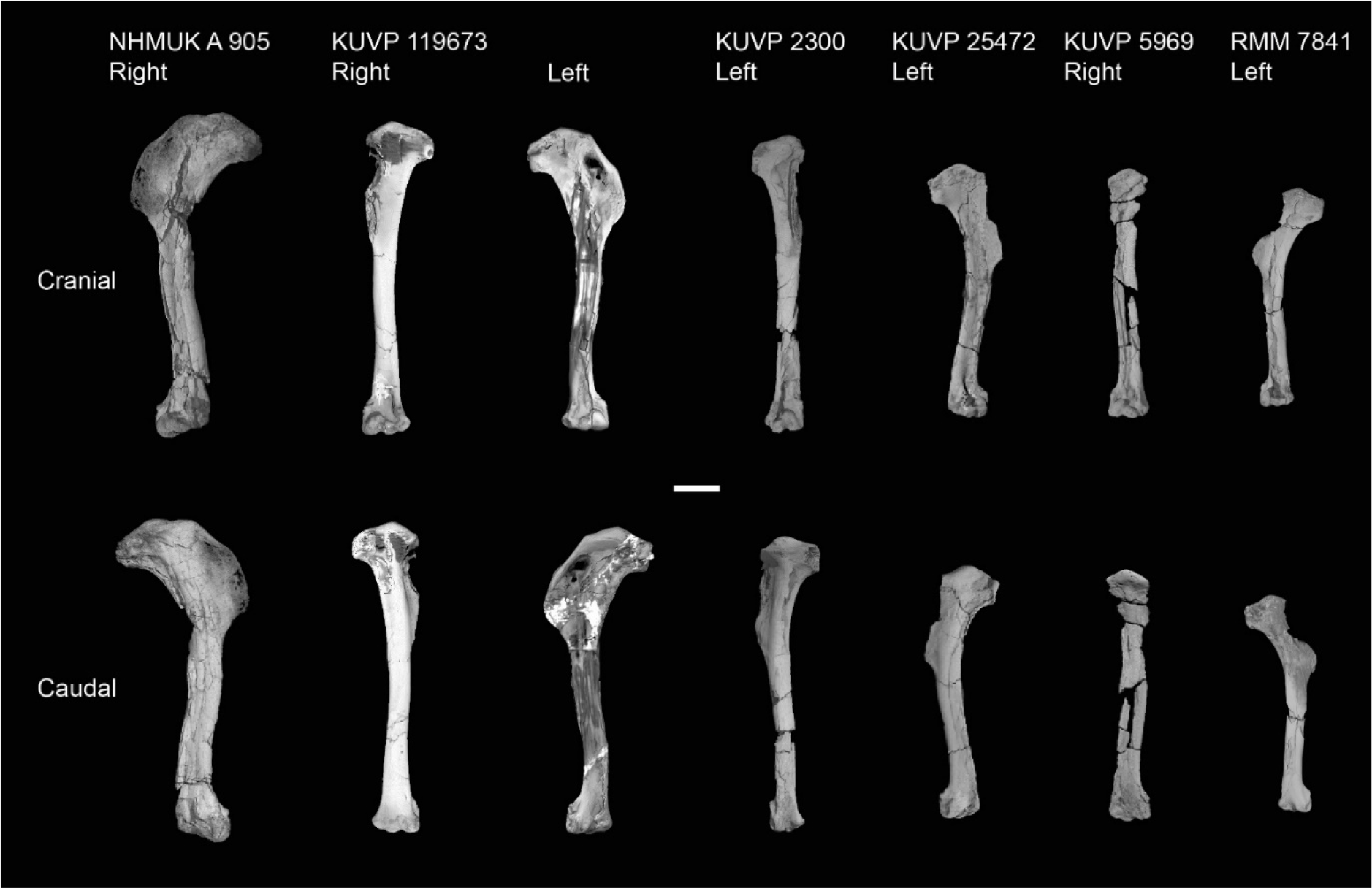
Size distribution of *Ichthyornis* humeri. Largely complete humeri included in this study in cranial and caudal views. Scale bar equals 1 cm.

**Table 6.** Measurements of the humerus of *Ichthyornis* specimens. Total length corresponds to the maximum proximodistal length of the humerus. Least circumference and least diameter correspond to the minimum circumference and diameter of the humeral shaft; midshaft width is provided for those specimens in which the preservational state precluded measuring these. Proximal craniocaudal width is measured as the maximum craniocaudal extension of the humeral head. Proximal dorsoventral width is measured as the maximum extension from the dorsal to the ventral humeral tubercles, including them. Midshaft width corresponds to the craniocaudal width of the humeral shaft halfway along its length. Asterisks (*) denote measurements that might be unreliable due to breakage or distortion, but are included for completeness. All measurements are in mm. -= not measurable.

| Specimen | Total length | Least circum. | Least diameter | Prox. Craniocaudal width | Prox. Dorsoventral width | Dist. Craniocaudal width | Dist. Dorsoventral width | Midshaft width |
| --- | --- | --- | --- | --- | --- | --- | --- | --- |
| ALMNH PV 1988.20.427.1 | - | - | - | 3.62 | 14.02 | - | - | - |
| ALMNH PV 1988.20.55 | - | - | - |  |  |  |  |  |
| BHI 6420 right | - | - | - | 3.49 | 15.83 | 6.94 | 9.70 | - |
| BHI 6420 left | 72.33* | - | - | 3.49 | 16.53 | 6.81 | 9.74 | 6.86* |
| BHI 6421 | - | - | - | - | - | - | 11.98 | 5.32 |
| FHSM VP 18702 right | - | - | - | 4.44 | 17.04 | - | - | - |
| FHSM VP 18702 left | - | - | - | - | - | 12.11 | 5.53 | 6.03 |
| KUVP 2300 | 65.23 | - | - | 3.68 | 12.08 | 6.32 | 7.82 | 4.50 |
| KUVP 5969 | 55.00 | - | - | 3.05* | 8.82* | 4.20* | 8.93 |  |
| KUVP 25469 | - | - | - | - | - | 6.94 | 11.27 | - |
| KUVP 25471 | 55.11 | - | - | 3.06 | 15.50 | 3.72* | 8.41* | 5.17* |
| KUVP 119673 right | 68.96 | 13.11 | 4.01 | 3.66 | 13.73 | 5.27 | 10.80 | - |
| KUVP 119673 left | 67.70 | - | - | 3.55 | 14.16 | 5.72 | 10.32 | 5.54* |
| NHMUK A 905 right | 70.61 | - | - | 4.08 | 16.76 | 5.60 | 11.37 | 6.87 |
| NHMUK A 905 left | - | - | - | 3.86 | 16.61 | - | - | - |
| RMM 7841 left | - | - | - | - | - | 4.57 | 7.59 |  |
| RMM 7841 right | 47.08 | - | - | 2.37 | - | 4.48 | 7.27 | 3.70 |
| RMM 5985 | - | - | - | - | - | 6.06 | 10.80 | - |

The proximal end of the humerus is best preserved in ALMNH PV98.20.427.1, which, despite lacking a deltopectoral crest and being broken distally, preserves the entire proximal end undistorted and in three dimensions (Fig. 16). The humeral head (caput humeri; Baumel & Witmer, 1993) is large, globose, and proximally convex in caudal view. It arches over the caudal surface of the humerus, exhibiting a subtle, hook-shaped apex curving medially at its dorsal end. The proximal development of the humeral head varies among the studied specimens: it is particularly well developed in ALMNH PV98.20.427.1 and BHI 6420, but is much less so in KUVP 119673. A marked and distinct straight ridge extends dorsoventrally from the dorsal tubercle, ending just dorsal to the capital incisure. This ridge crosses over the caudalmost extension of the humeral head and delimits its distal edge.

The ventral tubercle is well developed and distinct where preserved, but its shape varies among the different specimens. The shape is subcircular to ovoid in KUVP 25471 and FHSM VP 18702, but it is subquadrangular in ALMNH PV98.20.427.1 and KUVP 119673, and practically square in BHI 6420. This shape variation does not seem to be taphonomic in origin, since most of the specimens preserving the ventral tubercle are extremely well-preserved and do not show evidence of significant deformation.

The tricipital fossa (fossa tricipitalis; Baumel & Witmer, 1993) is extremely shallow and moderately excavated (Fig. 16), and is bound dorsally by a broad and marked dorsal ramus (crus dorsalis; Baumel & Witmer, 1993) extending distally from the ventral tubercle and ventrally by a thin and narrow ventral ramus (crus ventralis; Baumel & Witmer, 1993). This shallow tricipital fossa is moderately visible in ALMNH PV98.20.427.1, but it is significantly more marked in flattened specimens such as BHI 6420 and the left humerus of KUVP 119673, since both the dorsal and ventral rami become more apparent after deformation. A large and deep depression found just dorsal to the ventral process and ventrodistal to the humeral head might correspond to the secondary opening of the tricipital fossa as seen in certain extant birds such as *Chroicocephalus novaehollandiae*. This larger depression is bounded proximally by the proximal edge of the humerus, and it is clearly observable in all specimens preserving this region (Fig. 16). Neither of the two tricipital fossa depressions show evidence of being associated with pneumatic foramina; for example, the opening evident in the right humerus of KUVP 119673 appears to be the result of breakage.

The bicipital crest is well developed in *Ichthyornis,* extending distally from the ventral tubercle, and is quadrangular in caudal view and vaguely rhomboidal in ventral view.

Although the preservation of this structure in most of the studied specimens is too poor to allow clear observations, ALMNH PV98.20.427.1 preserves the undistorted bicipital crest in its entirety. The cranial surface of the bicipital crest is rounded and convex, but its caudal surface is flat or slightly concave. A large rounded and shallow depression is found on the proximal portion of the ventral surface of the crest, probably corresponding to the attachment point of m. bicipitalis (Watanabe et al., 2021). A minute but deep rounded pit is present on the caudal surface of the crest halfway along its length, and despite its small size, is clearly visible in ALMNH PV98.20.427.1 and in more flattened specimens such as the BHI 6420 and NHMUK A 905 humeri. This pit does not seem to connect to the internal, hollow cavity of the proximal humerus, and thus probably constitutes a nutrient foramen. An ovoid and deep pit is situated on the distal surface of the bicipital crest and is visible in all the specimens preserving this region of the humerus. Clarke (2004) interpreted this depression as the attachment point for m. scapulohumeralis caudalis (Baumel & Raikow, 1993), and considered it as one of the autapomorphies of *Ichthyornis*.

The deltopectoral crest is well preserved on the left BHI 6420 humerus, both NHMUK A 905 humeri, KUVP 2300, and both KUVP 119673 humeri. The crest is large and has an ovoid or semiquadrangular shape (Fig. 14). The crest is extremely thin, and it becomes progressively thinner towards its edge, where it is bounded by a narrow ridge extending from the proximal dorsal tubercle. The deltopectoral crest is flattened in most of the specimens, complicating the interpretation of any muscle scars—only a subtle concave depression across most of its caudal surface is apparent. Notably, two of the specimens preserve three-dimensional deltopectoral crests: KUVP 2300 and the right humerus of KUVP 119673 (Fig. 15). Although the deltopectoral crests in these specimens still exhibit moderate breakages and distortion, the shape and original orientation of the crest seems to be preserved, revealing that, in life, the crest would have been cranially or dorsocranially directed, similar to the condition in most crown birds (Serrano et al., 2020). KUVP 119673 is particularly remarkable in this regard, since it preserves both humeri, with the right element preserved in three dimensions and the left element flattened (Fig. 15). In the flattened humeri, such as the left humerus of KUVP 119673, BHI 6420 and NHMUK A 905, the deltopectoral crest appears to be dorsally directed, but the preservation of the right humerus of KUVP 119673 reveals that this is the product of taphonomic deformation. The deltopectoral crests of several flattened but otherwise exceptionally preserved fossil avialans, such as *Hongshanornis, Gansus*, and *Yanornis* (Zhou & Zheng, 2001; Chiappe et al., 2014; Wang et al., 2016b), as well as previously described specimens of *Ichthyornis* (Clarke, 2004), have traditionally been interpreted as being dorsally directed, which has led to the hypothesis that a cranially deflected morphology constitutes a synapomorphy of crown group birds (Clarke, 2004; Chiappe et al., 2014). Only the proposed ornithuran *Tingmiatornis* has been described as possessing a slightly cranially deflected deltopectoral crest, but this taxon has never been included in a phylogenetic analysis, and its affinities remain uncertain (Bono et al., 2016).

**Figure 15.**
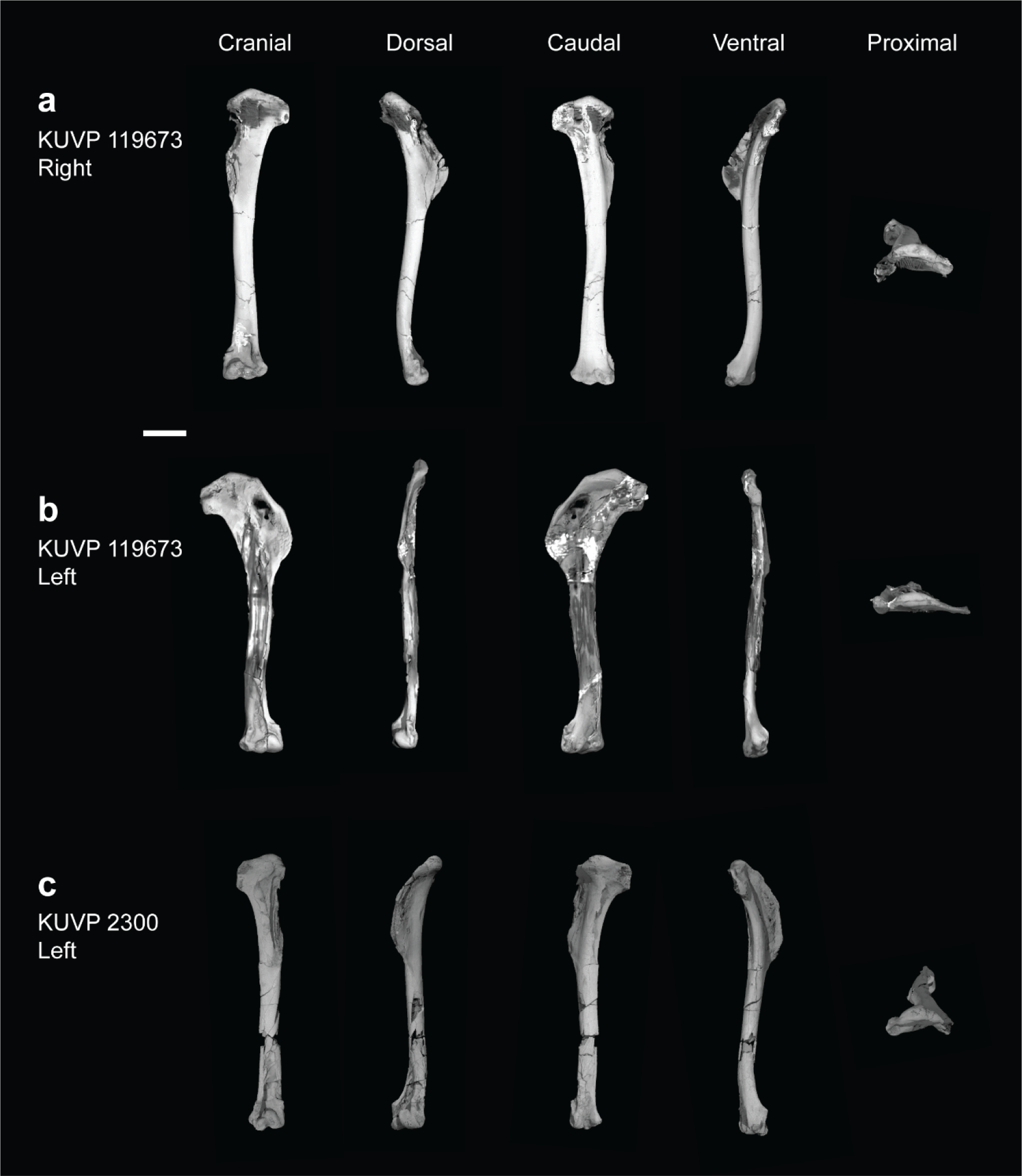
Three-dimensional morphology of *Ichthyornis humeri.* Showing the three-dimensionally preserved humerus of KUVP 119673 (a) and KUVP 2300 (c) in comparison with the flattened left humerus of KUVP 119673 (b), in cranial, dorsal, caudal, ventral and proximal views. Scale bar equals 1 cm.

**Figure 16.**
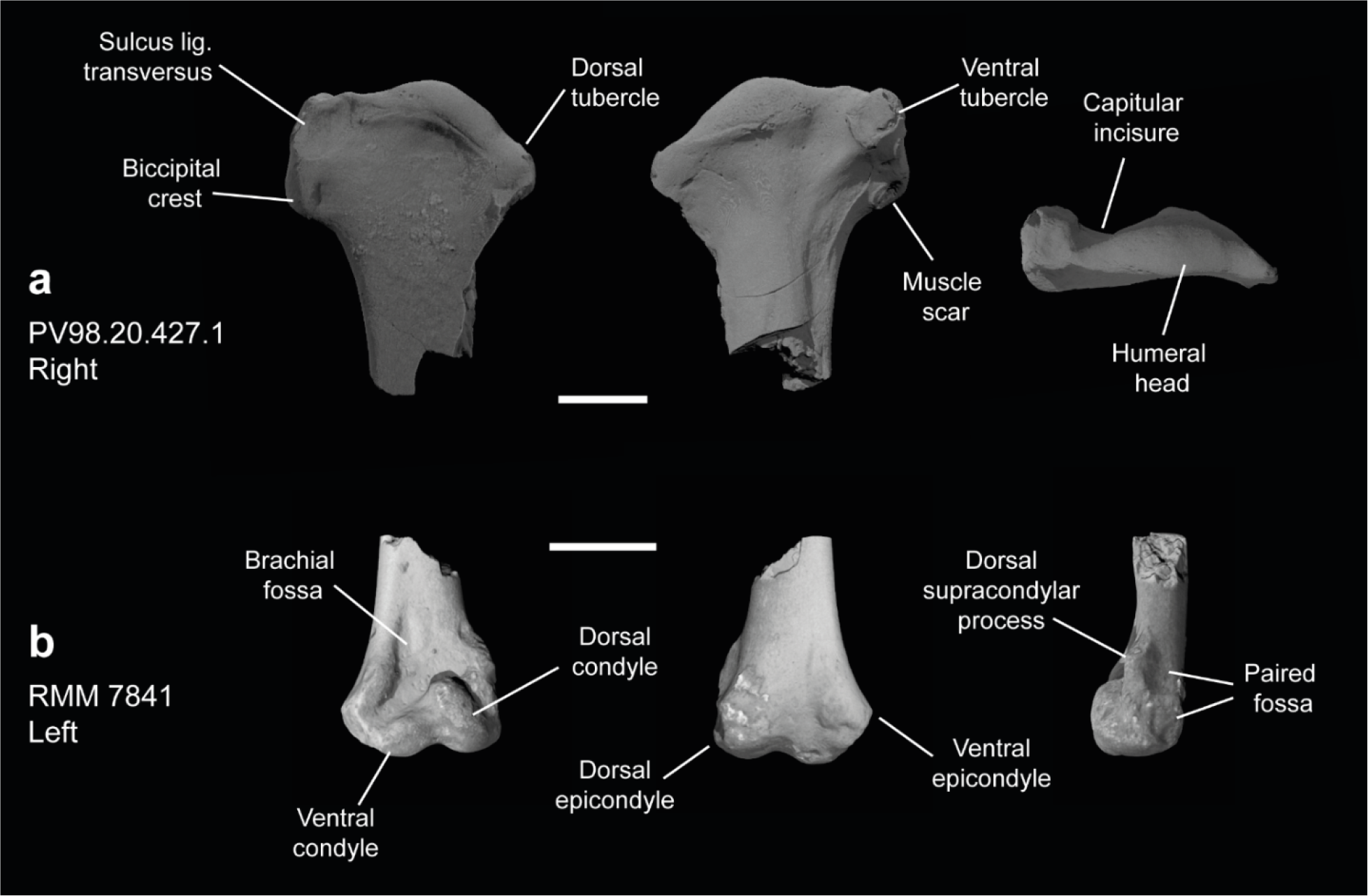
Detailed proximal and distal morphology of *Ichthyornis* humeri. (a) PV98.20.427.1 right proximal humerus in cranial, caudal and proximal views; and (b) RMM 7841 right distal humerus in cranial, caudal and lateral views. Scale bar equals 5 mm.

The new, exceptionally preserved humeri described here underscore the considerable impact of taphonomic distortion on the preservation of the deltopectoral crest, illustrating that at last a partially cranially-directed deltopectoral crest was already present in crownward non-neornithine avialans, and that caution is needed when interpreting the presence or absence of this morphology from flattened fossil remains.

On the cranial surface of the distal end of the humerus there are two well-developed dorsal and ventral condyles of similar length. The intercondylar incisure, running ventrodistally between both condyles, is shallow, and the distinction between the condyles is not clear (Fig. 16b). The dorsal condyle is more globose and is better developed than the ventral condyle, and it curves over the concave brachial depression (fossa m. brachialis; Baumel & Witmer, 1993). A conspicuous nutrient foramen is positioned near the centre of the brachial depression, just proximal to the dorsal condyle. This foramen is present on some, but not all, of the YPM humeri, sometimes being present only in one of the two humeri of an individual, as in the case of the holotype (Clarke, 2004). The dorsal condyle projects proximally, forming a low angle with the ventral condyle, which is almost parallel to the distal edge of the humerus. The ventral epicondyle shows two complex depressions on its ventrodistal end, and another shallow depression on the anterior surface between the ventral epicondyle and the ventral supracondylar tubercle. The morphology of this region is congruent with that described by Clarke (2004).

The dorsal supracondylar process is well developed and clearly preserved, extending further anterodorsally than in previously described specimens (Fig. 16b). The ventral supracondylar tubercle projects cranially but is subtler and much less developed than the dorsal supracondylar process. A shallow fossa is found just distal to the ventral supracondylar tubercle, which Clarke (2004) associated with the attachment point of the m. pronator superficialis. The proximal surface of the dorsal supracondylar process shows complex sculpturing, interpreted by Clarke (2004) as the attachment point for three different muscles.

A central depression runs through the midline on the caudal side of the humeral shaft, ending at the proximal end of the brachial depression. This feature, not mentioned by Clarke (2004), is probably taphonomic in origin. The posterior surface of the distal end of the humerus is completely flattened and crushed, and no morphological observations are possible in this region.

### ULNA

The ulna is preserved in eleven of the new specimens, although only FHSM VP-18702, KUVP 119673 and KUVP 5969 preserve the complete element (Fig. 17). The morphology of the anterior side of the bone is well preserved, but the posterior side is crushed and flattened, obscuring most relevant features. Measurements of the ulnae of the specimens included in this study are provided in Table 7.

**Figure 17.**
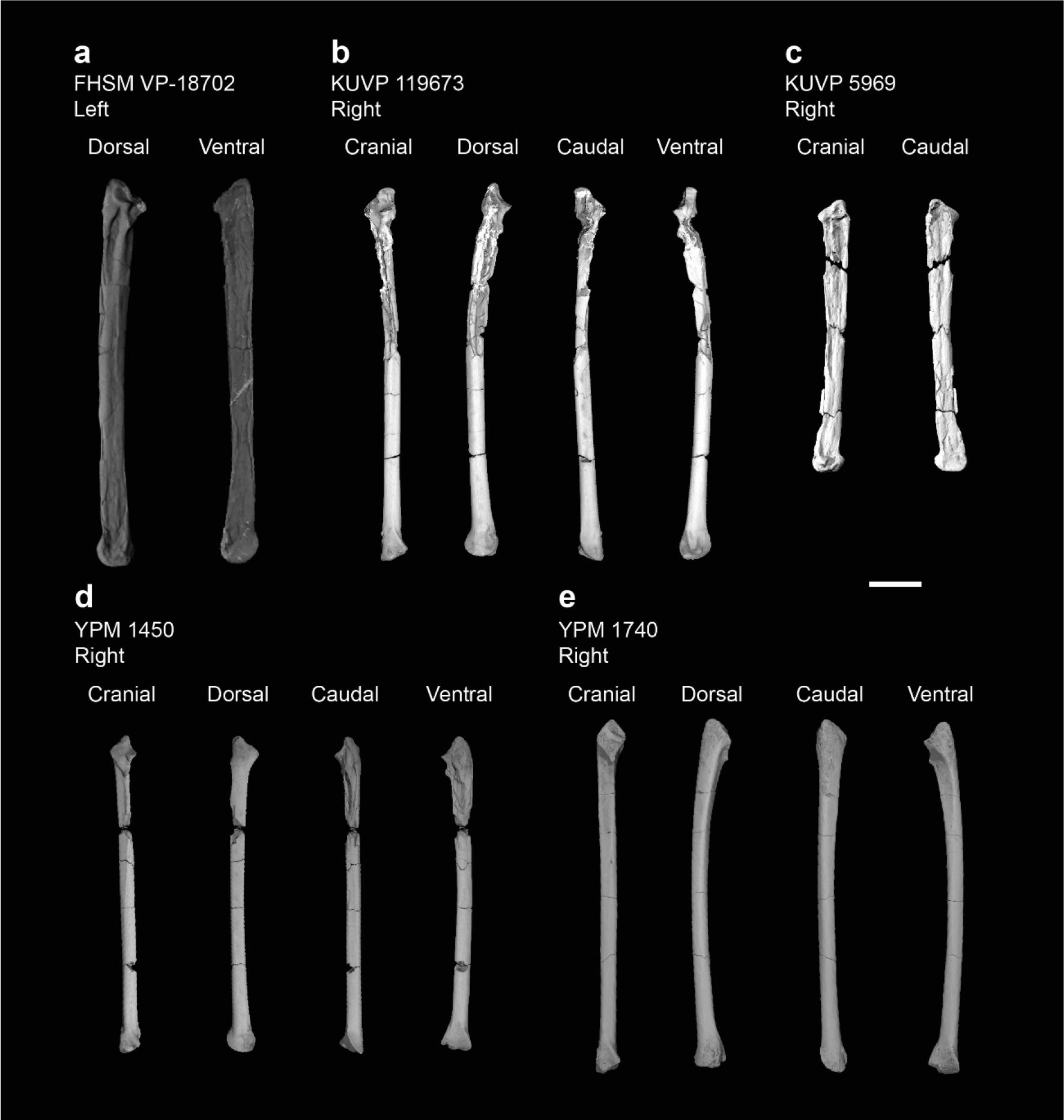
Ulnae of *Ichthyornis*. (a) FHSM VP-18702 in dorsal and ventral views, (b) KUVP 119673 in cranial, dorsal, caudal and ventral views, (c) KUVP 5969 in cranial and caudal views, (d) YPM 1450 and (e) YPM 1740 in cranial, dorsal, caudal and ventral views. Scale bar equals 1 cm.

**Table 7.** Measurements of the ulna of *Ichthyornis* specimens. Asterisks (*) denote measurements that might be unreliable due to breakage or distortion, but are included for completeness. All measurements are in mm. - = not measurable.

| Specimen | Total length | Prox. Craniocaudal width | Prox. Dorsoventral width | Dist. Craniocaudal width | Dist. Dorsoventral width | Midshaft width |
| --- | --- | --- | --- | --- | --- | --- |
| FHSM 18702 | 77.10 | - | 9.25* | - | 7.85* | 4.50* |
| KUVP 5969 | 52.49 | - | 6.23* | - | 6.29 | 3.79 |
| KUVP 119673 right | 71.90 | 4.94 | 7.31* | 4.97 | 6.90 | 3.20 |
| KUVP 199673 left | - | - | - | - | 7.24* | 4.54* |
| RMM 7841 | - | - | - | 4.54 | 4.84 | 2.23 |
| RMM 6201 | - | 6.25 | 6.67 | - | - | - |
| YPM 1450 | 61.51 | 5.05 | 5.75 | 4.24 | 5.71 | 2.84 |
| YPM 1740 | 68.45 | 5.33 | 5.74 | 5.56 | 6.10 | 3.08 |

The shaft of the FHSM VP-18702 ulna is only slightly curved, its general shape moderately straighter than that of the YPM specimens (Fig. 17). As noted by Clarke (2004), the bicipital tubercle is very large and extremely pointed, extending noticeably anteriorly (Fig. 18). The radial depression is a clear and distinct triangular depression adjacent to the bicipital tubercle.

**Figure 18.**
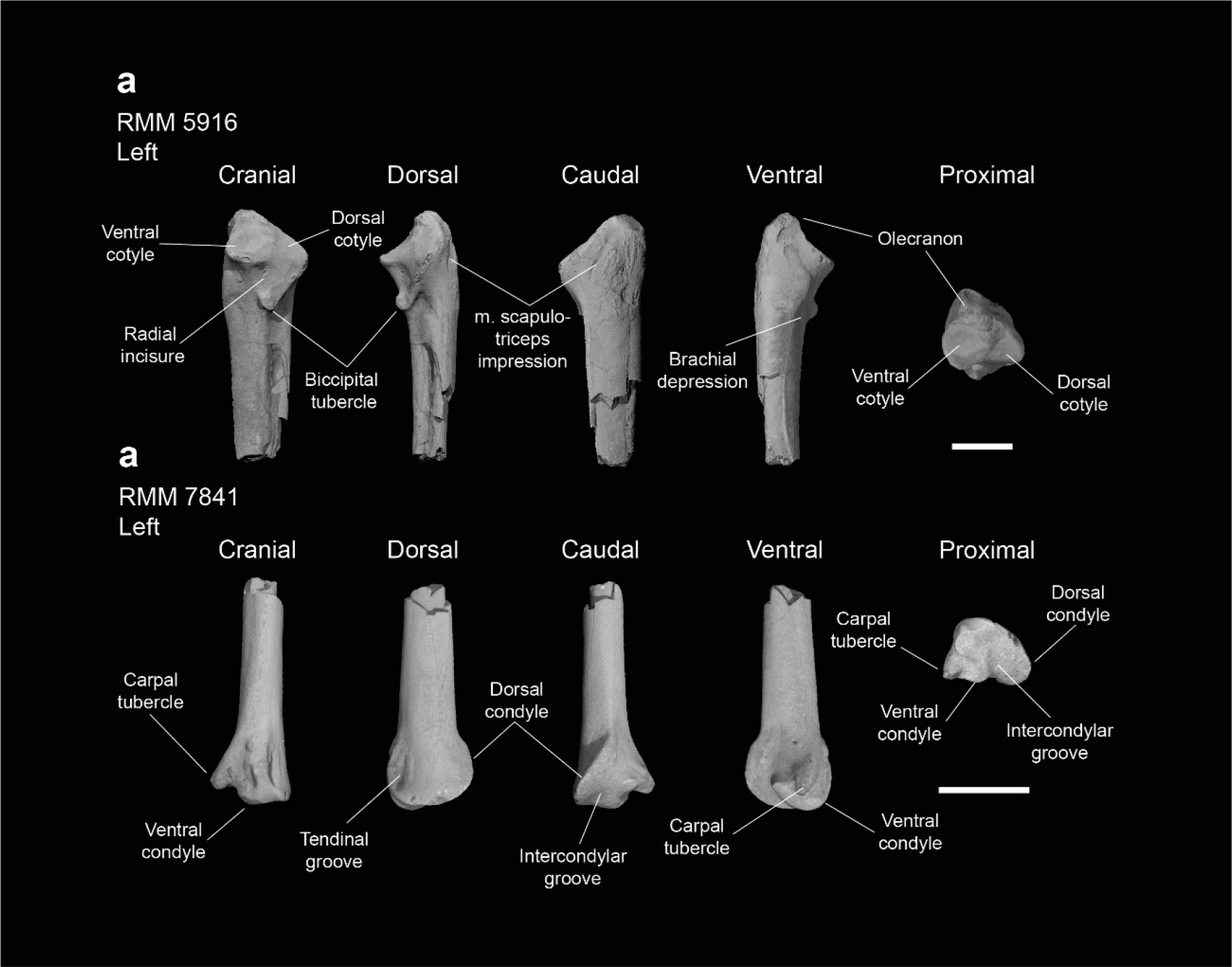
Detailed proximal and distal morphology of *Ichthyornis* ulnae. (a) RMM 5916 left proximal ulna and (b) RMM 7841 left distal ulna in cranial, dorsal, caudal, ventral and proximal views. Scale bar equals 5 mm.

The olecranon process is relatively well developed, with a slightly pointed proximal end, and shows a depression on its anterior side. Clarke (2004) describes a scar present on the ventral surface of the tip of the olecranon process, but this is not clear in FHSM VP-18702.

The dorsal cotylar process (proc. cotylaris dorsalis; Baumel & Witmer, 1993) is subtriangular and extends dorsally well beyond the body of the ulna. Both dorsal and ventral cotyles are well developed and of similar size, and there is no clear ridge separating them (Fig. 18).

From the bicipital process, a well-marked intermuscular line (linea intermusc.; Baumel & Witmer, 1993) extends along at least half the length of the shaft, but it becomes less evident after one third of its length (Fig. 18). A small nutrient foramen is found in line with this intramuscular line, in the same region where the line starts becoming less conspicuous.

Given the preservational state of the posterior side of the ulna, none of the follicular ligament scars identified by Clarke (2004) in most YPM specimens are visible in the FHSM VP-18702 ulna.

### RADIUS

The radius is preserved in ten of the new specimens included in this study. Of these, only the right radius of KUVP 119673 (divided into two fragments) and YPM 1741, and the left radius of FHSM VP-18702 and KUVP 25472 are complete. Of all the other fragmentary radii, the left proximal radius of KUVP 119673 and right distal radius of BHI 6420 and KUVP 25469 are the best preserved. Measurements of the radii of the specimens included in this study are provided in Table 8.

**Table 8.** Measurements of the radius of *Ichthyornis* specimens. Asterisks (*) denote measurements that might be unreliable due to breakage or distortion, but are included for completeness. All measurements are in mm. -= not measurable.

| Specimen | Total length | Prox. Craniocaudal width | Prox. Dorsoventral width | Dist. Craniocaudal width | Dist. Dorsoventral width | Midshaft width |
| --- | --- | --- | --- | --- | --- | --- |
| BHI 6420 | - | - | - | 3.19 | 6.63 | - |
| FHSM 18702 left | 72.20 | 3.14 | 4.52 | 4.578* | 6.92 | 3.403* |
| FHSM 18702 right | - | - | - | - | 6.318* | - |
| KUVP 5969 | - | - | - | - | - | - |
| KUVP 25469 | - | - | - | 3.87 | 7.56 | 3.12 |
| KUVP 25472 | 62.29 |  | 3.96 | - | 6.77 | 2.19 |
| KUVP 119673 right | 61.51 | 3.48 | 4.51 | - | 6.23 | 2.02 |
| KUVP 119673 left | - | - | 4.699* | - | - | - |
| YPM 1741 | 71.45 | 4.68 | 4.70 | 4.26 | 7.30 | 2.59 |

As described by Clarke (2004), the proximal end of the radius shows an ovoid humeral cotyle. Though Clarke describes this cotyle as slightly concave, it is completely flat in all the studied specimens save for YPM 1741. The ulnar facet is shallow and fairly indistinct, and a moderately developed and robust tubercle extends anteriorly opposite of the ulnar facet. A slight ridge extends distally from the tubercle, delimiting an ovoid fossa. This fossa was described by Clarke (2004) as equivalent to the very marked bicipital process of the ulna. Clarke noted a second conspicuous scar extending proximodistally from the tubercle; this is not visible in any of the studied specimens. Minor breakage in the shaft distal to the tubercle might have obscured this region.

The radial shaft is essentially straight, showing less curvature in anterior or posterior views than the YPM specimens. A clear and marked intermuscular line or ridge extends along most of the ventral surface of the radial shaft to the distal end of the bone. Clarke (2004) identified this structure and extensively discussed the possible muscular attachment points in this region.

The anterodorsal distal end of the radius shows a very conspicuous and marked tendinal groove (sulcus tendineus; Baumel & Witmer, 1993). The edge of the groove forms a ridge that runs across the ventral margin of the distal end of the radius towards the shaft.

Clarke (2004) described a very marked and prominent ovoid scar found on the posterior surface of the distal radius, in the middle of the ligamentous depression (depression ligamentosa; Baumel & Witmer, 1993). This scar is present in all of the studied specimens preserving the distal portion of the bone except for KUVP 25472, although it is very faint in KUVP 119673 and BHI 6420. This scar constitutes one of the autapomorphies of *Ichthyornis dispar*, although its absence in KUVP 25472 together with variation in its distinctness among the rest of the specimens may cast doubt on the extent to which it is truly diagnostic for *Ichthyornis*. A second, less-marked ovoid scar is found ventral to the first scar, at the base of a slightly developed posterior process, identified by Clarke (2004) as the ligamental process (tuberculum aponeurosis ventralis; Baumel & Witmer, 1993). The articular surface for the radial carpal is found dorsal to this process.

### ULNAR CARPAL

Three specimens included in this study contain the ulnar carpal bone: FHSM VP-18702 preserves the right ulnar carpal, ALMNH PV93.20.5 preserves both the left and right ulnar carpals, and KUVP 5969 preserves the left ulnar carpal. Only the ulnar carpal of SMM 2503 was previously known for *Ichthyornis*, though it was missing a portion of its dorsal ramus (crus breve; Baumel & Witmer, 1993) (Clarke, 2004). The condition of the ulnar carpal bones of both FHSM VP-18702 and ALMNH PV93.20.5 is exceptional, whereas the ulnar carpal of KUVP 5969 is missing its entire dorsal ramus. The morphology of the ulnar carpal bone in the new specimens is essentially identical to that described by Clarke (2004), with the exception of KUVP 5969.

The ulnar carpal is somewhat claw shaped in anterior view (Fig. 19), with a short, flattened dorsal ramus and a longer, more robust ventral ramus. The metacarpal incisure (incisura metacarpalis; Baumel & Witmer, 1993) is shallow and rounded. The articulation facet for the ulna is flat and subtriangular in dorsal view. Clarke (2004) described the tip of the dorsal ramus of SMM 2503 as missing, but the morphology of this structure seems to be essentially identical in both FHSM VP-18702 and ALMNH PV93.20.5, in which the region is slightly eroded, but without obvious signs of breakage.

**Figure 19.**
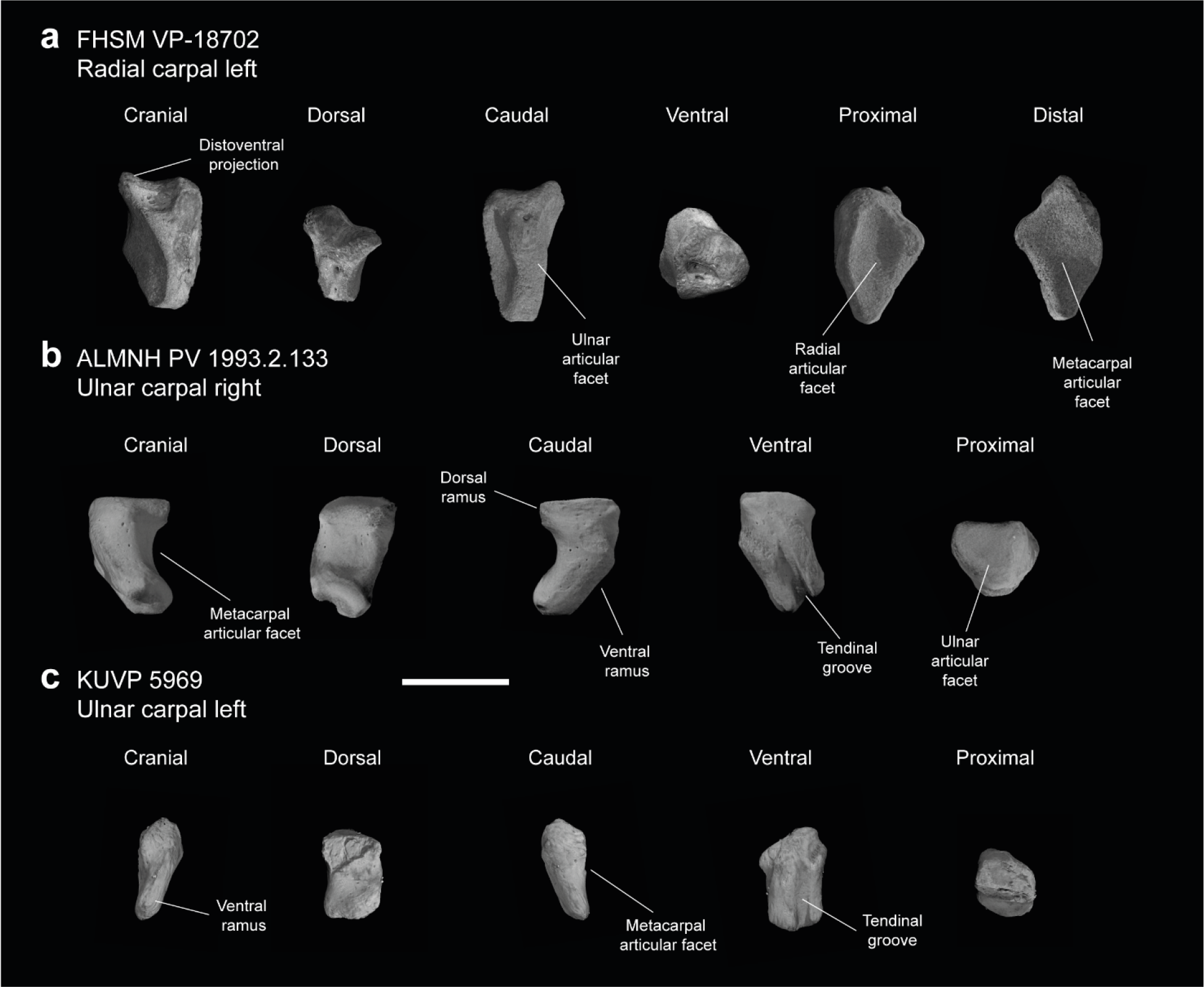
Free carpal bones of *Ichthyornis*. (a) FHSM VP-18702 left radial carpal, in cranial, dorsal, caudal, ventral, proximal and distal views; (b) ALMNH PV 1993.2.133 right ulnar carpal and (c) KUVP 5969 left ulnar carpal in cranial, dorsal, caudal, ventral and proximal views. Scale bar equals 5 mm.

The ventral ramus (crus longum; Baumel & Witmer, 1993) is elongate and recurved in FHSM VP-18702 and ALMNH PV93.20.5, with a markedly concave cranial surface and a moderately convex caudal surface, and becomes slightly wider at its ventral end. In contrast, the ventral ramus seems to be essentially straight in KUVP 5969, with both its dorsal and ventral ends on the same axis. This specimen exhibits flat cranial and caudal surfaces and lacks the ventral expansion present in the other specimens, instead slightly tapering ventrally. The ventral ramus of all three specimens preserves a deep and marked tendinal groove running across most of its flat caudal surface (Fig. 19). This tendinal groove originates at the base of the dorsal ramus of the ulnar carpal and extends to the midpoint of the ventral surface of the bone, at the articulation facet for the carpometacarpus. The groove is moderately proximodistally curved in FHSM VP-18702 and ALMNH PV93.20.5, but is completely straight in KUVP 5969. A round shallow pit is found on the cranial surface of the distalmost portion of the ventral ramus in both ALMNH PV93.20.5 radial carpals, but it seems to be absent in FHSM VP-18702 and KUVP 5969. The articular facet for the carpometacarpus, found on the cranial surface of the ventral ramus, is rounded with a moderately marked cranioventral depression in the two complete specimens; this depression is missing in KUVP 5969, in which this region is essentially flat.

The morphology of the ulnar carpal is poorly preserved in most Mesozoic avialans, preventing detailed comparisons, although it is very similar in *Iaceornis*. However, the dorsal ramus in *Iaceornis* is slightly reduced and its ventral ramus is moderately straighter than in *Ichthyornis*.

### RADIAL CARPAL

Following Clarke (2004), no radial carpal bones have been attributed to *Ichthyornis*, as YPM 1734, which bears a radial carpal, was reassigned as the holotype of *Iaceornis marshi* (Clarke, 2004). Two of the specimens studied here include exceptionally preserved radial carpals; BHI 6420 preserves the right radial carpal and FHSM VP-18702 preserves both the right and left elements. The morphology of the radial carpal is congruent in both specimens, and fills a gap in our knowledge of carpal anatomy among crownward stem birds, in which the radial carpal is rarely well-preserved or described.

All of the preserved radial carpals exhibit a dorsoventrally elongate shape, and are relatively compressed proximodistally. Their ventral surfaces are flat and concave, with a moderately developed distoventral projection. The elongated dorsal region of the radial carpal ends in a narrow tip. The flattened distal surface of the bone is rhomboidal in shape and slightly concave (Fig. 19), forming the articular surface for the carpometacarpus (facies articularis metacarpalis; Baumel & Witmer, 1993). This articular surface is delimited on its cranial side by a marked ridge running dorsoventrally from the dorsal end of the radial carpal to the tip of the distoventral projection. The proximal surface of the bone is a shallow ovoid depression (Fig. 19a), forming the articular surface for the radius (facies articularis radialis; Baumel & Witmer, 1993).

In caudal view, the radial carpal exhibits a wide and mostly flat ridge running dorsoventrally, delimiting on one side the proximal articular surface for the radius and the articular surface for the carpometacarpus on the distal side (Fig. 19a). This ridge continues into a process around the midpoint of its length, with two well marked foramina on the ventral side of this process. The ridge becomes flat and slightly concave dorsal to the process, corresponding to the articular surface for the ulna (facies articularis ulnaris; Baumel & Witmer, 1993).

A relatively wide and shallow sulcus for the tendon of m. ulnometacarpalis ventralis is found at the centre of the ventral surface of the radial carpal (Fig. 19a). The cranial end of this sulcus forms a subtle ridge running proximodistally from the base of the distoventral process to the proximal end of the sulcus. This ridge delimits a subtriangular depression with a considerable number of foramina in its cranial side. The more proximal region of the cranial side of the radial carpal preserves a flat, elevated and moderately concave surface, which roughly corresponds to the sulcus for the tendon of m. extensor carpi radialis (Mayr, 2014).

This sulcus runs dorsoventrally and maintains the same width for most of its length, enlarging slightly at its proximoventral end.

The morphology of the radial carpal has not been thoroughly described for most comparable fossil taxa, which makes comparisons with *Ichthyornis* difficult, but the overall proportions and morphology seen in *Ichthyornis* are quite similar to those of *Iaceornis* (Clarke, 2004) and *Presbyornis* (pers. obs.), suggesting that a comparable shape may have characterized the radial carpal of the ancestral neornithine. The radial carpal morphology varies greatly amongst extant crown birds (Mayr, 2014), but the morphology in *Ichthyornis* resembles most closely that of Anatidae and Pandionidae: these taxa exhibit a generally comparable elongate shape, together with the presence of a short but distinct distoventral projection and a rounded, concave, and porous cranial surface.

### CARPOMETACARPUS

Seven of the newly studied specimens preserve carpometacarpi, but the complete element is only represented in BHI 6420. FHSM VP-18702, KUVP 123459, 25469 and ALMNH PV93.20.5 preserve both the proximal and distal ends of the carpometacarpus but are missing part or most of the metacarpal shafts. MSC 34426 and RMM 3394 preserve only the distal portion of the element. The carpometacarpus of YPM 1724, already described by Clarke (2004) and included in this study, remains the most complete and best preserved carpometacarpus known for *Ichthyornis*. Measurements of the carpometacrpus of the specimens included in this study are provided in Table 9.

**Table 9.** Measurements of the carpometacarpus of *Ichthyornis* specimens. Proximal craniocaudal width is measured as the maximum craniocaudal extension of the proximal carpometacarpus, including the extensor process of metacarpal I. Proximal dorsoventral width corresponds to the maximum dorsoventral extension of the proximal carpometacarpus, including the pisiform process. Asterisks (*) denote measurements that might be unreliable due to breakage or distortion, but are included for completeness purposes. All measurements are in mm. -= not measurable.

| Specimen | Total length | Prox. Craniocaudal width | Prox. Dorsoventral width | Dist. Craniocaudal width | Dist. Dorsoventral width |
| --- | --- | --- | --- | --- | --- |
| ALMNH PV 1993.2.133 | - | - | - | 6.41 | 4.95 |
| BHI 6420 | 38.54 | 7.88 | 4.47* | 6.51 | 4.73 |
| FHSM VP 18702 | 36.03 | 7.24 | 6.29* | 6.55 | 4.42 |
| KUVP 25469 | 39.04* | 10.25 | 6.25 | 7.22 | 5.44 |
| KUVP 123459 | - | 9.63 | - | 6.05 | 4.74 |
| MSC 34426 | - | - | - | 6.60 | 4.39 |
| RMM 6202 | 28.21* | 9.98 | 6.94 | 6.87 | 4.74 |
| YPM 1742 | 39.58 | 10.99 | 5.18 | 6.56 | 5.24 |

The carpometacarpus in *Ichthyornis* is a proportionally long element, extending for about half of the length of the humerus or the ulna. Out of all the carpometacarpi studied, FHSM VP-18702 stands out as being shorter than other similarly proportioned specimens such as BHI 6420 and YPM 1724, representing only 87% of their maximum length. The FHSM VP-18702 carpometacarpus is divided into two proximal and distal fragments that seem to match perfectly, thus it seems that no part of its length is missing. Clarke (2004) reported certain carpometacarpi of differing lengths despite overall similar proportions, such as the relatively short carpometacarpus of YPM 1755 compared with the similarly proportioned but longer one of YPM 1773. but none of the other YPM specimens preserved enough material for further comparison. Comparison of BHI 6420 and FHSM VP-18702 reveals virtually equal lengths for several of their skeletal elements, such as the coracoid, the scapula, and the major manual digit phalanx (Tables 4, 5 and 10), and both specimens are otherwise comparable in their morphology and estimated mass (Fig. 30), seemingly confirming substantial variation in carpometacarpus length within *Ichthyornis*. Future work might clarify whether this disparity corresponds to intraspecific or interspecific variation.

**Table 10.** Measurements of the manual phalanx II:1 of *Ichthyornis* specimens. Total length and distal craniocaudal width measurements include the internal index process, when preserved. Asterisks (*) denote measurements that might be unreliable due to breakage or distortion, but are included for completeness. All measurements are in mm. - = not measurable.

| Specimen | Total length | Prox. Craniocaudal width | Prox. Dorsoventral width | Dist. Craniocaudal width | Dist. Dorsoventral width | Max. Craniocaudal width |
| --- | --- | --- | --- | --- | --- | --- |
| ALMNH PV<br>1993.2.133 | 22.03 | 5.96 | 4.80 | 5.60 | 3.37 | 7.32 |
| BHI 6420 | 23.81 | 4.99 | 2.98* | 6.41 | 3.09 | 8.11 |
| BHI 6421 | 20.93 | 3.96 | 4.91 | 5.80 | 3.95 | 6.90 |
| FHSM VP<br>18702 | 23.44* | 5.45* | 5.08 | 5.17* | 3.42* | 6.97 |
| KUVP<br>5969 | 15.63 | 3.33 | 2.74 | 3.39 | 2.36 | 4.91 |
| RMM<br>6201 | 19.97* | 4.36 | 4.36 | 4.97* | 3.74 | 7.08 |

As described by Clarke (2004), both the proximal and distal articular surfaces of the carpometacarpus bear a large number of small foramina, which can be observed in all the studied specimens except for FHSM VP-18702, although their absence in this specimen might be preservational. The carpal trochlea (trochlea carpalis; Baumel & Witmer, 1993) is large and well developed, and is flat in proximal view in all specimens. No groove is present between the lateral and medial condyles. Both condyles share a similar caudal extension and extend parallel to each other, contrary to the condition widespread among crown birds in which the medial condyle extends further caudally (Livezey & Zusi, 2006). Therefore, no distinct ulnocarpal articular facet is developed on the dorsal surface of the medial condyle, a condition roughly comparable to that of *Chauna*. The cranial surface of the proximal carpometacarpus is mostly flat, and the supratrochlear fossa (fossa supratrochlearis; Baumel & Witmer, 1993) is shallow and only moderately developed. The attachment point for the ligament ulnocarpo-metacarpale dorsale (Baumel & Raikow, 1993; Watanabe et al., 2021) is present just cranial to the supratrochlear fossa, visible as a shallow, rounded and faint depression in YPM 1724 and KUVP 25472, and much more deeply excavated and distinct in KUVP 25469. A rounded, pit-like scar is situated on the cranial edge of the lateral condyle just proximal to this depression in all the studied specimens, congruent with the attachment point for the m. ulnocarpalis dorsalis (Watanabe et al., 2021).

The pisiform process is situated at approximately the same level as the proximal end of the extensor process, whereas it is usually more distally situated in crown group birds (Clarke, 2004). It is short and robust, projecting slightly cranioventrally and overhanging slightly over the cranial carpal fovea, but not as far as in the surveyed charadriiforms. The pisiform appears to be proportionally larger in FHSM VP-18702, although the surrounding region is heavily distorted, which complicates an accurate comparison. The caudoproximal surface of the pisiform process is flat in all the studied specimens except FHSM VP-18702 and KUVP 25469, in which a minute and shallow fovea for the aponeurosis ventralis (Baumel & Raikow, 1993) is developed. A faint ridge extending distally from the pisiform process and reaching metacarpal III is visible in YPM 1724 and especially in KUVP 25469, but is not preserved in the rest of the specimens. A shallow oval scar is situated caudal to this ridge only in KUVP 25469, congruent with the attachment point for the ligament ulnocarpo-metacarpale ventrale (Watanabe et al., 2021). The infratrochlear fossa (fossa infratrochlearis; Baumel & Witmer, 1993) is shallow and poorly defined, as is the cranial carpal fovea.

Metacarpal I extends distally up to the distal extension of the proximal metacarpal symphysis as described by Clarke (2004) in YPM 1724, FHSM VP-18702, and KUVP 25469, although it seems to end slightly more proximally in KUVP 25472 and ALMNH PV93.20.5. The extensor process is subtriangular, and the cranial edge of metacarpal I is straight, running proximodistally between the cranialmost extension of the extensor process and the alular facet, subparallel to the main axis of the carpometacarpus. This is in contrast to most surveyed crown birds, in which the extensor process is significantly more cranially projected than the alular facet and the cranial edge of metacarpal I is positioned diagonal to the main axis of the carpometacarpus, as in *Puffinus lherminieri* or *Sterna hirundo*. The cranial edge of the process is thickened and the ventral surface is slightly concave, although it does not develop into a large concavity continuous with the cranial carpal fovea (fovea carpalis cranialis; Baumel & Witmer, 1993) as in Galliformes. In KUVP 25462 the ventral surface of the extensor process shows a large, round and deep depression (Fig 20); it is unclear if this is taphonomic in origin, although the surrounding proximal carpometacarpus is well-preserved. A similar rounded depression is present in this region in *Iaceornis*. The articular surface for the alular digit is flat and shelf-like, and is moderately dorsoventrally expanded, as in *Sterna hirundo*.

**Figure 20.**
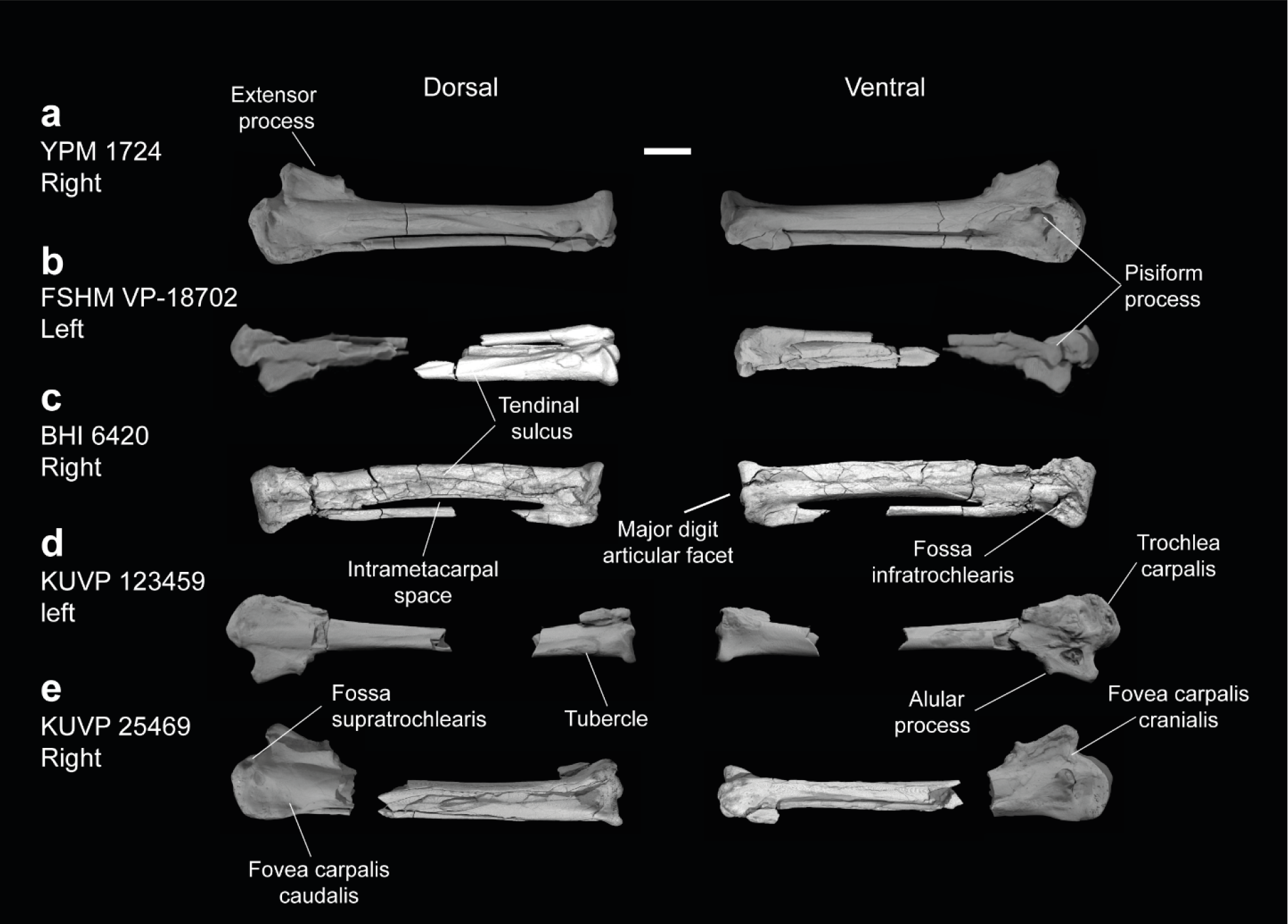
Carpometacarpi of *Ichthyornis*. (a) YPM 1724, (b) FHSM VP-18702, (c) BHI 6420, (d) KUVP 123459 and (e) KUVP 25469, in dorsal and ventral views. Scale bar equals 5 mm.

Metacarpal II is long, robust, and circular in cross-section. A flat scar is present on its proximal dorsocaudal surface in both YPM 1724 and KUVP 25469, starting in a position equivalent to the alular articular surface on the cranial surface. This scar was interpreted by Clarke (2004) as equivalent to the intermetacarpal process where developed, serving as an insertion point for the m. extensor metacarpi ulnaris. This scar runs parallel to the tendinal groove (sulcus tendineus; Baumel & Witmer, 1993) until the midpoint of metacarpal II. The scar appears to be entirely absent in all other studied specimens. The tendinal groove is wide and marked; it runs proximodistally across the cranial surface of metacarpal II until its midpoint, after which it is deflected caudally, wrapping around metacarpal II, in a similar manner to that of *Sterna hirundo*.

Metacarpal III is significantly reduced in both craniocaudal and dorsoventral diameter compared with metacarpal II, and it is mostly straight and parallel to it, defining an extremely narrow intermetacarpal space. Although most crownward Mesozoic euornitheans share an extremely reduced intermetacarpal space, the condition in *Ichthyornis* is narrower than in comparable taxa such as *Gansus* (Wang et al., 2016b) and *Iaceornis* (Clarke, 2004). Several crown bird groups with an elongated carpometacarpus also exhibit a straight metacarpal III, such as *Puffinus lherminieri*, *Sterna hirundo*, and *Presbyornis pervetus*, but such a reduced intermetacarpal space is uncommon, restricted mostly to ecologically specialized taxa such as Sphenisciformes and Gaviiformes. Metacarpal III is slightly craniocaudally compressed in its proximal region, but becomes progressively more circular in cross-section along its length. A tear-shaped and moderately ventrally developed tuberosity is visible on the ventral surface of metacarpal III, just proximal to the distal end of the proximal metacarpal symphysis, observable in YPM 1724 and KUVP 25469 and only faintly visible in FHSM VP-18702. A similar tuberosity appears to be present in *Iaceornis*, and is variably present among crown bird lineages, including *Crypturellus variegatus* and, combined with a shallow scar, *Sterna hirundo* and *Scolopax rusticola*, in which it is associated with a second attachment point for the ligament ulnocarpo-metacarpale ventrale (Watanabe et al., 2021).

Metacarpals II and III are almost equal in their distal extent, and the intermetacarpal symphysis is flat to moderately concave in cranial view. The dorsodistal terminus of metacarpal II defines a large and oval-shaped flat distal surface for the articulation of the major manual digit. This region bears a large and rounded dorsal process.

### PHALANX II-1

Five of the studied specimens preserve the first phalanx from the second digit, most of them in excellent condition: FHSM VP-18702 (left), BHI 6420 (left and right), BHI 6421 (right), ALMNH PV93.20.5 (right), KUVP 5969 (right) and RMM 6201 (right). These show a clear size distribution, with KUVP 5969, the smallest specimen, being only ∼66% of the length of the largest specimens, BHI 6420 and FHSM VP-18702 (Table 10).

The phalanx is elongate, and around 60% of the length of the carpometacarpus (Table 9). The element is dorsoventrally flattened and caudally expanded. The proximal articular surface is roughly quadrangular in proximal view, with a large, marked concave depression on its caudoventral margin for the major digit articular facet of the carpometacarpus. A robust and well-developed caudoproximally-directed process bounds this depression cranially. The development of this process varies among the studied specimens and apparently exhibits negative allometry: it is very short in FHSM VP-18702 and BHI 6421 and proportionally longer in ALMNH PV93.20.5 and RMM 6201 (Fig. 21). Clarke (2004) hypothesized that this process could represent a synapomorphy of *Ichthyornis dispar* + Neornithes, but this feature seems to be obscured in most comparable Mesozoic euornitheans, so this hypothesis cannot be properly assessed at present. A process comparable to that of *Ichthyornis* is widespread among extant birds, with a similar extension observable in, for example, *Gallus gallus* and *Stercorarius antarctica*.

**Figure 21.**
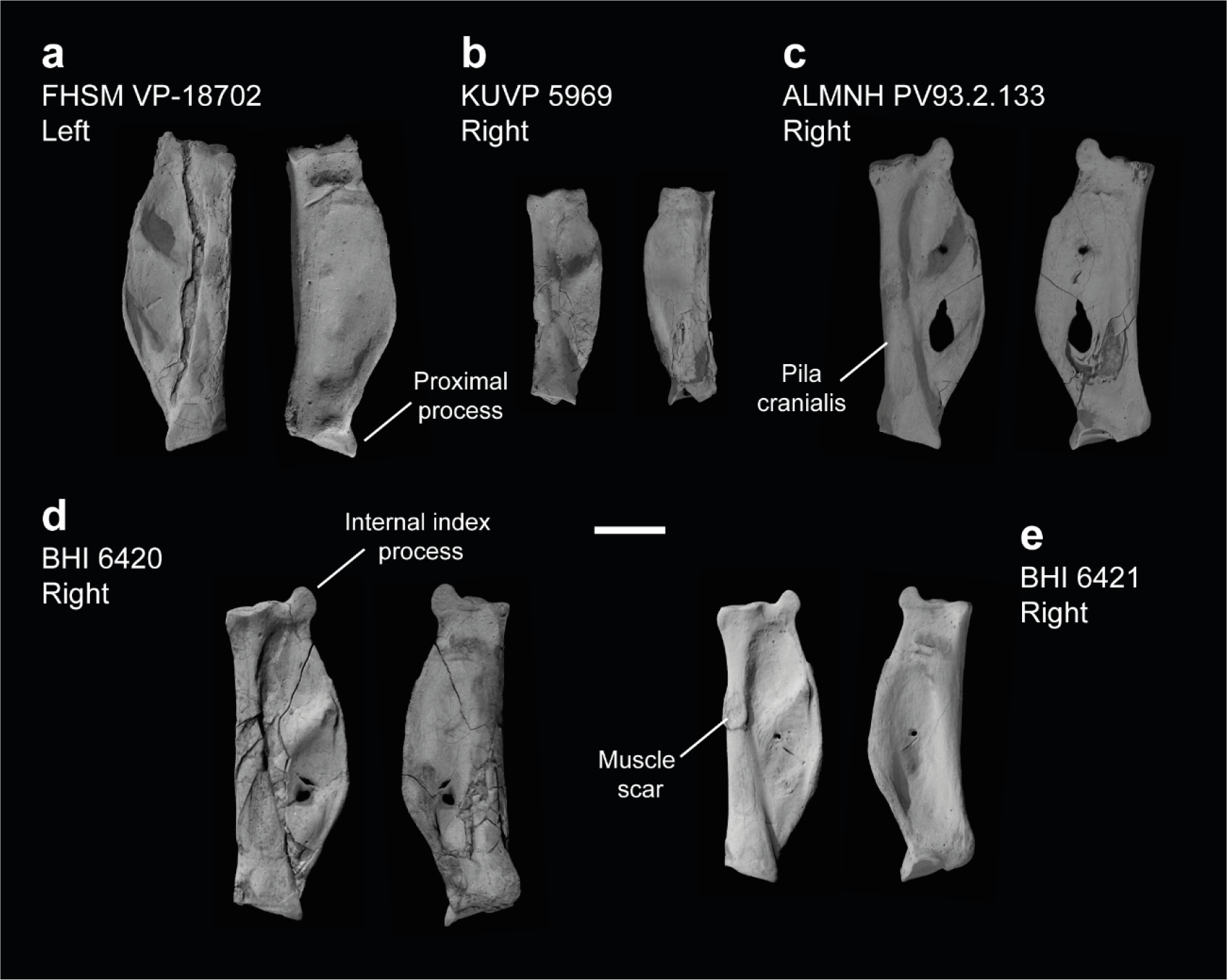
Manual phalanx II-1 of *Ichthyornis*. (a) FHSM VP-18702 left phalanx, (b) KUVP 5969 right phalanx, (c) ALMNH PV93.2.133 right phalanx, (b) BHI 6420 right phalanx and (e) BHI 6421 right phalanx, in dorsal (left) and ventral (right) views. Scale bar equals 5 mm.

A large tubercle is present just distal to the proximal articular surface on the cranioventral surface of the phalanx, likely for the distal implantation of the m. abductor digiti majoris (Baumel & Raikow, 1993). This tubercle extends into a marked ridge that delimits the ventral side of the cranial pillar (pila cranialis; Baumel & Witmer, 1993). The cranial pillar is robust and dorsoventrally wide, with a mostly flat cranial surface and a rounded dorsal surface. A very large oval muscle scar is present on the dorsal surface of the pila between 50% (ALMNH PV93.20.5) and 55% (BHI 6421) of its distal extent in all specimens except for KUVP 5969, in which a very shallow impression is only faintly observable. The craniocaudal width of the scar is approximately equal to the width of the dorsal surface of the cranial pillar in all specimens except BHI 6421, in which it is wider than the pillar. The proximal and cranial margins of the scar are raised, and its dorsal surface is flat. A similar scar has apparently only been described in *Yixianornis* (Clarke et al., 2006) among Mesozoic euornitheans, and no equivalent scar is present in this region for any surveyed crown bird.

The dorsal surface of the phalanx caudal to the cranial pillar shows two large rounded depressions, with the proximal depression larger than the distal one. These depressions, present in most extant birds examined, are not perforated, though the bone becomes very thin at their centre and both BHI 6420 and ALMNH PV93.20.5 show breakage in this region. The proximal depression shows a large foramen on its distal edge in several specimens, which penetrates the bone only in BHI 6421. Both depressions are separated by a wide caudodistally directed ridge (pila obliqua fossae; Livezey and Zusi, 2006) extending from a position roughly equivalent to the aforementioned scar, although this varies among the different specimens, and is situated distinctly proximal to the scar in BHI 6421. The caudal termination of this ridge is dorsally raised, with a marked groove on its caudal surface, likely for the passage of the m. interosseus ventralis tendon (Baumel & Raikow, 1993; Vazquez, 1995).

The ventral surface of the phalanx is mostly flat, showing a shallow and large depression extending proximodistally along most of its surface. A shallow and narrow groove is found on the distal portion of the ventral surface, running craniocaudally and reaching the caudal margin of the phalanx. This groove is mostly straight and horizontal in most specimens, but shows pronounced cranial curvature in ALMNH PV93.20.5 and distal curvature at its midpoint in FHSM VP-18702. The caudal edge of the phalanx forms a wide curve between the proximal and the distal articular surfaces, but in KUVP 5969 and, to a lesser degree, in FHSM VP-18702, this curve ends abruptly at around 90% of the phalanx length, distal to which the caudal edge becomes straight, similar to the condition in *Iaceornis marshi* (Clarke, 2004). In both specimens, this point is equivalent to the point where the ventral craniocaudal groove reaches the caudal edge of the phalanx. The caudal edge of the phalanx is slightly widened craniocaudally and dorsally recurved, forming a narrow ridge that expands around the midpoint of the caudal surface, where it meets the caudal end of the pila obliqua fossae.

The distal articular surface is dorsoventrally expanded with respect to the cranial pillar, and mostly flat, with a moderately developed condyle for phalanx II-2 in its centre. A short tubercle is present just proximal to the craniodorsal edge of the articular surface. A smaller but better-defined tubercle is present on the cranioventral edge of the articular surface in FHSM VP-18702 and BHI 6421, though it is broken in all other specimens. This tubercle seems to serve as an attachment point for the m. flexor digitorum superficialis where present, as in *Stercorarius antarctica* (Watanabe et al., 2021). A large distal internal index process can be observed on the caudal edge of the bone in all specimens except KUVP 5969, though only the base of the process is preserved in most of them. The process is rounded and convex on its caudal side and slightly concave on its cranial margin, reaching its maximum distal projection close to the cranial margin. It varies in size among specimens, and is proportionally largest in ALMNH PV93.20.5, extending further distally than in the other specimens. In KUVP 5969, which preserves the smallest phalanx II-1, the process is not developed, and only a limited distal tubercle is present in its place, in a similar manner to the condition in *Iaceornis* or *Yixianornis*. The presence of an internal index process has been considered one of the autapomorphies of *Ichthyornis dispar* (Clarke, 2004). Its absence in KUVP 5959 could be due to ontogenetic factors (e.g., if this specimen—the second smallest of those surveyed—is ontogenetically immature, perhaps the process had yet to develop). Although the ontogenetic development of the internal index process has not been previously investigated, we verified that it is absent in a *Macronectes giganteus* chick, but present and well developed in adults, lending further support to the identification of KUVP 5969 as a possible juvenile individual.

Alternatively, the lack of an internal index process in KUVP 5969 may indicate that this specimen represents a previously unrecognized taxon of small ornithurine from the Niobrara Formation. An internal index process is unique to *Ichthyornis* among known Mesozoic avialans, but its distribution is widespread within Neoaves, being present at least in Strisores, Columbiformes, Pterocliformes, Otididae, Gruidae, Charadriiformes, Procellariiformes, Suliformes, Pelecanidae, and Psittaciformes (Stegmann 1963, 1978). This raises the question of whether the process may represent a crown bird symplesiomorphy, or whether its presence among some extant neoavians represents the convergent acquisition of an *Ichthyornis*-like morphology in this extant clade. The large and rounded process in *Ichthyornis* is particularly reminiscent of the morphology exhibited by certain extant representatives of Charadriiformes, such as *Glareola pratincola* and *Sterna hirundo,* as well as certain Procellariiformes like *Puffinus lherminieri*.

### PHALANX II-2

The second phalanx of the major digit is preserved in three of the studied specimens, BHI 6420, KUVP 25469 and ALMNH PV93.20.5, all from the right manus. All three specimens are in exceptionally good condition, preserving clear muscle attachments not previously described for Mesozoic euornitheans, but only KUVP 25469 preserves the distalmost portion of the element (Fig. 22a). The phalanx is an elongate, narrow, and straight element with strongly craniocaudally expanded proximal and distal articular surfaces. The length of this phalanx is approximately equal to that of the proximal phalanx (Table 11). Both the cranial and the ventral surfaces of the shaft are deeply excavated.

**Figure 22.**
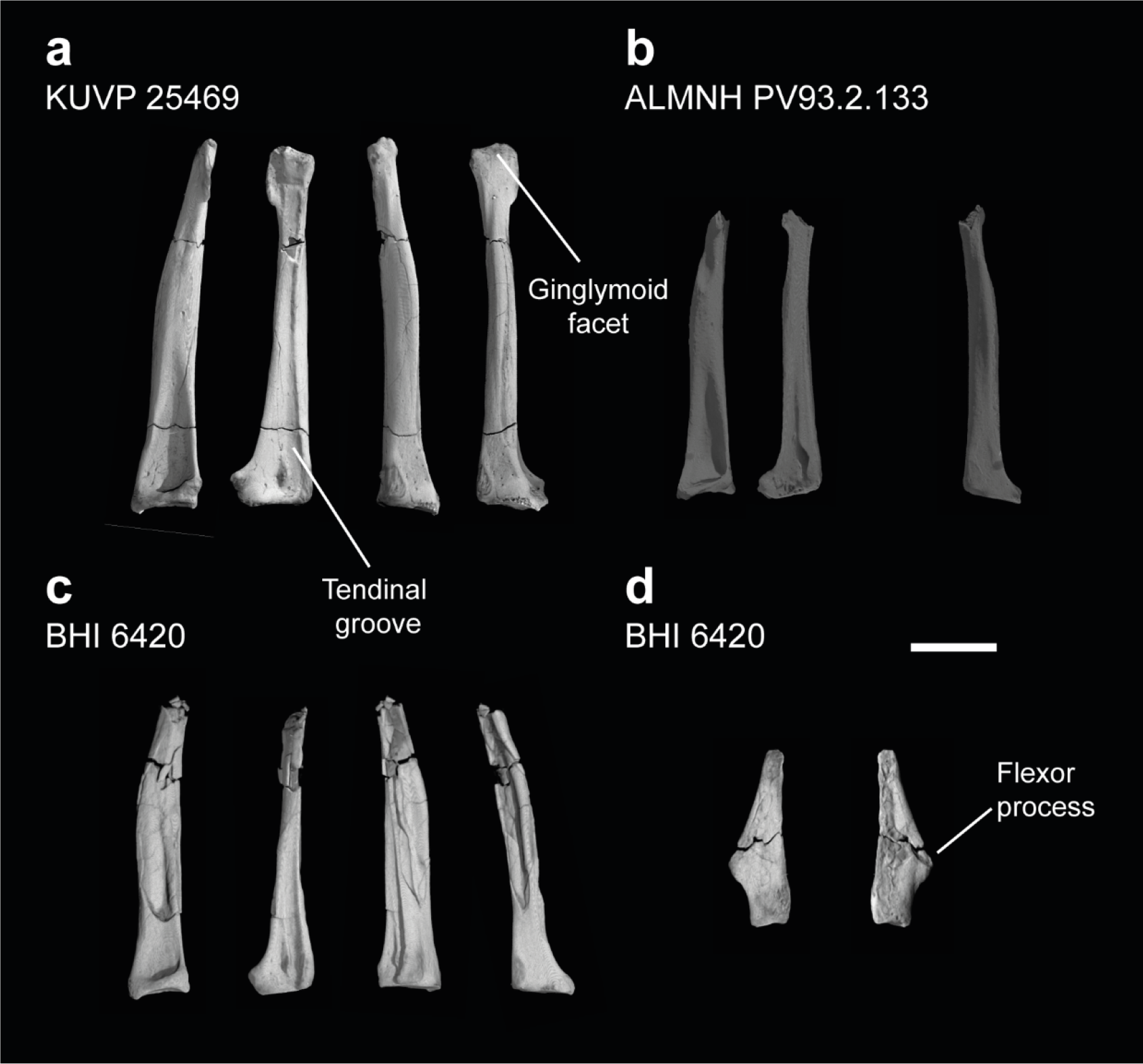
Manual phalanges II-2 and III-1 of *Ichthyornis*. (a) KUVP 25469 right II-2 phalanx, (b) ALMNH PV93.2.133 right II-2 phalanx and (c) BHI 6420 right II-2 phalanx in caudal, ventral, dorsal and cranial views; and (d) III-1 phalanx in dorsal and ventral views. Scale bar equals 5 mm.

**Table 11.** Measurements of the manual phalanx II:2 of *Ichthyornis* specimens. Asterisks (*) denote measurements that might be unreliable due to breakage or distortion, but are included for completeness purposes. All measurements are in mm. - = not measurable.

| Specimen | Total length | Prox. Craniocaudal width | Prox. Dorsoventral width | Dist. Craniocaudal width |
| --- | --- | --- | --- | --- |
| ALMNH PV<br>1993.2.133 | 17.91* | 3.43 | 3.87 | - |
| BHI 6420 | 17.46* | 3.91 | 3.49 | - |
| KUVP 25469 | 22.39 | 4.90 | 3.98 | 2.56 |

The proximal articular surface is mostly flat and shelf-like, with a shallow depression on its cranial side that matches the tuberosity developed on the distal articular surface of the proximal phalanx. A moderately developed tubercle is present just distal to the articular surface on the cranial surface of the phalanx, extending distally into a shallow ridge that delimits the ventral edge of the cranial surface of the phalanx, but does not reach the distal articular surface (Fig. 22). This tubercle has a flat proximal surface and preserves a rough cranial texture in all studied specimens, and it most likely corresponds to the implantation scar for the m. extensor longus digiti majoris (Baumel & Raikow, 1993). A similar but considerably better developed tubercle is present at a comparable position on the caudal surface of the bone, just distal to the proximal articular surface. This tubercle extends into a thin and caudally expanded ridge that delimits the caudal surface of the phalanx, reaching the distal region of the phalanx but becoming progressively shallower, and it articulates exactly with the internal index process of the proximal phalanx.

The dorsal surface of the phalanx shows a large and deep excavation on its proximal end for the attachment of the leading primary feather (Hieronymus, 2016). This excavation decreases in depth distally, becoming flat and indistinct on the distal third of the phalanx, contrary to the condition in most crown-group birds, in which this concavity extends along the whole length of the phalanx (Hudson et al., 1969,1972; Vazquez, 1995; Hieronymus, 2015). The ventral surface of the phalanx shows a shallow depression on its proximal portion but becomes essentially flat distally, with the exception of a deeply excavated groove, which extends along the whole length of the ventral surface. The cranial edge of the groove is delimited by the ridge extending distally from the m. extensor longus digiti majoris tubercle, and caudally by a shallow ridge (fig. 22a). On the proximal end of the groove, this caudal ridge extends into a thin and delicate tubercle or flange present on all three specimens. A large and round tubercle is present in this region in Alcidae for the distal implantation of the m. flexor digitorum superficialis and profundis (Watanabe et al., 2021), and the presence of the ventral groove in *Ichthyornis* indicates that the tendons of at least some of these muscles extended along the whole ventral margin of the phalanx. If that was the case, the flange at the proximal end of the groove may represent the base of a retinaculum covering both tendons in *Ichthyornis*. While in most crown birds both tendons attach only on the proximal region of phalanx II-2, the m. flexor digitorum superficialis extends to the distal end of the phalanx in Tinamidae and Galliformes (Hudson et al., 1964,1972) and into the proximal region of phalanx II-3 in extant birds that retain it, such as Anseriformes (Zusi & Bentz, 1968) and juvenile *Opisthocomus* (Hudson et al., 1964). The tendinal groove is much more developed and deeply excavated in *Ichthyornis* than in either group, and it is open distally, indicating the probable presence of a large and functional ungual phalanx.

The distal region of the phalanx is dorsoventrally compressed and craniocaudally expanded. A large flange-like projection is present along the caudal edge of the distal end, with a slightly concave ventral surface, which likely housed the distal implantation of the m. interosseous ventralis (Baumel & Raikow, 1993). The distal articular surface is only weakly ginglymoid (*contra* Clarke, 2004, in which it is described as well developed), but evidences the presence of a third phalanx on digit II forming an ungual, which is not preserved in the studied specimens (Fig. 22a). A second manual ungual phalanx is well developed in several volant Mesozoic euornitheans such as *Yixianornis, Iteravis,* and *Gansus* (Clarke et al., 2006; Liu et al., 2014; Zhou et al., 2014; Wang et al., 2016b), which show similar distal expansions on phalanx II-2. A greatly reduced third phalanx is present in Lithornithidae (Houde, 1988; Nesbitt & Clarke, 2016) and certain extant bird lineages, as previously mentioned. Two shallow projections are present on both sides of the groove for either M. flexor digitorum superficialis or profundis. Both projections preserve deep pits on their distal surfaces, likely for the collateral ligaments of the third phalanx.

### PHALANX III-1

The single phalanx of the minor digit is preserved only in BHI 6420. This element was previously only known for YPM 1775, in which both preserved phalanges lacked their distal portions (Clarke, 2004). In contrast, the distal portion is mostly preserved in BHI 6420.

The phalanx is a robust and elongate element, approximately half the length of phalanx II-1, and is dorsoventrally compressed (Fig. 22d). The proximal surface of the phalanx is wide and wedge-like, with a moderately developed tuberosity on its cranioventral surface, similar to the morphology described for *Iteravis* (Zhou et al., 2014). This tuberosity likely articulated both with the minor metacarpal and the proximal phalanx of the major digit. The cranial margin of the phalanx is mostly straight, but the caudal margin shows a large and well-developed flange-like flexor process halfway along the length of the phalanx, giving the phalanx a roughly triangular shape. The flexor process shows a marked pit on its dorsal surface for the implantation of the m. flexor digiti minoris (Baumel & Raikow, 1993). The dorsal surface of the phalanx is mostly flat, while the ventral surface is moderately concave. The distal end of the phalanx is tapered, though the distalmost portion appears to be broken (Fig. 22d).

The morphology of phalanx III-1 in *Ichthyornis* is similar to that of certain other crownward ornithurines such as *Gansus* and *Iteravis* (Liu et al., 2014; Zhou et al., 2014; Wang et al., 2016b), but more elongate and with a more strongly projected flexor process than in *Yixianornis* (Clarke et al., 2006). The morphology of the YPM 1775 III-1 phalanx was compared to that of Tinamidae by Clarke (2004), but despite the presence of a comparable flexor process, the phalanx is much more elongate in *Ichthyornis,* which could not be appreciated from the partially preserved phalanges in YPM 1775. While similar morphologies are developed across multiple crown-bird lineages such as Tinamidae or Anseriformes, the condition in *Ichthyornis* appears most similar to that of certain Lithornithidae such as *Calciavis grandei* (Nesbitt & Clarke, 2016) or some extant taxa like *Columba livia* (Columbiformes) or *Rynchops flavirostris* (Charadriiformes).

### PELVIC GIRDLE

The complete pelvic girdle is preserved in KUVP 119673, which preserves the fused ilia, ischia and pubes from both the left and right sides, disarticulated from the synsacrum (Fig. 23). KUVP 25469 preserves portions of the preacetabular ilium in association with the synsacrum, but these are extremely fragmentary. Amongst the YPM material, only one specimen preserves a semi-complete pelvic girdle, YPM 1732, in articulation with the synsacrum. Despite the large number of complete and partial synsacra found amongst the studied specimens (see above), no other specimen beyond KUVP 25469 preserves any pelvic remains, evidencing the weak fusion between the pelvic girdle and the sacral vertebrae in all but the largest specimens. Both sides of the KUVP 119673 pelvis are mediolaterally flattened and include radiopaque inclusions, which hampers the identification of some minor features such as muscle attachment impressions. Despite this, both sides preserve the entire surfaces of all the elements, except for the distal right ischium. Measurements of KUVP 119673 pelvic elements are provided in Table 12.

**Figure 23.**
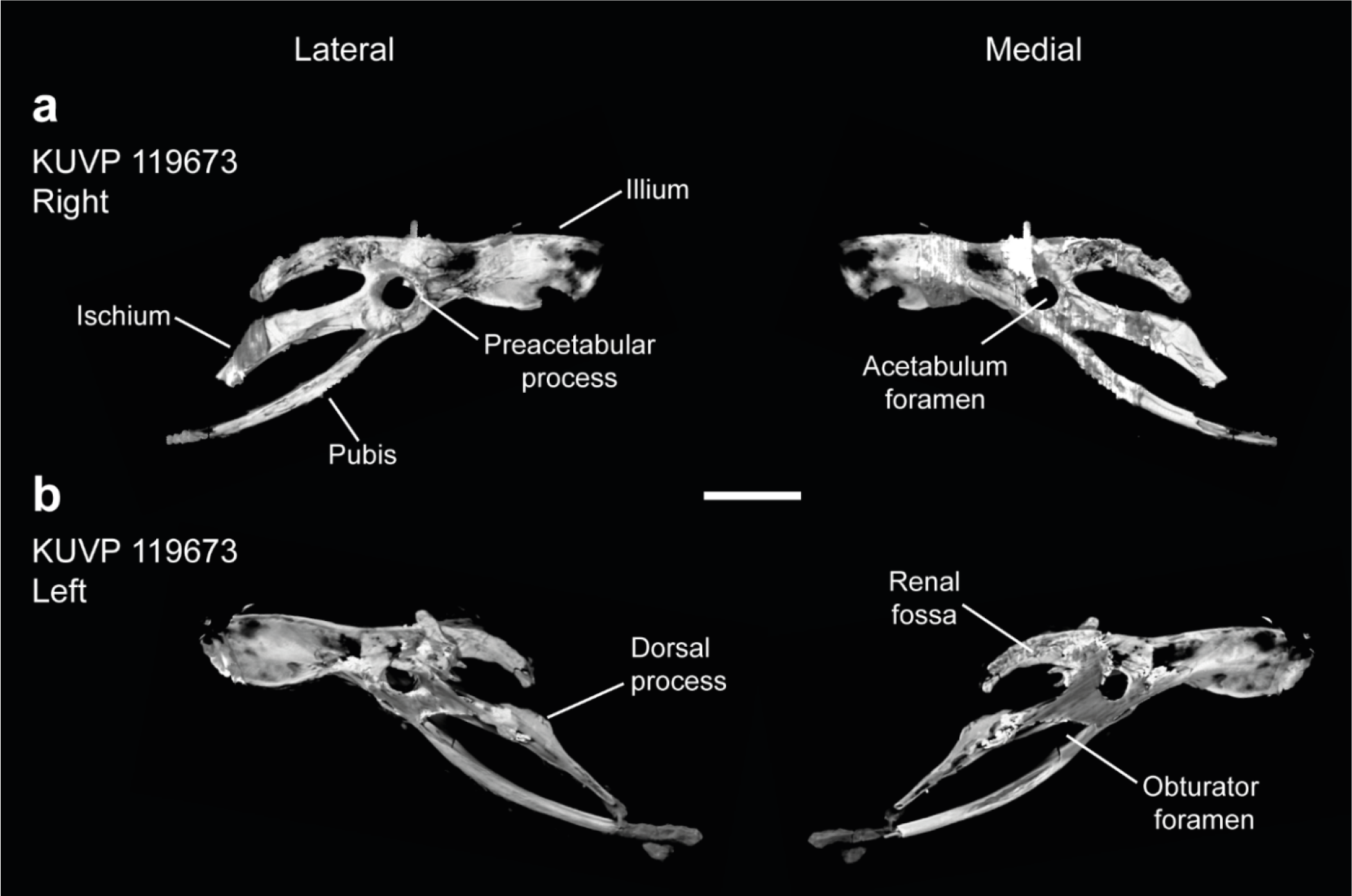
Pelves of *Ichthyornis* specimen KUVP 119673. (a) right pelvis and (b) left pelvis, in lateral and medial views. Scale bar equals 1 cm.

**Table 12.** Measurements of the pelvic elements of *Ichthyornis* specimens. Maximum length corresponds to the total craniocaudal length of the fused pelvic element. Ilium maximum height is measured as the maximum dorsoventral extension of the preacetabular iliac wing. Asterisks (*) denote measurements that might be unreliable due to breakage or distortion, but are included for completeness. All measurements are in mm. - = not measurable.

| Specimen | Max length | Ilium length | Ilium max height | Ischium length | Pubis length |
| --- | --- | --- | --- | --- | --- |
| KUVP 119673 Right | 56.27 | 35.04 | 7.63 | 25.03 | 34.40 |
| KUVP 119673 Left | 48.51* | 36.83 | 7.81 | 18.93* | 28.48* |

The ilium in YPM 1732 is missing most of its preacetabular and postacetabular portions, but the outline of these regions was illustrated by Marsh (1880). It is not clear whether this illustration was based on portions of the bones that have since been lost, or if the illustrations reflect hypothetical reconstructions (Clarke, 2004). Supporting the latter alternative, the iliac morphology preserved in KUVP 119673 differs considerably from that illustrated in Marsh (1880), particularly with regard to the shape of the postacetabular region. The preacetabular wing of the ilium is elongate, reaching toward the caudal end of the first sacral vertebra. The ilium shows rounded cranial and ventral margins and a mostly straight dorsomedial margin; it reaches its maximum dorsoventral height at around 40% of its preacetabular length measured from its rostral tip (Table 12), narrowing drastically just cranial to the acetabulum. No ossified tendons like those preserved in YPM 1732 are present on either side of the KUVP 119673 pelvis. The preacetabular lateral surface is strongly concave, with a shallow groove running craniocaudally through the centre of most of the lateral surface, defining a large attachment surface for the m. iliotrochantericus caudalis and cranialis, but no clear demarcation is visible between the attachment surfaces of both muscles. Both the dorsal and ventral margins of the preacetabular wing are thickened and slightly laterally recurved. The medial surface of the preacetabular wing is strongly convex, with a marked ridge, equivalent to the aforementioned lateral groove, running along most of its length. This ridge shows a moderate ventral curvature and forms the ventral side of the acetabulum where it meets the pubis, which seems continuous with this ridge medially. The acetabulum is large and circular, with its dorsal margin strongly thickened, extending into a moderately developed antitrochanteric process that does not show a clear lateral tip (Fig. 23).

The postacetabular ilium, which does not fuse with the ischium, is oriented dorsoventrally, *contra* Clarke (2004). Marsh (1880) illustrated a reconstructed laterally oriented and mediolaterally wide postacetabular ilium, but the illustration was noted by Clarke (2004) as differing from the preserved material. The postacetabular iliac wing in KUVP 119673 is short, reaching distally as far as the dorsal process of the ischium and being 70% of the length of the preacetabular wing. The postacetabular illium is dorsoventrally narrow, gently curving ventrally along its length, approaching but not fusing with the ischium, and defining an ovoid ilioischiadic space (foramen ilioischiadicum; Baumel & Witmer, 1993). The lateral surface of the postacetabular ilium is strongly convex, with a marked ridge running along its entire length, parallel to both the dorsal and ventral margins of the element. Conversely, the medial surface of the bone is strongly concave, defining a deeply excavated and elongated renal fossa (Livezey & Zusi, 2006). Given the reduced postacetabular iliac morphology, it is not clear whether the ilium contacted the caudal transverse processes of the synsacrum. In most other Mesozoic euornitheans, such as *Yixianornis* or *Yanornis* (Zhou & Zhang, 2001; Clarke et al., 2006), the postacetabular iliac wing is similarly dorsoventrally narrow, but in contrast to *Ichthyornis*, it does not show any significant dorsoventral curvature and it does not approach the dorsal process of the ischium. The morphology of the *Ichthyornis* ilium differs from that of *Yixianornis, Gansus* and *Iteravis* (Clarke et al., 2006; Liu et al., 2014; Zhou et al., 2014; Wang et al., 2016b), in which both pre- and postacetabular wings are of similar length, the preacetabular wing does not show significant variation in its dorsoventral width along its length, and the postacetabular wing is not markedly dorsoventrally recurved. The morphology in *Ichthyornis also* differs from that of Hesperornithes, in which the ilium has an extremely elongate and enlarged postacetabular wing (Bell & Chiappe, 2016, 2020). Conversely, the ilium of *Ichthyornis* is very similar to that of more stemward euornitheans like *Schizooura* (Zhou et al., 2012) and *Eogranivora* (Zheng et al., 2018), in which the postacetabular ilium is shortened (although more so in these taxa than in *Ichthyornis,* in which it extends for 50% of the length of the preacetabular wing), and the preacetabular wing is dorsoventrally constricted close to the acetabulum.

The ischium in *Ichthyornis* is extremely elongate, extending into a strap-like structure almost reaching the caudal end of the pubis (Fig. 23). The ischium shows a convex dorsal margin with a moderately developed and wide dorsal process near its midpoint. The dorsal process on both sides of KUVP 119673 is significantly shorter than the triangular and elongate flange-like extension visible in YPM 1732, but there are no signs of breakage. Its ventral margin is weakly concave, and no projection demarcating the obturator foramen area is visible, contrary to the condition described for YPM 1732 (Clarke, 2004). The caudal extension of the ischium is ventrally deflected and is strongly tapered, with its caudalmost extent, preserved only on the left side, almost reaching the pubis, but not completely closing the ischiopubic space. The lateral surface of the ischium is mostly flat, with a marked ridge extending from the antitrochanteric process and running across the whole length of the ischium. The ridge runs through the centre of the lateral surface proximally but becomes continuous with the ventral surface distally. An equivalent groove is developed on the medial surface, with a similar shape and extension to the lateral ridge. The ischium in *Ichthyornis* is essentially identical to that of *Gansus* and *Yixianornis* in its length and shape, with a similar extension of the dorsal process, which is shorter in *Iteravis*. The lateral ridge is present as well in most other crownward euornitheans, similarly extending along the whole length of the ischium in *Yixianornis* and *Iteravis*, but only reaching the midpoint of the ischium in *Gansus*.

The pubis is a robust and elongate rod-like element, and is longer than the entire craniocaudal length of the ilium. The pubis is strongly curved dorsoventrally, with its caudal tip pointing dorsally, and delimiting an almond-shaped ischiopubic space, with a very reduced caudal opening obscured by radiopaque inclusions on the left pelvis (Fig. 23b). This morphology of the ischiopubic space is similar to that of *Gansus* but contrasts with the condition in *Apsaravis*, in which the ischium and the pubis run parallel to each other and the space between both is greatly reduced. The pubis does not seem to taper along its length, although its caudalmost extent is not well preserved and it is unclear whether a moderate distal expansion existed as in *Iteravis* and *Gansus*. Since both pubes are separated and mediolaterally flattened, whether they exhibit any mediolateral curvature and contacted medially, as in *Yanornis, Yixianornis*, *Gansus* and *Iteravis*, cannot be directly observed. The pubes of YPM 1732 were illustrated by Marsh (1880) as not contacting medially, although since the only observable pubis of this specimen is missing most of its length this cannot be verified at present. As mentioned above, Marsh’s (1880) illustration was noted by Clarke (2004) to differ from the preserved material, and the morphology of the ilium illustrated by Marsh is strikingly different from that described in this study, casting additional doubt on Marsh’s interpretations of the pelvis. The lack of completely preserved pubic bones precludes inferences of the precise phylogenetic origins of the unfused pubic symphysis characteristic of crown birds, and, by extension, strong inferences regarding the maximum diameter of *Ichthyornis* eggs, despite earlier studies positing that *Ichthyornis* provides the earliest evidence of a fully open pubic symphysis homologous with that of crown birds (Mayr, 2017b). No preacetabular tubercle or pectineal process is developed on the cranial surface of the pubis, as noted by Clarke (2004), contrasting with the condition in Hesperornithes, in which this process is extremely well developed (Bell & Chiappe 2016, 2020).

### FEMUR

Five of the new specimens preserve femoral remains. FHSM VP-18702 preserves a complete but severely distorted and laterally flattened right femur, and beyond the three-dimensionally-preserved femoral head, no other features are discernible (Fig. 24c). KUVP 119673 preserves a craniocaudally flattened partial right femur, and most recognizable features apart from the general shape of the two distal condyles are observable. ALMNH PV93.20.5 preserves a complete and undistorted left femur exhibiting only minimal breakage, representing the best-preserved *Ichthyornis* femur known to date (Fig. 24a). ALMNH PV98.20.183.1 preserves the distal portion of the femur in three dimensions, although the surface of the bone is eroded, obscuring most features. ALMNH PV98.20.186 preserves only the proximal portion of the right femur, albeit in excellent condition. The morphology of the femur in the studied specimens is generally congruent with that described by Clarke (2004), although the exceptional quality of the new specimens allows for a more detailed description. Measurements of the femora of the specimens included in this study are provided in Table 13

**Figure 24.**
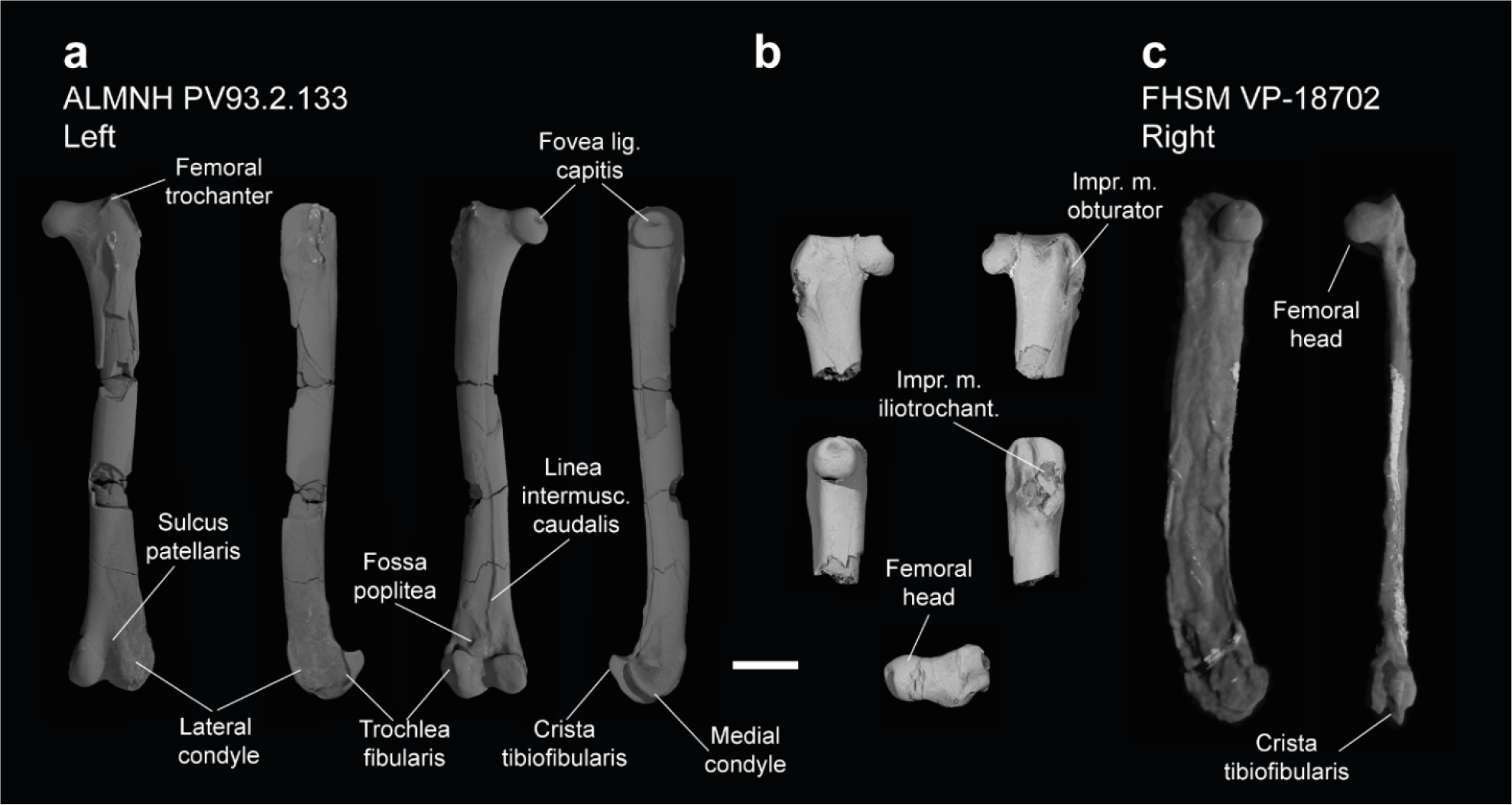
Femora of *Ichthyornis*. (a) ALMNH PV93.2.133 left femur in cranial, lateral, caudal and medial views; (b) proximal right femur of ALMNH PV98.20.186 in (clockwise order from upper left) cranial, caudal, lateral, dorsal and medial views and (c) right femur of FHSM VP-18702 in medial (left) and caudal (right) views. Scale bar equals 5 mm.

**Table 13.**
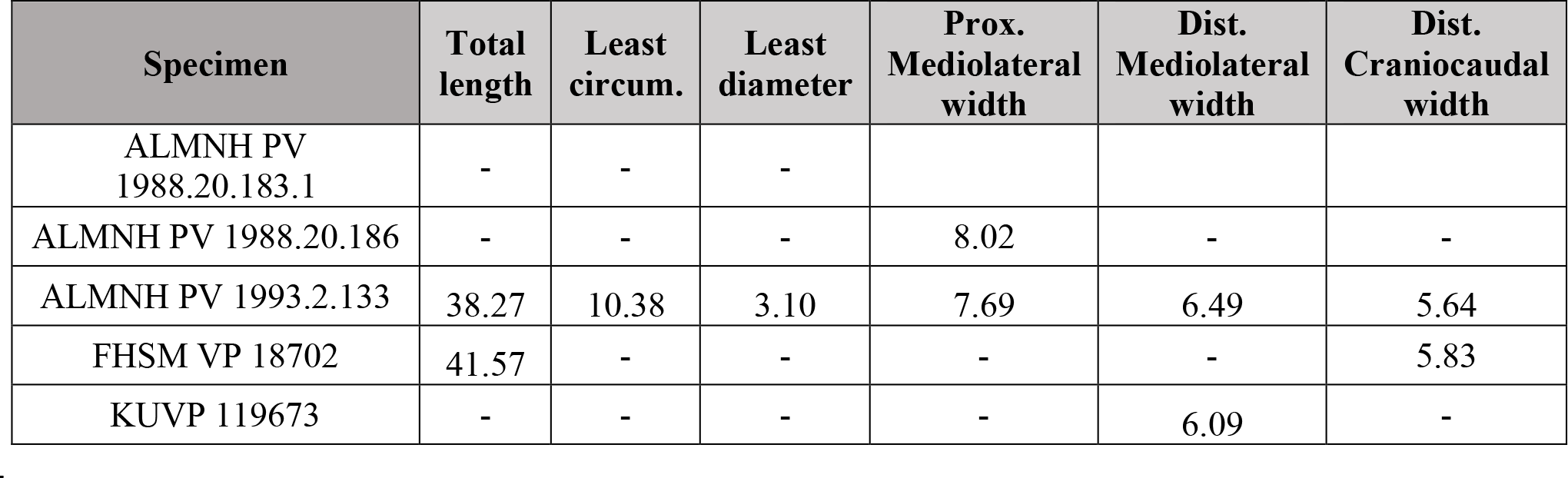
Measurements of the femur of *Ichthyornis* specimens. Least circumference and least diameter correspond to the minimum circumference and diameter of the humeral shaft; midshaft width is provided for those specimens in which the preservational state precluded measuring these. Proximal mediolateral width is measured as the maximum mediolateral extension of the femur, including the femoral head and the femoral trochanter. Midshaft width corresponds to the mediolateral width of the femoral shaft halfway along its length. Asterisks (*) denote measurements that might be unreliable due to breakage or distortion, but are included for completeness. All measurements are in mm. - = not measurable.

The femoral head is large and globose, with a slightly flattened articular facet for the acetabulum (facies artic. acetabularis; Baumel & Witmer, 1993), and a deeply excavated depression found on the ventral surface of the femoral neck (coll. fem.; Livezey & Zusi, 2006), where it meets the femoral shaft and the antitrochanteric articular facet. A round and deep capital ligament fossa is found in all three specimens, even in the extremely distorted femur of FHSM VP-18702. This fossa is present in many crownward euornitheans, such as *Gansus* (Wang et al., 2016b) and *Apsaravis* (Clarke & Norell, 2002), although it is absent in more stemward euornitheans such as *Similiyanornis* and *Abitusavis* (Wang et al., 2020). The femoral trochanter is poorly developed proximally, and it does not extend further dorsally than the femoral head, hence no notable trochanteric fossa is developed (Fig. 24a,b).

Although the development of the femoral trochanter is poorly characterized among stem euornitheans, the minimal dorsal extent of the trochanter in *Ichthyornis* is similar to the condition in *Patagopteryx* (Chiappe, 2002), *Gansus* (Wang et al., 2016b) and *Abitusavis* (Wang et al. 2020), but distinct from *Vorona* (Forster et al., 2002), *Apsaravis* (Clarke & Norell, 2002) *Iteravis* (Zhou et al., 2014) and Hesperornithes (Zinoviev, 2011; Bell & Chiappe, 2016, 2020; Bell et al., 2019), in which it extends beyond the femoral head. The dorsal extension of the femoral trochanter has been suggested to have a strong correlation with the swimming capabilities of water-dwelling birds, and in particular, a short femoral trochanter at the same level as the femoral head appears to be associated with foot-propelled swimming (Raikow, 1970, 1985; Zinoviez, 2011; Clifton et al., 2018; Bell et al., 2019). The condition in *Ichthyornis* is comparable to that of several crown birds exhibiting foot-propelled swimming, such as *Anas platyrhynchos* and *Anser albifrons* (Anseriformes)*, Puffinus lherminieri, Hydrobates leucorhous* and *Diomedea cauta* (Procellariiformes)*, Morus bassanus* and *Phalacrocorax carbo* (Suliformes) and *Phaethon lepturus* (Phaethontiformes), while it is slightly more developed in specialized foot-propelled diving taxa such as *Podiceps auritus* (Podicipediformes) and *Gavia arctica* (Gaviiformes; Bell et al., 2019). In contrast, the femoral trochanter is usually much more developed in taxa that are associated with aquatic habitats but which are less natatorial, such as *Chauna chavaria* (Anseriformes), *Rallus striatus* (Gruiformes), *Sterna hirundo* and *Rynchops flavirostris* (Laridae), and *Charadrius rubricollis* (Charadriidae), and in most land-dwelling birds, such as *Crypturellus variegatus* (Tinamidae) or *Gallus gallus* (Phasianidae). The trochanteric crest is short and does not extend cranially beyond the main body of the femur, contrary to most surveyed extant birds, with the exception of *Puffinus lherminieri*. The barely developed crest is visible in ALMNH PV93.20.5, extending from the dorsal onto the cranial surfaces of the femoral trochanter, but it is even less developed in ALMNH PV98.20.186. The cranial surface of the proximal femur is moderately concave and delimited laterally by the trochanteric crest, and its surface is pierced by numerous foramina in both ALMNH PV93.20.5 and ALMNH PV98.20.186. A large ovoid muscle attachment scar extends from this region into the femoral shaft, probably for the m. femorotibialis intermedius, delimiting on its lateral edge a shallow cranial intermuscular line, which extends from the distal end of the trochanteric crest but quickly becomes indistinct, contrary to the condition in *Gansus* (Wang et al., 2016b).

The lateral surface of the proximal femur preserves several deep and well-developed pits and grooves, which allow for a precise reconstruction of the muscular attachment points of the femoral trochanter (Fig. 24a, b). Three small scars are found on the trochanteric caudolateral surface; of these, the dorsalmost pit-like scar is the deepest, probably corresponding to the attachment point for the m. obturator (impr. m. obtur; Livezey & Zusi, 2006). Distal to this pit, there is a short longitudinal groove extending proximodistally, and just distal to that, there is a slightly shallower comma-shaped scar; these probably correspond to the attachments for the m. iliofemoralis externus and m. ischiofemoralis. A well-developed crest (apparently corresponding to the “trochanter minor” of Guetie. 1976) is situated just craniolateral to these scars; this crest is vaguely sigmoidal in shape, curving caudally on its distal end, and extending distally for about 11% of the total femoral length. Two deeply excavated consecutive grooves run proximodistally just parallel to the craniolateral surface of this crest (Fig. 24b); the proximal and more elongated one corresponds to the attachment for m. iliotrochantericus caudalis, while the shorter and more curved distal groove serves as the attachment for m. iliotrochantericus cranialis or medius (Baumel & Raikow, 1993).

The femoral shaft is only well preserved in ALMNH PV93.20.5, although multiple breakages along its length obscure the muscular scars on its surface. The shaft is mostly straight along its length, narrowing distally. It is only moderately curved caudally on its distal portion, where the shaft becomes significantly broader in cross-section (Fig. 24a). The caudal intermuscular line is visible and very marked along the distal half of the shaft, separating the attachment regions for the m. femorotibialis lateralis and medialis. Although it is possible that the intermuscular line extended further proximally, the breakages in the region make this difficult to assess. Contrary to the condition in all surveyed extant birds, the caudal intermuscular line in *Ichthyornis* does not divide into two distinct branches until very close to the distal femoral condyles, not delimiting a large popliteal plane. A large and very well marked foramen is present approximately at the midpoint along the shaft length, just lateral to the apparent proximal origin of the caudal intermuscular line; a similar foramen is variably present in several crown bird lineages, including anseriforms such as *Anas platyrhynchos*, and suliforms like *Morus bassanus* and *Phalacrocorax carbo*.

The distal femur is only well preserved in ALMNH PV93.20.5, since although ALMNH PV98.20.183.1 preserves this region, its bone surface is mostly eroded. The cranial surface of the distal femur shows a wide but shallow and poorly defined patellar groove, which continues into a deep intercondylar groove distally (Fig. 24a). The distribution of the patellar groove amongst non-neornithine euornitheans is poorly known due to preservation, although it is inferred to be absent in *Yixianornis* (Clarke et al., 2006), but present in *Apsaravis* (Clarke et al., 2006) and Hesperornithes such as *Parahesperornis* (Bell & Chiappe, 2020). The presence of a patellar groove is widespread among crown birds, but the wide and poorly defined groove in *Ichthyornis* is most similar to the condition in Procellariiformes like *Ardena tenuirostris* and Suliformes such as *Morus bassanus.* Both distal condyles are well developed, with the lateral condyle being larger and extending further distally than the medial condyle. The cranial surface of both tubercles is rounded and poorly defined, and no clear cranial ridges delimiting the patellar groove are developed on either condyle. The proximocranial termination of the medial condyle develops into a moderately developed tubercle, while the lateral condyle extends gradually from the cranial surface of the shaft. Two deep and pit-like adjacent impressions of similar size are present along the craniodistal edge of the lateral condyle (Fig. 24a). The one situated more cranially corresponds to the depression of the m. tibialis cranialis tendon, while the one situated more laterally corresponds to the impression of the lig. collaterale laterale (Baumel & Witmer, 1993).

The lateral surface of the lateral condyle is mostly eroded, but the fibular trochlea is well marked and developed, extending laterally from the lateral surface of the condyle along almost 1/5^th^ of the total distal width of the femur. The lateral extension of the fibular trochlea is not known in most non-neornithine euornitheans, but the condition in *Ichthyornis* is much less prominently developed than the condition in Hesperornithes or in extant diving birds, such as *Phalacrocorax carbo*. A small and poorly developed tubercle for the attachment of m. gastrocnemialis lateralis is present on the proximal end of the fibular trochlea, and just distal to it, a wide but shallow and poorly defined impression for the ansae m. iliofibularis is visible (Baumel & Witmer, 1993). The tibiofibular crest is large, subtriangular in lateral and medial views (Fig. 24a), and its caudal tip is slightly proximally curved, exhibiting a morphology particularly reminiscent of the condition in the hesperornithean *Fumicollis* (Bell & Chiappe, 2015, 2020), but distinct from that of other Hesperornithes, in which the crest is much more rounded. Among extant birds, the general shape of the tibiofibular crest is remarkably similar to that of the tinamou *Crypturellus variegatus* and the suliform *Phalacrocorax carbo*, while the morphology is much more rounded and less caudally extensive in most other surveyed crown birds. A shallow, rounded, and poorly marked depression is found just proximal to the tibiofibular crest, probably corresponding to the impression for the lig. cruciati caudalis (Baumel & Witmer, 1993). Immediately proximal to this depression, a short unidentified ridge extends laterodistally, and a small but marked tubercle is found just proximal to it.

The medial condyle lacks a distinct medial epicondyle, but a large shallow and rounded depression is present on its medial surface for the attachment of the medial collateral ligament. The medial condyle is mediolaterally wide and extends medially past the medial surface of the distal femur. Its distal surface is mostly flat, and the medial condyle shows a subquadrangular shape in caudal view. A moderately developed medial supracondylar crest extends from the medial condyle towards the shaft, meeting one of the distal termini of the caudal intermuscular line.

A shallow but well-defined ovoid popliteal fossa is present just proximal to the medial condyle on the caudal surface of the distal femur (Fig. 24a). The fossa is proximodistally short but lateromedially wide, and lies almost parallel to the distal surface of the femur. The presence of a popliteal fossa is uncertain in most non-neornithine euornitheans, but it is present in Hesperornithes such as *Parahesperornis* (Bell & Chiappe, 2002). A popliteal fossa of comparable depth and shape to that of *Ichthyornis* is variably present among crown birds, such as *Chroicocephalus novaehollandiae* (Laridae) and *Morus bassanus* (Sulidae). A large foramen is present at the deepest point of the popliteal fossa, similar to the condition in all surveyed crown birds. The caudal surface of the distal femur just proximal to the popliteal fossa is rugose and complex, with multiple ridges and depressions extending from the caudal intermuscular line, probably representing muscle or ligament attachment points, but these are unidentifiable at present. The caudodistal surface of the distal femur shows an extremely deep and distinct impression of the lig. cruciati cranialis between both distal condyles.

### TIBIOTARSUS

Five of the studied specimens preserve partial tibiotarsi, in all cases preserving the distal end of the element (Fig. 25). Only ALMNH PV93.20.5 preserves the proximal portion of the left tibiotarsus together with the distal end of the same bone. However, even that element is broken, and at least a small portion of the shaft appears to be missing since the fractured ends do not precisely match. As such, a precise estimate of tibiotarsus length, and therefore a definitive assessment of complete hindlimb proportions, is not presently possible. Despite the illustration of a complete tibiotarsus from the holotype of *Ichthyornis victor* in Marsh (1880), no complete tibiotarsi are currently known for *Ichthyornis*, and the element illustrated by Marsh might have been lost or broken (Clarke, 2004). Measurements of the tibiotarsal fragments from the specimens included in this study are provided in Table 14.

**Figure 25.**
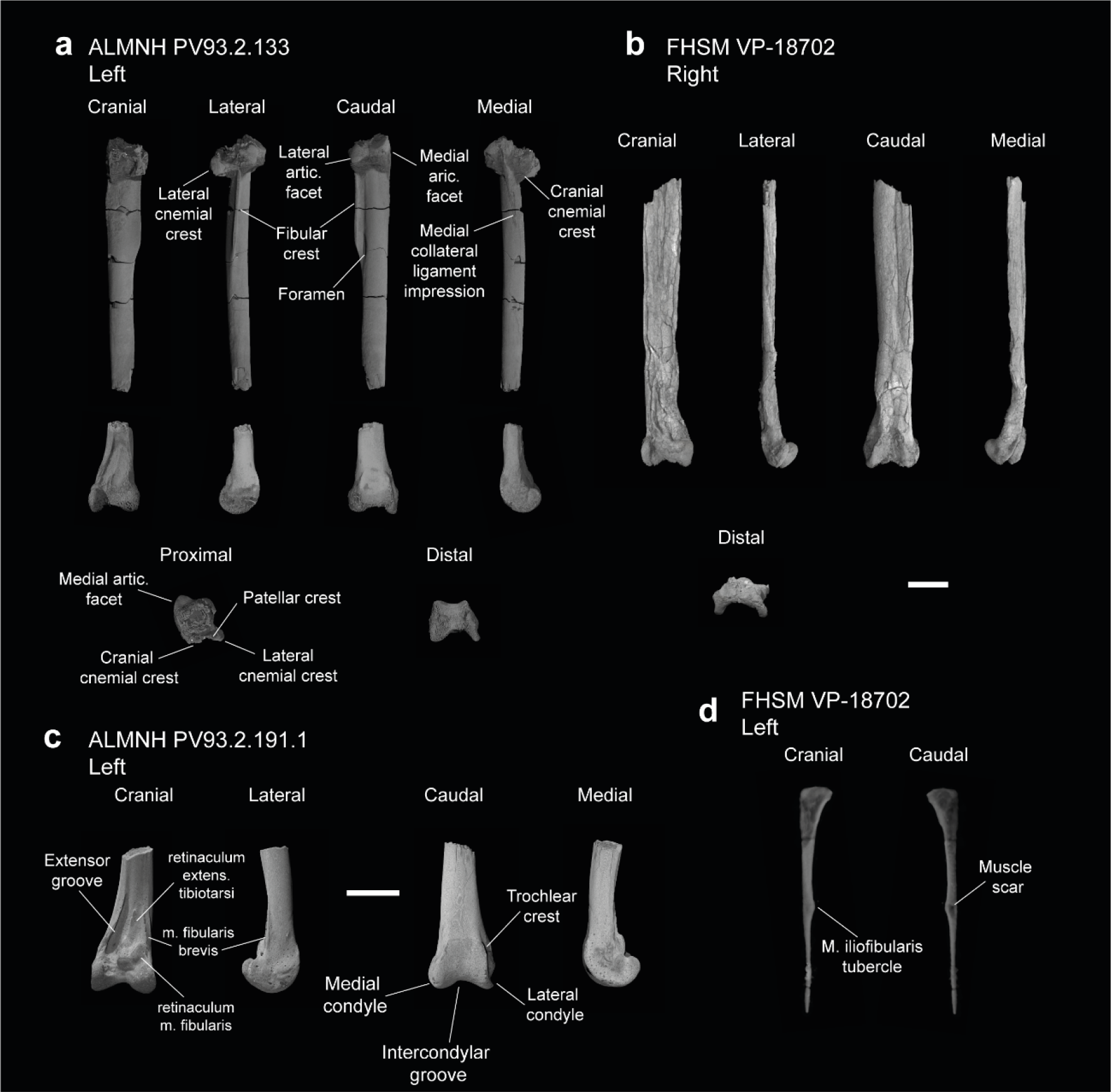
Tibiotarsi and fibula of *Ichthyornis*. (a) ALMNH PV93.2.133 left tibiotarsus proximal and distal fragments, (b) FHSM VP-18702 right tibiotarsus, (c) ALMNH PV93.2.191.1 left distal tibiotarsus and (c) FHSM VP-18702 right fibula. Scale bars equal 5 mm.

**Table 14.**
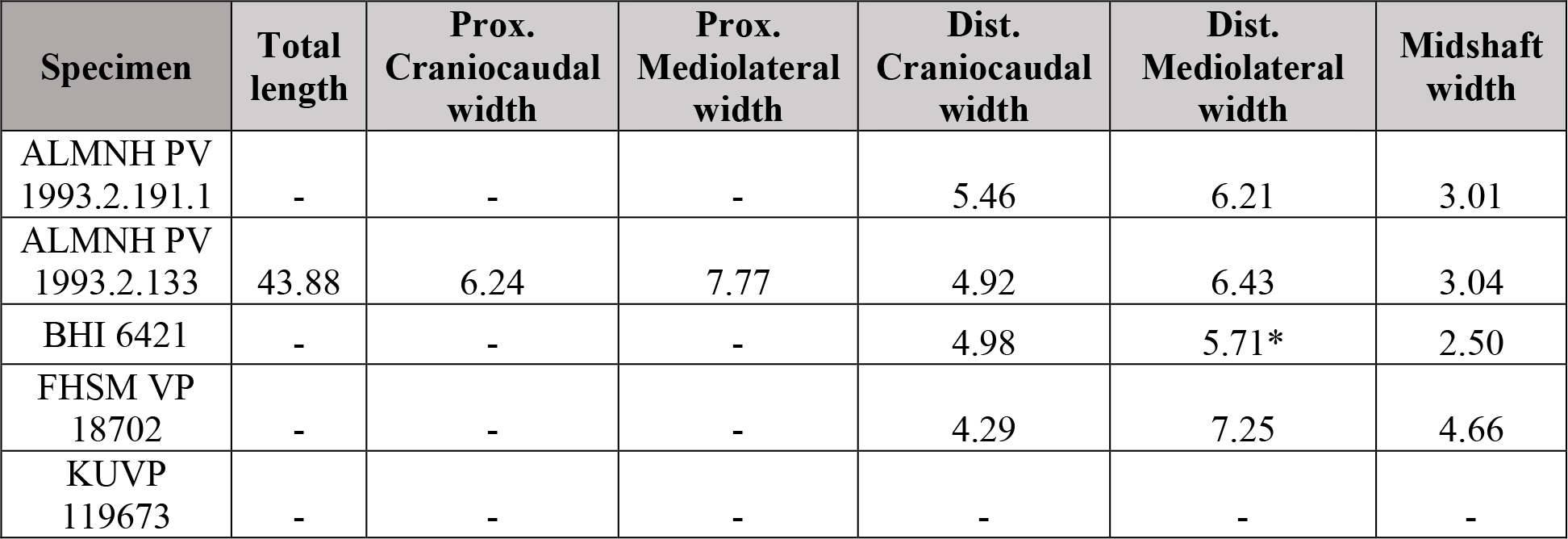
Measurements of the tibiotarsus of *Ichthyornis* specimens. Midshaft width corresponds to the craniocaudal or mediolateral width of the tibiotarsus shaft halfway through its length, depending on the preservation of each specimen. Asterisks (*) denote measurements that might be unreliable due to breakage or distortion, but are included for completeness. All measurements are in mm. - = not measurable.

The proximal end of the left tibiotarsus in ALMNH PV93.20.5 is crushed and distorted proximodistally, but preserves several features that have not been previously illustrated. The specimen may represent the best-preserved proximal tibiotarsus from *Ichthyornis* known to date. The proximal surface is damaged, but the patellar crest (crista patellaris; Baumel & Witmer, 1993) and the lateral and medial articular surfaces are well preserved (Fig. 25a). The width of the proximal portion of the tibiotarsus seems to be significantly greater than that of the shaft, in contrast to the condition in YPM 1450, though this might be caused by distortion of this region in ALMNH PV93.20.5. The medial articular surface is larger than the lateral articular surface (*contra* Clarke, 2004, where it is described as slightly smaller), and it extends further both proximally and caudally with respect to the lateral articular surface. The proximal surface of the medial articular surface is flat and ovoid in proximal view, and is slightly inclined proximodistally with respect to the main axis of the tibiotarsus. The lateral articular surface is more reduced in size and lacks a well-preserved proximal surface, though the preserved portion of the structure suggests a convex morphology as described by Clarke (2004). Contrary to Clarke (2004), a well-developed and obvious fibular crest is present, running along half of the preserved proximal portion of the tibiotarsus, and extending laterally as far as the lateral articular surface. The crest seems to be continuous with the lateral articular surface, contrary to the condition illustrated by Marsh (1880) in which the crest appears to be absent, but Marsh’s interpretation might have been a result of distortion of the proximal end of the tibiotarsus in YPM 1450. The cranial surface of the fibular crest is flat, but its caudal surface is slightly concave, with a thickened lateral edge. The morphology of the fibular crest is fairly similar to the condition in *Gansus*, particularly with regard to its lateral projection (Wang et al., 2016b). A large, deep foramen is present just medial to the distal terminus of the fibular crest. The foramen is found at the end of a short but deep sulcus running next to the medial end of the crest. A similar foramen is variably present in crown birds, but the condition in *Ichthyornis* is most similar to that of Anseriformes, particularly in *Anser albifrons*. A flexor fossa is not apparent, but this may be an artifact of poor preservation.

Two cnemial crests are present in ALMNH PV93.20.5, but only the lateral crest is well-preserved (Fig. 25). The cranial cnemial crest is missing its proximal and cranial ends, but the preserved portions indicate that it was large and robust. The cranial surface of the cnemial crest is subcircular in medial view, and its cranial extension is approximately equal to the shaft diameter. The preserved portion of the cranial cnemial crest does not show any lateral curvature, delimiting a completely flat gastrocnemial surface (facies gastrocnemialis; Baumel & Witmer, 1993). The lateral cnemial crest is thick and robust, slightly longer in its lateral extension than the cranial extension of the cranial cnemial crest, but it is proximodistally shorter, with its proximal end distal to that of the cranial crest. It shows a slight caudolateral curvature, with a flat cranial surface and a moderately convex caudal surface. No patellar crest is preserved, but given the limited proximal extension of the lateral cnemial crest, it was probably not strongly developed. Although distortion of the proximal region of the tibiotarsus might obscure the true proximal extent of both cnemial crests, their visible morphology is congruent with that described for YPM 1450 (Clarke, 2004), and contrasts with the condition in other crownward euornitheans like *Gansus* and *Iteravis,* in which the cnemial crests are moderately cranially projected.

The shaft is not completely preserved in any of the studied specimens, but ALMNH PV93.20.5, BHI 6421 and FHSM VP-18702 preserve large portions of it (Fig. 25). The shaft is mostly straight but shows a moderate lateral twist close to the distal end of the bone in BHI 6421, and reaches its narrowest point distally at around 75% of its length. While no impressions are visible in ALMNH PV93.20.5, a shallow and narrow fibular impression runs across the length of the shaft in BHI 6421, reaching the distal portion of the element.

A flat, oval-shaped impression is visible on the medial surface of the shaft, just medial to the cranial cnemial crest. This impression is congruent with that of the ligamentum collateralis medialis (Baumel & Witmer, 1993), and extends into a shallow ridge that runs across the medial surface of the shaft, with a similar distal extension to that of the fibular crest.

The distal portion of the tibiotarsus is well preserved in ALMNH PV93.20.5, ALMNH PV93.2.191.1, and BHI 6421, and is considerably craniocaudally flattened in FHSM VP-18702 (Fig. 25). The observable morphology is congruent in all of these specimens, and agrees well with that previously reported for *Ichthyornis* (Clarke, 2004), although the specimens described here are the first undistorted examples reported thus far. The extensor groove (sulcus extensorius; Baumel & Witmer, 1993) is deep and marked, running moderately obliquely to the main axis along the medial side of the shaft, terminating just proximal to the medial condyle. The medial edge of the groove is delimited by a thin but sharp ridge, which terminates just proximal to the lateral condyle. No supratendinal bridge is present in any of the studied specimens. A supratendinal bridge is not preserved in any of the YPM *Ichthyornis* specimens either, but Clarke (2004) did not dismiss its possible presence given the poor state of preservation of the YPM specimens. The exceptionally preserved ALMNH specimens and BHI 6421 indicate that the lack of a supratendinal bridge in *Ichthyornis* is genuine (Fig. 25). This structure therefore optimizes as a synapomorphy of the most exclusive clade composed of Neornithes and the most crownward-known stem birds based on its presence in *Iaceornis marshi* (Gauthier, 1986; Martin, 1985, Cracraft, 1988; Mayr & Clarke, 2003; Clarke, 2004) and crown birds.

Several tubercles, impressions and scars are visible on the cranial surface of the distal tibiotarsus in all of the studied specimens. The largest of these, a large subtriangular concave surface, is situated just lateral to the extensor groove (Fig. 25c). This surface is sharply raised on its distal end, extending cranially along at least 60% of the lateral condyle’s length. This surface might correspond to the implantation for the retinaculum extensorium tibiotarsi and the passage of the m. extensor digitorum longus (Baumel & Raikow, 1993), as interpreted by Clarke (2004). On the laterodistal end of this concave surface there is a raised ovoid tubercle which extends only slightly further cranially (Fig. 25). It likely corresponds to the implantation region for the retinaculum m. fibularis (Baumel & Raikow, 1993). Lateral to both, there is a shallow groove which extends from the lateral side of the fibular groove, turning progressively medially closer to the proximal end of the extensor groove and then laterally closer to the proximal edge of the lateral condyle, with a marked foramen on its distal end. This groove is delimited by two sharp ridges along most of its length and corresponds with the impression of the m. fibularis brevis. The morphology of this region is not well described in the literature for most Mesozoic euornitheans, but the muscle impressions and tubercles in *Ichthyornis* seem to correspond broadly with the three tubercles described in *Gansus* (Wang et al., 2016b). Despite presumed changes in muscular configuration associated with the evolutionary origin of a supratendinal bridge (Hutchinson, 2002), the muscle implantation arrangement is remarkably similar to the condition in extant Laridae such as *Chroicocephalus* and *Sterna*.

Both distal condyles are very similar in size, shape, and distal extent, and are approximately round in lateral and medial view (Fig. 25). The lateral condyle is slightly wider and broader than the medial condyle, which in turn extends slightly further cranially. The intercondylar groove is wide and shallow, and the proximal ends of both condyles gradually slope towards the midline of the cranial surface. The medial surface of the medial condyle is concave and excavated, with a moderately developed medial epicondyle. The lateral surface of the lateral condyle is mostly flat, and no lateral epicondyle is present. In caudal view, raised ridges (trochlear crests) extending from the distal edges of both condyles delimit a well-developed and subquadrangular caudal tarsometatarsal articulation (trochlea cartilaginis tibialis, Baumel & Witmer, 1993). The proximal edge of the trochlea is almost completely horizontal. Proximal to the trochlea, a shallow groove running parallel to the edge of the trochlea is visible in ALMNH PV93.2.191.1 and BHI 6421 (Fig. 25c).

### FIBULA

The fibula has never been previously recovered or described for *Ichthyornis*, but three fibulae are preserved between two of the new specimens described here. In FHSM VP-18702 both fibulae are preserved in disarticulation, with an almost complete right fibula and the proximal portion of the left fibula. KUVP 119673 preserves a left fibula which is disarticulated, but in association with the distal tibiotarsus. Most of this specimen’s proximal morphology is taphonomically distorted (Fig 25d).

The preservation of the fibulae in the new specimens does not allow a comprehensive description, as several features are either obscured or indistinct. The fibula is very slender and elongate. In all three preserved fibulae, the element appears rather flat and mediolaterally compressed, but it is unclear whether this is taphonomic in nature or representative of the original morphology. The lateral surface of the proximal fibula in FHSM VP-18702 appears mostly flat to slightly convex, but a shallow tubercle seems to be developed at its midpoint.

The medial surface is shallowly excavated and mostly concave. A small foramen seems to be developed on the medial surface (Fig. 25). It is unclear whether this is opening represents an artifact of preservation, although a similarly excavated foramen is present in this region in *Parahesperornis* (Bell & Chiappe, 2020). The proximal end of the fibula is subtriangular in lateral view, with a slightly convex articular surface for the femur and a relatively large, caudally directed crest. The mediolateral extension of the fibula at its widest point is approximately equal to the maximum width of the tibiotarsal shaft.

The fibular shaft is thick and subtriangular in cross section along its proximal half, with a slightly narrower caudal edge conferring a keeled shape (Fig. 25). A marked and distinct tubercle for the m. iliofibularis (Baumel & Witmer, 1993) is developed halfway along the preserved portion of the shaft. As in other Ornithurae (including crown birds; Chiappe, 1996), the tubercle is caudally directed and shows a deep excavation on its lateral surface both in FHSM VP-18702 and KUVP 119673. The width of the shaft remains mostly constant proximal to the m. iliofibularis tubercle, but distal to the tubercle the shaft tapers considerably and becomes extremely thin and rod-like. None of the specimens preserve the complete length of the fibular shaft, but it would have probably extended more than halfway along the length of the tibiotarsus and into the distal region of the element, since fibular impressions are apparent on the lateral surface of the distal tibiotarsus in BHI 6421. This inferred length of the fibula seems longer than that reported for *Gansus* and *Iteravis* (Liu et al., 2014; Zhou et al., 2014; Wang et al., 2016b), and is substantially longer than that of *Yixianornis* and *Yanornis* (Zhou & Zhang, 2001; Clarke et al., 2006; Wang et al., 2020), which both show highly reduced fibulae. The general morphology and distal extent of the *Ichthyornis* fibulae appear remarkably similar to those of the Common Tern (*Sterna hirundo*), which also shows an expanded, convex proximal end and a very elongate fibular shaft.

### TARSOMETATARSUS

Five of the studied specimens include at least one tarsometatarsus (Fig. 26). Of these, only FHSM VP-18702 preserves a complete element, and, although this element is highly distorted, it apparently represents the first occurrence of a complete tarsometatarsus for *Ichthyornis*. Although the remaining tarsometatarsi are broken to varying extents, four of them from the ALMNH collections are exceptionally well-preserved: PV93.20.5 preserves both the proximal and distal portions of the left tarsometatarsus (though a small part of the shaft is missing), PV88.20.375.3 includes only the proximal end, and PV88.20.180 and PV88.20.181 are both distal tarsometatarsi. Measurements of the tarsometatarsi of the specimens included in this study are provided in Table 15.

**Figure 26.**
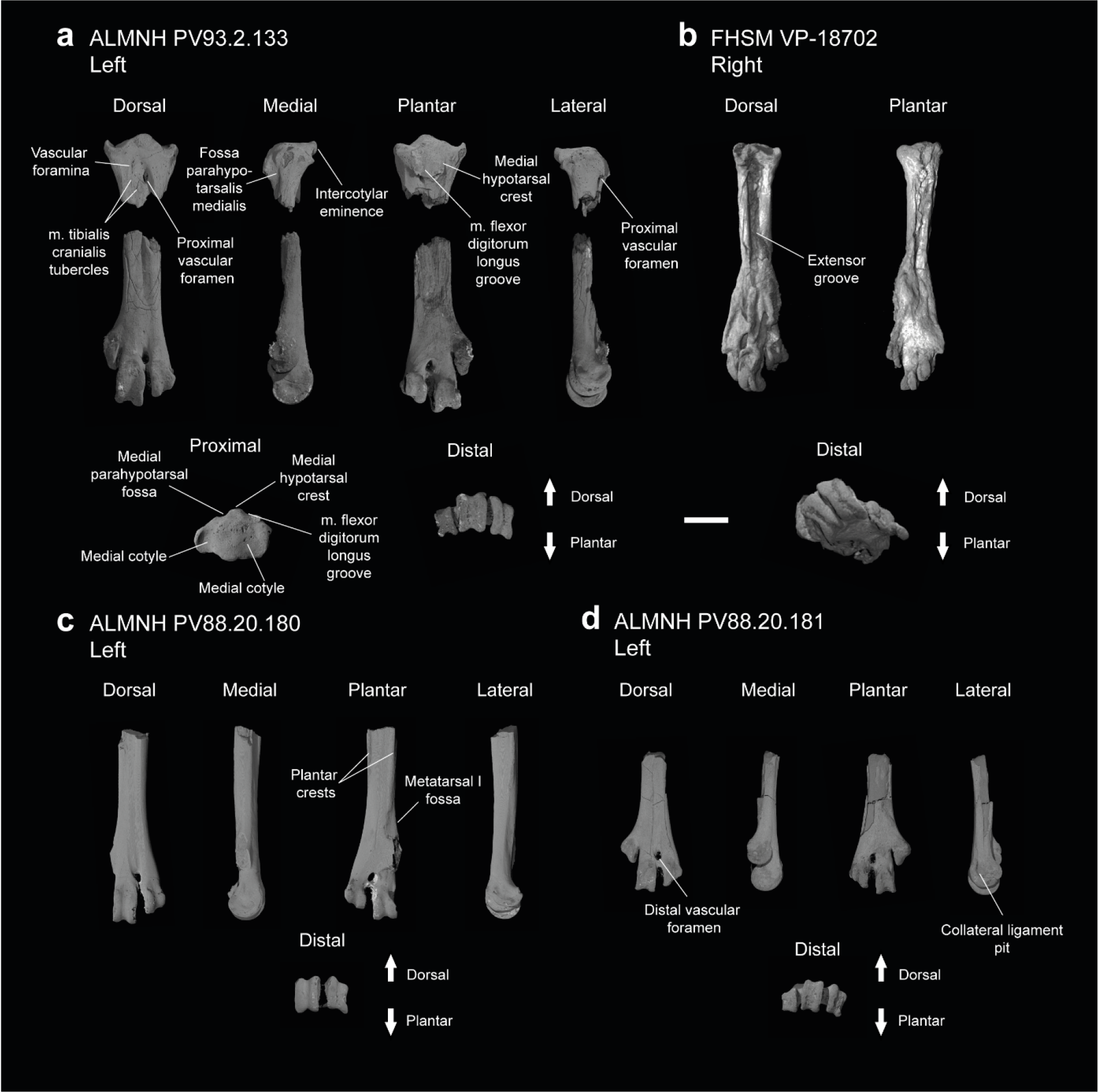
Tarsometatarsi of *Ichthyornis*. (a) ALMNH PV93.2.133 left tarsometatarsus proximal and distal fragments, (b) FHSM VP-18702 right tarsometatarsus, (c) ALMNH PV88.20.180 left distal tarsometatarsus and (c) ALMNH PV88.20181 left distal tarsometatarsus. Scale bar equals 5 mm.

**Table 15.** Measurements of the tarsometatarsus of *Ichthyornis* specimens. Total length corresponds to the maximum proximodistal extension of the tarsometatarsus, measured from the proximal intercotylar eminence to the distalmost point of metatarsal III trochlea. Proximal dorsoplantar width measurements include the hypotarsus, when preserved. Distal dorsoplantar width corresponds to the maximum dorsoplantar extension of metatarsal III trochlea. Distal mediolateral width is measured from the medialmost point of metatarsal II trochlea to the lateralmost point of metatarsal IV trochlea. Midshaft width corresponds to the mediolateral width of the tarsometatarsus shaft halfway through its length. Trochleae widths correspond to the maximum mediolateral extension of metatarsal II, III and IV trochleae. Asterisks (*) denote measurements that might be unreliable due to breakage or distortion, but are included for completeness. All measurements are in mm. -= not measurable.

| Specimen | Total length | Prox. Dorsoplantar width | Prox. Mediolateral width | Dist. Dorsoplantar width | Dist. Mediolateral width | Midshaft width | Trochlea II width | Trochlea III width | Trochlea IV width |
| --- | --- | --- | --- | --- | --- | --- | --- | --- | --- |
| FHSM VP 18702 | 29.13 | 4.21 | 5.69* | - | 7.78* | 3.78 | - | 1.93* | 1.75* |
| ALMNH PV 1988.20.180 | - | - | - | 3.82 | - | 3.19 | - | 2.51 | 2.20 |
| ALMNH PV 1988.20.181 | - | - | - | 3.67 | 7.66 | 2.88 | 1.68 | 2.32 | 1.88 |
| ALMNH PV 1988.20.375.3 | - |  |  |  |  |  |  |  |  |
| ALMNH PV 1993.2.133 | - | 4.84 | 7.24 | 3.72 | 7.43 | 3.42 | 1.87 | 2.57 | 1.96 |

The tarsometatarsus in *Ichthyornis* is completely fused, with both tarsals and metatarsals completely indistinguishable from one-another except in the distal region, where the articular surfaces for the pedal phalanges of metatarsals II-IV are distinct. The whole element is short and robust, with the tarsometatarsus of FHSM VP-18702 measuring 60% of the length of the femur (Fig. 26b; Tables 13 and 15). The tarsometatarsus exhibits reduced mediolateral expansion at both the proximal and distal regions in relation to its shaft. The generally stout form of the *Ichthyornis* tarsometatarsus is reminiscent of the tarsometatarsi of *Yanornis* and *Yixianornis* (Clarke et al., 2006), and differs from the more elongate condition in the more closely related *Iteravis* and *Gansus* (Liu et al., 2014; Zhou et al., 204; Wang et al., 2016b). Metatarsal I is not preserved in any of the studied specimens, indicating that it probably separated easily after death.

The proximal region of the tarsometatarsus is best preserved in ALMNH PV93.20.5, with no apparent distortion or breakage. The proximal end of metatarsal III is situated plantar to those of metatarsals II and IV. The intercotylar eminence (eminentia intercotylaris; Baumel & Witmer, 1993) is moderately developed, projecting only slightly proximally with respect to the lateral and medial edges of the tarsometatarsal cotyles (Fig. 26a). The dorsal edge of the eminence extends slightly dorsally beyond the rest of the proximal surface of the tarsometatarsus, and is slightly proximodistally recurved in lateral view, defining a marked and globose surface on its proximal end in dorsal view. Both cotyles are shallow and concave in proximal view, and, contrary to Clarke (2004), the lateral cotyle has a larger proximal surface than the medial cotyle. The medial cotyle is situated slightly more proximally than the lateral cotyle, is more deeply concave, and is delimited by a thickened ridge dorsally and medially, with the ridge projecting slightly less proximally than the intercotylar eminence on its medial side. A similar but less-developed ridge is present on the lateral cotyle, which ends distal to that of the medial condyle. The proximal extension of the medial cotylar edge does not seem to be evident in the YPM tarsometatarsi, as illustrated and described previously (Clarke, 2004), and seems to be broken in ALMNH PV88.20.375.3.

Several foramina are present in the proximal depression between metatarsals, as described by Clarke (2004). A large foramen penetrates the bone in the space between metatarsals III and IV, and opens plantarly just distal to the hypotarsus (Fig. 26a). Similar large, proximal vascular foramina are also found in some Mesozoic euornitheans such as *Gansus* and *Apsaravis* (Clarke & Norell, 2001; Wang et al., 2016b), as well as among crown-birds such as *Sterna hirundo* and *Puffinus lherminieri*, but they differ from the condition in Hesperornithes, in which these do not perforate the tarsometatarsus (Bell & Chiappe, 2020). An additional, previously undescribed minute foramen is present between metatarsals III and IV, proximal to the large foramen. Clarke (2004) described a series of three foramina present between metatarsals II and III, but only two can be seen in this region in ALMNH PV93.20.5. Contrary to Clarke (2004), no corresponding foramen is present opposite to these on the plantar surface of the new tarsometatarsi.

The extensor groove is deeply excavated and wide, extending for most of the length of the shaft. Only FHSM VP-18702 preserves the whole length of the extensor groove, but its poor preservation distorts the morphology of this region (Fig. 26b). The proximal portion of the extensor groove is well-preserved in ALMNH PV93.20.5. and PV88.20.375.3. The two tubercles described by Clarke (2004) for the implantation of m. tibialis cranialis are present in both specimens but are less distinct than in YPM 1739. The tubercle on the lateral surface of metatarsal II, situated at the same height as the large foramen, is a shallow and slightly concave scar in ALMNH PV93.20.5, and it is barely observable in PV88.20.375.3, instead of representing a clear tubercle. The tubercle on the dorsal surface of metatarsal III is large and clearly developed in ALMNH PV93.20.5, although its morphology would be better described as a raised, flat, ovoid surface rather than a tubercle. A pair of tubercles is developed in the extensor groove in *Gansus* (Wang et al., 2016b), while a single one is present in Hesperornithes (Bell & Chiappe, 2016, 2020).

The hypotarsus and the proximal plantar surface are well preserved in ALMNH PV93.20.5, but eroded and crushed in ALMNH PV88.20.375.3 and FHSM VP-18702. Although YPM 1739 preserves part of the hypotarsus, several of the distinctive features present in the newly described specimens are not preserved in the YPM material (Clarke, 2004). The hypotarsus in *Ichthyornis* is a roughly quadrangular patch of bone, situated mostly behind the lateral cotyle (Fig. 26a). It shows a very limited plantar projection, with a flattened plantar surface showing only very shallow grooves and ridges. The hypotarsus extends distally for about 10% of the length of the tarsometatarsus, terminating just proximal to the plantar opening of the proximal vascular foramen in a sharp depression. A shallow groove is visible running proximodistally, interpreted here as the sulcus for the m. flexor digitorum longus (Hutchinson, 2002; Mayr, 2016). A weakly developed ridge is present on the lateral edge of the groove; it has a flat lateral surface, congruent with the lateral hypotarsal crest as described by Clarke (2004). The medial side of the groove is delimited by a wide and robust ridge, likely homologous with the medial hypotarsal crest (crista medialis flexoris digitorum longus; Mayr, 2016). The crest is situated just plantar to the intercotylar eminence, projecting further plantarly than the lateral ridge, and has a flattened plantar surface. The distal end of the medial crest is broken off in ALMNH PV93.20.5, but it is clear that it had a sharp distal end and did not extend into a medianoplantar crest (crista medianoplantarus; Baumel & Witmer, 1993). The medial surface of the medial crest is concave and moderately excavated, forming a proximodistally directed groove in a similar position to the medial parahypotarsal fossa (fossa parahypotarsalis medialis; Baumel & Witmer; 1993) that serves as an attachment point for the m. flexor hallucis brevis (Vanden Berge & Zweers, 1993). The morphology of the hypotarsus in *Ichthyornis* differs from the rudimentary condition in other crownward euornitheans like *Gansus, Yixianornis,* and *Changmaornis* (Wang et al., 2013) as well as Hesperornithes (Bell & Chiappe, 2016, 2020), in which the hypotarsus, if developed, shows no distinct ridges or grooves. In contrast, the condition in *Ichthyornis* is not dissimilar from that of Lithornithidae, in which a single shallow groove for the m. flexor digitorum longus is medially bounded by a wide, weakly projecting medial crest, which in turn delimits a concave surface on its medial side (Houde, 1988; Nesbitt & Clarke, 2016). This condition is also reminiscent of the morphology seen in the possible early euornithean *Kaririavis* (Carvalho et al., 2021), in which a single tendinal groove for the m. flexor digitorum longus is developed, although in this taxon the hypotarsus exhibits a subtriangular shape, and instead of exhibiting a sharp distal terminus, it extends into an elongated ridge continuing along most of the plantar surface of the tarsometatarsus. The morphology exhibited by *Kaririavis* might suggest the independent development of a hypotarsus in this taxon, especially in light of its uncertain, yet probably stemward phylogenetic position, and the complex combination of plesiomorphic and derived features it exhibits (Carvalho et al., 2021). Among crown birds, the hypotarsal morphology of *Ichthyornis* is most reminiscent of taxa with asulcate hypotarsi like Cathartidae and Cariamiformes (Mayr, 2016). In these, while the hypotarsus is significantly more plantarly projected and shows a flattened proximal surface, the medial and lateral crests are poorly developed, the sulcus for the m. flexor digitorum longus is extremely shallow, and the medial surface of the medial crest is concave and excavated. While this morphology has arisen independently among several crown bird lineages (Mayr, 2016), the similarity between Lithornithidae and *Ichthyornis* may indicate that this condition is plesiomorphic for crown group birds.

The shaft of the tarsometatarsus is mostly straight, with its narrowest point situated about halfway between the proximal and distal ends of the bone. On the dorsal surface, the extensor groove becomes progressively shallower, but remains visible along the whole length of the shaft. The groove is bounded laterally and medially by two shallow, subparallel ridges deriving from the distal termini of the medial and lateral cotyles respectively, which extend into the articular trochleae of metatarsals III and IV. The plantar surface of the shaft is mostly flat and featureless. Moderately extended lateral and medial plantar crests are visible in FHSM VP-18702 and in ALMNH PV88.20.180, bounding the flexor groove (sulcus flexorius; Baumel & Witmer, 1993). The lateral plantar crest terminates halfway along the length of the shaft, but a shallow ridge extends from its distal tip to the lateral surface of the metatarsal IV trochlea. The medial plantar crest extends further distally, terminating at the proximal edge of the metatarsal I fossa, which is only well-preserved in ALMNH PV88.20.180 (Fig. 26c). The fossa is shallow and poorly excavated, but it is clearly divided into a proximal and a distal concave region, separated by a weakly developed horizontal ridge. A marked intermuscular line runs proximodistally across the midline of the plantar surface of the shaft, starting just distal to the distal terminus of the lateral plantar crest. As described by Clarke (2004), the intermuscular line runs parallel to the main axis of the shaft before turning laterally and meeting with the metatarsal IV trochlea.

Metatarsals III and IV delimit a very large, ovoid distal vascular foramen situated far distally, at a similar height to the metatarsal II trochlea and with its distal margin at a similar position to the proximal margin of the metatarsal III trochlea. The dorsal opening of the foramen is found at the distal end of a deep sulcus formed between both metatarsals, and is seemingly continuous with the extensor groove in ALMNH PV88.20.180. However, the groove between the metatarsals is less marked proximally and does not contact the extensor groove in the other newly described specimens that preserve this region. The foramen shows a single plantar opening of similar size, but rounder in shape, at the same height along the shaft. This foramen has two plantar exits in most crown birds, but the single plantar opening is found in several Mesozoic euornitheans, including *Gansus*, *Apsaravis* and the Hesperornithes (Clarke & Norell, 2002; Wang et al., 2016b; Bell & Chiappe 2016; 2020). The distal vascular foramen is weakly or partially enclosed distally, as previously described by Clarke (2004), with the distal bone margin being very narrow and extremely porous.

Of the fused metatarsals, metatarsal II is the shortest and most proximally situated, with its distal end at a height close to that of the proximal end of the metatarsal III trochlea. Metatarsals III and IV show a similar distal extension, with metatarsal III extending slightly further distally. The metatarsal trochleae are essentially coplanar and aligned with the major dorsoplantar axis of the tarsometatarsal shaft, defining a flat and featureless plantar surface of the distal tarsometatarsus. Only the metatarsal II trochlea is slightly more plantarly positioned. This morphology differs from that found in most other Mesozoic euornitheans, including *Yanornis, Changmaornis, Gansus*, *Iteravis*, *Apsaravis,* and Hesperornithes (Zhou & Zhang, 2001; Clarke & Norell, 2002; Wang et al., 2013, 2016; Liu et al., 2014; Zhou et al., 2014; Bell & Chiappe 2016, 2020), in which metatarsal II is plantarly deflected. In *Gansus*, metatarsal IV is more plantarly situated than metatarsal III as well, defining a deeply concave plantar surface, with the cranial surface of the metatarsal II trochlea situated at the midpoint between the dorsal and plantar surfaces of metatarsal II (Wang et al., 2016b). The relative dorsoplantar position of the distal terminus of each metatarsal is highly variable among crown-group birds, but the morphology in *Ichthyornis* is reminiscent of the condition in Procellariiformes such as *Puffinus lherminieri* and *Daption capense*, though the flat and featureless distal plantar surface is more similar to Tinamiformes such as *Crypturellus variegatus*. The trochleae of metatarsals II–IV all show a very well-developed ginglymoid morphology, with deep collateral ligament pits on their lateral and medial surfaces. In metatarsal II (most clearly visible in ALMNH PV88.20.181), the ginglymoid condition is slightly less developed than in metatarsal III and IV, with the sulcus between both lateral and medial ridges of the trochlea being very shallow, and the lateral ridge projecting significantly less distally than the medial ridge. The ginglymoid morphology of the metatarsal II trochlea contrasts with the condition in other euornitheans, such as *Gansus* and *Apsaravis*, in which the articular surface of metatarsal II is rounded (Clarke & Norell, 2002; Wang et al., 2016b). The trochlea of metatarsal III is the widest and proximodistally longest, differing from the condition in Hesperornithes, in which the trochlea of metatarsal IV is the largest (Bell & Chiappe, 2020). The distal terminus of metatarsal IV is dorsolaterally oriented, and its trochlea is highly asymmetrical, with the medial trochlear ridge terminating well dorsal to the lateral trochlear ridge. The lateral surface of the trochlea extends into an elongated flange that continues further plantarly than the metatarsal II trochlea.

### PEDAL PHALANGES

No complete pes is known for *Ichthyornis*, and only a single pedal phalanx is preserved amongst the YPM material (Clarke, 2004). Two of the newly described specimens include pedal phalanges: FHSM VP-18702 preserves four phalanges and ALMNH PV93.20.5 preserves a single phalanx; these are all disarticulated and none represent ungual phalanges (Fig. 29). Unfortunately, definitively establishing which digits they belonged to, and what position they occupied within those digits, is challenging. Measurements of the pedal phalanges of the specimens included in this study are provided in Table 16.

**Table 16.** Measurements of the pedal phalanges of *Ichthyornis* specimens. Asterisks (*) denote measurements that might be unreliable due to breakage or distortion, but are included for completeness. All measurements are in mm. -= not measurable.

| Specimen | Total length | Proximal width | Distal width |
| --- | --- | --- | --- |
| FHSM VP 18702:<br>Phalanx 1 | 17.19 | 2.76 | 1.81* |
| FHSM VP 18702:<br>Phalanx 2 | 9.17 | 2.14 | 1.74 |
| FHSM VP 18702:<br>Phalanx 3 | 12.95 | 2.76* | 1.94 |
| FHSM VP 18702:<br>Phalanx 4 | 13.52 | 2.33 | 1.82 |
| ALMNH PV 1993.2.133 | 12.39 | 2.38 | 1.85 |

The largest phalanx from FHSM VP-18702 is badly flattened and lacks most of its proximal articular surface (Fig. 27a). The phalanx is completely straight, with no lateromedial curvature. The distal articular surface shows a strong ginglymoid condition, with deep pits on its lateral and medial surfaces for the collateral ligaments. This phalanx is greatly elongated compared with the FHSM VP-18702 tarsometatarsus (the phalanx is 60% as long as the tarsometatarsus; Tables 15 and 16). Such elongated phalanges are not known among any other Mesozoic euornitheans, even in those with shortened tarsometatarsi like *Yixianornis* or elongated pedal phalanges like *Gansus* (Clarke et al., 2006; Wang et al., 2016b), in which the proximal digit III phalanx (the longest phalanx) represents around 40% of the total length of the tarsometatarsus. A similar proportion of around 40% is found in Hesperornithes such as *Hesperornis* and *Parahesperornis*, for which the proximal phalanx of digit IV is the longest one. Proportions comparable to *Ichthyornis* between the longest phalanx and the tarsometatarsus are found in several predominantly aquatic crown bird lineages, including Anseriformes, such as *Anas platyrhynchos* (56%), Gruiformes like *Fulica atra* (55%) or Suliformes like *Morus bassanus* (62%). The lack of a well-preserved proximal articular surface, together with the significant mediolateral compression of FHSM VP-18702 complicates the identification of the position of this phalanx. Its extremely elongate condition together with the width of its proximal end (greater than that of the metatarsal III trochlea) suggest that it most likely represents the proximal phalanx of digits II or III.

**Figure 27.**
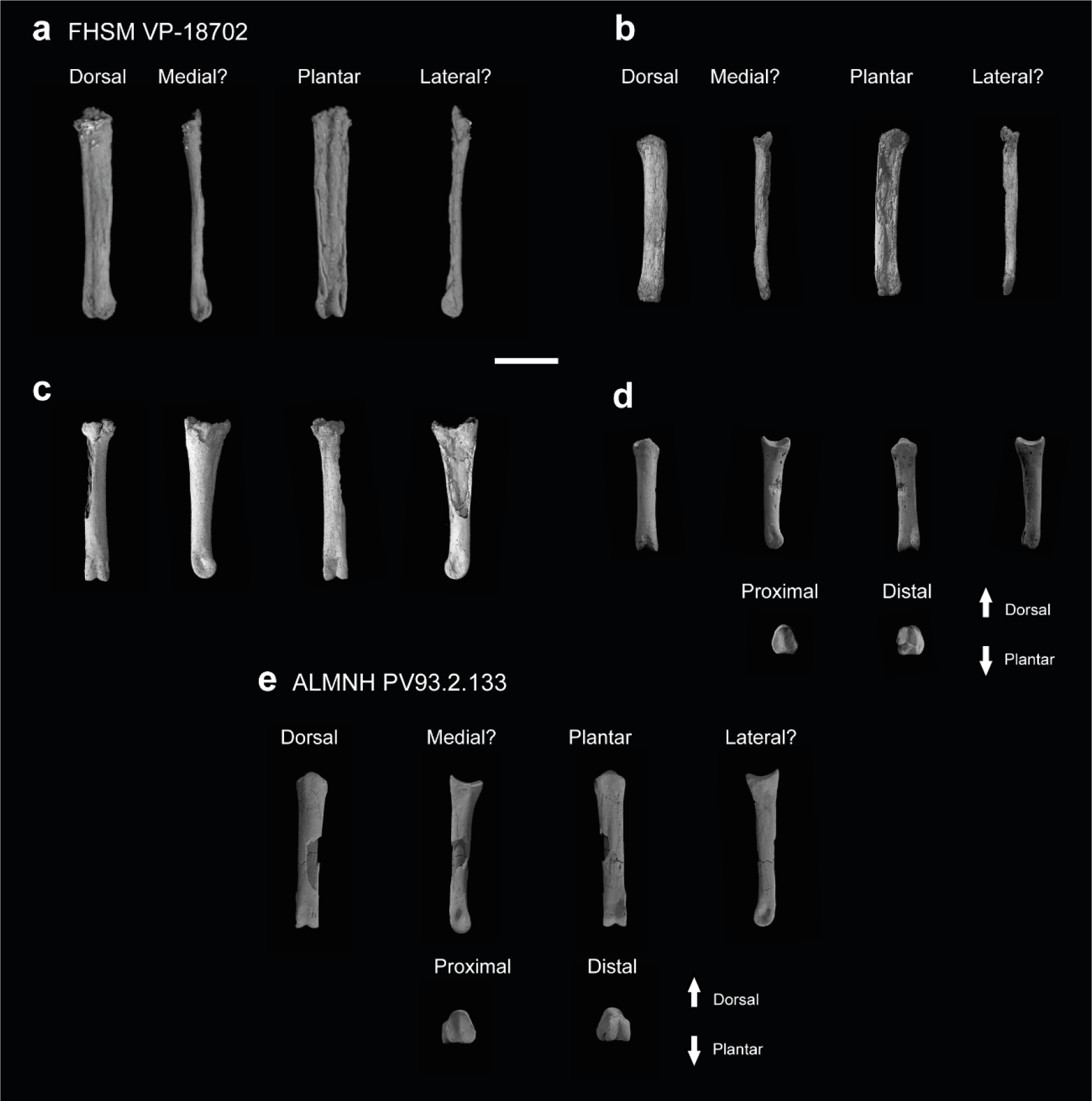
Pedal phalanges of *Ichthyornis*. FHSM VP-18702 (a) phalanx III:1?, (b) phalanx III:2?, (c) phalanx IV:1? (c) phalanx IV:2?; ALMNH PV93.2.133 (d) phalanx IV:1?. Scale bar equals 5 mm.

The second-longest phalanx of FHSM VP 18702 is also dorsoventrally flattened, and while its proximal articular surface is preserved, the distal trochlea is greatly distorted (Fig. 27b). The phalanx shows limited mediolateral curvature, but this appears to be a taphonomic artifact. The phalanx is elongate, and is 82% of the length of the longest phalanx (Table 16). The proximal articular surface, though distorted, appears mostly symmetrical, with two similarly sized cotyles, which suggest that it does not represent the proximal phalanx of digits II or IV. Its proximal end has a similar width to the distal trochlea of the longest phalanx, which suggests that it is most likely the second phalanx of digit III.

The third-longest phalanx is only slightly shorter than the second, and it is well preserved, mostly three-dimensionally (Fig. 27c). The dorsal surface of the phalanx is mostly flat, but the plantar surface is moderately arched. The shaft is mostly straight with no significant proximodistal reduction in width, though the proximal articular surface is significantly wider than the shaft. The proximal articular surface is mediolaterally crushed, but preserves both articular cotyles, which appear slightly asymmetrical. Despite some taphonomic compression, the width of the proximal articular surface is greater than that of the distal trochlea of the longest phalanx. However, the proximal articular surface appears to be considerably wider than the trochleae of metatarsals II and IV as well, which, despite being badly compressed, are not wider than the trochlea of metatarsal III in other specimens (Table 16) . This suggests that this phalanx might represent the second phalanx of digits II or IV, for which no proximal phalanges are preserved, complicating direct comparisons. The distal articular surface of the phalanx is well-preserved and strongly ginglymoid.

The shortest phalanx of FHSM VP-18702 extends for only 52% of the length of the longest phalanx (Table 16). It is extremely well preserved, with the only noticeable blemish being the fact that its distal articular surface is slightly eroded (Fig. 27d). The shaft is mostly straight, and only narrows proximodistally to a slight degree such that the proximal surface is not significantly wider than the shaft. Its plantar surface is more strongly arched than that of the third-longest phalanx, and the ginglymoid morphology of its distal articular surface is more weakly developed. The proximal articular surface preserves two symmetrical and equally sized trochleae. The dorsal proximal margin does not exhibit a strong extensor tubercle, but robust flexor tubercles are found on the plantar surface. The short and stout proportions of this phalanx are congruent with its identity as the second or third phalanges of Digit IV, which exhibits a greater number of non-ungual phalanges (four) than the rest of the digits in most crown birds and in crownward avialans such as *Gansus* (Wang et al., 2016) and *Apsaravis* (Li et a., 2014).

The only pedal phalanx belonging to ALMNH PV93.20.5 is well-preserved, missing only part of the bone surface of the shaft (Fig. 27e). The phalanx is similar in length to the third-longest phalanx of FHSM VP-18702 (Table 16). The shaft is mostly straight with subparallel lateral and medial surfaces, with no significant reduction in width towards the distal end, and the ventral surface of the shaft is only moderately arched. The proximal articular surface, only slightly mediolaterally wider than the shaft, shows two slightly asymmetrical cotyles. The dorsal surface extends well proximally, but does not form a clear extensor tubercle. The ventral surface has only weakly-developed flexor tubercles. The width of the proximal articular surface is similar to that of the ALMNH PV93.20.5 metatarsal IV trochlea, which, together with its slightly asymmetrical cotyles, suggests it represents the proximal phalanx of digit IV. The distal articular surface its strongly ginglymoid, with very deep pits for the collateral ligaments.

### PHYLOGENETIC RESULTS

To assess the referral of the new specimens to *Ichthyornis,* all well-represented specimens were initially incorporated into phylogenetic analyses as distinct operational taxonomic units (OTUs). Despite multiple morphological differences among several of the new specimens, all the new specimens were recovered in an exclusive monophyletic group with YPM 1450, the holotype of *Ichthyornis,* in analyses using the Wang et al. (2020) dataset. Analyses employing the Huang et al. (2016) dataset recovered all *Ichthyornis* specimens in a polytomy comprising the holotype and the clade formed by Hesperornithes + crown birds; however, forcing all the *Ichthyornis* specimens to form an exclusive monophyletic group requires only a single additional step. All subsequent analyses were performed using a single combined OTU for *Ichthyornis*.

As discussed in the Methods, *Apsaravis* was identified as a wildcard taxon, and its phylogenetic position was remarkably variable depending on the optimality criteria used. Analyses excluding *Apsaravis* recovered much more consistent results, and therefore, these will be the results discussed below. See the Supplemental Information for results including *Apsaravis*.

Parsimony analysis using the Huang et al. dataset returned 2 most parsimonious trees with 563 steps (consistency index = 0.513, retention index = 0.798). The crownward portion of the strict consensus tree is well resolved, although bootstrap support values are relatively low overall (Fig. 28b). *Ichthyornis* is recovered stemward of Hesperornithes, and *Iaceornis* is recovered crownward of both, as the sister taxon to crown birds (Neornithes). Bayesian analysis of the Huang et al. (2016) dataset recovered an identical topology of the crownward portion of the phylogeny, with very high posterior probabilities supporting the positions of *Ichthyornis*, Hesperornithes and crown birds (Fig. 28a). Alternative topological arrangements in which *Ichthyornis* is crownward to Hesperornithes, or forms an exclusive clade with them, require 2 or 4 additional steps, respectively. Topological differences between parsimony and Bayesian analyses are more pronounced among more stemward euornitheans, but neither method recovers a monophyletic Hongshanornithidae or Songlingornithidae.

**Figure 28.**
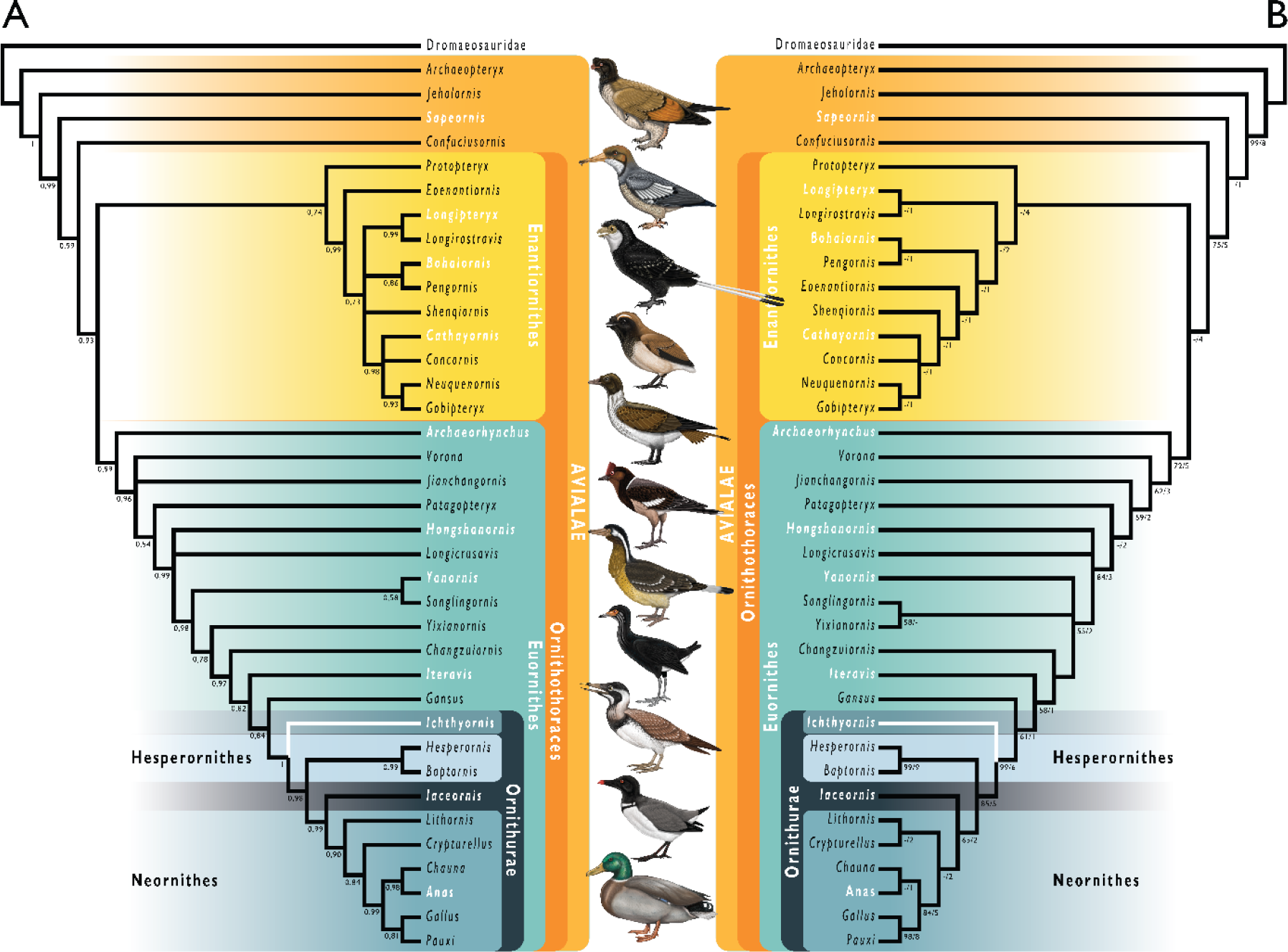
Phylogenetic results of analyses using a modified version of the morphological matrix from Huang et al. (2016), excluding *Apsaravis ukhaana*. (A) Results from Bayesian inference; node values indicate Bayesian posterior probabilities. (B) Results of parsimony analyses: strict consensus of two most parsimonious trees, branch values indicate bootstrap support values (left of bar) and Bremer decay indices (right of bar). Taxonomic names in white indicate taxa illustrated in the figure, the corresponding illustration is indicated by the superscript numbers. Branches in white indicate the position of *Ichthyornis dispar*, the focal taxon of this study. Illustrations courtesy of R. Olive, used with permission.

Parsimony analysis using the Wang et al. (2020) dataset returned 10 most parsimonious trees with 1395 steps (consistency index = 0.281, retention index = 0.661). Similar to the previous analyses, the crownward region of the tree is well resolved, but bootstrap values are low (Fig. 29b). *Ichthyornis* is recovered stemward to Hesperornithes, which emerge as the sister taxon to the crown group. Bayesian inference recovers an identical topology for the crownward portion of the tree, with very high posterior support values (Fig. 29a). Alternative topological arrangements in which *Ichthyornis* is crownward to Hesperornithes, or forms an exclusive clade with them, require 3 or 5 additional steps, respectively. Relationships among stemward euornitheans differ substantially between the parsimony and the Bayesian inference results, with Bayesian analyses recovering *Vorona* and *Patagopteryx* in an exclusive clade, as well as a monophyletic Songlingornithidae, despite recent iterations of this dataset recovering a monophyletic Yanornithidae instead (Wang et al., 2020b, 2021). Notably, the position of *Iteravis* varies under alternative optimality criteria, emerging stemward of *Gansus* in the parsimony analysis but within Songlingornithidae under Bayesian inference.

**Figure 29.**
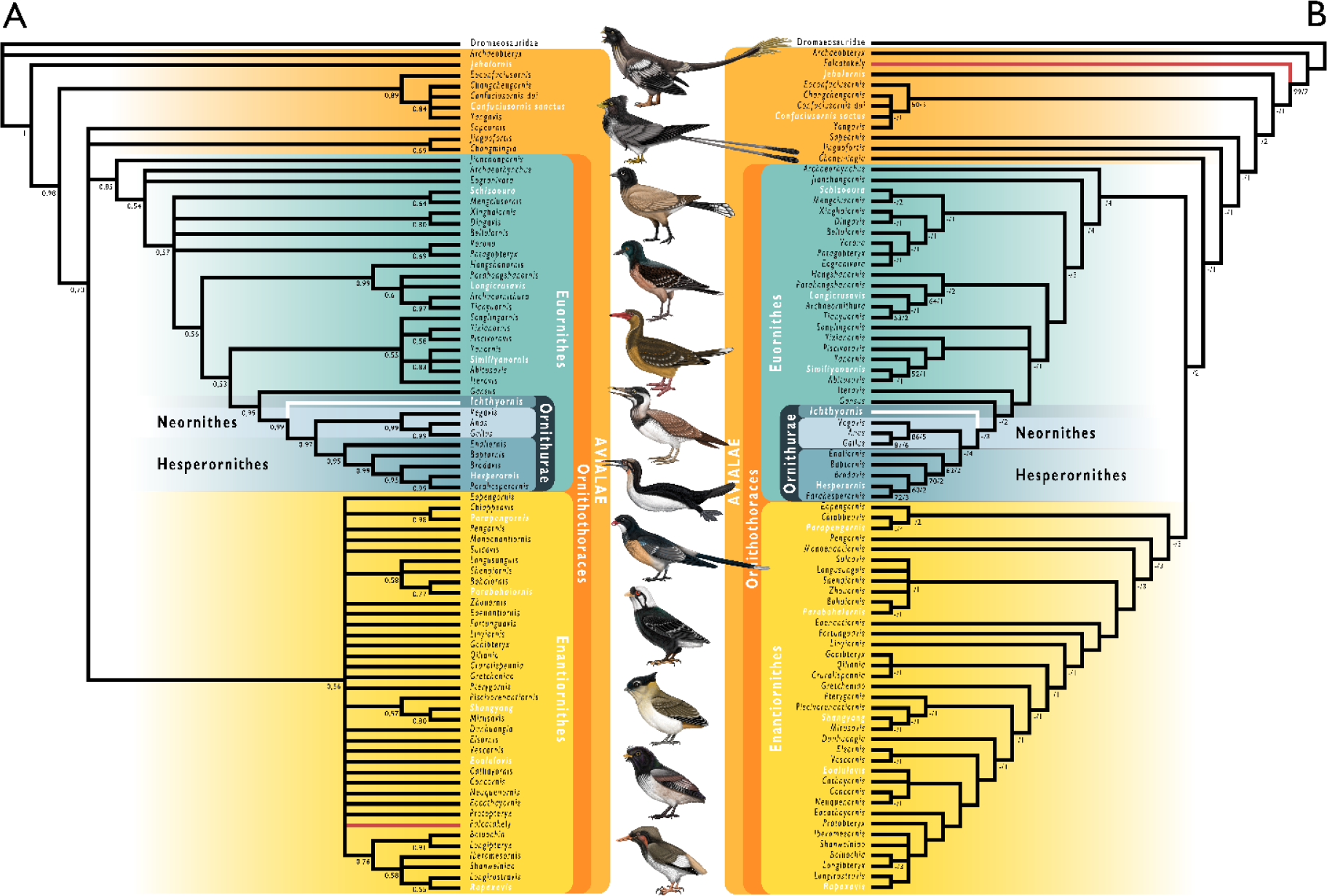
Phylogenetic results of analyses using a modified version of the morphological matrix from Wang et al. (2020), excluding *Apsaravis ukhaana*. (A) Results from Bayesian inference; node values indicate Bayesian posterior probabilities. (B) Results from parsimony analyses; strict consensus of ten most parsimonious trees, branch values indicate bootstrap support values (left of bar) and Bremer decay indices (right of bar). Taxonomic names in white indicate taxa illustrated in the figure. Branches in red indicate the position of *Falkatakely forsterae*, highlighting remarkable lability in its phylogenetic position recovered from analyses employing different phylogenetic optimality criteria. Branches in white indicate the position of *Ichthyornis dispar*, the focal taxon of this study. Illustrations courtesy of R. Olive, used with permission.

### BODY MASS ESTIMATES

We estimated the total body mass for several of the new *Ichthyornis* specimens following the equations from Field et al. (2013). Several different measurements were taken for these estimates (Fig. 30), providing a total mass distribution for our sample between 104.43 g and480.67 g. As reported by Field et al. (2013), these measurements recover estimates with variable accuracy, and measurements of the coracoid provide particularly precise estimates of volant bird body mass. The coracoid happens to be the most commonly preserved skeletal element in our sample, and measurements of maximum diameter of the coracoid humeral articulation facet, maximum coracoid lateral length, and the coracoid shaft least width could be obtained from ten, five and nine specimens respectively (Table 4). Maximum humeral length could be taken for seven specimens, although it this proxy recovered systematically higher mass estimates than the coracoid measurements, suggesting that the humeri of *Ichthyornis* were proportionally long (Fig. 30; Table 6). In contrast, the total tarsometatarsal length, which could only be measured for FHSM VP-18702 (Table 15), recovered a mass estimate notably smaller than those derived from any other measurement. However, tarsus length is highly ecologically variable, making it a particularly poor predictor of volant bird body mass (Field et al. 2013); as such, we considered this estimate unreliable. Where specimens preserved both right and left skeletal elements, recovered mass estimates were averaged (Fig. 30). In certain cases, specimens preserving both paired elements produced notably different body mass estimates, such as BHI 6420 and NHMUK A 905. Both of these specimens are flattened, and the differing estimates might be caused by taphonomic deformation, although the possibility that these specimens could represent composites of multiple individuals cannot be ruled out.

**Figure 30.**
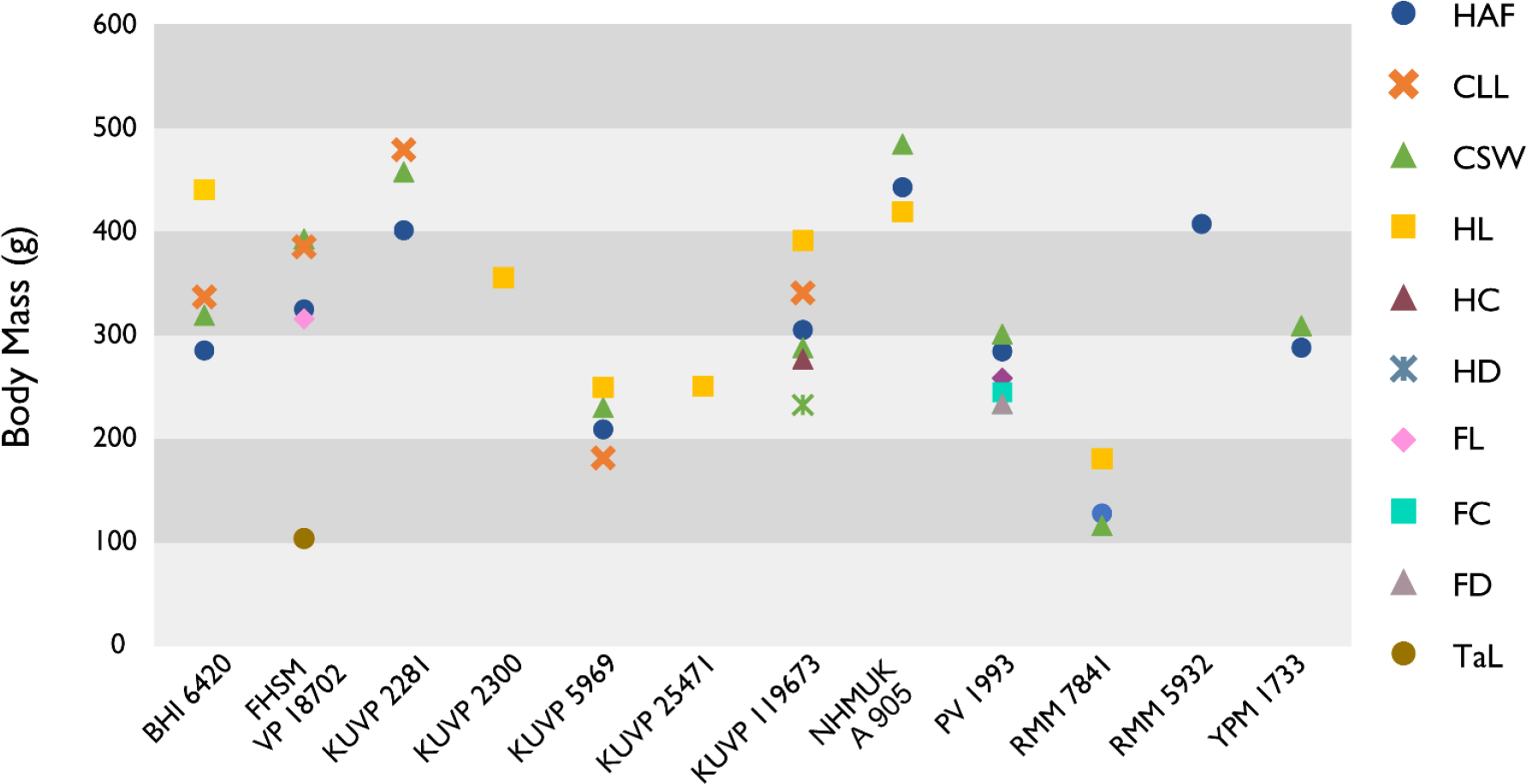
Body mass estimates for selected *Ichthyornis* specimens. Body mass correlates represented, in order of increasing Percent Prediction Error (after Field et al., 2013) are: maximum diameter of the coracoid’s humeral articulation facet (HAF), maximum coracoid lateral length (CLL), least coracoid shaft width (CSW), maximum humerus length (HL), least humerus shaft circumference (HC), least humerus shaft diameter in cranial view (HD), maximum femur length (FL), least femur shaft circumference (FC), least femur shaft diameter in cranial view (FD) and maximum tarsometatarsus length (TaL).

Overall, the body mass estimates provide a relatively wide mass distribution for *Ichthyornis*, with the largest specimens being between 2.5 and 4.2 times the size of the smallest, depending on the measurements taken. NHMUK A 905 (4,26.64 g to 486.18 g) and KUVP 2281 (402.65 g to 459.27 g) were recovered as the largest individuals, while KUVP 5969 (182.48 g to 250.80 g) and RMM 7841 (117.15 g to 181.79 g) were the smallest. As discussed above, the studied specimens exhibit probable evidence of ontogenetic variation, particularly in the case of KUVP 5969, which might represent a juvenile individual; as such, the small body size estimate for these specimens may be indicative of an early ontogenetic stage.

## DISCUSSION

### NOVEL MORPHOLOGICAL INFORMATION

This work reveals the preservation of several skeletal elements previously unknown for *Ichthyornis*, as well as considerable novel morphological information for skeletal elements previously represented only by fragmentary or poorly preserved specimens. Amongst the new material, the vertebral series stands out as especially significant, as only a few isolated vertebrae have been previously described (Marsh, 1880; Clarke, 2004). Only three YPM specimens, YPM 1450, 1732 and 1733, preserve any significant vertebral material, but in all cases, the presacral and caudal vertebrae are isolated and fragmentary. As described above, three of the new specimens, KUVP 25472, KUVP 119674, and ALMNH PV93.2.133, preserve a substantial portion of the vertebral column, revealing in unprecedented detail the vertebral morphology of *Ichthyornis*. While the incomplete nature of the specimens currently makes it impossible to establish the total number and exact position of all vertebrae, the preservation of the partial cervical series in KUVP 119673 and the almost complete thoracic series in KUVP 25472 allow for a much more precise estimate than was previously possible. These specimens illustrate that *Ichthyornis* had at least 10 cervical and 10 thoracic vertebrae (Fig. 4, 5 & 6), a count similar to those of closely related taxa such as *Gansus* (Wang et al., 2016b) and *Yixianornis* (Clarke et al., 2006). The vertebral morphology of the new specimens is very similar to that described by Marsh (1880) and Clarke (2004), although the new specimens provide a better characterization of the cervicothoracic transition.

The synsacrum of *Ichthyornis* was previously known from only three YPM specimens, which notably diverged in the number of fused sacral vertebrae, exhibiting either 10 or 12 ankylosed sacrals Clarke (2004). Here, the synsacrum is preserved in four of the new specimens, and three of them preserve the element in its entirety (Fig. 7). As described above, the new material reveals a pattern of vertebral fusion not dissimilar to that exhibited by some extant birds, such as *Gallus gallus* and *Ardea cinerea*, during post-hatching ontogeny (Bui & Larsson, 2021; Watanabe, pers. comm.). The new specimens show the presence of variably ankylosed sacral vertebrae on the anterior end of the sacrum, and the fusion of additional vertebrae to the caudal end of the element. While this variability is presumably ontogenetic, there seems to be no close correlation between the size of the specimens and the degree of fusion they exhibit, recalling similarly idiosyncratic fusion patterns across the skeleton in other Mesozoic dinosaurs (Longrich & Field, 2012; Bailleul et al., 2016, Griffin et al., 2021), and potentially pointing towards greater intraspecific or interspecific variation than previously recognized (see discussion of intraspecific variation below). Two of the new specimens preserve caudal vertebrae, although these do not reveal significant new morphological information, and no pygostyle is preserved among the studied material. This element remains unknown for *Ichthyornis*.

The ribs of *Ichthyornis* have not been previously described in detail. Both vertebral and sternal ribs are preserved in the new specimens, and, although well preserved in several instances, none appear to be entirely complete (Fig. 8). Considering this abundance of costal material, the absence of any preserved uncinate processes is striking. The sternum is preserved in three of the new specimens, and although all of them are variably distorted, together they provide an unprecedented level of information on the sternal morphology of *Ichthyornis* (Fig. 9), our understanding of which was previously based on very incomplete remains (Fig. 10). The caudal portion of the sternum was previously very poorly known, and the new specimens, particularly KUVP 119673, reveal the presence of short, rounded, and open sternal incisures, dissimilar to those of other crownward euornitheans such as *Gansus* (Wang et al., 2016b). The new specimens also provide the first detailed view on furcular morphology in *Ichthyornis*, as only fragments of this element have been previously reported.

Contrary to the hypothesis presented by Clarke (2004), *Ichthyornis* exhibits a clear hypocleideum extending from the clavicular symphysis (Fig. 11), a feature not previously found in any crownward euornitheans. The omal ends of the furcula also differ from the condition described by Clarke (2004), tapering omally instead of terminating in a blunt end.

Nine of the new specimens preserve one or both coracoids. In several cases, such as BHI 6420, FHSM VP-18702, KUVP 119673, and KUVP 2281, these represent the best-preserved coracoids known for *Ichthyornis*, exceeding the completeness and quality of preservation seen in any YPM specimens (Fig. 12). The new material shows that both the acrocoracoid and procoracoid processes were more strongly developed and hooked than previously recognized, together forming a claw-like shape, although these are broken in multiple specimens (Clarke, 2004). The scapula is preserved in five of the new specimens, which exhibit surprising variation in the morphology of the acromion process, the extremely reduced morphology of which was previously considered a diagnostic feature of *Ichthyornis* by Clarke (2004). Although the acromion is always small, it is variably absent, rounded, pointed and triangular, or hooked and recurved (Fig. 13). This variability might illustrate a greater degree of intraspecific variation than has been previously recognized (see below), and calls the diagnostic utility of this character for *Ichthyornis* into question.

The humerus is the most commonly reported skeletal element for *Ichthyornis*, although all previously described specimens are variously flattened. Despite the abundance of previously described *Ichthyornis* humeri (Marsh 1880; Harrison, 1973; Olson, 1975; Lucas & Sullivan, 1982; Fox, 1984; Clarke, 2004; Porras-Muzquiz et al., 2014; Shimada & Wilson, 2016), the twelve new specimens described here that bear humeri provide a wealth of novel morphological data (Fig. 14). Particularly remarkable is the three-dimensional preservation of two humeri, the left humerus of KUVP 2300 and the right humerus of KUVP 119673, which reveal a craniodorsally oriented deltopectoral crest (Fig. 15), and a curved, sigmoidal humeral shaft, previously only reported by Shimada & Wilson (2016), who attributed this novel shape to either taphonomic distortion of the specimen or interspecific variation. The preservation of both a flattened left and a three-dimensional right humerus in KUVP 119673 reveal that these important features are genuine and easily obscured by taphonomic flattening, which emphasizes the need for caution in interpreting the morphology of flattened avialan fossils. In contrast to the exceptionally preserved humeri described here, neither the ulna nor the radius is especially well represented among the new specimens, and no previously undescribed morphological details of these elements are discernible. The free carpal elements are well represented amongst the new material, and we describe the radial carpal bone of *Ichthyornis* for the first time as well as the first complete ulnar carpals (Fig. 19). The morphology of the radial carpal is similar to that of *Iaceornis* and is reminiscent of the radial carpal of the stem-anseriform *Presbyornis* (pers. obs.).

Although several of the new specimens preserve carpometacarpi, the carpometacarpus of YPM 1714, previously described by Marsh (1880) and Clarke (2004), remains the best-preserved *Ichthyornis* carpometacarpus known to date (Fig. 20). Here, the extremely short carpometacarpus of the large individual FHSM VP-18702 confirms a pattern of variation in carpometacarpus length among otherwise similarly sized and proportioned individuals, as initially noted by Clarke (2004). The morphology of the manual phalanges preserved amongst the new material is virtually identical to that of previously described *Ichthyornis* material (Fig. 21 & 22), with the exception of the proximal phalanx of the major digit of KUVP 5969, which lacks an internal index process (see intraspecific variation below). The exceptional level of preservation of the new specimens may facilitate a detailed investigation of the muscular and ligament attachments of the hand in future work.

Only one of the new specimens, KUVP 119673, preserves pelvic remains, but these constitute the most complete pelves known for *Ichthyornis* (Fig. 23). Although partially complete pelves in anatomic connection were previously known from YPM 1732, significant portions of the ilium and pubis were missing, and their anatomy was inferred in illustrations by Marsh (1880). The well preserved pelves of KUVP 119673 show a markedly different morphology from that reconstructed by Marsh (1880), with narrow and blade-like postacetabular ilia similar to those of other crownward euornitheans such as *Gansus* (You et al., 2006; Wang et al., 2016b).

Hindlimb material for *Ichthyornis* is much scarcer than forelimb material within the YPM collections, and the same is true for the new specimens. Only one specimen, ALMNH PV93.2.133, preserves a mostly complete, three-dimensional hindlimb, although multiple other specimens include fragmentary tibiotarsi and tarsometatarsi. The femur of ALMNH PV93.2.133 is three-dimensionally preserved and constitutes the best-preserved femur of *Ichthyornis* known so far (Fig. 24). Importantly, discrete features of its morphology do not differ substantially from those described by Marsh (1880) and Clarke (2004) based on taphonomically flattened YPM material. The proximal tibiotarsus of ALMNH PV93.2.133, although fragmentary and somewhat distorted, preserves several features not previously observed in *Ichthyornis* (Fig. 25). Two of the new specimens preserve fibulae, and although these are poorly preserved, they represent the first fibulae described for the taxon (Fig. 25d). Multiple specimens include well-preserved tarsometatarsi, and among them FHSM VP-18702 is particularly remarkable for exhibiting the first complete *Ichthyornis* tarsometatarsus (Fig. 26). The excellent preservation of the tarsometatarsi of ALMNH PV93.2.133 and MSC 13214 enables for the first time the description of a rudimentary hypotarsus, which illustrates a remarkable combination of stem-and crown bird-like morphological features. The pedal phalanges of *Ichthyornis* are described in detail for the first time (Fig. 27); although it is impossible to infer their position with certainty, precluding a complete reconstruction of foot anatomy for *Ichthyornis*, the proportions of these phalanges in comparison with the rest of hindlimb elements suggest a greatly enlarged pes, presumably adapted for foot-propelled swimming.

Together, the 40 new specimens reported here provide us with the most complete view of *Ichthyornis* postcranial morphology to date, greatly updating our understanding of this taxon. This new material allows an almost complete reconstruction of the *Ichthyornis* postcranial skeleton (Fig. 31), making *Ichthyornis* one of the most complete and best-preserved Mesozoic avialans known to date (Brocklehurst et al., 2012; Pittman et al., 2020b). In addition to their completeness, many *Ichthyornis* fossil remains are notable for their exceptional three-dimensional preservation, a condition rarely seen among Mesozoic avialans, which are usually flattened, crushed, or otherwise distorted (Field et al., 2018a; Hu et al., 2020). The unprecedented completeness and quality of these remains should allow for extensive reconstructions of soft tissue, and explorations of functional morphology and biomechanics at a level of detail previously impossible for crownward Mesozoic avialans.

**Figure 31.**
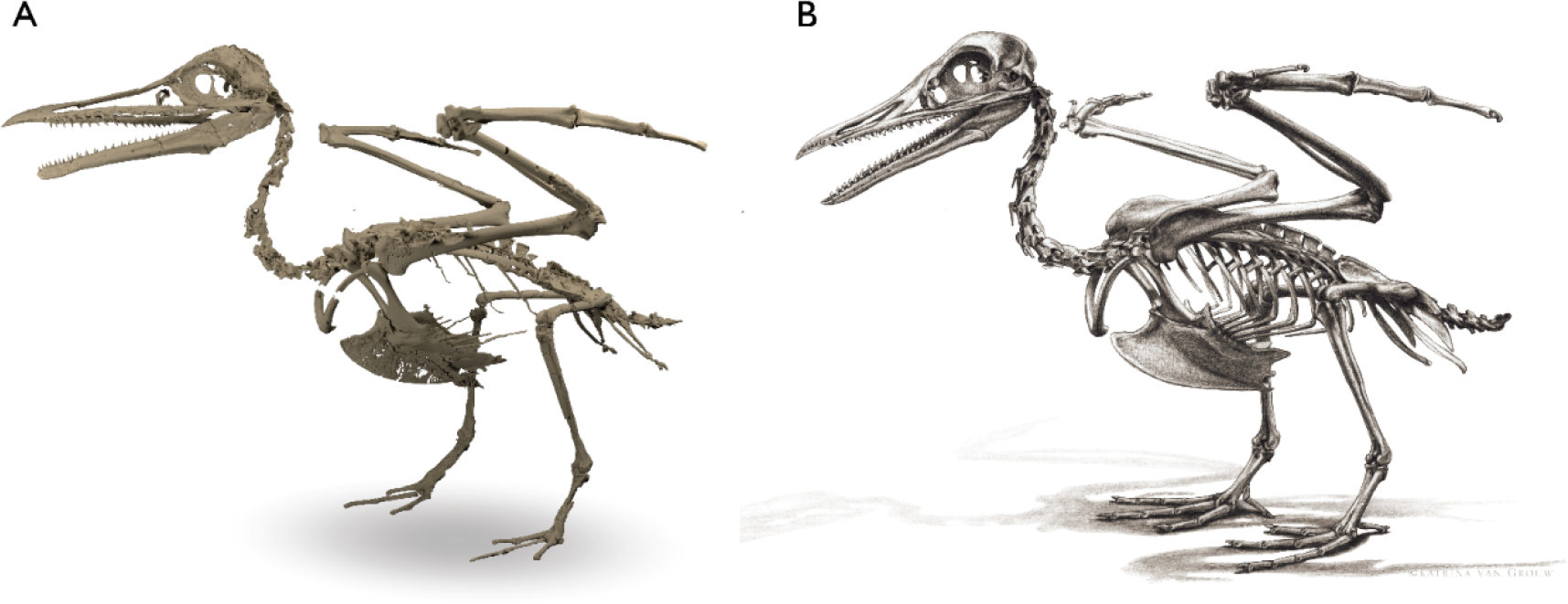
Skeletal reconstructions of *Ichthyornis dispar*. (A) three-dimensional skeletal reconstruction using a composite of the specimens described in this study; (B), complete skeletal reconstruction of *Ichthyornis dispar*, including reconstructed fragmentary and missing elements, courtesy of K. van Grouw, used with permission.

Although reconstructing the ecology and lifestyle of *Ichthyornis* is not the primary aim of this work, the new material described here enables some new inferences into its biology.

*Ichthyornis* shows similar wing proportions to volant extant birds, especially marine soaring taxa. The brachial index (BI, *sensu* Nudds et al., 2006) of *Ichthyornis* could only be measured for KUVP 5969 and KUVP 119673, since these are the only specimens preserving both complete humeri and ulnae. In KUVP 119673, the BI is 0.94, similar to taxa such as terns (e.g., *Sterna hirundo*; 0.89), gulls (e.g., *Chroicocephalus novaehollandiae*; 0.89), shearwaters (e.g., *Puffinus lherminieri*; 0.99) and tropicbirds (e.g., *Phaethon lepturus*; 0.92) (see Supplemental Information). In contrast, in KUVP 5969 the BI is 1.05, more similar to the proportions of non-marine taxa such as geese (e.g., *Anser albifrons*; 1.04) and coots (e.g., *Fulica atra*; 1.01). As discussed above, KUVP 5969 might represent a juvenile individual, and its distinct proportions, with a humerus longer than the ulna, might be related to its early developmental stage.

None of the studied three-dimensional *Ichthyornis* forelimb elements show any evidence of the flattened morphology observed in specialized wing-propelled divers (Habib, 2010; Watanabe et al., 2021). Instead, *Ichthyornis* appears to exhibit several hindlimb features related to swimming, and several anatomical features may point towards it having exhibited a paddle-swimming ecology. These include a reduced femoral trochanter, a character state associated with foot propelled-diving (Zinoviev, 2011; Clifton et al., 2018; Bell et al., 2019) and observed in taxa such as *Anas platyrhynchos*, *Puffinus lherminieri*, *Phaethon lepturus*, *Phalacrocorax carbo* and *Morus bassanus*, but absent in non-natatorial taxa such as *Sterna hirundo* or *Charadrius rubricollis*, in land-dwelling birds such as *Gallus gallus* and wing-propelled swimmers such as *Alca torda*. While the leg proportions of *Ichthyornis* cannot be completely reconstructed due to a lack of entirely complete tibiotarsi, the extremely short tarsometatarsus and elongated pedal phalanges are similar to the proportions observed in foot-propelled swimmers like *Anas platyrhynchos* and *Phalacrocorax carbo*. Only the ecological preferences of a few crownward euornitheans have been assessed in detail, particularly those of *Gansus* (You et al., 2006; Li et al., 2011; Nudds et al., 2013) and the extremely specialized diving Hesperornithes (Bell et al., 2019). In the case of *Gansus,* which is recovered in a phylogenetic position just stemward of *Ichthyornis* in the present study, the interpretation of its ecology is complicated by a unique combination of characters, with aspects of its forelimb morphology similar to Apodiformes, but hindlimb proportions reminiscent of aquatic taxa as Podicipedidae or Phalacrocoracidae (Nudds et al., 2013). The extremely elongated pes of *Ichthyornis* is similar to that of *Gansus*, but the rest of the hindlimb elements are proportionally much shorter in *Ichthyornis*, particularly the tarsometatarsus, resulting in a pes that would have been proportionally much larger. Understanding exactly how this hindlimb anatomy would have related to swimming capabilities will require a more in-depth functional investigation, but these observations minimally lend support to the interpretation of *Ichthyornis* as capable foot-propelled swimmer and marine soarer.

### MORPHOLOGICAL VARIATION IN *ICHTHYORNIS*

Although eight different species of *Ichthyornis* have been previously named, five of these were synonymized into *I. dispar* by Clarke (2004), while two were split into the separate genera *Guildavis* and *Apatornis*. Clarke (2004) found no basis for distinguishing different species within the genus *Ichthyornis*, though recognized a considerable range of size variation and moderate morphological disparity among the YPM *Ichthyornis* specimens. The substantial degree of variation within the YPM sample and the incomplete nature of much of the material made it impossible to rule out the hypothesis of considerable intraspecific variation, thus, there was no firm basis for the separation of *Ichthyornis* into distinct species-level taxa.

Within the YPM collections, only two specimens are smaller than the holotype (YPM 1450), and most of the YPM sample (85.7%) includes specimens substantially larger than the holotype (Clarke, 2004). Clarke illustrated a continuous size distribution among the YPM specimens on the basis of humeral and ulnar measurements, illustrating the absence of clearly distinguishable discrete size classes (Clarke, 2004). Among the specimens included in the current study that preserve complete humeri, 62.5% represent specimens with a substantially greater humeral length than the *Ichthyornis* holotype (with their humeri being at least 90% or more of the total humeral length of the largest known individual), 25% are of a similar size to the holotype, and only one specimen (RMM 7841) is notably smaller than the holotype (Table 6). One of the new specimens, BHI 6420, preserves what appears to be the longest *Ichthyornis* humerus recorded so far, exceeding the length of the largest previously recorded individual, YPM 9685 (see description). Humerus length provides a reasonably, though not exceptionally, accurate proxy for body mass among extant flying birds (Field et al., 2013; see Body Mass Estimation section), and the humeral size distributions discussed above are matched by similar size patterns for all of the other long bones examined (see measurements in Supplemental Information).

Fossil remains identified as belonging to *Ichthyornis* have been described from a stratigraphic interval spanning over 10 million years, and Clarke (2004) observed size differences apparently related to stratigraphic provenance. The oldest specimens among the YPM material, from the Cenomanian (approximately 95 MYA), seem to have been amongst the largest described, but the younger specimens from the Turonian (approximately 93.9 MYA) and the Coniacian (approximately 89.8 MYA) are substantially smaller, although it is unclear whether these observations reflect an evolutionary anagenetic signal. In contrast, a broad size distribution apparent among specimens deriving from the Santonian (approximately 86.3 MYA) and the Campanian (approximately 83.5 MYA), the interval from which the holotype of *I. dispar*, the majority of the YPM material, and all of the specimens described here derive. Similarly-sized individuals are found among the specimens deriving from the Santonian Smoky Hill Member of the Niobrara Formation and the Campanian Mooreville Formation, and both very large (BHI 6420, KUVP 2281, NHMUK A905, RMM 5932) and very small (KUVP 5969, RMM 7841) specimens are known from both localities. See the Body Mass Estimation section below for detailed information on the size distribution of the specimens described here.

As discussed above, most of the new specimens described in this work were referred to *Ichthyornis* based on the presence of nine unambiguous diagnostic features previously put forward by Clarke (2004), although we observe a previously unappreciated amount of variation for a selection of these characters. In cases where diagnostic features were not preserved, our referrals were based on morphological similarity to previously published *Ichthyornis* material (Table 1). Despite some morphological variation within our sample, when several of the most complete specimens (BHI 6420, 6421, FHSM VP-18702, KUVP 2281, KUVP 5969, KUVP 25469, KUVP 25472, KUVP 119673, NHMUK A 905 and ALMNH PV93.2.133) were included as distinct operational taxonomic units in our phylogenetic analyses, they all formed a clade with the *I. dispar* holotype, YPM 1450, supported by strong node support values across alternative optimality criteria (see Phylogenetic Analysis section). All specimens formed a polytomy within this clade, and no consistent internal relationships were recovered among the different specimens included in the analyses. On the basis of these analyses, we considered it appropriate to refer all of these specimens to *I. dispar*.

As detailed above, the morphology of the acromion process of the scapula—one of the autapomorphies of *I. dispar* suggested by Clarke (2004)—shows a greater degree of variation among the new specimens than had previously been observed in *Ichthyornis*, with only KUVP 2281 and KUVP 119673 exhibiting the extremely diminutive and pointed acromion supposedly diagnostic for *I. dispar*. In contrast, several specimens show differing morphologies that do not seem to be related to taphonomic distortion, such as a longer hooked process in FHSM VP-18702, a larger and more globose acromion on BHI 6420 and the right scapula of NHMUK A 905, and an essentially absent, undeveloped acromion on the left scapula of NHMUK A 905. The oval scar described by Clarke (2004) as present on the distal radius of *I. dispar* is only faintly visible in FHSM VP-18702, KUVP 25469, and KUVP 25472, and it is conspicuously absent in BHI 6420 and KUVP 119673, despite the excellent preservation of these specimens. Clarke (2004) discussed variability in the position and development of this character that might be related to size, with the scar apparently being fainter in smaller individuals. However, this does not seem to hold true among the new specimens evaluated here, particularly for BHI 6420, which appears to be among the largest *Ichthyornis* individuals recorded. Lastly, the presence of an internal index process—another possible autapomorphy of *I. dispar*—is observable in all the specimens preserving the first phalanx of the major manual digit, even when broken (as in RMM 6201), with the clear exception of KUVP 5969, in which this process is undeveloped (discussed above). We consider it most likely that the unusual morphology of KUVP 5969 is related to its small size and probable early ontogenetic stage, as this feature is absent in early ontogenetic stages of some extant marine birds such as *Macronectes giganteus*. Alternatively, the complete lack of an internal index process might point towards KUVP 5969 belonging to another species closely related to *I. dispar*.

Beyond this variability in the diagnostic characters of *Ichthyornis dispar*, the specimens included in this study show additional morphological variation. Excluding variability that might be attributed to preservational factors, such as the curvature of the humeral shaft or the orientation of the deltopectoral crest, two of the elements exhibiting the most striking differences among individuals are the synsacrum and the carpometacarpus. As discussed in the description, the four preserved synsacra show an apparent pattern of vertebral fusion similar to that of certain extant bird groups, which might point towards these specimens representing different ontogenetic stages. However, this does not seem to correlate well with size, with smaller specimens like ALMNH PV93.2.133 showing a greater degree of fusion and a larger number of postacetabular sacral vertebrae than the larger KUVP 119673, and a comparable degree of fusion to that of the much larger KUVP 25469. Additionally, some individuals of similar size, such as FHSM VP-18702 and KUVP 119673, exhibit differing degrees of synsacral fusion. A comparable pattern in which larger specimens were noted to exhibit morphology consistent with a relatively early ontogenetic stage was previously noted by Clarke (2004), with reference to the axis of YPM 1775, the mandible of YPM 1735, and the ulna of YPM 1740. The extent to which this mismatch between size and apparent levels of ontogenetic maturity reflects intraspecific or interspecific variation is challenging to assess at present, but recalls similar patterns observed in other groups of fossil and extant archosaurs (Bailleul et al., 2016; Griffin et al., 2021), and patchy stratigraphic sampling precludes our ability to evaluate any potential anagenetic patterns. Notable variation in adult size has been previously inferred to be ancestral to archosaurs, and documented in many stem-birds (Sander & Klein, 2005; Hone, 2016; Griffin & Nesbitt, 2016; Carr, 2020; Chapelle et al., 2021). Whether *Ichthyornis* exhibited comparable variability in adult size is difficult to establish, but this interpretation provides a possible explanation for the considerable variation in size and morphology we observe, despite the near-crown position of *Ichthyornis* and its similar growth patterns to extant birds (Chinsamy et al., 1998).

As discussed above, the basis for recognizing both *Apatornis* and *Guildavis* as distinct from *Ichthyornis* rests solely on sacral morphology, as no other skeletal remains are associated with those taxa Clarke (2004). The new specimens, which despite their differing synsacral anatomy are otherwise diagnosable as *I. dispar* on the basis of additional skeletal material, raise the distinct possibility that both *Guildavis* and *Apatornis* might fall within the range of variation of *Ichthyornis*. The validity of *Apatornis* and *Guildavis* may therefore necessitate future reevaluation.

The morphology of the carpometacarpus is generally consistent among the new specimens, with the exception of that of FHSM VP-18702, which exhibits a markedly smaller carpometacarpus than that seen in similarly sized individuals such as BHI 6420. A similar observation was reported by Clarke (2004) for a single individual, YPM 1755, which exhibits a shorter carpometacarpus than other specimens of similar size.

Of all the specimens included in this study, KUVP 5969 is the most morphologically distinct, exhibiting an intriguing number of differences with respect to the other individuals. In addition to the absence of an internal index process as noted above, the glenoid facet of KUVP 5969 lies further towards the omal end of the coracoid with respect to the scapular cotyle than in any other specimen, with the position of the sternal end of the glenoid omal to that of the scapular cotyle. Although the coracoid of KUVP 5969 is small and relatively poorly preserved, this feature does not seem to be related to taphonomic distortion of the element, and is not present on the even smaller coracoid of RMM 7841. KUVP 5969 also shows a proportionally short humerus; indeed, it is the only specimen in which the humerus is shorter than the ulna (Tables 6 and 7). The humerus tends to be particularly elongated in soaring marine birds, and although its proportional length varies during ontogeny, its longitudinal growth seems to stop by the time of fledgling (Watanabe, 2017, 2018). However, a similar condition has been observed in the stemward euornithean *Archaeorhynchus*, in which at least one early-stage juvenile specimen shows an ulna that is shorter than the humerus, with it being subequal or longer in other later stage subadult and adult specimens (Foth et al., 2021). The radial carpal of KUVP 5969 is poorly preserved, but its morphology differs from that of other specimens as well, with a practically straight ventral ramus showing very little curvature. The intriguing suite of morphological differences exhibited by KUVP 5969, which is otherwise similar to other *Ichthyornis* specimens, might be related to its relatively small size (it is the second-smallest of the new specimens). However, at present it is unclear whether these differences might be related to a relatively early ontogenetic stage (with KUVP 5969 representing a rare juvenile individual), or whether they might be related to further intraspecific or interspecific variation. Unfortunately, with the exception of the coracoid of RMM 7841, other specimens of comparably small size are too fragmentary to evaluate whether they show a similar morphology to that of KUVP 5969.

The level of morphological variability within the *Ichthyornis* specimens described here is moderate overall, and most individuals exhibit congruent morphologies. For all of the features noted to exhibit some degree of variation, such variation was only observed in single specimens, complicating the study of broader patterns of variation within *Ichthyornis*. Neither the variable features noted here nor those identified by Clarke (2004) appear to exhibit any correlation with temporal or geographic provenance, and size differences appear to be the major factor influencing morphological variability in most cases, other than in the synsacrum. As a consequence, it is not currently possible to establish whether any of these morphological differences reflect interspecific variation, and we follow Clarke (2004) in considering that there is no basis for the establishment of distinct *Ichthyornis* species at present. The use of quantitative techniques such as geometric morphometrics has previously been used to evaluate the degree of intraspecific variation in fossil species, and to discern between the variability caused by taphonomic factors and that related to plausible intra-and interspecific biological variability (Hedrick et al., 2019; Lefebvre et al., 2020). Such an approach might be applicable in future studies in order to evaluate potential patterns of interspecific and intraspecific variation in *Ichthyornis*.

### PHYLOGENETIC POSITION

As discussed above, despite the presence of several morphological differences among the new *Ichthyornis* specimens, all of them were recovered very close to the *Ichthyornis* holotype YPM 1450 in phylogenetic analyses that treated the new specimens as distinct operational taxonomic units. In these analyses, the synapomorphies supporting an *Ichthyornis* clade corresponded to the features previously highlighted as diagnostic for *Ichthyornis* by Clarke (2004). These included features like amphicoelous vertebrae, the shape of the distal ulna, and the presence of an internal index process on the II:1 manual phalanx, but also include several features of the skull and lower jaws, the presence of crossed coracoid sulci in the sternum, the hooked acrocoracoid process, the elongated scapula, the shape of the humeral deltopectoral crest and the short femoral trochanteric crest (see Supplemental Information for the complete list of synapomorphies).

All of our phylogenetic analyses recovered a similar topology for the crownward portion of the tree, regardless of the dataset or method used (see Figs. 28 and 29 for the main results from Bayesian and Parsimony analyses, and Supplemental Information for additional results). In all cases, *Ichthyornis* was recovered in a position stemward of Hesperornithes, a position previously inferred on the basis of iterations of the Wang et al. (2020) dataset (O’Connor et al., 2011, 2020; Wang et al. 2017, 2019; Atterholt et al., 2018; Field et al., 2018a), but never previously recovered from analyses using previous versions of the Huang et al. (2016) dataset, which have always recovered *Ichthyornis* closer to the avian crown group (Chiappe, 2002; Clarke, 2004; You et al., 2006; Huang et al,. 2016; Torres et al. 2021).

Notably, two recent iterations of the Wang et al. dataset recovered *Ichthyornis* crownward to Hesperornithes (Wang et al., 2020b) or in a polytomy with Hesperornithes and the crown bird group (Wang et al., 2021). The more crownward position of Hesperornithes recovered in our analyses appears to be well supported by several synapomorphies absent in *Ichthyornis*, including completely heterocoelous cervical vertebrae, pneumatic foramina piercing the centra of the mid-cranial cervicals, the narrow shape of the sternal rostrum, non-tapering tibiotarsal condyles, and several skull and lower jaw characters previously discussed by Field et al. (2018a).

Notably, *Iaceornis* is also recovered crownward to both *Ichthyornis* and Hesperornithes in analyses employing the updated Huang et al. (2016) dataset, despite its remarkable morphological similarity to *Ichthyornis* (Marsh, 1880; Clarke, 2004). None of the synapomorphies recovered for the Hesperornithes + *Iaceornis* + crown group clade are preserved in *Iaceornis*, from which no cranial or axial material are yet known. Only two synapomorphies are uniquely shared between *Iaceornis* and the crown group: the presence of raised intermuscular ridges in the sternum and a large and projected extensor process on the carpometacarpus. The crownward position of *Iaceornis* in relation to both Hesperornithes and *Ichthyornis* highlights the need for a more detailed characterization of its morphology in future work.

The monophyly of Ornithurae (see clade definitions) is well supported by several apomorphies. These include the presence of costal facets on the sternum, clear caudal or caudolateral sternal processes instead of closed fenestrae, a relatively elongate extensor process of the carpometacarpus and an intermetacarpal process present as a scar, the presence of a flexor process on the III:1 manual phalanx, subparallel pubes and ischia, the presence of a renal fossa on the postacetabular illium and the presence of two proximal vascular foramina on the tarsometatarsus. Several of these morphological features were previously recovered as synapomorphies of Neornithes, particularly those related to the shape of the sternum and the pelvis (Clarke, 2004; Huang et al., 2016; Wang et al., 2017, 2019). These structures were previously poorly preserved in *Ichthyornis,* and mostly absent in Hesperornithes due to their extremely derived and specialized anatomy (Bell & Chiappe, 2016, 2020). The presence of these features in *Ichthyornis* therefore pushes the inferred origin of these features to the ancestral lineage subtending Ornithurae.

The topology of the stemward portion of euornithean phylogeny is less stable between alternative analyses using both datasets, although major clades such as Hongshanornithidae are mostly consistently recovered. Notably, the recently defined clade Yanornithidae is not recovered using the Wang et al. dataset despite being supported in recent iterations of this matrix (Wang et al., 2020b; 2021), and instead Songlingornithidae, including *Yixianornis* and *Songlingornis* is recovered. *Iteravis*, traditionally found crownward of these taxa, is recovered as part of Songlingornithidae for the first time in our Bayesian analyses using the Wang et al. (2021) dataset, although the posterior probabilities supporting this clade are low. Despite its exclusion from these analyses due to its unstable position, the inclusion of *Apsaravis* has a notable impact on the tree topologies we recover (see Supplemental Information). When included, *Apsaravis* is recovered either as a stemward euornithean, in a polytomy with Hongshanornithidae, or very close to crown birds, either slightly stemward or just crownward of *Ichthyornis* (see Supplemental Information), due to an unusual combination of character states (Clarke & Norell, 2002). We suggest that this taxon is in need of a detailed reinvestigation in order to further understand its unusual morphology, especially in light of its remarkable state of preservation.

Although the study of enantiornithine phylogenetic relationships is beyond the scope of this study, it is worth highlighting the extremely variable position of *Falcatakely*, which was originally recovered as an enantiornithine by O’Connor et al. (2020) in a Bayesian analysis using a previous iteration of the Wang et al. (2020) dataset. Our analyses recover an identical position under Bayesian inference, but when our dataset is analyzed under parsimony, *Falcatakely* is recovered as a non-ornithothoracean avialan, just crownward of *Archaeopteryx* (Fig. 28b). Obviously, more complete skeletal material will be necessary to fully resolve the phylogenetic position of this intriguing Maastrichtian taxon (Field, 2020).

## CONCLUSION

This study has reevaluated the postcranial morphology of the crownward avialan *Ichthyornis* on the basis of 40 new fossil specimens, making it among the most substantial additions to our knowledge of non-neornithine avialan postcranial morphology. Much of the new material is exceptionally well-preserved, including multiple elements preserved in three-dimensions and illustrating previously unknown impressions of soft tissue attachments. We hope the novel observations and data provided here will help facilitate future insights into the functional morphology of this key near-crown avialan, and a deeper understanding of the functional and anatomical origins of the crown bird skeleton.

Our work has illustrated a substantial range of morphological variation within *Ichthyornis*, including probable evidence of ontogenetic variation. Virtually every element of the postcranial skeleton of *Ichthyornis* is represented among the new specimens, providing an unprecedentedly detailed look into the three-dimensional skeletal morphology of a near-crown avialan. Indeed, the new material illustrates that much of the *Ichthyornis* postcranium falls within the range of anatomical variation of crown birds, and indicates that several features previously considered unique to the bird crown group, such as a cranially deflected deltopectoral crest or the presence of a developed hypotarsus, were present in *Ichthyornis*.

The new material described here should help resolve ongoing uncertainty regarding the relative phylogenetic placement of *Ichthyornis*, Hesperornithes, and Neornithes, with important implications for the crownward-most portion of avialan phylogeny. We illustrate that, irrespective of the phylogenetic dataset used, *Ichthyornis* resolves in a position stemward to Hesperornithes, and our results emphasize the unresolved phylogenetic positions of taxa such as *Apsaravis* and the importance of reevaluating obscure crownward avialans like *Iaceornis*. Additionally, we provide phylogenetic definitions for several key avialan subclades, and hope that these help facilitate unambiguous discourse as future work continues to resolve the systematics of Cretaceous avialans.

In light of this work, as well previous monographic descriptions by Clarke (2004) and Marsh (1880), *Ichthyornis* inarguably ranks among the most completely known fossil avialans of any age. We hope that the new anatomical and phylogenetic information provided for *Ichthyornis* in the present work provides a useful substrate for continued progress on investigations into the biology of the crownward-most portion of the avian stem group.

## ACKNOWLEDGEMENTS

We thank L. Rieppel and J. Secord for discussions on the historical significance of *Ichthyornis*, M. Wills for helpful guidance, D. Ehret, J. Ebersole, M. Lowe, D. Brinkman, and V. Rhue for collections assistance, M. Fox, J. Maisano, M. Colbert, and K. Smithson for assistance with CT scanning, J. Watanabe for discussions on skeletal ontogeny, and K. Super for the discovery and donation of FHSM VP-18702. We thank A. Maltese for photographs of recently discovered comparative material. We thank as well R. Olive for his Mesozoic avialan illustrations. We are indebted to K. van Grouw for assistance with the three-dimensional skeletal reconstruction of *Ichthyornis* and for her stunning skeletal illustration.

## SUPPLEMENTAL INFORMATION

Additional data about the studied specimens, their provenance and their identification as *Ichthyornis dispar*, as well as a list of comparative extant material is provided in the Supplemental Information. Detailed scanning parameters, phylogenetic methods, phylogenetic results and description of the characters of morphological phylogenetic matrices are provided in the Supplemental Information as well. Supplemental figures include detailed illustrative plates of the main specimens included in this study with all their preserved elements to scale, as well as 16 supplemental phylogenetic trees.

## DATA AVAILABILITY STATEMENT

The data generated in this this study is included directly in the article and Supplementary Material. The digital 3D models of the specimens are available on MorphoSource under the following link: DOI.ORG/XXXX/XXXX

