## Supplemental data for "40 new specimens of *Ichthyornis* provide unprecedented insight into the postcranial morphology of crownward stem group birds"

#### CONTENTS

I: Institutional abbreviations

II: Specimen content summary and provenance data

III: Scan parameters

IV: Comparative specimens

V: Phylogenetic analyses

VI: Synapomorphies diagnosing key clades

VII: Morphological character descriptions

VIII: Supplementary references

#### I: Institutional abbreviations

**ALMNH:** Vertebrate Paleontology Collection, Alabama Museum of Natural History, University of Alabama, Tuscaloosa, AL, USA

**BHI:** Black Hills Institute of Geological Research, Hill City, SD, USA

**FHSM:** Sternberg Museum of Natural History, Fort Hays State University, Hays, KS, USA

**KUVP:** Vertebrate Paleontology Division, University of Kansas Biodiversity Institute & Natural History Museum, Lawrence, KS, USA

**MSC & RMM**: McWane Science Center (formerly Red Mountain Museum), Birmingham, AL, USA

**NHMUK**: National History Museum, London, UK

**UMZC**: University of Cambridge Museum of Zoology, Cambridge, UK

**YPM:** Yale Peabody Museum, Yale University, New Haven, NY, USA

#### II: Specimen content summary and provenance data

Here we list all the new *Ichthyornis* specimens included in this study. A brief summary of the anatomical elements represented, their state of preservation, and additional relevant information is provided for each specimen. All available provenance, locality and stratigraphic data for each specimen are listed; locality data for specimens described in Field et al., (2018) were previously provided in that work.

The apomorphies that allow the referral of each specimen to *Ichthyornis* are listed, based on those suggested by Clarke (2004): (1) a single large pneumatic foramen situated on the anteromedial surface of the quadrate, (2) amphicoelous cervical vertebrae, (3) free caudal vertebrae exhibiting well-developed and elongated prezygapophyses, (4) scapula exhibiting an extremely diminutive acromion process, (5) pit-shaped fossa on the distal end of the bicipital crest of the humerus, (6) ulnar trochlear surface equal in length across its caudal and distal surfaces, (7) oval scar located on the caudoventral surface of the distal radius, (8) a large tubercle developed close to the articular surface of phalanx II:1 in the carpometacarpus and (9) presence of an internal index process in the distal end of manual phalanx II:1. Where none of these apomorphies are preserved, specimens were referred to *Ichthyornis* on the basis of morphological congruence with other diagnosed *Ichthyornis* specimens.

An abbreviated summary of this section is provided in Table 1 of the main text.

**ALMNH PV 1988.20.180:** The specimen comprises a single isolated left distal tarsometatarsus. The element is well preserved in three dimensions and undistorted. The shaft is broken close to the proximal portion of the element, and the trochlea of metatarsal I is missing.

The specimen does not preserve any previously identified apomorphies of *Ichthyornis* and is diagnosed on the basis of its morphological similarity to more complete specimens, particularly ALMNH PV 1993.2.133 and FHSM VP-18702.

The specimen derives from the Mooreville Chalk Formation (early Campanian, Late Cretaceous), from the Ada-3 site near "Robert Wilson property" in Dallas County, Alabama, US.

**ALMNH PV 1988.20.181:** The specimen comprises a single isolated left distal tarsometatarsus. The element is well preserved in three-dimensions, although part of the shaft is slightly flattened and missing some of the external bone surface. The shaft is broken close to its midpoint. All metatarsal trochleae are preserved, although slightly eroded on their distal surfaces.

The specimen does not preserve any previously identified apomorphies of *Ichthyornis* and is diagnosed on the basis of its morphological similarity to more complete specimens, particularly ALMNH PV 1993.2.133, FHSM VP-18702 and YPM 1739.

The specimen derives from the Mooreville Chalk Formation (early Campanian, Late Cretaceous), from the Ada-3 site near "Robert Wilson property" in Dallas County, Alabama, US.

**ALMNH PV 1988.20.183.1:** The specimen includes a single isolated left distal femur. The bone is preserved in three dimensions and is undistorted, but most of the bone surface is eroded and poorly preserved. The shaft is broken close to its distal end.

The specimen does not preserve any previously identified apomorphies of *Ichthyornis* and is diagnosed on the basis of its morphological similarity to more complete specimens, particularly ALMNH PV 1993.2.133 and YPM 1450.

The specimen derives from the the Mooreville Chalk Formation (early Campanian, Late Cretaceous), from the Ada-3b site near "David Wilson property" in Dallas County, Alabama, US.

**ALMNH PV 1988.20.186:** The specimen comprises a single isolated right proximal femur. The element is preserved in three dimensions and is undistorted, with only part of the femoral trochanter broken off. The shaft is broken very close to the proximal end of the femur.

The specimen does not preserve any previously identified apomorphies of *Ichthyornis* and is diagnosed on the basis of its morphological similarity to more complete specimens, particularly ALMNH PV 1993.2.133 and FHSM VP-18702.

The specimen derives from the Mooreville Chalk Formation (early Campanian, Late Cretaceous), from the Ada-3b site near "David Wilson property" in Dallas County, Alabama, US.

**ALMNH PV 1988.20.375.3:** The specimen includes a single isolated right proximal tarsometatarsus. The element is preserved in three dimensions and is undistorted, but most of the bone surface is eroded and poorly preserved. The shaft is preserved almost in its entirety and is broken very close to its distal end. Despite the poor condition of the bone, the hypotarsus is well-preserved.

The specimen does not preserve any previously identified apomorphies of *Ichthyornis* and is diagnosed on the basis of its morphological similarity to more complete specimens, particularly ALMNH PV 1993.2.133, FHSM VP-18702 and YPM 1739.

The specimen derives from the Mooreville Chalk Formation (early Campanian, Late Cretaceous), from the Ada-3 site near "David Wilson property" in Dallas County, Alabama, US.

**ALMNH PV 1988.20.427.1:** The specimen includes a single isolated left proximal humerus. The element is exceptionally well-preserved in three-dimensions and is undistorted. Most of the shaft is missing, being broken close to the proximal end of the bone, and only the base of the deltopectoral crest is preserved.

The specimen is diagnosed as *Ichthyornis* based on the presence of a pit-shaped fossa on the distal end of the bicipital crest (5).

The specimen derives from the Mooreville Chalk Formation (early Campanian, Late Cretaceous), from the Ada-3 site near "David Wilson property" in Dallas County, Alabama, US.

**ALMNH PV 1988.20.5:** The specimen includes two isolated elements, a portion of left proximal humerus and an unidentified bone fragment, which might correspond to a fragment of a vertebra. The humerus is preserved in three-dimensions, but it exhibits multiple breakages. Only part of the humeral head and the ventral tubercle are preserved, and the proximal surface of the ventral tubercle is eroded. The possible vertebral fragment is poorly preserved and lacks most recognizable features, but given its elongated shape it could potentially correspond to a portion of the pygostyle or synsacrum.

The specimen is diagnosed as *Ichthyornis* based on the presence of a pit-shaped fossa on the distal end of the bicipital crest (5).

The specimen derives from the Mooreville Chalk Formation (early Campanian, Late Cretaceous) from the Ada-3c site near Harrell Station in Dallas County, Alabama, US.

**ALMNH PV 1988.26.1:** The specimen comprises a single isolated element, probably a fragmentary pedal phalanx. The shaft is broken, but the proximal portion of the bone is well-preserved, including a ginglymoid articulation surface. An ossified tendon seems to be attached to the bone close to the proximal end of the shaft.

The specimen derives from the Mooreville Chalk Formation (early Campanian, Late Cretaceous) from Alabama, US.

**ALMNH PV 1993.2.191.1:** The specimen comprises a single isolated right distal tibiotarsus. The bone is in exceptional condition and is three-dimensionally preserved. Most of the shaft is missing, being broken close to the distal end of the tibiotarsus.

The specimen does not preserve any previously identified apomorphies of *Ichthyornis* and is diagnosed on the basis of its morphological similarity to more complete specimens, particularly ALMNH PV 1993.2.133, YPM 1450 and YPM 1732.

The specimen derives from the Mooreville Chalk Formation (early Campanian, Late Cretaceous) from the Ada-3UA-10 site near Harrell Station in Dallas County, Alabama, US.

**ALMNH PV 1993.2.133:** The specimen comprises a disarticulated partial skeleton (Sup. Fig. 1 and 2). The elements preserved are three cervical vertebrae (including the atlas), four thoracic vertebrae, a partial synsacrum divided into two portions, a single proximal portion of a thoracic rib, a right coracoid, the right omal end of the furcula, the distal portion of the left ulna, both ulnar carpals, a complete right carpometacarpus divided into two fragments, right manual phalanges II-1 and II-2, a complete left femur, the proximal and distal ends of the left tibiotarsus, the proximal and distal ends of the left tarsometatarsus and a single pedal phalanx. Most of the elements are preserved in exceptional condition in three dimensions and do not exhibit a substantial level of deformation; however, most of the long bones are broken or missing portions of their shafts.

The specimen is diagnosed as *Ichthyornis* based on the presence of an ulnar trochlear surface equal in length across its caudal and distal surfaces (6), a large tubercle developed close to the articular surface of the II:1 phalanx in the carpometacarpus (8) and the presence of an internal index process on the distal end of manual phalanx II:1.

The specimen derives from the Mooreville Chalk Formation (early Campanian, Late Cretaceous) from the Harrell Station Site, ~1.5 miles south-southeast of Harrell, SW ¼, Section 29, T. 17 N., R. 9 E, Marion Junction Quadrangle, Dallas County, Alabama, US. Based on co-occurring nanoflora, the site is confidently assigned to Nanofossil Zone CC-18a, the lower half of the *Aspidolithus parcus* Zone. This corresponds to an early (though not earliest) Campanian age (Field et al., 2018).

**BHI 6420:** The specimen comprises a partial disarticulated skeleton preserving the thoracic region. The elements preserved are both coracoids (the right one divided in two fragments), the left omal end of the furcula, both scapulae, both humeri (both broken in two pieces along their shafts), the proximal end of the right ulna, the complete right radius (divided in two fragments), the right radial carpal, the complete right carpometacarpus, right and left manual phalanges II-1 and right manual phalanges II-2 and III-1. Most of the elements are flattened and show multiple signs of breakage, although most long elements are preserved in their entirety despite being broken into several fragments.

The specimen is diagnosed as *Ichthyornis* based on the presence of a scapula exhibiting an extremely diminutive acromion process (4), a pit-shaped fossa on the distal end of the bicipital crest of the humerus (5), a large tubercle developed close to the articular surface of phalanx II:1 on the carpometacarpus (8) and the presence of an internal index process on the distal end of manual phalanx II:1 (9). Despite being extremely diminutive, the shape of the acromion process of the scapula is globose and differs from that of other *Ichthyornis* specimens. The distal portion of the ulna is too distorted to evaluate its proportions and the oval scar on the caudoventral surface of the distal radius is either eroded or absent.

The specimen derives from the smoky Hill Member of the Niobrara Formation from Gove County, Kansas (US).

**BHI 6421:** The specimen includes multiple isolated cranial and postcranial elements. Cranial and mandibular elements from this specimen were previously described by Field et al. (2018). The postcranial elements preserved comprise three partial thoracic vertebrae, most of the shaft and the distal portion of the right humerus, the distal end of the right ulna, a complete right manual phalanx II-1 and the shaft and distal portion of the right tibiotarsus. The quality of preservation of the different postcranial elements is variable, with a flattened and fragmentary humerus and ulna, but well-preserved three-dimensional vertebrae, manual phalanx and tibiotarsus.

The specimen is diagnosed as *Ichthyornis* based on the presence of a single large pneumatic foramen situated on the anteromedial surface of the quadrate (1), an ulnar trochlear surface equal in length across its caudal and distal surfaces (6) and an internal index process on the distal end of manual phalanx II:1 (9).

Specimen collected by Dave Tanking in Gove County, Kansas (US), about 2 miles east of Monument Rocks. Specimen was from high in section, above the *Inoceramus grandis* zone and within the *I. platinus* zone, placing the site within the upper Smoky Hill Member of the Niobrara Formation.

**FHSM VP 18792:** The specimen comprises various associated matrix blocks with encased fossil material, as well as extracted isolated elements, preserving a fairly complete partial skeleton (Sup. Fig. 3, 4 and 5). The cranial and mandibular elements from this specimen were described by Field et al. (2018). The main matrix block preserves most cranial elements, two fragmentary cervical vertebrae, the synsacrum, multiple complete and fragmentary ribs, including at least one thoracic rib, the complete right coracoid and the omal portion of the left coracoid, the proximal portion of the right humerus and the distal portion of the left humerus, the complete left radius and the distal portion of the right radius, the left ulna, the right radial carpal, the proximal portion of the left carpometacarpus and the complete right femur. An associated but separate block preserves a complete left scapula, the distal portion of the left tibiotarsus and the complete right tarsometatarsus. A third block preserves a complete and three-dimensional sternum, the distal portion of the left carpometacarpus, a single pedal phalanx and several fragmentary ribs. The elements extracted from the matrix are the left radial carpal, the right ulnar carpal, the left manual phalanx II-1, and three pedal phalanges. The preservation of the different elements is variable; all axial elements are poorly preserved and most long bones are complete but heavily flattened and distorted, including the radius, ulna and femur. In contrast, pectoral and manual elements are exceptionally well-preserved in three dimensions.

The specimen is diagnosed as *Ichthyornis* based on the presence of a single large pneumatic foramen situated on the anteromedial surface of the quadrate (1), amphicoelous cervical vertebrae (2), an extremely diminutive acromion process of the scapula (4), a pit-shaped fossa on the distal end of the bicipital crest of the humerus (5), an oval scar located on the caudoventral surface of the distal radius (7), a large tubercle developed close to the articular surface of phalanx II:1 of the carpometacarpus (8), and an internal index process on the distal end of manual phalanx II:1 (9). Despite being extremely diminutive, the shape of the acromion process of the scapula is hooked and slightly elongated, differing from that of other known *Ichthyornis* specimens. The distal portion of the ulna is too distorted to evaluate its proportions. Despite most of the internal index process being broken off, the base of the process is clearly preserved.

The specimen derives from the base of MU 10 in the Smoky Hill Member of the Niobrara Formation (middle Santonian stage, Late Cretaceous), near Castle Rock in southeast Gove Country, Kansas, US (Coordinates 38.852575 -100.165942).

**KUVP 2281:** The specimen includes two isolated elements, a left coracoid and a left scapula (Sup. Fig. 6). Both elements are in an exceptional state of preservation, exhibiting three-dimensional morphology and minimal breakage, with only the caudalmost end of the scapula missing.

The specimen is diagnosed as *Ichthyornis* based on the presence of an extremely diminutive acromion process of the scapula (4).

The specimen derives from the Smoky Hill Member of the Niobrara Formation (middle Santonian stage, Late Cretaceous) in Kansas, US.

**KUVP 2300:** The specimen comprises a single isolated complete left humerus (Sup. Fig. 6). The element is in good condition and is three-dimensionally preserved, although it exhibits substantial breakage along the shaft and in the deltopectoral crest.

The specimen is diagnosed as *Ichthyornis* on the basis of the presence of a pit-shaped fossa on the distal end of the bicipital crest of the humerus (5).

The specimen derives from the Smoky Hill Member of the Niobrara Formation (middle Santonian stage, Late Cretaceous), near Hill Creek River, Kansas, US.

**KUVP 5969:** The specimen includes several forelimb elements (Sup. Fig. 6). The right humerus, radius and ulna are mounted in anatomical connection and embedded in plaster. The isolated elements include the right coracoid (broken into two pieces), a fragmentary left ulnar carpal and the complete right manual phalanx II-1. The preservation of all the elements is relatively poor, with all long bones exhibiting substantial flattening, and many relevant regions of the coracoid and humerus eroded or broken.

The specimen does not preserve any previously identified apomorphies of *Ichthyornis* and is diagnosed on the basis of its morphological similarity to more complete specimens, particularly with FHSM VP 18702 and YPM 1450. The distal portion of the ulna is too distorted to evaluate its proportions. Notably, manual phalanx II:1 lacks an internal index process (9) despite not showing any breakage; together with several other aspects of its morphology, this may indicate that this specimen represents a juvenile individual (see discussion in main text).

The specimen derives from the Smoky Hill Member of the Niobrara Formation (middle Santonian stage, Late Cretaceous), near Smoky Hill River, Logan County, Kansas, US.

**KUVP 25469:** The specimen preserves several isolated elements (Sup. Fig. 7), including a complete synsacrum, the right omal end of the furcula, the distal portion of the right humerus, most of the shaft and the distal portion of the right radius, a complete right carpometacarpus (divided into two portions) and a complete right manual phalanx II-2. The preservation of most elements is exceptional and three-dimensional, although the synsacrum and humerus exhibit substantial flattening and distortion.

The specimen is diagnosed as *Ichthyornis* on the basis of the presence of an oval scar located on the caudoventral surface of the distal radius (7).

The specimen derives from the Smoky Hill Member of the Niobrara Formation (middle Santonian stage, Late Cretaceous), from the locality Cove-21, Kansas, US.

**KUVP 25471:** The specimen includes a single extremely fragmentary thoracic vertebra and a complete left humerus (Sup. Fig. 7). The vertebra preserves only part of its centrum and the caudal articular surface. The humerus is flattened and lacks most of the deltopectoral crest.

The specimen is diagnosed as *Ichthyornis* based on the presence of a pit-shaped fossa on the distal end of the bicipital crest of the humerus (5).

The specimen derives from the Smoky Hill Member of the Niobrara Formation (middle Santonian stage, Late Cretaceous) in Logan County, Kansas, US.

**KUVP 25472:** The specimen comprises a partially complete axial series, a complete left radius and most of the shaft and proximal portion of the right ulna (Sup. Fig. 7). The axial series includes five cervical vertebrae, nine thoracic vertebrae, including two preserved in articulation, and three caudal vertebrae, two of them preserved in articulation. The radius and both the thoracic and caudal vertebrae are preserved in three dimensions and in exceptional condition. The ulna and most of the cervical vertebrae exhibit varying degrees of flattening, distortion and breakage.

The specimen is diagnosed as *Ichthyornis* on the basis of the presence of amphicoelous cervical vertebrae (2) and free caudal vertebrae exhibiting well-developed and elongated prezygapophyses (3). It is difficult to verify whether the radius exhibits an oval scar on its distal caudoventral surface, but this region might be slightly eroded or undeveloped.

The specimen was collected by M.C. Bonner and derives from the Smoky Hill Member of the Niobrara Formation (middle Santonian stage, Late Cretaceous), from the locality of KU-LOG-39 in Logan County, Kansas, US.

**KUVP 119673:** The specimen preserves a fairly complete partial skeleton and is comprised of a main matrix block with encased fossil elements and several extracted isolated elements (Sup. Fig. 8, 9 and 10). The cranial and mandibular elements from this specimen were described by Field et al. (2018). The main block preserves a partially complete axial series, of which most of the vertebrae are articulated or in close association; the vertebrae preserved are eight cervical vertebrae (at least four of which are smashed together and difficult to discern), two anterior thoracic vertebrae, a complete synsacrum and three isolated caudal vertebrae. The main block preserves a complete sternum as well, multiple complete and fragmentary ribs, including at least one thoracic rib, the complete left humerus, both right and left pelves (disarticulated from the synsacrum), a partial right femur preserving most of the shaft and part of the distal end, a distal fragmentary left tibiotarsus and the left fibula. The isolated elements extracted from the block are a single thoracic vertebra, the complete right coracoid, the right scapula, the complete right humerus, the complete right and left radii (the right element broken into two fragments) and the complete right ulna and most of the shaft and distal portion of the left ulna. The quality of preservation is variable. Many of the elements still encased within the main block are largely flattened (sternum, pelves, right humerus) and fragmentary (femur, tibiotarsus). Most of the elements extracted from the block are preserved in three dimensions and in exceptional condition (coracoid, radii, right humerus). The differential preservation of the right and left humeri is remarkable, allowing a detailed exploration of the effects of taphonomic distortion on humeral morphology in *Ichthyornis* (see main text). Many of the elements of this specimen preserve metallic radiopaque inclusions that complicated the scanning process. These were especially prevalent in the sternum, the coracoid and the right humerus.

The specimen is diagnosed as *Ichthyornis* based on the presence of a single large pneumatic foramen situated on the anteromedial surface of the quadrate (1), amphicoelous cervical vertebrae (2), free caudal vertebrae exhibiting well-developed and elongated prezygapophyses (3), an extremely diminutive acromion process in the scapula (4), a pit-shaped fossa on the distal end of the bicipital crest of the humerus (5), and an ulnar trochlear surface equal in length across its caudal and distal surfaces (6). It is difficult to verify whether the radius exhibits the presence of an oval scar on its caudoventral distal surface, but this region might be slightly eroded or undeveloped.

The specimen was collected by J.D. Stewar in 1992 from the Smoky Hill Member of the Niobrara Formation (Santonian Stage, Late Cretaceous), W1/2 Sec. II, T16S, R28W, near the center of the boundary between N.W. and S.W. quarters, Cathouse Ranch, Lane County, Kansas, US (Field et al., 2018).

**KUVP 123459:** The specimen includes a left partial carpometacarpus broken into proximal and distal fragments and missing part of the shaft of metacarpal II, and the distal end of the right ulna. Both elements are preserved in three-dimensions and in exceptional condition despite breakage.

The specimen is diagnosed as *Ichthyornis* based on the presence of a large tubercle developed close to the articular surface of phalanx II:1 on the carpometacarpus (8).

The specimen derives from the Smoky Hill Member of the Niobrara Formation (middle Santonian stage, Late Cretaceous), from the locality KU-LOG-D52 near Fox Canyon in Logan County, Kansas, US.

**MSC 13214:** The specimen comprises a single isolated right tarsometatarsus, broken into proximal and distal fragments and missing a portion of the shaft. The preservation quality is exceptional, exhibiting minimal flattening and distortion, preserving the hypotarsus and all three metatarsal trochleae.

The specimen does not preserve any previously identified apomorphies of *Ichthyornis* and is diagnosed on the basis of its morphological similarity to more complete specimens, particularly ALMNH PV 1993.2.133, FHSM VP-18702 and YPM 1739.

The specimen derives from the Mooreville Chalk Formation (early Campanian, Late Cretaceous) from the Agr-7 site in Greene County, Alabama, US.

**MSC 13868:** The specimen includes a single isolated bone fragment of uncertain identity, possibly a phalanx belonging to the first pedal digit. The element is preserved in three-dimensions and does not seem to exhibit any distortion, although it is broken at its (probable) proximal end.

The specimen derives from the Mooreville Chalk Formation (early Campanian, Late Cretaceous) from the Agr-11 site in Greene County, Alabama, US.

**MSC 34426:** The specimen includes a single isolated partial left carpometacarpus, preserving the distal end of the bone and most of the metacarpal II shaft. The specimen is in exceptional condition, is vpreserved in three-dimensions, and does not exhibit any substantial distortion.

The specimen is diagnosed as *Ichthyornis* based on the presence of a large tubercle developed close to the articular surface of phalanx II:1 in the carpometacarpus (8).

The specimen derives from the deposits of the Mooreville Chalk Formation (early Campanian, Late Cretaceous) from Alabama, US.

**MSC 34427:** The specimen is comprised of several forelimb elements. The elements preserved are the left coracoid (broken into three fragments, preserving the entire omal portion of the bone but only parts of the shaft and the sternal end), the distal end of the left ulna, and the proximal and distal ends of the left carpometacarpus. Additionally, three bone fragments may correspond to portions of the shaft of the carpometacarpus. All elements are preserved in three-dimensions but in poor condition, with multiple breakages and mostly eroded bone surfaces.

The specimen is diagnosed as *Ichthyornis* based on the presence of a large tubercle developed close to the articular surface of phalanx II:1 on the carpometacarpus (8) and an ulnar trochlear surface equal in length across its caudal and distal surfaces (6).

The specimen derives from the Mooreville Chalk Formation (early Campanian, Late Cretaceous) from Alabama, US.

**NHMUK A 905:** The specimen comprises three associated blocks of matrix with encased fossil material, as well as extracted isolated elements, preserving most of the pectoral region and several forelimb elements. The specimen preserves a partially complete sternum, the medial portion of the furcula, the complete right coracoid and most of the left coracoid, both complete scapulae, the complete right humerus and the proximal half of the left humerus, and the proximal portion of the right radius. Despite generally good preservation, all elements except the furcula and the radius are flattened and severely distorted.

The specimen is diagnosed as *Ichthyornis* based on the presence of an extremely diminutive acromion process on the scapula (4) and a pit-shaped fossa on the distal end of the bicipital crest of the humerus (5). Despite being extremely diminutive, the acromion processes of the two scapulae are slightly variable and differ from each other and from other *Ichthyornis* specimens, although these variations may be the result of taphonomic distortion.

The specimen derives from the Niobrara Formation (middle Santonian stage, Late Cretaceous), Kansas, US.

**RMM 2841:** The specimen includes a single isolated left distal carpometacarpus, preserving most of the distal end of the element and part of the metacarpal II shaft, which is broken approximately at its midpoint. The distal part of the element is preserved in three dimensions, but most of the shaft is flattened and broken.

The specimen is diagnosed as *Ichthyornis* based on the presence of a large tubercle developed close to the articular surface of phalanx II:1 on the carpometacarpus (8).

The specimen derives from the Mooreville Chalk Formation (early Campanian, Late Cretaceous) from the Agr-7 site in Greene County, Alabama, US.

**RMM 3394:** The specimen includes a single isolated left distal carpometacarpus, broken close to the midpoint of the metacarpal II shaft. The element is preserved in exceptional condition and in three dimensions.

The specimen is diagnosed as *Ichthyornis* based on the presence of a large tubercle developed close to the articular surface of phalanx II:1 on the carpometacarpus (8).

The specimen derives from the deposits of the Lower Mooreville Chalk Formation (early Campanian, Late Cretaceous) from the Apn-1 site in Pickens County, Alabama, US.

**RMM 5794:** The specimen comprises a single fragmentary proximal right humerus. The element preserves only a portion of the humeral head and the humeral shaft, but it is missing most recognizable features. The element is preserved in three dimensions but exhibits substantial breakage.

The specimen does not preserve any previously identified apomorphies of *Ichthyornis* and is diagnosed on the basis of its morphological similarity to more complete specimens, particularly KUVP 2300 and 119673 and YPM 1450.

The specimen derives from the deposits of the Mooreville Chalk Formation (early Campanian, Late Cretaceous) from the Agr-17 site in Greene County, Alabama, US.

**RMM 5895:** The specimen comprises several fragmentary forelimb elements. The elements preserved are the distal portion of the left humerus, a partial left radius, preserving the proximal region and most of the shaft, and left manual phalanx II:1. The humerus is poor condition and flattened, but both the radius and the manual phalanx are preserved in three dimensions and in good condition, despite minor breakage.

The specimen is diagnosed as *Ichthyornis* based on the presence of an internal index process in the distal end of manual phalanx II:1 (9).

The specimen derives from the deposits of the Mooreville Chalk Formation (early Campanian, Late Cretaceous) from the Agr-4 site in Greene County, Alabama, US.

**RMM 5916:** The specimen comprises a single isolated proximal left ulna. Most of the shaft of the ulna is missing, but the proximal end of the element is preserved in three dimensions and in exceptional condition, not exhibiting any substantial deformation or breakage.

The specimen does not preserve any previously identified apomorphies of *Ichthyornis* and is diagnosed on the basis of its morphological similarity to more complete specimens, particularly KUVP 119673 and YPM 1450 and 1740.

The specimen derives from the Mooreville Chalk Formation (early Campanian, Late Cretaceous) from the Agr-17 site in Greene County, Alabama, US.

**RMM 5937:** The specimen includes a single isolated right coracoid broken into two fragments. The omal end and most of the sternal end of the element are well preserved in three dimensions, but part of the shaft is missing.

The specimen does not preserve any previously identified apomorphies of *Ichthyornis* and is diagnosed on the basis of its morphological similarity to more complete specimens, particularly KUVP 2281, 119673 and YPM 1733.

The specimen derives from the Mooreville Chalk Formation (early, but not earliest, Campanian, Late Cretaceous) from the Ada-3 site near Harrell Station, in Dallas County, Alabama, US.

**RMM 6200:** The specimen comprises a fragmentary left radius and ulna. The radius is broken into two fragments comprising the proximal portion and part of the shaft, and the ulna preserves only the distal portion of the element. All elements are well preserved and in three dimensions, although the bone surface of the distal ulna is slightly eroded.

The specimen is diagnosed as *Ichthyornis* based on the presence of an ulnar trochlear surface equal in length across its caudal and distal surfaces (6).

The specimen derives from the deposits of the Mooreville Chalk Formation (early Campanian, Late Cretaceous) from the Agr-17 site in Greene County, Alabama, US.

**RMM 6201:** The specimen is comprised of a single isolated right manual phalanx II:1. The element is mostly complete, but the distal portion of the bone is broken, and the bone surface is slightly eroded.

The specimen is diagnosed as *Ichthyornis* based on the presence of an internal index process on the distal end of manual phalanx II:1 (9). Despite most of the internal index process being broken off, the base of the process is clearly preserved.

The specimen derives from the deposits of the Mooreville Chalk Formation (early Campanian, Late Cretaceous) from the Agr-17 site in Greene County, Alabama, US.

**RMM 6202:** The specimen includes a single almost complete right carpometacarpus. The element is broken into three major fragments, the proximal portion, the shaft of metacarpal II and the distal portion of the element. Additional fragments probably correspond to portions of metacarpal III. Despite the breakage, the element is preserved in good condition and in three dimensions, not exhibiting any major flattening or distortion.

The specimen is diagnosed as *Ichthyornis* based on the presence of a large tubercle developed close to the articular surface of phalanx II:1 in the carpometacarpus (8).

The specimen derives from the Mooreville Chalk Formation (early Campanian, Late Cretaceous) from the Agr-17 site in Greene County, Alabama, US.

**RMM 7841:** The specimen comprises several isolated pectoral and forelimb fragments. The elements preserved are the omal portion of the right coracoid, the complete right humerus and a portion of the shaft and the distal end of the left humerus, and the distal portion of the left ulna. Both the coracoid and ulna are well preserved in three dimensions, but both humeri are flattened and exhibit major distortion and breakage.

The specimen is diagnosed as *Ichthyornis* based on the presence of a pit-shaped fossa on the distal end of the bicipital crest of the humerus (5) and an ulnar trochlear surface equal in length across its caudal and distal surfaces (6).

The specimen derives from the Mooreville Chalk Formation (early Campanian, Late Cretaceous) from the Agr-4 site in Greene County, Alabama, US.

**RMM 7842:** The specimen includes a single fragmentary right humerus. The element is broken into three fragments: a portion of the humeral head, the proximal half of the shaft (broken at its midpoint and including the base of the deltopectoral crest), and the distal half of the shaft including the distal end of the element. The bone is three-dimensionally preserved and in good condition despite the breakage, but part of the bone surface is slightly eroded.

The specimen does not preserve any apomorphic features and is diagnosed as *Ichthyornis* on the basis of morphological similarity, particularly with KUVP 2300 and 119673 and YPM 1450.

The specimen derives from the deposits of the Mooreville Chalk Formation (early Campanian, Late Cretaceous) from Greene County, Alabama, US.

**RMM 7844:** The specimen includes two badly preserved bone fragments. These probably correspond to the distal portion of the right humerus and part of the humerus shaft. The elements are flattened and exhibit major distortion, and most of the bone surface is eroded.

The specimen does not preserve any previously identified apomorphies of *Ichthyornis* and is diagnosed on the basis of its morphological similarity to more complete specimens, particularly KUVP 2300 and 119673, and YPM 1450.

The specimen derives from the deposits of the Mooreville Chalk Formation (early Campanian, Late Cretaceous) from the Agr-4 site in Greene County, Alabama, US.

**Previously described specimens**

Here we list several previously described *Ichthyornis* specimens, discussed in the main text for comparative purposes. The elements of each specimen included in this study are described, but more detailed descriptions of these specimens were provided by Marsh (1880) and Clarke (2004). Provenance data for all YPM material discussed has been previously published (e.g., Clarke 2004).

**YPM 1450:** This is the holotype of *Ichthyornis dispar* (Marsh, 1880; Clarke, 2004). The specimen comprises a partially complete skeleton, including a partial skull, mandibles, cervical, thoracic and sacral vertebrae, a partial sternum, ribs, coracoid, humerus, ulna, radius, carpometacarpus, femur, tibiotarsus and several additional fragments. In this study we incorporated the partial sternum and the right ulna. The sternum preserves only the cranial end of the element and part of the dorsal surface, and it is crushed, distorted, and poorly preserved. In contrast, the right ulna is complete and undistorted and exhibits only minor breakage.

As the holotype, YPM 1450 is the specifier for the name *Ichthyornis*, and exhibits amphicoelous cervical vertebrae (2), the presence of a pit-shaped fossa on the distal end of the bicipital crest of the humerus (5), an ulnar trochlear surface equal in length across its caudal and distal surfaces (6), an oval scar located on the caudoventral surface of the distal radius (7) and a large tubercle close to the articular surface of manual phalanx II:1 in the carpometacarpus (8).

**YPM 1724:** This specimen was previously identified as *Ichthyornis victor* (Marsh, 1880; Clarke, 2004), and comprises a single right carpometacarpus. The element is exceptionally well preserved in three dimensions and does not exhibit any distortion or substantial breakage.

The specimen is diagnosed as *Ichthyornis* on the basis of the presence of a large tubercle close to the articular surface of manual phalanx II:1 on the carpometacarpus (8).

**YPM 1733:** This specimen was previously identified as *Ichthyornis victor* (Marsh 1880; Clarke, 2004), and comprises a partial skeleton including cervical vertebrae, coracoid, scapula, humerus, radius, synsacrum and part of the ilium. This study incorporates the right coracoid, which is three-dimensionally preserved, but exhibits minor breakage and several missing portions on its omal and sternal ends.

The specimen is diagnosed as *Ichthyornis* on the basis of the presence of amphicoelous cervical vertebrae (2) and a scapula exhibiting an extremely diminutive acromion process (4).

**YPM 1740:** This specimen is the holotype of the now invalid *Ichthyornis validus* (Marsh, 1880; Clarke, 2004). It includes a single isolated right ulna. The element is complete and does not exhibit any breakage or distortion. The proximal portion of the element appears to be slightly eroded or undeveloped, as described by Clarke (2004).

The specimen is diagnosed as *Ichthyornis* on the basis of the presence of an ulnar trochlear surface equal in length across its caudal and distal surfaces (6).

**YPM 1741:** This specimen was previously identified as *Ichthyornis victor* (Marsh, 1880; Clarke, 2004). The specimen preserves several pectoral and forelimb elements, including the coracoid, scapula, humerus and radius. In this study we incorporated the right radius, which is three-dimensionally preserved and in exceptional condition and does not show any substantial breakage or distortion.

The specimen is diagnosed as *Ichthyornis* on the basis of the presence of an oval scar located on the caudoventral surface of the distal radius (7).

**II: Scan Parameters**

All ALMNH, BHI, FHSM, KUVP, MSC, and RMM specimens were scanned at the University of Texas High-Resolution X-ray CT Facility (UTCT). NHMUK A 905 and all comparative extant taxa were scanned at the Cambridge Biotomography Centre (CBC). All YPM specimens were scanned at the Center for Nanoscale Systems at Harvard (CNS). Scanning parameters were as follows. All scanned material was digitally segmented and rendered using VGStudio Max 3.3.0 and 3.4.5.

**ALMNH PV 1988.20.180, 1988.20.181, 1988.20.183.1, 1988.20.186, 1988.20.375.3 and 1988.20.427.1** (scanned together): NSI scanner at UTCT. Feinfocus High Power source, 170 kV, 0.15 mA, aluminum filter, Perkin Elmer detector, 0.25 pF gain, 1 fps, 1x1 binning, no flip, source to object 172.0 mm, source to detector 1316.735 mm, continuous CT scan, no frames averaged, 0 skip frames, 3000 projections, 6 gain calibrations, 5 mm calibration phantom, data range [-2.5, 45.0] (grayscales adjusted from NSI defaults), beam-hardening correction = 0.1. Voxel size = 0.0181 mm. Total slices = 1236.

**ALMNH PV 1988.20.5, 1988.26.1 and 1993.2.191.1** (scanned together): NSI scanner at UTCT. Feinfocus High Power source, 170 kV, 0.15 mA, aluminum filter, Perkin Elmer detector, 0.25 pF gain, 1 fps, 1x1 binning, no flip, source to object 182.0 mm, source to detector 1316.735 mm, continuous CT scan, no frames averaged, 0 skip frames, 3000 projections, 6 gain calibrations, 5 mm calibration phantom, data range [-3.0, 40.0] (grayscales adjusted from NSI default values), beam-hardening correction = 0.1. Voxel size = 0.0204 mm. Total slices = 1620.

**ALMNH PV 1993.2.133 (hindlimb elements):** NSI scanner at UTCT. Feinfocus source, high power, 170 kV, 0.16 mA, 1 aluminum filter, Perkin Elmer detector, 0.25 pF gain, 1 fps (999.911 ms integration time), no binning, no flip, source to object 175 mm, source to detector 1316.685 mm, continuous CT scan, no frames averaged, 0 skip frames, 3600 projections, 6 gain calibrations, 5 mm calibration phantom, data range [-2.5, 53] (grayscale range adjusted from NSI defaults), beam-hardening correction = 0.05. Voxel size = 0.0203 mm. Total slices = 1879.

**ALMNH PV 1993.2.133 (forelimb and torso elements):** NSI scanner at UTCT. Feinfocus source, high power, 170 kV, 0.16 mA, 1 aluminum filter, Perkin Elmer detector, 0.25 pF gain, 1 fps (999.911 ms integration time), no binning, no flip, source to object 175 mm, source to detector 1316.685 mm, continuous CT scan, no frames averaged, 0 skip frames, 3600 projections, 6 gain calibrations, 5 mm calibration phantom, data range [-2.5, 53] (grayscale range adjusted from NSI defaults), beam-hardening correction = 0.1. Voxel size = 0.0203 mm. Total slices = 1563.

**BHI 6420 (coracoid, humeri, carpometacarpus and manual elements):** NSI scanner at UTCT. Feinfocus source, high power, 190 kV, 0.15 mA, aluminum filter, Perkin Elmer detector, 0.25 pF gain, 1 fps (999.911 ms integration time), no binning, no flip, source to object 222 mm, source to detector 1316.544 mm, continuous CT scan, no frames averaged, 0 skip frames, 2873 projections, 6 gain calibrations, 5 mm calibration phantom, data range [-1, 15] (grayscale range adjusted from NSI defaults), beam-hardening correction = 0.1. Voxel size = 0.031 mm. Total slices = 1813.

**BHI 6420 (coracoid, scapulae, ulna, radii, carpal and manual elements, fibula):** NSI scanner at UTCT. Feinfocus source, high power, 190 kV, 0.15 mA, aluminum filter, Perkin Elmer detector, 0.25 pF gain, 1 fps (999.911 ms integration time), no binning, no flip, source to object 230 mm, source to detector 1316.544 mm, continuous CT scan, no frames averaged, 0 skip frames, 2873 projections, 6 gain calibrations, 5 mm calibration phantom, data range [-1, 15] (grayscale range adjusted from NSI defaults), beam-hardening correction = 0.1. Voxel size = 0.0328 mm. Total slices = 1842.

**BHI 6421:** NSI scanner at UTCT. Feinfocus source, high power, 190 kV, 0.15 mA, aluminum filter, Perkin Elmer detector, 0.25 pF gain, 1 fps (999.911 ms integration time), no binning, no flip, source to object 230 mm, source to detector 1316.544 mm, continuous CT scan, no frames averaged, 0 skip frames, 2873 projections, 6 gain calibrations, 5 mm calibration phantom, data range [-1, 15] (grayscale range adjusted from NSI defaults), beam-hardening correction = 0.1. Voxel size = 0.0328 mm. Total slices = 1842.

**FHSM VP 18702 (main block):** NSI scanner at UTCT. Feinfocus source, high power, 170 kV, 0.225 mA, 1 aluminum filter, Perkin Elmer detector, 0.25 pF gain, 1 fps (999.911 ms integration time), no binning, no flip, source to object 597.681 mm, source to detector 1316.685 mm, continuous CT scan, 2 frames averaged, 0 skip frames, 2550 projections, 6 gain calibrations, 15 mm calibration phantom, data range [-5, 50] (grayscale range adjusted from NSI defaults), beam-hardening correction = 0.4. Voxel size = 0.1163 mm. Total slices = 1811.

**FHSM VP 18702 (block containing scapula, tibiotarsus and tarsometatarsus):** NSI scanner at UTCT. Feinfocus source, high power, 170 kV, 0.180 mA, 1 aluminum filter, Perkin Elmer detector, 0.25 pF gain, 1 fps (999.911 ms integration time), no binning, no flip, source to object 282.463 mm, source to detector 1316.685 mm, continuous CT scan, 2 frames averaged, 0 skip frames, 2460 projections, 6 gain calibrations, 5 mm calibration phantom, data range [-0.3, 4.6] (grayscale range adjusted from NSI defaults), beam-hardening correction = 0.05. Voxel size = 0.0447 mm. Total slices = 1767.

**FHSM VP 18702 (block containing sternum, distal carpometacarpus and pedal phalanx):** NSI scanner at UTCT. Feinfocus source, high power, 170 kV, 0.225 mA, 1 aluminum filter, Perkin Elmer detector, 0.25 pF gain, 1 fps (999.911 ms integration time), no binning, no flip, source to object 341.114 mm, source to detector 1316.685 mm, continuous CT scan, 2 frames averaged, 0 skip frames, 2550 projections, 6 gain calibrations, 15 mm calibration phantom, data range [-0.75, 9] (grayscale range adjusted from NSI defaults), beam-hardening correction = 0.15. Voxel size = 0.058 mm. Total slices = 1727.

**FHSM VP 18702 (carpal and manual elements, pedal phalanges):** NSI scanner at UTCT. Feinfocus source, high power, 170 kV, 0.15 mA, 1 aluminum filter, Perkin Elmer detector, 0.25 pF gain, 1 fps (999.911 ms integration time), no binning, no flip, source to object 142.92 mm, source to detector 1316.685 mm, continuous CT scan, no frames averaged, 0 skip frames, 3600 projections, 6 gain calibrations, 5 mm calibration phantom, data range [-2.8, 30] (grayscale range adjusted from NSI defaults), beam-hardening correction = 0.05. Voxel size = 0.013 mm. Total slices = 1820.

**KUVP 2281, 2300 and 119673 (scapula)** (scanned together): NSI scanner at UTCT. Feinfocus source, high power, 200 kV, 0.12 mA, no filter, Perkin Elmer detector, 0.5 pF gain, 2 fps (499.893 ms integration time), no binning, no flip, source to object 109.3 mm, source to detector 993 mm, continuous CT scan, no frames averaged, 0 skip frames, 3600 projections, 6 gain calibrations, 5 mm calibration phantom, data range [-7, 62] (rescaled from NSI default), beam-hardening correction = 0.25. Voxel size = 0.035 mm. Total slices = 1893.

**KUVP 5969 (block containing humerus, radius and ulna):** NSI scanner at UTCT. Feinfocus source, high power, 220 kV, 0.23 mA, no filter, Perkin Elmer detector, 0.5 pF gain, 2 fps (499.893 ms integration time), no binning, no flip, source to object 202 mm, source to detector 993 mm, continuous CT scan, no frames averaged, 0 skip frames, 3600 projections, 6 gain calibrations, 15 mm calibration phantom, data range [-4, 13] (rescaled from NSI default), beam-hardening correction = 0.125. Voxel size = 0.0643 mm. Total slices = 1619.

**KUVP 5969 (coracoid, carpal and manual elements):** NSI scanner at UTCT. Feinfocus source, high power, 220 kV, 0.23 mA, no filter, Perkin Elmer detector, 0.5 pF gain, 2 fps (499.893 ms integration time), no binning, no flip, source to object 202 mm, source to detector 993 mm, continuous CT scan, no frames averaged, 0 skip frames, 3600 projections, 6 gain calibrations, 15 mm calibration phantom, data range [-4, 13] (rescaled from NSI default), beam-hardening correction = 0.125. Voxel size = 0.0643 mm. Total slices = 1619.

**KUVP 25469:** NSI scanner at UTCT. Feinfocus source, high power, 200 kV, 0.120 mA, no filter, Perkin Elmer detector, 0.5 pF gain, 2 fps (499.893 ms integration time), no binning, no flip, source to object 75 mm, source to detector 993 mm, continuous CT scan, no frames averaged, 0 skip frames, 3600 projections, 6 gain calibrations, 5 mm calibration phantom, data range [-4, 98] (rescaled from NSI default), beam-hardening correction = 0.25. Voxel size = 0.0241 mm. Total slices = 1864.

**KUVP 25471 (vertebra) and 123459 (distal ulna)** (Scanned together): NSI scanner at UTCT. Feinfocus source, high power, 200 kV, 0.145 mA, no filter, Perkin Elmer detector, 0.5 pF gain, 2 fps (499.893 ms integration time), no binning, no flip, source to object 124.123 mm, source to detector 1090.684 mm, continuous CT scan, no frames averaged, 0 skip frames, 3600 projections, 6 gain calibrations, 5 mm calibration phantom, data range [-10, 126] (rescaled from NSI default), beam-hardening correction = 0.3. Voxel size = 0.0344 mm. Total slices = 1827.

**KUVP 25471 (humerus), 25472 (radius and ulna), 123459 (carpometacarpus)** (scanned together): NSI scanner at UTCT. Feinfocus source, high power, 200 kV, 0.145 mA, no filter, Perkin Elmer detector, 0.5 pF gain, 2 fps (499.893 ms integration time), no binning, no flip, source to object 124.123 mm, source to detector 1090.684 mm, continuous CT scan, no frames averaged, 0 skip frames, 3600 projections, 6 gain calibrations, 5 mm calibration phantom, data range [-10, 126] (rescaled from NSI default), beam-hardening correction = 0.3. Voxel size = 0.0344 mm. Total slices = 1827.

**KUVP 25472 (vertebrae):** NSI scanner at UTCT. Feinfocus source, high power, 200 kV, 0.145 mA, no filter, Perkin Elmer detector, 0.5 pF gain, 2 fps (499.893 ms integration time), no binning, no flip, source to object 71.071 mm, source to detector 1090.684 mm, continuous CT scan, no frames averaged, 0 skip frames, 3600 projections, 6 gain calibrations, 5 mm calibration phantom, data range [-17, 107] (rescaled from NSI default), beam-hardening correction = 0.225. Voxel size = 0.0198 mm. Total slices = 916.

**KUVP 25472 (vertebrae and fragments), 119673 (vertebra)** (scanned together): NSI scanner at UTCT. Feinfocus source, high power, 200 kV, 0.145 mA, no filter, Perkin Elmer detector, 0.5 pF gain, 2 fps (499.893 ms integration time), no binning, no flip, source to object 71.071 mm, source to detector 1090.684 mm, continuous CT scan, no frames averaged, 0 skip frames, 3600 projections, 6 gain calibrations, 5 mm calibration phantom, data range [-6, 80] and [-20, 540] (rescaled from NSI default), beam-hardening correction = 0.15. Voxel size = 0.0198 mm. Total slices = 844.

**KUVP 119673 (main block):** NSI scanner at UTCT. Feinfocus source, high power, 200 kV, 0.14 mA, 1 aluminum filter, Perkin Elmer detector, 0.5 pF gain, 2 fps (499.893 ms integration time), no binning, no flip, source to object 202 mm, source to detector 993 mm, continuous CT scan, no frames averaged, 0 skip frames, 7400 projections, 6 gain calibrations, 15 mm calibration phantom, data range [-10, 446] or [-10, 969] (rescaled from NSI default), beam-hardening correction = 0.35. Voxel size = 0.0643 mm. Total slices = 1758.

**KUVP 119673 (coracoid):** NSI scanner at UTCT. Feinfocus source, high power, 200 kV, 0.12 mA, no filter, Perkin Elmer detector, 0.5 pF gain, 2 fps (499.893 ms integration time), no binning, no flip, source to object 59.65 mm, source to detector 993 mm, continuous CT scan, no frames averaged, 0 skip frames, 3600 projections, 6 gain calibrations, 5 mm calibration phantom, data range [-40, 1500] (rescaled from NSI default), beam-hardening correction = 0.35. Voxel size = 0.0192 mm. Total slices = 1821.

**KUVP 119673 (humerus, ulna and radii):** NSI scanner at UTCT. Feinfocus source, high power, 200 kV, 0.145 mA, no filter, Perkin Elmer detector, 0.5 pF gain, 2 fps (499.893 ms integration time), no binning, no flip, source to object 52.804 mm, source to detector 1090.684 mm, continuous CT scan, no frames averaged, 0 skip frames, 3600 projections, 6 gain calibrations, 5 mm calibration phantom, data range [-10, 246] and [-107.943871, 1487.616821] (rescaled from NSI default), beam-hardening correction = 0.25. Voxel size = 0.0148 mm. Total slices = 1323.

**KUVP 119673 (quadrate, vertebra and fragments):** NSI scanner at UTCT. Feinfocus source, high power, 200 kV, 0.145 mA, no filter, Perkin Elmer detector, 0.5 pF gain, 2 fps (499.893 ms integration time), no binning, no flip, source to object 52.804 mm, source to detector 1090.684 mm, continuous CT scan, no frames averaged, 0 skip frames, 3600 projections, 6 gain calibrations, 5 mm calibration phantom, data range [-10, 246] and [-107.943871, 1487.616821] (rescaled from NSI default), beam-hardening correction = 0.25. Voxel size = 0.0148 mm. Total slices = 1323.

**MSC 34427:** NSI scanner at UTCT. Feinfocus source, high power, 200 kV, 0.13 mA, no filter, Perkin Elmer detector, 0.5 pF gain, 2 fps (499.893 ms integration time), no binning, no flip, source to object 133.681 mm, source to detector 1091.513 mm, continuous CT scan, no frames averaged, 0 skip frames, 3600 projections, 6 gain calibrations, 5 mm calibration phantom, data range [-10, 300] (grayscale range adjusted from NSI defaults), beam-hardening correction = 0.4. Voxel size = 0.0147 mm. Total slices = 1095.

**NHMUK A 905 (block containing sternum):** Nikon 49 Metrology XT H 225 ST High Resolution CT Scanner at CBC. 155 kV, 260 µA, copper filter, source to object 213.079 mm, source to detector 1173.528 mm, 3142 projections, 2 frames per projection, Voxel size = 0.0363 mm. Total slices = 1764

**NHMUK A 905 (block containing humerus):** Nikon 49 Metrology XT H 225 ST High Resolution CT Scanner at CBC. 160 kV, 200 µA, copper filter, source to object 239.774 mm, source to detector 1173.528 mm, 1080 projections, 2 frames per projection, Voxel size = 0.0409 mm. Total slices = 1856

**NHMUK A 905 (block containing furcula and radius):** Nikon 49 Metrology XT H 225 ST High Resolution CT Scanner at CBC. 120 kV, 135 µA, copper filter, source to object 166.597 mm, source to detector 1173.528 mm, 1080 projections, 2 frames per projection, Voxel size = 0.0284 mm. Total slices = 1999

**NHMUK A 905 (block containing coracoid and scapula):** Nikon 49 Metrology XT H 225 ST High Resolution CT Scanner at CBC. 120 kV, 135 µA, copper filter, source to object 182.669 mm, source to detector 1173.528 mm, 1080 projections, 2 frames per projection, Voxel size = 0.0311 mm. Total slices = 1883

**NHMUK A 905 (scapula, coracoid and humerus):** Nikon 49 Metrology XT H 225 ST High Resolution CT Scanner at CBC. 120 kV, 135 µA, copper filter, source to object 187.085 mm, source to detector 1173.528 mm, 1080 projections, 2 frames per projection, Voxel size = 0.0319 mm. Total slices = 1999

**RMM 3394, 7844 and MSC 13868** (scanned together): NSI scanner at UTCT. Feinfocus source, high power, 200 kV, 0.13 mA, no filter, Perkin Elmer detector, 0.5 pF gain, 2 fps (499.893 ms integration time), no binning, no flip, source to object 133.681 mm, source to detector 1091.513 mm, continuous CT scan, no frames averaged, 0 skip frames, 3600 projections, 6 gain calibrations, 5 mm calibration phantom, data range [-10, 300] (grayscale range adjusted from NSI defaults), beam-hardening correction = 0.3. Voxel size = 0.0147 mm. Total slices = 942.

**RMM 2841 and MSC 13217** (scanned together): NSI scanner at UTCT. Feinfocus source, high power, 200 kV, 0.13 mA, no filter, Perkin Elmer detector, 0.5 pF gain, 2 fps (499.893 ms integration time), no binning, no flip, source to object 133.681 mm, source to detector 1091.513 mm, continuous CT scan, no frames averaged, 0 skip frames, 3600 projections, 6 gain calibrations, 5 mm calibration phantom, data range [-10, 300] (grayscale range adjusted from NSI defaults), beam-hardening correction = 0.3. Voxel size = 0.0147 mm. Total slices = 942.

**RMM 5794, RMM 7843 and MSC 13214** (scanned together): NSI scanner at UTCT. Feinfocus source, high power, 200 kV, 0.13 mA, no filter, Perkin Elmer detector, 0.5 pF gain, 2 fps (499.893 ms integration time), no binning, no flip, source to object 147 mm, source to detector 1091.513 mm, continuous CT scan, no frames averaged, 0 skip frames, 3600 projections, 6 gain calibrations, 5 mm calibration phantom, data range [-10, 300] (grayscale range adjusted from NSI defaults), beam-hardening correction = 0.3. Voxel size = 0.0188 mm. Total slices = 1188.

**RMM 5895 and 7842** (scanned together): NSI scanner at UTCT. Feinfocus source, high power, 200 kV, 0.13 mA, no filter, Perkin Elmer detector, 0.5 pF gain, 2 fps (499.893 ms integration time), no binning, no flip, source to object 149 mm, source to detector 1091.513 mm, continuous CT scan, no frames averaged, 0 skip frames, 3600 projections, 6 gain calibrations, 5 mm calibration phantom, data range [-10, 216] (grayscale range adjusted from NSI defaults), beam-hardening correction = 0.2. Voxel size = 0.0194 mm. Total slices = 1813.

**RMM 5916, 6201 and MSC 34426** (scanned together): NSI scanner at UTCT. Feinfocus source, high power, 200 kV, 0.13 mA, no filter, Perkin Elmer detector, 0.5 pF gain, 2 fps (499.893 ms integration time), no binning, no flip, source to object 133.681 mm, source to detector 1091.513 mm, continuous CT scan, no frames averaged, 0 skip frames, 3600 projections, 6 gain calibrations, 5 mm calibration phantom, data range [-10, 216] (grayscale range adjusted from NSI defaults), beam-hardening correction = 0.2. Voxel size = 0.0147 mm. Total slices = 1577.

**RMM 5937 and 6202** (scanned together): NSI scanner at UTCT. Feinfocus source, high power, 200 kV, 0.13 mA, no filter, Perkin Elmer detector, 0.5 pF gain, 2 fps (499.893 ms integration time), no binning, no flip, source to object 143 mm, source to detector 1091.513 mm, continuous CT scan, no frames averaged, 0 skip frames, 3600 projections, 6 gain calibrations, 5 mm calibration phantom, data range [-10, 243] (grayscale range adjusted from NSI defaults), beam-hardening correction = 0.2. Voxel size = 0.0175 mm. Total slices = 981.

**RMM 6200 and 7841** (scanned together): NSI scanner at UTCT. Feinfocus source, high power, 200 kV, 0.13 mA, no filter, Perkin Elmer detector, 0.5 pF gain, 2 fps (499.893 ms integration time), no binning, no flip, source to object 170 mm, source to detector 1091.513 mm, continuous CT scan, no frames averaged, 0 skip frames, 3600 projections, 6 gain calibrations, 5 mm calibration phantom, data range [-10, 216] (grayscale range adjusted from NSI defaults), beam-hardening correction = 0.3. Voxel size = 0.0258 mm. Total slices = 1869.

**YPM 1450 and 1740 (scanned together):** XTek CT XT H 225 Scanner at Harvard CNS. 255µA, 75kV, 1 second exposure, average 2 frames, 3200 views, no ring artifact correction, tungsten target, no filter.

**YPM 1724 and 1733 (scanned together):** XTek CT XT H 225 Scanner at Harvard CNS. 90µA, 95kV, 1 second exposure, average 2 frames, 3200 views, no ring artifact correction, tungsten target, no filter.

**YPM 1741:** XTek CT XT H 225 Scanner at Harvard CNS. 90µA, 95kV, 1 second exposure, average 2 frames, 3200 views, no ring artifact correction, tungsten target, no filter.

**IV: Comparative specimens**

Here we list all extant taxa specimens studied first-hand for comparative purposes in this work. Taxa mentioned in the main text and not listed here were compared based on the available literature. The listed specimens were studied from either first-hand examination or from CT-scan data. Uncatalogued specimens will eventually be added to the UMZC collections. Forelimb and hindlimb measurements for relevant specimens are provided in Supplemental Table 1.

**UMZC 187.AA** – *Alca torda*

**UMZC 191.B** – *Fratercula artica*

**UMZC 192.A** – *Uria sp.*

**UMZC 203** – *Gavia arctica*

**UMZC 206.B** – *Podiceps auritus*

**UMZC 209.A** – *Podica senegalensis*

**UMZC 211.A** – *Chauna chavaria*

**UMZC 224** – *Anas platyrhynchos*

**UMZC 242.EA** – *Anser albifrons*

**UMZC - 263** – *Fregata minor*

**UMZC 261.CA** – *Phaethon lepturus*

**UMZC 262.F** – *Morus bassanus*

**UMZC 265.A** – *Phalacrocorax carbo*

**UMZC 265.B** – *Phalacrocorax carbo*

**UMZC 268.a** – *Rynchops flavirostris*

**UMZC 269.BA** – *Sterna sumatrana*

**UMZC 271.B** – *Chlidonias niger*

**UMZC 272.A** – *Larus marinus*

**UMZC 274.1** – *Larus fuscus*

**UMZC 274.c** – *Chroicocephalus novaehollandiae*

**UMZC 284.AA** – *Puffinus iherminieri*

**UMZC 287.A** – *Macronectes giganteus*

**UMZC 287.D** – *Hydrobates leucorhous*

**UMZC 289.b** – *Daption capense*

**UMZC 290.A** – *Diomedea cauta*

**UMZC 298.a** – *Fulica atra*

**UMZC 304.A** – *Rallus striatus*

**UMZC 320.I** – *Charadrius rubricollis*

**UMZC 336.C** – *Ardea alba*

**UMZC 338.E** – *Eurypyga helias*

**UMZC 402** – *Gallus gallus*

**UMZC 404.B** – *Crypturellus variegatus*

**UMZC 411.b** – *Columba livia*

**UMZC 493.F** – *Podargus strigoides*

**UMZC 517.M** – *Cathartes aura*

**UMZC 907** – *Caprimulgus europaeus*

**Uncatalogued** – *Sterna hirundo*

**Uncatalogued** – *Macronectes giganteus (juvenile)*

**V: Phylogenetic analyses**

Phylogenetic methods used in this study are detailed in the Materials and Methods section. Phylogenetic results are described and discussed in the main text. Here we list all the analyses we performed, and the most relevant scoring changes and additions to the previously published morphological matrices of Huang et al. (2016) and Wang et al. (2020).

We incorporated five additional characters from Field et al. (2018) into both morphological matrices, and all taxa already present in both datasets were scored for these. Where these characters could not be scored, we incorporated the character codings from O'Connor et al. (2020), which incorporated those of Field et al. (2018) into a previous version of the Wang et al. (2020) dataset (Wang & Zhou, 2019).

No additional taxa were added to the Huang et al. (2016) dataset, but *Gansus zheni* was deleted based on it being considered synonymous with *Iteravis huchzermeyeri* (Wang et al. 2018). We therefore considered it to represent another specimen of the same taxon, so all character scorings present for *Gansus zheni* but missing for *Iteravis huchzermeyeri* were added to the latter. Additionally, several *Iteravis* characters were re-scored based on Wang et al. (2018). A single character from *Gansus* was re-scored based on Wang et al. (2016). All previous skull characters of *Jianchangornis microdonta* were re-scored as missing, after its skull was reported to be a chimera (O'Connor, 2019). Character 63 of *Longipteryx chaoyanensis* was found to be mistakenly scored as "8" in the Huang et al. (2016) dataset, despite the character having only two states. The complete list of changes is as follows:

*Iteravis huchzermeyeri*: 7:0, 8:0, 51:0, 54:2, 61:2/3, 76:2/3, 95:1, 102:0, 107:1, 109:0, 110:0, 111:1, 112:0, 113:1, 114:1, 115:0, 124:0, 125:0, 130:0, 136:0/1, 140:3, 155:0, 157:0, 165:0, 169:0, 176:0, 178:1, 180:1, 197:2, 222:0

*Gansus yumenensis*: 69: 1

*Jianchangornis microdonta*: 2:?, 4:?, 5:?, 6:?, 44:?

*Longipteryx chaoyanensis:* 63:1

In addition to the mistaken scoring for *Longipteryx chaoyanensis* mentioned above, multiple scoring errors were found in the Huang et al. (2016) dataset. These issues were corrected when found, but an exhaustive overhaul of that dataset is beyond the scope of this study. Many of these issues were already present in a previous version of that dataset from Liu et al. (2014), but not in Clarke (2004) or Clarke et al. (2006). Notably, five characters, 81, 157, 193, 151 and 214, were consistently scored for more states than they are stated to include. When possible, these characters were re-scored for all taxa based on the literature and previous versions of the dataset that were free of these issues; otherwise, they were scored as missing.

*Falkatakely forsterae* was added to the Wang et al. (2020) dataset; since it was used in an identical version of the dataset in O'Connor et al. (2020), no character re-scorings were needed. *Iberomesornis romerali* was present in previous versions of this dataset (Wang et al. 2017; Field et al. 2018) but absent in more recent versions of the matrix after Wang et al. (2018) added 18 new characters, so it was incorporated and scored for all new characters. Several taxa incorporated into recent versions of this dataset (Wang et al. 2018a,b, 2020, 2021; Wang & Zhou 2019) were not present in the version used by Field et al. (2018) nor in that of O'Connor (2020) and were scored for all of the new characters from Field et al. (2018). Changes and re-scorings were as follows:

*Iberomesornis romerali*: 263:?, 264:0, 265:?, 266:1, 267:?, 268:?, 269:1, 270?, 271:1, 272:0, 273:?, 274:?, 275:?, 276:0, 277:?, 278:?, 279:?, 280:?, 281:?, 282:?, 283:?, 284:?, 285:?

*Yangavis confucii*: 281:0, 282:?, 283:1, 284:0, 285:?

*Shangyang graciles*: 281:0, 282:?, 283:0, 284:?, 285:?

*Jinguofortis perplexus*: 281:0, 282:?, 283:0, 284:?, 285:?

*Mirusavis parvus*: 281:?, 282:?, 283:?, 284:?, 285:?

*Mengsciusornis dentatus*: 281:0, 282:?, 283:1, 284:?, 285:?

*Similiyanornis brevipectus*: 281:1, 282:?, 283:0, 284:?, 285:?

*Abitusavis lii*: 281:1?, 282:?, 283:0, 284:?, 285:?

*Chiappeavis magnapremaxillo*: 281:1, 282:?, 283:?, 284:0, 285:?

*Parapengornis eurycaudatus*: 281:0, 282:?, 283:0, 284:0, 285:?

*Eogranivora edentulata*: 281:1, 282:?, 283:?, 284:?, 285:?

*Gretcheniao sinensis*: 281:?, 282:?, 283:?, 284:?, 285:?

*Xinghaiornis lini*: 281:0, 282:?, 283:1, 284:0, 285:?

*Dingavis longmaxilla*: 281:0, 282:?, 283:1, 284:0, 285:?

*Piscivoroenantiornis inusitatus*: 281:0, 282:0, 283:0, 284:?, 285:?

Given the notable degree of morphological differences among the *Ichthyornis* specimens included in this study (see morphological descriptions and discussion), several of the new *Ichthyornis* specimens were scored as distinct operational taxonomic units (OTUs) in both matrices. Only well-preserved and/or partially complete specimens were scored; these were: AMLMNH PV 1993.2.133, BHI 6420 and 6421, FHSM VP 18702, KUVP 2281, 5969, 25469, 25472 and 119673. Additionally, the *Ichthyornis dispar* holotype YPM 1450 was scored as a separate OTU based on the literature to evaluate the affinities and phylogenetic relationships of the new specimens.

All specimens were combined into a single OTU after the results of the previous analyses recovered an exclusive clade including the *Ichthyornis* holotype, or in a polytomy comprising the holotype of *Ichthyornis* and the clade formed by Hesperornithes + crown birds (see supplementary trees). The differences between this combined *Ichthyornis dispar* OTU with that scored in previous versions of these datasets are as follows:

*Ichthyornis* Huang et al. (2016) dataset: 74:1, 80:0, 81:1/2, 83:1, 102:0, 104:0&1, 112:0/1, 143:2/3, 151:1, 160:0, 164:0, 166:1, 214:1, 215:2, 216:0

*Ichthyornis* Wang et al. (2020) dataset: 63:0, 64:4&5, 65:0, 66:1, 79:0, 97:0, 100:2, 105:0, 107:1, 111:2, 112:1, 113:0, 118:1, 128:1, 147:1, 153:2, 159:1&2, 160:1, 176:1, 186:1, 205:1, 208:1, 218:0, 228:1, 246:1, 263:1, 266:1, 273:0, 276:1

All analyses were performed under both Maximum Parsimony and Bayesian analytical frameworks, as described in the "Materials & Methods" section. Additionally, analyses were run both including and excluding *Apsaravis ukhaana*, after it was identified as a wildcard taxon with a high impact on the resulting tree topology and support values. The complete list of phylogenetic trees included in this supplement is as follows:

**Supplementary Tree 1:** Strict consensus of the 20 most parsimonious trees (best score 599) from the Huang et al. (2016) matrix, with all *Ichthyornis* specimens scored as distinct OTUs and including *Apsaravis*. Bootstrap values above 50% are shown. Consensus Tree Length: 672 steps; Consistency Index: 0.435; Retention Index: 0.749.

**Supplementary Tree 2:** Strict consensus of the 40 most parsimonious trees (best score 571) from the Huang et al. (2016) matrix, with all *Ichthyornis* specimens scored as distinct OTUs and excluding *Apsaravis*. Bootstrap values above 50% are shown. Consensus Tree Length: 580 steps; Consistency Index: 0.503; Retention Index: 0.805.

**Supplementary Tree 3:** 50% majority rule consensus tree derived from a Bayesian inference analysis of the Huang et al. (2016) matrix, with all *Ichthyornis* specimens scored as distinct OTUs and including *Apsaravis*. Posterior probability values are shown.

**Supplementary Tree 4:** 50% majority rule consensus tree derived from a Bayesian inference analysis of the Huang et al. (2016) matrix, with all *Ichthyornis* specimens scored as distinct OTUs and excluding *Apsaravis*. Posterior probability values are shown.

**Supplementary Tree 5:** Strict consensus of the 10 most parsimonious trees (best score 1448) from the Wang et al. (2020) matrix, with all *Ichthyornis* specimens scored as distinct OTUs and including *Apsaravis*. Bootstrap values above 50% are shown. Consensus Tree Length: 1463 steps; Consistency Index: 0.269; Retention Index: 0.674.

**Supplementary Tree 6:** Strict consensus of the 10 most parsimonious trees (best score 1417) from the Wang et al. (2020) matrix, with all *Ichthyornis* specimens scored as distinct OTUs and excluding *Apsaravis*. Bootstrap values above 50% are shown. Consensus Tree Length: 1429 steps; Consistency Index: 0.276; Retention Index: 0.679.

**Supplementary Tree 7:** 50% majority rule consensus tree derived from a Bayesian inference analysis of the Wang et al. (2020) matrix, with all *Ichthyornis* specimens scored as distinct OTUs and including *Apsaravis*. Posterior probability values are shown.

**Supplementary Tree 8:** 50% majority rule consensus tree derived from a Bayesian inference analysis of the Wang et al. (2020) matrix, with all *Ichthyornis* specimens scored as distinct OTUs and excluding *Apsaravis*. Posterior probability values are shown.

**Supplementary Tree 9:** Strict consensus of the 16 most parsimonious trees (best score 588) from the Huang et al. (2016) matrix, with a single combined *Ichthyornis* OTU and including *Apsaravis*. Bootstrap values above 50% are shown. Consensus Tree Length: 704 steps; Consistency Index: 0.411; Retention Index: 0.703.

**Supplementary Tree 10:** Strict consensus of the 2 most parsimonious trees (best score 560) from the Huang et al. (2016) matrix, with a single combined *Ichthyornis* OTU and excluding *Apsaravis*. Bootstrap values above 50% are shown. Consensus Tree Length: 563 steps; Consistency Index: 0.513; Retention Index: 0.798.

**Supplementary Tree 11:** 50% majority rule consensus tree derived from a Bayesian inference analysis of the Huang et al. (2016) matrix, with a single combined *Ichthyornis* OTU and including *Apsaravis*. Posterior probability values are shown.

**Supplementary Tree 12:** 50% majority rule consensus tree derived from a Bayesian inference analysis of the Huang et al. (2016) matrix, with a single combined *Ichthyornis* OTU and excluding *Apsaravis*. Posterior probability values are shown.

**Supplementary Tree 13:** Strict consensus of the 30 most parsimonious trees (best score 1432) from the Wang et al. (2020) matrix, with a single combined *Ichthyornis* OTU and including *Apsaravis*. Bootstrap values above 50% are shown. Consensus Tree Length: 1776 steps; Consistency Index: 0.222; Retention Index: 0.545.

**Supplementary Tree 14:** Strict consensus of the 10 most parsimonious trees (best score 1395) from the Wang et al. (2020) matrix, with a single combined *Ichthyornis* OTU and excluding *Apsaravis*. Bootstrap values above 50% are shown. Consensus Tree Length: 1404 steps; Consistency Index: 0.281; Retention Index: 0.661.

**Supplementary Tree 15:** 50% majority rule consensus tree derived from a Bayesian inference analysis of the Wang et al. (2020) matrix, with a single combined *Ichthyornis* OTU and including *Apsaravis*. Posterior probability values are shown.

**Supplementary Tree 16:** 50% majority rule consensus tree derived from a Bayesian inference analysis of the Wang et al. (2020) matrix, with a single combined *Ichthyornis* OTU and excluding *Apsaravis*. Posterior probability values are shown.

#### VI: Synapomorphies Diagnosing Key Clades

Here we list the synapomorphies diagnosing key clades from the analyses using both Huang et al. (2016) and Wang et al. (2020) morphological matrices. The synapomorphies shown here were optimized based on trees using the combined *Ichthyornis* OTU and excluding *Apsaravis* (see Materials & Methods).

##### Autapomorphic character combinations for *Ichthyornis*

Inferred under the results of maximum parsimony and Bayesian analysis using the Huang et al. (2016) matrix

- Osseous external naris considerably smaller than the antorbital fenestra (char. 12: 1 > 0)
- Dentary unforked posteriorly, or with a weakly developed dorsal ramus (char. 42: 1 > 0)
- Frontal/parietal suture fused (char. 51: 0 > 1)
- Sternum, coracoidal sulci crossed on midline (char. 75: 1 > 2)
- Ulna, distal end, dorsal condyle, dorsal trochlear surface, extent along posterior margin approximately equal in extent (char. 133: 0 > 1)
- Ulnar carpal present (char. 137: 0 > 1)
- Manual digit II, phalanx 2, "internal index process" (Stegmann, 1978) present on posterodistal edge (char. 153: 0 > 1)

Inferred under the results of maximum parsimony and Bayesian analysis using the Wang et al. (2020) matrix

- Premaxillae in adults completely fused (char. 1: 1 > 2)
- Cervical vertebrae variably dorsoventrally compressed, amphicoelous (biconcave: flat to concave articular surfaces) (char. 51: 1 > 0)
- Acrocoracoid process of coracoid hooked medially (char. 87: 0 > 1)
- Scapula, length as long as or longer than humerus (char. 97: 0 > 1)
- Sternum, coracoidal sulci crossed on midline (char. 115: 1 > 2)
- Major digit (II), phalanx 1, internal index process (Stegmann, 1978) present on caudodistal edge (char. 170: 0 > 1)

- Femoral trochanteric crest does not project beyond femoral head (char. 202: 1 > 2)
- Humerus, deltopectoral crest, distal end recedes abruptly with the humeral shaft (char. 253: 0 > 1)

**Inferred synapomorphies under the results of maximum parsimony and Bayesian analysis using the Huang et al. (2016) matrix**

**Ornithurae**

- Sternum, posterior margin exhibiting distinct posteriorly projected medial and/or lateral processes (char. 78: 2 > 1)
- Metacarpal I, anteroproximally-projected muscular process tip of process just surpasses the distal articular facet for phalanx 1 in anterior extent (char. 143: 1 > 2)
- Metacarpal I, shelf-like distal articulation with phalanx I (char. 145: 0 > 1)
- Ischium and pubis subparallel, pubis posteriorly directed (char. 158: 0 > 1)
- Postacetabular ilium, ventral surface, renal fossa developed (char. 164: 0 > 1)
- Ischium less than 1/3 of pubis length, extends farther than end of ischium (char. 167: 0 > 1)
- Pubis compressed mediolaterally (char. 168: 0 > 1)
- Pubes non-contacting distally (char. 169: 0 > 1)
- Tarsometatarsus, two proximal vascular foramina (char. 195: 1 > 2)
- Manual phalanx III:1, flexor tubercle present (char. 214: 0 > 1)
- Manual digit III, two phalanges (char. 215: 3 > 1)

**Hesperornithes + *Iaceornis* + Neornithes**

- Coronoid ossification absent (char. 18: 0 > 1)

- Cervical vertebrae heterocoelous anterior (i.e., mediolaterally concave, dorsoventrally convex) and posterior (i.e., mediolaterally convex, dorsoventrally concave) surfaces (char. 52: 1 > 2)
- Premaxilla, ventral surface covered with a smooth palatal shelf (char. 222: 0 > 1)
- Absent transverse ridges of bone (socketing) along potentially region of maxilla and dentary occupied by embryonic dental lamina (potentially dentigerous region) (char. 223: 0 > 1)
- Anterior margin of upper temporal region formed by reduced postorbital ossification directed ventrally but not laterally (char. 224: 0 > 1)
- Posterior margin of upper temporal region formed by reduced extension of squamosal enclosing a smaller adductor region and with a minor anterior extent (char. 225: 0 > 1)

***Iaceornis* + Neornithes**

- Sternum: raised, paired intermuscular ridges (linea intermuscularis; Baumel and Witmer, 1993) parallel to sternal midline present (char. 77: 0 > 1)
- Metacarpal I, anteroproximally-projected muscular process tip of extensor process conspicuously surpasses articular facet by approximately the width of facet, producing a pronounced knob (char. 143: 3 > 4)

**Neornithes**

- Sternum, pneumatic foramina present in the depressions (loculi costalis; Baumel and Witmer, 1993) between rib articulations (processi articularis sternocostalis; Baumel and Witmer [1993]) (char. 74: 0 > 1)
- Coracoid, pneumatized, Present (char. 91)
- Tibia/tarsal formed condyles, medial condyle projecting farther anteriorly than lateral (char. 181: 1 > 0)

**Inferred synapomorphies under the results of maximum parsimony and Bayesian analysis using the Wang et al. (2020) matrix**

**Avialae**

- No synapomorphies

**Euornithes**

- Procoracoid process present on coracoid (char. 89: 0 > 1)
- Sternal carina near to, or projecting rostrally from, the cranial border of the sternum (char. 110: 1 > 0)
- Humeral head globe shaped, craniocaudally convex (char. 122: 0 > 1)
- Proximal extension of metacarpal III: level with metacarpal II (char. 162: 1 > 0)
- Proximal phalanx of major digit (II) flat and craniocaudally expanded (char. 169: 0 > 1)
- Tibiotarsus twice the length of tarsometatarsus or more (char. 257: 1 -> 0)
- Omal tip of furcula tapered (char. 276: 0 > 1)

**Ornithurae**

- Sternum exhibiting costal facets (char. 116: 0 > 1)
- Deltopectoral crest of the humerus projected cranially (char. 128: 0 > 1)
- Ulnar carpal with well developed rami, U-shaped to V-shaped (char. 152: 0 -> 1)
- Ungual phalanx of major digit absent (char. 172: 0 > 1)
- Ischium more than two-thirds the length of the pubis (char. 188: 0 > 1)
- Pubic shaft laterally compressed throughout its length (char. 196: 0 > 1)
- Pubic apron absent (absence of symphysis) (char. 197: 0 > 1)
- Femur with distinct fossa for the capital ligament (char. 199: 0 > 1)
- Tarsometatarsal with two proximal vascular foramina (char. 226: 1 > 2)

- Tarsometatarsal intercotylar eminence well developed, low and rounded (char. 228: 0 > 1)

**Hesperornithes + Neornithes**

- Maxillary process of the premaxilla subequal or longer than the facial contribution of the maxilla (char. 2: 0 > 1)
- One or more pneumatic foramina piercing the centra of mid-cranial cervicals, caudal to the level of the parapophysis-diapophysis (char. 50: 1 > 0)
- Thoracic vertebrae series completely heterocoelous (char. 57: 0 > 1)
- Rostral margin of the sternum broad and rounded absent (char. 114: 1 > 0)
- Tibia/tarsal-formed condyles, no tapering of either condyle (char. 217: 0 > 1)
- Premaxilla, ventral surface covered with a smooth palatal shelf (char. 282: 0 > 1)
- Transverse ridges of bone (socketing) along potentially region of maxilla and dentary occupied by embryonic dental lamina (potentially dentigerous region) absent (char. 283: 0 > 1)
- Anterior margin of upper temporal region formed by reduced postorbital ossification directed ventrally but not laterally (char. 284: 0 > 1)
- Posterior margin of upper temporal region formed by reduced extension of squamosal enclosing a smaller adductor region and with a minor anterior extent (char. 285: 0 > 1)

**Neornithes**

- Sacral vertebrae, 15 or more ankylosed vertebrae (char. 64: 5 > 6)
- Ilium/ischium, distal co-ossification to completely enclose the ilioischiadic fenestra (char. 179: 0 > 1)

- Ischium with a proximodorsal (or proximocaudal) process present, developed as a small flange or raised scar contacting/fused with pubis and demarcating the obturator foramen distally (char. 190: 0 > 1)
- Extensor canal on tibiotarsus groove bridged by an ossified supratendinal bridge (char. 214: 1 > 2)
- Tarsometatarsus, projected surface and/or grooves on proximocaudal surface (associated with the passage of tendons of the pes flexors in Neornithes; hypotarsus) at least one groove enclosed by bone caudally (char. 229: 2 > 3)

#### VII: Morphological Character Descriptions

**Matrix from Huang et al. (2016) including additional characters added by Field et al. (2018).**

1. Premaxillae: (0) unfused in adults; (1) fused anteriorly in adults, posterior nasal [frontal] processes not fused to each other; (2) frontal processes completely fused as well as anterior premaxillae. (Ordered).

2. Premaxillary teeth: (0) present throughout premaxillae; (1) absent; (2) anteriorly reduced, posteriorly present; (3) anteriorly present, posteriorly reduced;

3. Maxillary teeth: (0) present; (1) absent.

4. Dentary teeth: (0) present; (1) absent.

5. Tooth crown serration: (0) present; (1) vestigial or absent.

6. Dentaries: (0) joined proximally by ligaments; (1) joined by bone.

7. Mandibular symphysis, two strong grooves forming an anteriorly-opening “v” in ventral view: (0) absent; (1) present.

8. Facial margin: (0) primarily formed by the maxilla, with the maxillary process of the premaxilla restricted to the anterior tip; (1) maxillary process of the premaxilla extending 1/2 facial margin; (2) maxillary process of the premaxilla extending more than 1/2 of facial margin. (Ordered).

9. Nasal [frontal] process of premaxilla: (0) short; (1) long, closely approaching frontal.

10. Nasal process of maxilla, dorsal ramus: (0) prominent, exposed medially and laterally; (1) absent or reduced to slight medial, and no lateral, exposure.

11. Nasal process of maxilla, participation of ventral ramus in anterior margin of antorbital fenestra in lateral view: (0) present, extensive; (1) small dorsal projection of the maxilla participates in the anterior margin of the antorbital fenestra, descending process of the nasals contacts premaxilla to exclude maxilla from narial margin; (2) no dorsal projection of maxilla participates in anterior margin of the antorbital fenestra. (Ordered).

12. Osseous external naris: (0) considerably smaller than the antorbital fenestra; (1) larger.

13. Ectopterygoid: (0) present; (1) absent.

14. Articulation between vomer and pterygoid: (0) present, well developed; (1) reduced, narrow process of pterygoid passes dorsally over palatine to contact vomer; (2) absent, pterygoid and vomer do not contact.

15. Palatine and pterygoid: (0) long, anteroposteriorly-overlapping, contact; (1) short, primarily dorsoventral, contact.

16. Palatine contacts: (0) maxillae only; (1) premaxillae and maxillae.

17. Vomer contacts premaxilla: (0) present; (1) absent.

18. Coronoid ossification: (0) present; (1) absent.

19. Projecting basisphenoid articulation with pterygoid: (0) present; (1) absent.

20. Basipterygoid processes: (0) long; (1) short (articulation with pterygoid sub-equal to, or longer than, amount projected from the basisphenoid rostrum).

21. Basisphenoid-pterygoid articulations: (0) located basal on basisphenoid; (1) located markedly anterior on basisphenoid (parasphenoid rostrum) such that the articulations are subadjacent on the narrow rostrum (the “rostropterygoid articulation” of Weber, 1993).

22. Basisphenoid/pterygoid articulation, orientation of contact: (0) anteroventral; (1) mediolateral; (2) entirely dorsoventral.

23. Pterygoid, articular surface for basisphenoid: (0) concave “socket,” or short groove enclosed by dorsal and ventral flanges; (1) flat to convex; (2) flat to convex facet, stalked, variably projected. (Ordered).

24. Pterygoid, kinked: (0) present, surface for basisphenoid articulation at high angle to axis of palatal process of pterygoid; (1) absent, articulation in line with axis of pterygoid.

25. Osseous interorbital septum (mesethmoid): (0) absent; (1) present.

26. Osseous interorbital septum (mesethmoid): (0) restricted to posterior or another just surpassing premaxillae/frontal contact in rostral extent does not surpass posterior edge of external nares in rostral extent; (1) extending rostral to posterior extent of frontal processes of premaxillae and rostral to posterior edge of external nares.

27. Eustachian tubes: (0) paired and lateral; (1) paired, close to cranial midline; (2) paired and adjacent on midline or single anterior opening.

28. Eustachian tubes ossified: (0) absent; (1) present.

29. Squamosal, ventral or "zygomatic” process: (0) variably elongate, dorsally enclosing otic process of the quadrate and extending anteroventrally along shaft of this bone, dorsal head of quadrate not visible in lateral view; (1) short, head of quadrate exposed in lateral view.

30. Orbital process of quadrate, pterygoid articulation: (0) pterygoid broadly overlapping medial surface of orbital process (i.e., “pterygoid ramus”); (1) restricted to anteromedial edge of process.

31. Quadrate, orbital process: (0) pterygoid articulates with anterior-most tip; (1) pterygoid articulation does not reach tip; (2) pterygoid articulation with no extent up orbital process, restricted to quadrate corpus. (Ordered).

32. Quadrate/ pterygoid contact: (0) as a facet, variably with slight anteromedial projection cradling base; (1) condylar, with a well-projected tubercle on the quadrate.

33. Quadrate, well-developed tubercle on anterior surface of dorsal process: (0) absent; (1) present.

34. Quadrate, quadratojugal articulation: (0) overlapping; (1) peg and socket articulation.

35. Quadrate, dorsal process, articulation: (0) with squamosal only; (1) with squamosal and prootic.

36. Quadrate, dorsal process, development of intercotylar incisure between prootic and squamosal cotylae: (0) absent, articular surfaces not differentiated; (1) two distinct articular facets, incisure not developed; (2) incisure present, “double headed.”

37. Quadrate, mandibular articulation: (0) bicondylar articulation with mandible; (1) tricondylar articulation, additional posterior condyle or broad surface.

38. Quadrate, pneumaticity: (0) absent; (1) present.

39. Quadrate, cluster of pneumatic foramina on posterior surface of the tip of dorsal process: (0) absent; (1) present.

40. Quadrate, pneumatization, large, single pneumatic foramen: (0) absent; (1) posteromedial surface of corpus.

41. Articular pneumaticity: (0) absent; (1) present.

42. Dentary strongly forked posteriorly: (0) unforked, or with a weakly developed dorsal ramus; (1) strongly forked with the dorsal and ventral rami approximately equal in posterior extent.

43. Splenial, anterior extent: (0) splenial stops well posterior to mandibular symphysis; (1) extending to mandibular symphysis, though non-contacting; (2) extending to proximal tip of mandible, contacting on midline.

44. Mandibular symphysis, anteroposteriorly extensive, flat to convex, dorsal-facing surface developed: (0) absent, concave; (1) flat surface developed.

45. Mandibular symphysis, symphysial foramina: (0) absent; (1) present.

46. Mandibular symphysis, symphysial foramen/foramina: (0) single; (1) paired.

47. Mandibular symphysis, symphysial foramen/foramina: (0) opening on posterior edge of symphysis; (1) opening on dorsal surface of symphysis.

48. Meckel’s groove: (0) not completely covered by splenial, deep and conspicuous medially; (1) covered by splenial, not exposed medially.

49. Anterior external mandibular fenestra: (0) absent; (1) present.

50. Jugal/postorbital contact: (0) present, (1) absent.

51. Frontal/parietal suture (0) open; (1) fused.

52. Cervical vertebrae: (0) variably dorsoventrally compressed, amphicoelous ("biconcave": flat to concave articular surfaces); (1) anterior surface heterocoelous (i.e., mediolaterally concave, dorsoventrally convex), posterior surface flat; (2) heterocoelous anterior (i.e., mediolaterally concave, dorsoventrally convex) and posterior (i.e., mediolaterally convex, dorsoventrally concave) surfaces. (Ordered).

53. Thoracic vertebrae (with ribs articulating with the sternum), one or more with prominent hypapophyses: (0) absent; (1) present. (This character does not address the presence of hypapophyses on transitional vertebrae, or “cervicothoracics”, that do not have associated ribs that articulate with the sternum [e.g., Gauthier, 1986; Chiappe, 1996]. In contrast, in Neornithes, well-developed hypapophyses are developed well into the thoracic series, on vertebrae with ribs articulating with the sternum.)

54. Thoracic vertebrae, count: (0) 12 or more; (1) 11; (2) 10 or fewer. (Ordered).

55. Thoracic vertebrae: (0) at least part of series with subround, central articular surfaces (e.g., amphicoelous/opisthocoelous) that lack the dorsoventral compression seen in heterocoelous vertebrae; (1) series completely heterocoelous.

56. Thoracic vertebrae, parapophyses: (0) rostral to transverse processes; (1) directly ventral to transverse processes (close to midpoint of vertebrae).

57. Thoracic vertebrae, centra, length, and midpoint width: (0) approximately equal in length and midpoint width; (1) length markedly greater than midpoint width.

58. Thoracic vertebrae, lateral surfaces of centra: (0) flat to slightly depressed; (1) deep, emarginate fossae; (2) central ovoid foramina.

59. Thoracic vertebrae with ossified connective tissue bridging transverse processes: (0) absent; (1) present.

60. Notarium: (0) absent; (1) present.

61. Sacral vertebrae, number ankylosed: (0) less than 7; (1) 7; (2) 8; (3) 9; (4) 10; (5) 11 or more; (6) 15 or more (Chiappe, 1996). (Ordered).

62. Sacral vertebrae, series of short vertebrae, with dorsally-directed parapophyses just anterior to the acetabulum: (0) absent; (1) present, three such vertebrae; (2) present, four such vertebrae. (Ordered).

63. Free caudal vertebrae, number: (0) more than 8; (1) 8 or less.

64. Caudal vertebrae, chevrons, fused on at least one anterior caudal: (0) present; (1) absent.

65. Free caudals; length of transverse processes: (0) subequal to width of centrum; (1) significantly shorter than centrum width.

66. Anterior free caudal vertebrae: (0) elongate pre/post-zygapophyses; (1) pre- and post-zygapophyses short and variably non-contacting; (2) prezygapophyses clasping the posterior surface of neural arch of preceding vertebra, postzygapophyses negligible. (Ordered).

67. Distal caudals: (0) unfused; (1) fused.

68. Fused distal caudals, morphology: (0) fused element length equal or greater than 4 free caudal vertebrae; (1) length less than 4 caudal vertebrae; (2) less than 2 caudal vertebrae in length. (Ordered).

69. Ossified uncinate processes: (0) absent; (1) present and unfused to ribs; (2) fused to ribs. (Ordered).

70. Gastralia: (0) present; (1) absent.

71. Ossified sternal plates: (0) unfused; (1) fused, flat; (2) fused, with slightly raised midline ridge; (3) fused with projected carina. (Ordered).

72. Carina or midline ridge: (0) restricted to posterior half of sternum; (1) approaches anterior limit of sternum.

73. Sternum, dorsal surface, pneumatic foramen (or foramina): (0) absent; (1) present.

74. Sternum, pneumatic foramina in the depressions (loculi costalis; Baumel and Witmer, 1993) between rib articulations (processi articularis sternocostalis; Baumel and Witmer [1993]): (0) absent; (1) present.

75. Sternum, coracoidal sulci spacing on anterior edge: (0) widely-separated mediolaterally, (1) adjacent; (2) crossed on midline.

76. Sternum, number of processes for articulation with the sternal ribs: (0) three; (1) four, (2) five; (3) six; (4) seven or more. (Ordered).

77. Sternum: raised, paired intermuscular ridges (linea intermuscularis; Baumel and Witmer, 1993) parallel to sternal midline: (0) absent; (1) present.

78. Sternum, posterior margin, distinct posteriorly projected medial and/or lateral processes: (0) absent (directly laterally projected zyphoid processes developed but not considered homologues as these are copresent with the posterior processes in the new clade); (1) with distinct posterior processes; (2) midpoint of posterior sternal margin connected to medial posterior processes to enclose paired fenestra. (Ordered).

79. Clavicles: (0) fused; (1) unfused.

80. Interclavicular angle (clavicles elongate): (0) greater than, or equal, to 90 degrees; (1) less than 90 degrees.

81. Furcula, hypocleideum: (0) absent; (1) a tubercle; (2) an elongate process. (Ordered).

82. Furcula, laterally excavated: (0) absent; (1) present.

83. Furcula, dorsal (omal) tip: (0) flat or blunt tip; (1) with a pronounced posteriorly pointed tip.

84. Furcula, ventral margin of apophysis: (0) curved, angling; (1) with a truncate or squared base.

85. Scapula and coracoid: (0) fused; (1) unfused.

86. Scapula and coracoid articulation: (0) pit-shaped scapular cotyla developed on the coracoid, and coracoidal tubercle developed on the scapula ("ball and socket" articulation); (1) scapular articular surface of coracoid convex; (2) flat.

87. Coracoid, procoracoid process: (0) absent; (1) present.

88. Coracoid: (0) height approximately equal mediolateral dimension; (1) height more than twice width, coracoid "strut-like."

89. Coracoid, lateral margin: (0) straight to slightly concave; (1) convex.

90. Coracoid, dorsal surface (= posterior surface of basal maniraptoran theropods): (0) strongly concave; (1) flat to convex.

91. Coracoid, pneumatized: (0) absent; (1) present.

92. Coracoid, pneumatic foramen: (0) proximal; (1) distal.

93. Coracoid, lateral process: (0) absent; (1) present.

94. Coracoid, ventral surface, lateral intermuscular line or ridge: (0) absent; (1) present.

95. Coracoid, glenoid facet: (0) dorsal to, or at approximately same level as, acrocoracoid process/ “biceps tubercle”; (1) ventral to acrocoracoid process.

96. Coracoid, acrocoracoid: (0) straight; (1) hooked medially.

97. Coracoid, n. supracoracoideus passes through coracoid: (0) present; (1) absent.

98. Coracoid, medial surface, area of the foramen n. supracoracoideus (when developed): (0) strongly depressed; (1) flat to convex.

99. Angle between coracoid and scapula at glenoid: (0) more than 90 degrees; (1) 90 degrees or less.1 probably but see orientation of scapular cotyla.

100. Scapula, posterior end: (0) wider or approximately the same width as proximal dorsoventral shaft width; (1) tapering distally.

101. Scapula: (0) straight, (1) dorsoventrally curved.

102. Scapula, length: (0) shorter than humerus, (1) as long as or longer than the humerus.

103. Scapula, acromion process: (0) projected anteriorly to surpass the articular surface for coracoid (facies articularis coracoidea; Baumel and Witmer, 1993); (1) projected less anteriorly than the articular surface for coracoid.

104. Scapula, acromion process: (0) straight; (1) laterally hooked tip.

105. Humerus and ulna, length: (0) humerus longer than ulna; (1) ulna and humerus approximately the same length; (2) ulna significantly longer than humerus. (Ordered).

106. Humerus, proximal end, head in anterior or posterior view: (0) strap-like, articular surface flat, no proximal midline convexity; (1) head domed proximally.

107. Humerus, proximal end, proximal projection: (0) dorsal edge projected farthest; (1) midline projected farthest.

108. Humerus, ventral tubercle and capital incisure: (0) absent; (1) present.

109. Humerus, capital incisure: (0) an open groove; (1) closed by tubercle associated with a muscle insertion just distal to humeral head.

110. Humerus, anterior surface, well-developed fossa on midline making proximal articular surface appear v-shaped in proximal view: (0) absent; (1) present.

111. Humerus, "transverse groove": (0) absent; (1) present, developed as a discreet, depressed scar on the proximal surface of the bicipital crest or as a slight transverse groove.

112. Humerus, deltopectoral crest: (0) projected dorsally (in line with the long axis of humeral head); (1) projected anteriorly.

113. Humerus, deltopectoral crest: (0) less than shaft width; (1) same width; (2) dorsoventral width greater than shaft width. (Ordered).

114. Humerus, deltopectoral crest, proximoposterior surface: (0) flat to convex; (1) concave.

115. Humerus, deltopectoral crest: (0) not perforate; (1) with a large fenestra.

116. Humerus, bicipital crest, pit-shaped scar/fossa for muscular attachment on anterodistal, distal or posterodistal surface of crest: (0) absent; (1) present.

117. Humerus, bicipital crest, pit-shaped fossa for muscular attachment: (0) anterodistal on bicipital crest; (1) directly ventrodistal at tip of bicipital crest; (2) posterodistal, variably developed as a fossa.

118. Humerus, bicipital crest: (0) little or no anterior projection; (1) developed as an anterior projection relative to shaft surface in ventral view; (2) hypertrophied, rounded tumescence. (Ordered).

119. Humerus, proximal end, one or more pneumatic foramina: (0) absent; (1) present.

120. Humerus, distal condyles: (0) developed distally; (1) developed on anterior surface of humerus.

121. Humerus, long axis of dorsal condyle: (0) at low angle to humeral axis, proximodistally oriented; (1) at high angle to humeral axis, almost transversely oriented.

122. Humerus, distal condyles: (0) sub-round, bulbous; (1) weakly defined, "strap-like."

123. Humerus, distal margin: (0) approximately perpendicular to long axis of humeral shaft; (1) ventrodistal margin projected significantly distal to dorsodistal margin, distal margin angling strongly ventrally (sometimes described as a well-projected flexor process).

124. Humerus, distal end, compressed anteroposteriorly and flared dorsoventrally: (0) absent; (1) present.

125. Humerus, brachial fossa: (0) absent; (1) present, developed as a flat scar or as a scar-impressed fossa.

126. Humerus, ventral condyle: (0) length of long axis of condyle less than the same measure of the dorsal condyle; (1) same or greater.

127. Humerus, demarcation of muscle origins (e.g., m. extensor metacarpi radialis in Neornithes) on the dorsal edge of the distal humerus: (0) no indication of origin as a scar, a pit, or a tubercle; (1) indication as a pit-shaped scar or as a variably projected scar-bearing tubercle or facet.

128. Humerus, distal end, posterior surface, groove for passage of m. scapulotriceps: (0) absent; (1) present.

129. Humerus, m. humerotricipitalis groove: (0) absent; (1) present as a ventral depression contiguous with the olecranon fossa.

130. Ulna, cotylae: (0) dorsoventrally adjacent; (1) widely separated by a deep groove.

131. Ulna, dorsal cotyla convex: (0) absent; (1) present.

132. Ulna, distal end, dorsal condyle, dorsal trochlear surface developed as a semilunate ridge: (0) absent; (1) present.

133. Ulna, distal end, dorsal condyle, dorsal trochlear surface, extent along posterior margin: (0) less than transverse measure of dorsal trochlear surface; (1) approximately equal in extent.

134. Ulna, bicipital scar: (0) absent; (1) developed as a slightly-raised scar; (2) developed as a conspicuous tubercle.

135. Ulna, brachial scar: (0) absent; (1) present.

136. Radius, ventroposterior surface: (0) smooth; (1) with muscle impression along most of surface; (2) deep longitudinal groove.

137. Ulnar carpal: (0) absent; (1) present.

138. Ulnar carpal: (0) “heart-shaped,” little differentiation into short dorsal and ventral rami; (1) V-shaped, well-developed dorsal and ventral rami.

139. Ulnar carpal, ventral ramus (crus longus, Baumel and Witmer, 1993): (0) shorter than dorsal ramus (crus brevis); (1) same length as dorsal ramus; (2) longer than dorsal ramus.

140. Semilunate carpal and metacarpals: (0) no fusion; (1) incomplete proximal fusion; (2) complete proximal fusion; (3) complete proximal and distal fusion. (Ordered).

141. Semilunate carpal, position relative to metacarpal I: (0) over 1/2 or more of proximal surface; (1) over less than 1/2 proximal surface. (Ordered)

142. Metacarpal III, anteroposterior diameter as a percent of same dimension of metacarpal II: (0) approximately equal or greater than 50%; (1) less than 50%.

143. Metacarpal I, anteroproximally-projected muscular process: (0) absent no distinct process visible; (1) small knob at anteroproximal tip of metacarpal; (2) tip of process just surpasses the distal articular facet for phalanx 1 in anterior extent; (3) tip of extensor process conspicuously surpasses articular facet by approximately half the width of facet, producing a pronounced knob; (4) tip of extensor process conspicuously surpasses articular facet by approximately the width of facet, producing a pronounced knob. (Ordered).

144. Metacarpal I, anterior surface: (0) roughly hourglass-shaped proximally, at least moderately expanded anteroposteriorly, and constricted just before flare of articulation for phalanx 1; (1) anterior surface broadly convex.

145. Metacarpal I, distal articulation with phalanx I: (0) ginglymoid; (1) shelf.

146. Pisiform process: (0) absent; (1) present.

147. Carpometacarpus, ventral surface, supratrochlear fossa deeply excavating proximal surface of pisiform process: (0) absent; (1) present.

148. Intermetacarpal space (between metacarpals II and III), (0) reaches proximally as far as the distal end of metacarpal I; (1) terminates distal to end of metacarpal I.

149. Carpometacarpus, distal end, metacarpals II and III, articular surfaces for digits: (0) metacarpal II sub-equal or surpasses metacarpal III in distal extent; (1) metacarpal III extends farther.

150. Intermetacarpal process or tubercle: (0) absent; (1) present as scar; (2) present as tubercle or flange. (Ordered).

151. Manual digit II, phalanx 1: (0) subcylindrical to subtriangular; (1) strongly dorsoventrally compressed, flat caudal surface.

152. Manual digit II, phalanges: (0) length of phalanx II-1 less than or equal to that of II-2; (1) longer.

153. Manual digit II, phalanx 2, "internal index process" (Stegmann, 1978) on posterodistal edge: (0) absent; (1) present (Clarke and Chiappe, 2001).

154. Ilium, ischium, pubis, proximal contact in adult: (0) unfused; (1) partial fusion (pubis not ankylosed); (2) completely fused. (Ordered).

155. Ilium/ischium, distal coossification to completely enclose the ilioischiadic fenestra: (0) absent; (1) present.

156. Ischium: (0) forked (dorsal process present); (1) straight, no dorsal process.

157. Ischium, dorsal process: (0) does not contact ilium; (1) contacts ilium.

158. Ischium and pubis: (0) not subparallel, pubis directed ventrally; (1) subparallel, pubis posteriorly directed; (2) pubis appressed to ischium. (Ordered).

159. Laterally projected process on ischiadic peduncle (antitrochanter): (0) directly posterior to acetabulum; (1) posterodorsal to acetabulum.

160. Preacetabular pectineal process (Baumel and Witmer, 1993): (0) absent; (1) present as a small flange; (2) present as a well-projected flange. (Ordered).

161. Preacetabular ilium: (0) approach on midline, open, or cartilaginous connection; (1) coossified, dorsal closure of "iliosynsacral canals".

162. Preacetabular ilium extends anterior to first sacral vertebrae: (0) no free ribs overlapped; (1) one or more ribs overlapped.

163. Postacetabular ilium: (0) dorsoventrally oriented; (1) mediolaterally oriented.

164. Postacetabular ilium, ventral surface, renal fossa developed: (0) absent; (1) present.

165. Ilium, m. cuppedicus fossa as broad, mediolaterally-oriented surface directly anteroventral to acetabulum: (0) present; (1) surface absent, insertion variably marked by a small entirely lateral fossa anterior to acetabulum.

166. Ischium, posterior demarcation of the obturator foramen: (0) absent; (1) present, developed as a small flange or raised scar contacting / fused with pubis and demarcating the obturator foramen distally.

167. Ischium, length relative to that of pubis: (0) 1/3 or greater total pubis length extends posterior to end of ishium; (1) less than 1/3 pubis extends farther than end of ishium.

168. Pubis: (0) sub-oval in cross section; (1) compressed mediolaterally.

169. Pubes, distal contact: (0) contacting, variably coossified into symphysis; (1) non-contacting.

170. Distal end of pubis: (0) expanded, flared; (1) straight, subequal, in proportion with rest of pubis.

171. Femur, fossa for insertion of lig. capitis femoris: (0) absent; (1) present.

172. Femur, posterior trochanter: (0) present, developed as a slightly projected tubercle or flange; (1) hypertrophied, "shelf-like" conformation (in combination with development of the trochanteric shelf; see Hutchinson, 2001); (2) absent (Chiappe, 1991). (Ordered).

173. Femur, lesser and greater trochanters: (0) separated by a notch; (1) developed as a single trochanteric crest.

174. Femur, patellar groove: (0) absent; (1) present.

175. Femur: (0) ectocondylar tubercle and lateral condyle separated by deep notch; (1) ectocondylar tubercle and lateral condyle form single trochlear surface.

176. Femur, posterior projection of the lateral border of the distal end, continuous with lateral condyle: (0) absent; (1) present.

177. Laterally-projected fibular trochlea: (0) absent; (1) present, developed as small notch; (2) a shelf-like projection. (Ordered).

178. Femur, popliteal fossa: (0) a groove open distally and bounded medially and laterally by narrow condyles; (1) closed distally by expansion of both condyles (primarily the medial).

179. Calcaneum and astragalus: (0) unfused to each other or tibia in adult; (1) fused to each other, unfused to tibia; (2) complete fused to each other and tibia. (Ordered).

180. Tibia, cnemial crest(s): (0) lateral crest only; (1) lateral and anterior crests developed.

181. Tibia/tarsal formed condyles: (0) medial condyle projecting farther anteriorly than lateral; (1) equal in anterior projection.

182. Tibia/tarsal formed condyles, extensor canal: (0) absent; (1) an emarginate groove; (2) groove bridged by an ossified supratendinal bridge. (Ordered).

183. Tibia/tarsal formed condyles, tuberositas retinaculi extensoris (Baumel and Witmer, 1993) indicated by short medial ridge or tubercle proximal to the condyles close to the midline and a more proximal second ridge on the medial edge: (0) absent; (1) present.

184. Tibia/tarsal formed condyles, mediolateral widths: (0) medial condyle wider; (1) approximately equal; (2) lateral condyle wider. (Ordered).

185. Tibia/tarsal formed condyles: (0) gradual sloping medial constriction of condyles; (1) no medial tapering of either condyle.

186. Tibia/tarsal formed condyles, intercondylar groove: (0) mediolaterally broad, approximately 1/3 width of anterior surface; (1) less than 1/3 width of total anterior surface.

187. Tibia, extension of articular surface for distal tarsals/tarsometatarsus: (0) no posterior extension of trochlear surface, or restricted to distal-most edge of posterior surface; (1) well-developed posterior extension, sulcus cartilaginis tibialis of Neornithes (Baumel and Witmer, 1993), distinct surface extending up the posterior surface of the

tibiotarsus; (2) with well-developed, posteriorly projecting, medial and lateral crests. (Ordered).

188. Tibia, distal-most mediolateral width: (0) wider than mid-point of shaft, giving distal profile a weakly developed triangular form; (1) approximately equal to shaft width, no distal expansion of whole shaft, although condyles may be variably splayed mediolaterally.

189. Fibula: (0) reaches tarsal joint articulating into distinct socket formed between the proximal tarsals and the tibia; (1) reduced in length, does not reach tarsal joint.

190. Distal tarsals and metatarsals, fusion: (0) distal tarsals fuse to metatarsals; (1) distal tarsals fuse to metatarsals and proximal metatarsals coossify; (2) distal tarsals fuse to metatarsals, and metatarsals fuse to each other proximally and distally; (3) extreme distal fusion, distal vascular foramen closed (Martin, 1983; Cracraft, 1986). (Ordered)

191. Metatarsal V: (0) present; (1) absent.

192. Metatarsal III: (0) proximally in plane with II and IV; (1) proximally displaced plantarly, relative to metatarsals II and IV.

193. Tarsometatarsus, intercotylar eminence: (0) absent; (1) well developed, globose.

194. Tarsometatarsus, projected surface or grooves on proximoposterior surface (associated with the passage of tendons of the pes flexors in Neornithes; hypotarsus): (0) absent; (1) developed as posterior projection with flat posterior surface; (2) projection, with distinct crests and grooves; (3) at least one groove enclosed by bone posteriorly. (Ordered).

195. Tarsometatarsus, proximal vascular foramen(foramina): (0) absent; (1) one, between metatarsals III and IV; (2) two. (Ordered).

196. Metatarsal I: (0) straight; (1) curved or distally deflected but not twisted, ventral surface convex “J shaped”; (2) deflected and twisted such that the ventromedial surface is concave proximal to trochlear surface for phalanx I. (Ordered).

197. Metatarsal II tubercle (associated with the insertion of the tendon of the m. tibialis cranialis in Neornithes): (0) absent; (1) present, on approximately the center of the proximodorsal surface of metatarsal II; (2) present, developed on lateral surface of metatarsal II, at contact with metatarsal III or on lateral edge of metatarsal III. (Ordered).

198. Metatarsal II, distal plantar surface, fossa for metatarsal I (fossa metatarsi I; Baumel and Witmer, 1993): (0) absent; (1) shallow notch; (2) conspicuous ovoid fossa. (Ordered).

199. Metatarsal II, articular surface for first phalanx: (0) ginglymoid; (1) rounded.

200. Metatarsals, relative mediolateral width : (0) metatarsal IV approximately the same width as metatarsals II and III; (1) metatarsal IV narrower than MII and MIII; (2) metatarsal IV greater in width than either metatarsal II or III.

201. Metatarsals, comparative trochlear width: (0) II approximately the same size as III and/or IV; (1) II wider than III and/or IV; (2) II narrower than III and/or IV.

202. Distal vascular foramen: (0) simple, with one exit; (1) forked, two exits (plantar and distal) between metatarsals III and IV.

203. Metatarsal III, trochlea in plantar view, proximal extent of lateral and medial edges of trochlea: (0) absent, trochlear edges approximately equal in proximal extent; (1) present, lateral edge extends farther.

204. Metatarsal II, distal extent of metatarsal II relative to metatarsal IV: (0) approximately equal in distal extent; (1) metatarsal II shorter than metatarsal IV, but

reaching distally farther than base of metatarsal IV trochlea; (2) metatarsal II shorter than metatarsal IV, reaching distally only as far as base of metatarsal IV trochlea. (Ordered).

205. Predentary element: (0) absent; (1) present.

206. Pygostyle, well developed transverse processes throughout length of pygostyle: (0) absent; (1) present.

207. Sternum: distal-most expansion or flare of the posterolateral process: (0) absent, tip of process subequal in width with the more proximal part of the process; (1) present, strongly flared, significantly broader than the more proximal part of the process.

208. Sternum, xiphoid process: (0) present; (1) absent.

209. Sternal incisures, anterior extent relative to rib articulations: (0) terminate sub-equal with posterior-most rib facet; (1) terminate well posterior to the posterior-most rib facet.

210. Sternal carina, location of apex: (0) posterior to anterior margin; (1) equal or surpassing anterior margin.

211. Coracoid, width at the sternal articulation: (0) broad flares markedly; (1) very narrow.

212. Coracoid, glenoid facet, position relative to the scapular cotyla: (0) extends to sternal margin of the cotyla in lateral view; (1) over half of the glenoid facet lies omal to the cotyla in lateral view.

213. Manual digit I, distal extent relative to metacarpal II (including ungual, I:2 if present): (0) digit I projects distal to distal end of metacarpal II; (1) terminates close to the terminus of metacarpal II; (2) over half the length of the metacarpal II but does not reach its distal end; (3) equal to, or less, than one half of the length of metacarpal II.

214. Manual phalanx III:1,flexor tubercle: (0) absent or indistinct, not developed as a tubercle; (1) present .

215. Manual digit III, number of phalanges: (0) four; (1) two; (2) one; (3) three.

216. Ilium, length of preacetabular ilium compared to postactabular ilium: (0) preactabular longer; (1) equal; (2) postacetabular ilium longer. (Ordered).

217. Pedal digits, relative length: (0) digit III longest, followed by digit IV, digit II shortest; (1) digit III almost equal to digit IV, digit II shortest; (2) digit IV longest, then digit III, digit II shortest. (Ordered)

218. Hallux, claw to phalanx proportions, 1:1: (0) equal to or longer; (1) shorter.

219. Pedal digit II, proportions of II:3 to II:1: (0) II: 3 subequal to or longer than II:1 and; (1) II:3 shorter.

220. Tail feathers, number of associated with free caudals: (0) more than 14, (1) none, but two associated with pygostyle; (2) none, but more than two associated with pygostyle.

**Characters added by Field et al. (2018):**

221. Premaxillae, extent along margin of jaw: (0) 1⁄3 or less of total length of rostrum; (1) between 1⁄3 and 1⁄2 of total length of rostrum; (2) over 1⁄2 of total length of rostrum. (Ordered)

222. Premaxillae, ventral surface: (0) deeply concave with exposed pair of neurovascular foramina; (1) covered with a smooth palatal shelf.

223. Transverse ridges of bone (socketing) along the region of maxilla and dentary occupied by embryonic dental lamina (potentially dentigerous region): (0) present; (1) absent.

224. Anterior margin of upper temporal region: (0) formed by extensive postorbital ossification with a primarily mediolateral, horizontal orientation, a flat upper table, and a posterior deflection of the tip; (1)formed by reduced postorbital ossification directed ventrally but not laterally.

225. Posterior margin of upper temporal region: (0) formed by extensive, ridged and buttressed squamosal process extending first laterally and then rostrally, with rostral portion elongate (longer than tall) and about the same length as the lateral extent; (1) formed by reduced extension of squamosal enclosing a smaller adductor region and with a minor anterior extent.

**Matrix from Wang et al. (2020), including additional characters added by Field et al. (2018)**

1. Premaxillae in adults: unfused (0); fused only rostrally (1); completely fused (2). (Ordered)

2. Maxillary process of the premaxilla: restricted to its rostral portion (0); subequal or longer than the facial contribution of the maxilla (1).

3. Frontal process of the premaxilla: short (0); relatively long, approaching the rostral border of the antorbital fenestra (1); very long, extending caudally near the level of lacrimals (2).

4. Premaxillary teeth: present throughout (0); present but rostral tip edentulous (1); present but restricted to rostral portion (2); absent (3).

5. Caudal margin of naris: far rostral than the rostral border of the antorbital fossa (0); nearly reaching or overlapping the rostral border of the antorbital fossa (1).

6. Naris longitudinal axis: considerably shorter than the long axis of the antorbital fossa (0); subequal or longer (1).

7. Maxillary teeth: present (0); absent (1).

8. Dorsal (ascending) ramus of the maxilla: present with two fenestrae (the promaxilllary and maxillary fenestra) (0); present with one fenestra (1); infenestrated (2); ramus absent (3). (Ordered)

9. Caudal margin of choana: located rostrally, not overlapping the region of the orbit (0); displaced caudally, at the same level or overlapping the rostral margin of the orbit (1).

10. Rostral margin of the jugal: away from the caudal margin of the naris (0); or very close to (leveled with) the caudal margin of the naris (1).

11. Contact between palatine and maxilla/premaxilla: palatine contact maxilla only (0); contacts premaxilla and maxilla (1).

12. Vomer and pterygoid articulation: present, well developed (0); reduced, narrow process of pterygoid passes dorsally over palatine to contact vomer (1); absent, pterygoid and vomer do not contact (2).

13. Jugal process of palatine: present (0); absent (1).

14. Contact between palatine and pterygoid: long, craniocaudally overlapping contact (0); short, primarily dorsoventral contact (1).

15. Contact between vomer and premaxilla: present (0); absent (1).

16. Ectopterygoid: present (0); absent (1).

17. Postorbital: present (0); absent (1).

18. Contact between postorbital and jugal: present (0); absent (1).

19. Quadratojugal: sutured to the quadrate (0); joined through a ligamentary articulation (1).

20. Lateral, round cotyla on the mandibular process of the quadrate (quadratojugal articulation): absent (0); present (1).

21. Contact between the quadratojugal and squamosal: present (0); absent (1).

22. Squamosal incorporated into the braincase, forming a zygomatic process: absent (0); present (1).

23. Squamosal, ventral or "zygomatic" process: variably elongate, dorsally enclosing otic process of the quadrate and extending cranioventrally along shaft of this bone, dorsal head of quadrate not visible in lateral view (0); short, head of quadrate exposed in lateral view (1).

24. Frontal/parietal suture in adults: open (0); fused (1).

25. Quadrate orbital process (pterygoid ramus): broad (0); sharp and pointed (1).

26. Quadrate pneumaticity: absent (0); present (1).

27. Quadrate: articulating only with the squamosal (0); articulating with both prootic and squamosal (1).

28. Otic articulation of the quadrate: articulates with a single facet (squamosal) (0); articulates with two distinct facets (prootic and squamosal) (1); articulates with two distinct facets and quadrate differentiated into two heads (2). (Ordered)

29. Quadrate distal end: with two transversely aligned condyles (0); with a triangular, condylar pattern, usually composed of three distinct condyles (1).

30. Basipterygoid processes: long (0); short (articulation with pterygoid subequal to, or longer than, amount projected from the basisphenoid rostrum) (1).

31. Pterygoid, articular surface for basipterygoid process: concave “socket”, or short groove enclosed by dorsal and ventral flanges (0); flat to convex (1); flat to convex facet, stalked, variably projected (2). (Ordered)

32. Eustachian tubes: paired, lateral, and well-separated from each other (0); paired, close to each other and to cranial midline or forming a single cranial opening (1).

33. Osseous interorbital septum (mesethmoid): absent (0); present (1).

34. Dentary teeth: present (0); absent (1).

35. Dentary tooth implantation: teeth in individual sockets (0); teeth in a communal groove (1).

36. Symphysial portion of dentaries: unfused (0); fused (1).

37. Deeply notched rostral end of the mandibular symphysis: absent (0); present (1).

38. Mandibular symphysis, symphyseal foramina: absent (0); single (1); paired (2).

39. Mandibular symphysis, symphyseal foramen/foramina: opening on caudal edge of symphysis (0); opening on dorsal surface of symphysis (1).

40. Small ossification present at the rostral tip of the mandibular symphysis (intersymphysial ossification): absent (0); present (1).

41. Caudal margin of dentary strongly forked: unforked, or with a weakly developed dorsal ramus (0); strongly forked with the dorsal and ventral rami approximately equal in caudal extent (1).

42. Mandibular ramus sigmoidal such that the rostral tip is dorsally convex and the caudal end is dorsally concave: absent (0); present (1).

43. Cranial extent of splenial: stops well caudal to mandibular symphysis (0); extending to mandibular symphysis, though non-contacting (1); extending to proximal tip of mandible, contacting on midline (2). (Ordered)

44. Meckel's groove (medial side of mandible): not completely covered by splenial, deep and conspicuous medially (0); covered by splenial, not exposed medially (1).

45. Rostral mandibular fenestra: absent (0); present (1).

46. Caudal mandibular fenestra: present (0); absent (1).

47. Articular pneumaticity: absent (0); present (1).

48. Teeth: serrated crowns (0); unserrated crowns (1).

49. Atlantal hemiarches in adults: unfused (0); fused, forming a single arch (1).

50. One or more pneumatic foramina piercing the centra of mid-cranial cervicals, caudal to the level of the parapophysis-diapophysis: present (0); absent (1).

51. Cervical vertebrae: variably dorsoventrally compressed, amphicoelous ("biconcave": flat to concave articular surfaces) (0); cranial surface heterocoelous (i.e., mediolaterally concave, dorsoventrally convex), caudal surface flat or slightly concave (1); heterocoelous cranial (i.e., mediolaterally concave, dorsoventrally convex) and caudal (i.e., mediolaterally convex, dorsoventrally concave) surfaces (2). (Ordered)

52. Prominent carotid processes in the intermediate cervicals: absent (0); present (1).

53. Postaxial cervical epipophyses: prominent, projecting further back from the postzygapophysis (0); weak, not projecting further back from the postzygapophysis, or absent (1).

54. Keel-like ventral surface of cervical centra: absent (0); present (1).

55. Prominent (50% or more the height of the centrum's cranial articular surface) ventral processes of the cervicothoracic vertebrae: absent (0); present (1).

56. Thoracic vertebral count: 13-14 (0); 11-12 (1); fewer than 11 (2). (Ordered)

57. Thoracic vertebrae: at least part of series with subround, central articular surfaces (e.g., amphicoelous/opisthocoelous) that lack the dorsoventral compression seen in heterocoelous vertebrae (0); series completely heterocoelous (1).

58. Caudal thoracic vertebrae, centra, length and midpoint width: approximately equal in length and midpoint width (0); length markedly greater than midpoint width (1).

59. Wide vertebral foramen in the mid-caudal thoracic vertebrae, vertebral foramen/articular cranial surface ratio (vertical diameter) larger than 0.40: absent (0); present (1).

60. Hyposphene-hypantrum accessory intervertebral articulations in the thoracic vertebrae: present (0); absent (1).

61. Lateral side of the thoracic centra: weakly or not excavated (0); deeply excavated by a groove (1); excavated by a broad fossa (2).

62. Cranial thoracic vertebrae, parapophyses: located in the cranial part of the centra of the thoracic vertebrae (0); located in the central part of the centra of the thoracic vertebrae (1).

63. Notarium: absent (0); present (1).

64. Sacral vertebrae, number ankylosed (synsacrum): less than 7 (0); 7 (1); 8 (2); 9 (3); 10 (4); 11 or more (5); 15 or more (6). (Ordered)

65. Synsacrum, procoelous articulation with last thoracic centrum (deeply concave facet of synsacrum receives convex articulation of last thoracic centrum): absent (0); present (1).

66. Cranial vertebral articulation of first sacral vertebra: approximately equal in height and width (0); wider than high (1).

67. Series of short sacral vertebrae with dorsally directed parapophyses just cranial to the acetabulum: absent (0); present, three such vertebrae (1); present, four such vertebrae (2). (Ordered)

68. Convex caudal articular surface of the synsacrum: absent (0); present (1).

69. Degree of fusion of distal caudal vertebrae: fusion absent (0); few vertebrae partially ankylosed (intervening elements are well-discernable) (1); vertebrae completely fused into a pygostyle (2). (Ordered)

70. Free caudal vertebral count: more than 35 (0); 35-26 (1); 25 - 20 (2); 19-9 (3); 8 or less (4). (Ordered)

71. Procoelous caudals: absent (0); present (1).

72. Distal caudal vertebra prezygapophyses: elongate, exceeding the length of the centrum by more than 25% (0); shorter (1); absent (2). (Ordered)

73. Free caudals, length of transverse processes: approximately equal to, or greater than, centrum width (0); significantly shorter than centrum width (1).

74. Proximal haemal arches: elongate, at least 3 times longer than wider (0); shorter (1); absent (2). (Ordered)

75. Pygostyle: longer than or equal to the combined length of the free caudals (0); shorter (1).

76. Cranial end of pygostyle dorsally forked: absent (0); present (1).

(rescored as 1).

77. Cranial end of pygostyle with a pair of laminar, ventrally projected processes: absent (0); present (1).

78. Distal constriction of pygostyle: absent (0); present (1).

79. Ossified uncinate processes in adults: absent (0); present and free (1); present and fused (2).

80. Uncinate process, orientation: perpendicular to rib (0); angled dorsally defining an acute angle with the rib (1).

81. Gastralia: present (0); absent (1).

82. Coracoid shape: rectangular to trapezoidal in profile (0); strut-like (1).

83. Coracoid and scapula articulation: pit-shaped scapular cotyla developed on the coracoid, and coracoidal tubercle developed on the scapula ("ball and socket" articulation) (0); scapular articular surface of coracoid convex (1); (2) flat.

84. Scapula: articulated at the shoulder (proximal) end of the coracoid (0); well below it (1).

85. Coracoid, humeral articular (glenoid) facet: dorsal to acrocoracoid process/"biceps tubercle" (0); ventral to acrocoracoid process (1).

86. Humeral articular facets of the coracoid and the scapula: placed in the same plane (0); forming a sharp angle (1).

87. Coracoid, acrocoracoid: straight (0); hooked medially (1).

88. Laterally compressed shoulder end of coracoid, with nearly aligned acrocoracoid process, humeral articular surface, and scapular facet, in dorsal view: absent (0); present (1).

89. Procoracoid process on coracoid: absent (0); present (1).

90. Lateral margin of coracoid: concave (0); nearly concave to straight for most part and the convex portion is restricted at sternal end, which measures less than half the width of sternal end (1); strongly convex, and the convex portion measuring more than half the sternal end (2).

91. Broad, deep fossa on the dorsal surface of the coracoid (dorsal coracoidal fossa): absent (0); present (1).

92. Supracoracoidal nerve foramen of coracoid: centrally located (0); displaced toward (often as an incisure) the medial margin of the coracoid (1); displaced so that it no longer passes through the coracoid (absent) (2). (Ordered).

93. Coracoid, medial surface, strongly depressed elongate furrow at the level of the passage of n. supracoracoideus: absent (0); present (1).

94. Supracoracoidal nerve foramen, location relative to dorsal coracoidal fossa: above fossa (0); inside fossa (1).

95. Coracoid, sternolateral corner: unexpanded (0); expanded (1); well-developed squared-off lateral process (sternocoracoidal process) (2); present and with a distinct omal projection (hooked) (3).

96. Scapular shaft: straight, both dorsal and ventral margins straight (0); straight shaft with convex dorsal margin and straight ventral margin (1); the scapular shaft sagittally curved (2).

97. Scapula, length: shorter than humerus (0); as long as or longer than humerus (1).

98. Scapular acromion process: in lateral or costal view, strongly projecting craniodorsally, forming a large angle with the proximal shaft of the scapular (0); nearly parallel to the shaft of the scapular (1).

99. Scapula, acromion process: projected cranially surpassing the articular surface for coracoid (0); projected less cranially than the articular surface for coracoid (1).

100. Scapula, acromion process, in costal or lateral aspect: straight and tapered toward cranial end (0); barely tapered with a blunt end (1); laterally hooked tip (2).

101. Proximal end of scapula, pit between acromion and humeral articular facet (scapular fossa): absent (0); present (1).

102. Costal surface of scapular blade with prominent longitudinal furrow: absent (0); present (1).

103. Scapular caudal end: blunt (may or may not be expanded) (0); sharply tapered (1).

104. Furcular, shape: boomerang-shaped (0); V to Y-shaped (1); U-shaped (2).

105. Furcula interclavicular angle: approximately 90° (0); less than 70° (1). The interclavicular angle is measured as the angle formed between three points, one at the omal end of each rami and the apex located at the clavicular symphysis.

106. Dorsal and ventral margins of the furcula: subequal in width (0); ventral margin distinctly wider than the dorsal margin so that the furcular ramus appears concave laterally (1).

107. Hypocleideum: absent (0); present as a tubercle or short process (1); present as an elongate process approximately 30% rami length (2); hypertrophied, exceeding 50% rami length (3). (Ordered)

108. Sternum: unossified (0); partially ossified, coracoidal facets cartilaginous (1); fully ossified (2).

109. Ossified sternum: two flat plates (0); single flat element (1); single element, with slightly raised midline ridge (2); single element, with projected carina (3).

110. Sternal carina: near to, or projecting rostrally from, the cranial border of the sternum (0); not reaching the cranial border of the sternum (1).

111. Sternum, caudal margin, number of paired caudal trabecula: none (0); one (1); two (2).

112. Sternum, outermost trabecula, shape: tips terminate cranial to caudal end of sternum (0); tips terminate at or approaching caudal end of sternum (1); tips extend caudally past the termination of the sternal midline (2).

113. Prominent distal expansion in the outermost trabecula of the sternum: absent (0); present, simple bulb-like (1); fan-shaped expansion (2); triangular expansion with an acute medial angle (3); branched (4).

114. Rostral margin of the sternum broad and rounded: absent (0); present (1).

115. Sternum, coracoidal sulci spacing on cranial edge: widely separated mediolaterally (0); adjacent (1); crossed on midline (2).

116. Costal facets of the sternum: absent (0); present (1).

117. Sternal costal processes: three (0); four (1); five (2); six (3); seven (4); eight (5). (Ordered)

118. Sternal midline, caudal end: blunt W-shape (0); V-shape (1); elongate straight projection (xiphoid process) (2); xiphoid process slightly flared mediolaterally (3); xiphoid process distal end strongly flared with prominent medial and lateral projections (4); rounded (5).

119. Sternum, caudal half, paired enclosed fenestra: absent (0); present (1).

120. Sternum, dorsal surface, pneumatic foramen (or foramina): absent (0); present (1).

121. Proximal and distal humeral ends: twisted (0); expanded nearly in the same plane (1).

122. Humeral head: concave cranially and convex caudally (0); globe shaped, craniocaudally convex (1).

123. Proximal margin of the humeral head concave in its central portion, rising ventrally and dorsally: absent (0); present (1).

124. Humerus, proximocranial surface, well-developed circular fossa on midline: absent (0); present (1).

125. Humerus with distinct transverse ligamental groove: absent (0); present (1).

126. Humerus, ventral tubercle projected caudally, separated from humeral head by deep capital incision: absent (0); present (1).

127. Pneumatic fossa in the caudoventral corner of the proximal end of the humerus: absent or rudimentary (0); well developed (1).

128. Humerus, deltopectoral crest: projected dorsally (the plane of the crest is coplanar to the cranial surface of the humerus) (0); projected cranially (1).

129. Humerus, deltopectoral crest: less than shaft width (0); approximately same width (1); prominent and subquadrangular (i.e., subequal length and width) (2).

130. Humerus, deltopectoral crest, perforated by a large fenestra: absent (0); present (1).

131. Humerus, bicipital crest: little or no cranial projection (0); developed as a cranial projection relative to shaft surface in ventral view (1); hypertrophied, rounded tumescence (2).

132. Humerus, distal end of bicipital crest, pit-shaped fossa for muscular attachment: absent (0); craniodistal on bicipital crest (1); directly ventrodistal at tip of bicipital crest (2); caudodistal, variably developed as a fossa (3).

133. Distal end of the humerus very compressed craniocaudally: absent (0); present (1).

134. Humerus, demarcation of muscle origins (e.g., m. extensor metacarpi radialis in Neornithes) on the dorsal edge of the distal humerus: no indication (0); a pit or a tubercle (1); a variably projected scar-bearing tubercle (dorsal supracondylar process) (2).

135. Well-developed brachial depression on the cranial face of the distal end of the humerus: absent (0); present (1).

136. Well-developed olecranon fossa on the caudal face of the distal end of the humerus: absent (0); present (1).

137. Humerus, distal end, caudal surface, groove for passage of m. scapulotriceps: absent (0); present (1).

138. Humerus, m. humerotricipitalis groove: absent (0); present as a well-developed ventral depression contiguous with the olecranon fossa (1).

139. Humerus, distal margin: approximately perpendicular to long axis of humeral shaft (0); ventrodistal margin projected significantly distal to dorsodistal margin, distal margin angling strongly ventrally (sometimes described as a well-projected flexor process) (1).

140. Humeral distal condyles: mainly located on distal aspect (0); on cranial aspect (1).

141. Humerus, long axis of dorsal condyle: at low angle to humeral axis, proximodistally oriented (0); at high angle to humeral axis, almost transversely oriented (1).

142. Humerus, distal condyles: subround, bulbous (0); weakly defined, “straplike” (1).

143. Humerus, ventral condyle: length of long axis of condyle less than the same measure of the dorsal condyle (0); same or greater (1).

144. Ulna: shorter than humerus (0); nearly equivalent to or longer than humerus (1).

145. Ulnar shaft, radial-shaft/ulnar-shaft ratio: larger than 0.70 (0); smaller than 0.70 (1).

146. Ulna, cotylae: dorsoventrally adjacent (0); widely separated by a deep groove (1).

147. Ulna, dorsal cotyla strongly convex: absent (0); present (1).

148. Ulna, bicipital scar: absent (0); developed as a slightly raised scar (1); developed as a conspicuous tubercle (2).

149. Proximal end of the ulna with a well-defined area for the insertion of m. brachialis anticus: absent (0); present (1).

150. Semilunate ridge on the dorsal condyle of the ulna: absent (0); present (1).

151. Shaft of radius with a long longitudinal groove on its ventrocaudal surface: absent (0); present (1).

152. Ulnar carpal: heart-shaped with little differentiation into short rami (0); U-shaped to V-shaped, well-developed rami (1).

153. Ulnar carpal, ventral ramus: shorter than dorsal ramus (crus brevis) (0); same length as dorsal ramus (1); longer than dorsal ramus (2).

154. Semilunate carpal and proximal ends of metacarpals in adults: unfused (0); semilunate fused to the alular (I) metacarpal (1); semilunate fused to the major (II) and minor (III) metacarpals (2); fusion of semilunate and all metacarpals (3). Any specimen that is inferred to be a juvenile should be scored as a "?" in order to account for the possibility of ontogenetic change.

155. Semilunate carpal, position relative to the alular metacarpal (I): over entire proximal surface (0); over less than one-half proximal surface or no contact present (1).

156. Carpometacarpus, proximal ventral surface: flat (0); raised ventral projection contiguous with minor metacarpal (1); pisiform process forming a distinct peg-like projection (2).

157. Carpometacarpus, ventral surface, supratrochlear fossa deeply excavating proximal surface of pisiform process: absent (0); present (1).

158. Round-shaped alular metacarpal (I): absent (0); present (1).

159. Alular metacarpal (I), extensor process: absent, no cranioproximally projected muscular process (0); present, tip of extensor process just surpassed the distal articular facet for phalanx 1 in cranial extent (1); tip of extensor process conspicuously surpasses articular facet by approximately half the width of facet, producing a pronounced knob (2); tip of extensor process conspicuously surpasses articular facet by approximately the width of facet, producing a pronounced knob (3). (Ordered)

160. Alular metacarpal (I), distal articulation with phalanx I: ginglymoid (0); shelf (1); ball-like (2).

161. Metacarpal III, craniocaudal diameter as a percentage of same dimension of metacarpal II: approximately equal or greater than 50% (0); less than 50% (1).

162. Proximal extension of metacarpal III: level with metacarpal II (0); ending distal to proximal surface of metacarpal II (1).

163. Intermetacarpal process or tubercle on metacarpal II: absent (0); present as scar (1); present as tubercle or flange (2).

164. Intermetacarpal space: absent or very narrow (0); at least as wide as the maximum width of minor metacarpal (III) shaft (1).

165. Intermetacarpal space: reaches proximally as far as the distal end of metacarpal I (0); terminates distal to end of metacarpal I (1).

166. Distal end of metacarpals: unfused (0); partially or completely fused (1).

167. Minor metacarpal (III) projecting distally more than the major metacarpal (II): absent (0); present (1).

168. Alular digit (I), phalanx 1, distal extension relative to the major metacarpal (II): beyond the distal end of major metacarpal (0); approximately equal in distal extension (1); shorter than the distal end but beyond half of the major metacarpal (2); terminating less than half of the major metacarpal (3). (Ordered)

169. Proximal phalanx of major digit (II): of normal shape (0); flat and craniocaudally expanded (1).

170. Major digit (II), phalanx 1, “internal index process” (Stegmann, 1978) on caudodistal edge: absent (0); present (1).

171. Second phalanx of major digit (II): longer than proximal phalanx (0); shorter than or equivalent to proximal phalanx (1).

172. Ungual phalanx of major digit (II): present (0); absent (1).

173. Ungual phalanx of major digit (II): larger or subequal to other manual unguals (0); smaller than the alular ungual but larger than that of the minor (III) digit, and the ungual of the minor digit may or may not present (1); smaller than the unguals of the alular and minor digits (2).

174. Proximal phalanx of the minor digit (III) much shorter than the remaining non-ungual phalanges of this digit: absent (0); present (1).

175. Ungual phalanx of minor digit (III): present (0); absent (1).

176. Length of manus (semilunate carpal + major metacarpal and digit) relative to humerus: longer (0); subequal (1); shorter (2). (Ordered)

177. Intermembral index = (length of humerus + ulna)/(length of femur + tibiotarsus): less than 0.7, flightless (0); between 0.7 and 0.9 (1); between 0.9 and 1.1 (2); greater than 1.1 (3).

178. Pelvic elements in adults, at the level of the acetabulum: unfused or partial fusion (0); completely fused (1).

179. Ilium/ischium, distal co-ossification to completely enclose the ilioischiadic fenestra: absent (0); present (1).

180. Preacetabular process of ilium twice as long as postacetabular process: absent (0); present (1).

181. Preacetabular ilium: approach on midline, open, or cartilaginous connection (0); co-ossified, dorsal closure of “iliosynsacral canals” (1).

182. Ilium, m. cuppedicus fossa as broad, mediolaterally oriented surface directly cranioventral to acetabulum: present (0); surface absent, insertion variably marked by a small entirely lateral fossa cranial to acetabulum (1).

183. Preacetabular pectineal process (Baumel and Witmer, 1993): absent (0); present as a small flange (1); present as a well-projected flange (2). (Ordered)

184. Small acetabulum, acetabulum/ilium length ratio equal to or smaller than 0.11: absent (0); present (1).

185. Prominent antitrochanter: caudally directed (0); caudodorsally directed (1).

186. Postacetabular process shallow, less than 50% of the depth of the preacetabular wing at the acetabulum: absent (0); present (1).

187. Iliac brevis fossa: present (0); absent (1).

188. Ischium: two-thirds or less the length of the pubis (0); more than two-thirds the length of the pubis (1).

189. Obturator process of ischium: prominent (0); reduced or absent (1).

190. Ischium, caudal demarcation of the obturator foramen: absent (0); present, developed as a small flange or raised scar contacting/fused with pubis and demarcating the obturator foramen distally (1).

191. Ischium with a proximodorsal (or proximocaudal) process: absent (0); present (1).

192. Ischiadic terminal processes forming a symphysis: present (0); absent (1).

193. Orientation of proximal portion of pubis: cranially to subvertically oriented (0); retroverted, separated from the main synsacral axis by an angle ranging between $65^{\circ}$ and $45^{\circ}$ (1); more or less parallel to the ilium and ischium (2). (Ordered)

194. Pubic pedicel: cranioventrally projected (0); ventrally or caudoventrally projected (1).

195. Pubic pedicel of ilium very compressed laterally and hook-like: absent (0), present (1).

196. Pubic shaft laterally compressed throughout its length: absent (0); present (1).

197. Pubic apron: present (0); absent (absence of symphysis) (1).

198. Pubic foot: flaring into simple round shape (0); triangular shape with a pointed caudal tip and caudoventrally directed with respect to the distal pubic shaft (1); the caudal tip recurved caudodorsally with respect to the distal pubic shaft (2); absent (3).

199. Femur with distinct fossa for the capital ligament: absent (0); present (1).

200. Femoral neck: present (0); absent (1).

201. Femoral anterior trochanter: separated from the greater trochanter (0); fused to it, forming a trochanteric crest with a laterally curved edge (1); fused to it, forming a trochanteric crest with a flattened edge (2).

202. Femoral trochanteric crest: projects proximally beyond femoral head (0); equal in proximal projection (1); does not project beyond femoral head (2).

203. Femoral posterior trochanter: present, developed as a slightly projected tubercle or flange (0); hypertrophied, “shelf-like” conformation (1); absent (2).

204. Femur with prominent patellar groove: absent (0); present as a continuous extension onto the distal shaft (1); present and separated from the shaft by a slight ridge, giving it a pocketed appearance (2).

205. Femur: ectocondylar tubercle and lateral condyle separated by deep notch (0); ectocondylar tubercle and lateral condyle contiguous but without developing a tibiofibular crest (1); tibiofibular crest present, defining laterally a fibular trochlea (2). (Ordered)

206. Caudal projection of the lateral border of the distal end of the femur, proximal and contiguous to the ectocondylar tubercle/tibiofibular crest: absent (0); present (1).

207. Femoral popliteal fossa distally bounded by a complete transverse ridge: absent (0); present (1).

208. Fossa for the femoral origin of m. tibialis cranialis: absent (0); present (1).

209. Tibia, calcaneum, and astragalus: unfused or poorly co-ossified (sutures still visible) (0); complete fusion of tibia, calcaneum, and astragalus (1).

210. Round proximal articular surface of tibiotarsus: absent (0); present (1).

211. Tibiotarsus, proximal articular surface: flat (0); angled so that the medial margin is elevated with respect to the lateral margin (1).

212. Tibiotarsus, cnemial crests: absent (0); present, one (1); present, two (2).

213. Tibia, caudal extension of articular surface for distal tarsals/tarsometatarsus: absent, articular restricted to distalmost edge of caudal surface (0); well-developed caudal extension, sulcus cartilaginis tibialis of Neornithes, distinct surface extending up the caudal surface of the tibiotarsus (1); with well-developed, caudally projecting medial and lateral crests (2). (Ordered)

214. Extensor canal on tibiotarsus: absent (0); present as an emarginate groove (1); groove bridged by an ossified supratendinal bridge (2). (Ordered)

215. Tibia/tarsal-formed condyles: medial condyle projecting farther cranially than lateral condyle (0); equal in cranial projection (1).

216. Tibia/tarsal-formed condyles, mediolateral widths: medial condyle wider (0); approximately equal (1); lateral condyle wider (2). (Ordered).

217. Tibia/tarsal-formed condyles: gradual sloping of condyles towards midline of tibiotarsus (0); no tapering of either condyle (1).

218. Proximal end of the fibula: prominently excavated by a medial fossa (0); nearly flat (1).

219. Fibula, tubercle for m. iliofibularis: craniolaterally directed (0); laterally directed (1); caudolaterally or caudally directed (2). (Ordered)

220. Fibula, distal end reaching the proximal tarsals: present (0); absent (1).

221. Distal tarsals in adults: free (0); completely fused to the metatarsals (1). Any specimen that is inferred to be a juvenile should be scored as a "?" in order to account for the possibility of ontogenetic change.

222. Metatarsals II-IV, intermetatarsal fusion: absent or minimal co-ossification (0); partial fusion, sutural contacts easily discernible (1); completely or nearly completely fused, sutural contacts absent or poorly demarcated (2). (Ordered)

223. Proximal end of metatarsus: plane of articular surface perpendicular to longitudinal axis of metatarsus (0); strongly inclined dorsally (1).

224. Metatarsal V: present (0); absent (1).

225. Proximal end of metatarsal III: in the same plane as metatarsals II and IV (0); plantarly displaced with respect to metatarsals II and IV (1).

226. Tarsometatarsal proximal vascular foramen/foramina: absent (0); one between metatarsals III and IV (1); two (2).

227. Metatarsals, relative mediolateral width: metatarsal IV approximately the same width as metatarsals II and III (0); metatarsal IV narrower than metatarsals II and III (1); metatarsal IV greater in width than either metatarsal II or III (2).

228. Well-developed tarsometatarsal intercotylar eminence: absent (0); present, low and rounded (1); present, high and peaked (2).

229. Tarsometatarsus, projected surface and/or grooves on proximocaudal surface (associated with the passage of tendons of the pes flexors in Neornithes; hypotarsus): absent (0); developed as caudal projection with flat caudal surface (1); projection, with distinct crests and grooves (2); at least one groove enclosed by bone caudally (3). (Ordered)

230. Plantar surface of tarsometatarsus excavated: absent (0); present (1).

231. Tarsometatarsal distal vascular foramen completely enclosed by metatarsals III and IV: absent (0); present (1).

232. Metatarsal I: straight (0); J-shaped, the articulation of the hallux is located on the same plane as the attachment surface of the metatarsal I (1); J-shaped; the articulation of the hallux is perpendicular to the attachment surface (2); the distal half of the metatarsal I is laterally deflected so that the laterodistal surface is concave (3).

233. Metatarsal II tubercle (associated with the insertion of the tendon of the m. tibialis cranialis in Neornithes): absent (0); present, on approximately the center of the

proximodorsal surface of metatarsal II (1); present, developed on lateral surface of metatarsal II, at contact with metatarsal III or on lateral edge of metatarsal III (2). (Ordered)

234. Metatarsal II, distal plantar surface, fossa for metatarsal I: absent (0); shallow notch (1); conspicuous ovoid fossa (2). (Ordered)

235. Relative position of metatarsal trochleae: trochlea III more distal than trochleae II and IV (0); trochlea III at same level as trochlea IV, both more distal than trochlea II (1); trochlea III at same level as trochleae II and IV (2); distal extent of trochlea III intermediate to trochlea IV and II where trochlea IV projects furthest distally (3).

236. Metatarsal II, distal extent of metatarsal II relative to metatarsal IV: approximately equal in distal extent (0); metatarsal II shorter than metatarsal IV but reaching distally farther than base of metatarsal IV trochlea (1); metatarsal II shorter than metatarsal IV, reaching distally only as far as base of metatarsal IV trochlea (2).

237. Distal tarsometatarsus, trochlea in distal view: aligned in a single plane (0); metatarsal II slightly displaced plantarly with respect to III and IV (1); metatarsal II strongly displaced plantarly in respect to III and IV, such that there is little or no overlap in medial view (2)

238. Trochlea of metatarsal II broader than the trochlea of metatarsal III: absent (0); present (1).

239. Metatarsal III, trochlea in plantar view, proximal extent of lateral and medial edges of trochlea: trochlear edges approximately equal in proximal extent (0); medial edge extends farther (1).

240. Distal end of metatarsal II strongly curved medially: absent (0); present (1).

241. Digit IV phalanges in distal view, medial trochlear rim enlarged with respect to lateral trochlear rim: absent (0); present (1); greatly enlarged with the lateral trochlea reduced to a rounded peg (2).

242. Completely reversed hallux (arch of ungual phalanx of digit I opposing the arch of the unguals of digits II-IV): absent (0); present (1).

243. Size of claw of hallux relative to other pedal claws: shorter, weaker, and smaller (0); similar in size (1); longer, more robust, and larger (2).

244. Alula: absent (0); present (1).

245. Fan-shaped feathered tail composed of more than two elongate rectrices: absent (0); present (1).

246. Sternum, outermost trabecula: mainly parallel to the long axis of the sternum (0); clearly directed laterally (1).

247. Distal end of furcula relative to sternal margin of coracoid: proximal to or level with the sternal margin of the coracoid (0); well beyond the sternal end of the coracoid (1). When coracoid and furcula are not remained in natural position, then their proximodistal lengths are compared.

248. Scapula and coracoid: fused (0); unfused (1).

249. Scapula, acromion process length relative to the length of the humeral articular facet: less than half (0); nearly equivalent (1); longer but less than two times (2); more than two times longer (3); (Ordered)

250. Alular digit (I), phalanx 1: longer than the phalanx 1 of digit II (0); shorter than or equivalent to the phalanx 1 of digit II (1).

251. Coracoid, width of the sternal end relative to the length along the shaft: approximately half or greater (0); between half to 1/3 (1); less than 1/3 (2).

252. Coracoid, sternal margin: convex (0); nearly straight (1); concave (2);

253. Humerus, deltopectoral crest, distal end recedes abruptly with the humeral shaft: present (0); absent (1)

254. Tibia/tarsal-formed condyles, intercondylar groove: mediolaterally broad, approximately 1/3 with of anterior surface (0); less than 1/3 width of anterior surface (1).

255. Metatarsal IV, distal extension of the metatarsal IV relative to the metatarsal III: shorter and proximal to the proximal margin of the trochleae III (0); shorter but reaching distally further than the proximal margin of the trochleae III (1); approximately equal or surpassing the trochleae III (2)

256. Reduced claw in digit IV: absent (0); present (1).

257. The length ratio between tibiotarsus and tarsometatarsus: 2 or larger (0); between 2 and 1.6 (1); smaller than 1.6 (2). When distal tarsals are not fused with metatarsals, metatarsal III length is used.

258. Pedal digit, penultimate phalanx, longer than preceding phalanges in each digit: absent (0); present (1).

259. Proximal phalanx of hallux, the longest non-ungual phalanx: absent (0); present (1).

260. Phalanx in digit IV: not as follows (0), the second and the third phalanges reduced and significantly shorter than the fourth phalanx (1), as before but with the proximal phalanx reduced to be nearly equal in length with the second and third phalanx (2).

261. Pedal digit III claw, length relative to the tarsometatarsus: less than 20% the length of tarsometatarsus (0); 20% – 40% (1); extremely elongated and measuring more than 40% the length of tarsometatarsus (2); When metatarsals are not fused with distal tarsals, the length of metatarsal III is used. The length of ungual represents the linear distance between proximal end (position equivalent to flexor process) and tip of the sheath. If the sheath is not preserved or disarticulated with bony ungual, it should be scored as "?". (Ordered)

262. Post-dentary mandible: not as follows (0); dorsal margin concave and ventral margin convex (1); sigmoid (2).

263. Premaxilla, preorbital portion occupying 60% or more the skull length: absent (0); present (1).

264. Pedal digit II: not as follows (0); much robust than the other digits (1)

265. Alular digit (I): long, exceeding the distal end of the major metacarpal (0); subequal (1); short, not surpassing this metacarpal (2). (Ordered)

266. Tarsometatarsus, length compared to femur length: 0.6 or fewer (0); 0.8-0.6 (1); 1-0.8 (2);1 or greater (3).

267. Quadratojugal, shape: without horizontal process posterior to ascending process (reversed "L" shape) (0); with horizontal posterior process (i.e., inverted 'T' or 'Y' shape) (1).

268. Jugal and quadratojugal, fusion: absent (0); present, the two bones are not distinguishable from one another (1). (Ordered)

269. Caudal vertebrae, change in morphology of free caudals along the tail: present, with distinct transition point from shorter centra with long transverse processes proximally to

longer centra with small or no transverse processes distally (0); absent, vertebrae homogeneous in shape, without transition point (1).

270. Caudal vertebrae, location of transition point along the tail: begins distal to the 10th caudal vertebra (0); between the 7th and 10th caudal vertebra (1); proximal to the 7th caudal vertebra (2) (Ordered).

271. Coracoid and scapula, angle between bones at glenoid: greater than or equivalent to 90° (0); smaller than 90° (1).

272. Ischium, distal end: continuous with the proximal shaft, rendering the cranial. margin of the ischium straight or weakly convex (0); directed cranioventrally toward the pubis, rendering the cranial margin concave (1).

273. Ischium, caudal margin with a dorsal process: located proximal to or close to the midpoint of the caudal margin (0); distal to the midpoint of the caudal margin (1).

274. Sternum, cranial margin with a pair of craniolateral processes: absent (0); present (1).

275. Premaxilla corpus: dorsoventral height greater than or equal to craniocaudal length (0); dorsoventral height smaller than craniocaudal length (1).

276. Furcula, omal tip: blunt or expanded (0); tapered (1).

277. Alular metacarpal, cranial margin: expanded cranioproximally and craniodistally, and constricted just before flare of articulation with digit (0); broadly convex (1).

278. Alular digit, phalanx 1: straight (0); bowed (1).

279. Major digit, phalanx 2: straight (0); bowed (1).

280. Minor digit: with four phalanges (0); less than four phalanges (1).

**Characters added by Field et al. (2018):**

281. Premaxillae, extent along margin of jaw: 1⁄3 or less of total length of rostrum (0); between 1⁄3 and 1⁄2 of total length of rostrum (1); over 1⁄2 of total length of rostrum (2). (Ordered)

282. Premaxillae, ventral surface: deeply concave with exposed pair of neurovascular foramina (0); covered with a smooth palatal shelf (1).

283. Transverse ridges of bone (socketing) along the region of maxilla and dentary occupied by embryonic dental lamina (potentially dentigerous region): present (0); absent (1).

284. Anterior margin of upper temporal region: formed by extensive postorbital ossification with a primarily mediolateral, horizontal orientation, a flat upper table, and a posterior deflection of the tip (0); formed by reduced postorbital ossification directed ventrally but not laterally (1).

285. Posterior margin of upper temporal region: formed by extensive, ridged and buttressed squamosal process extending first laterally and then rostrally, with rostral portion elongate (longer than tall) and about the same length as the lateral extent (0); formed by reduced extension of squamosal enclosing a smaller adductor region and with a minor anterior extent (1).

#### SUPPLEMENTAL FIGURES

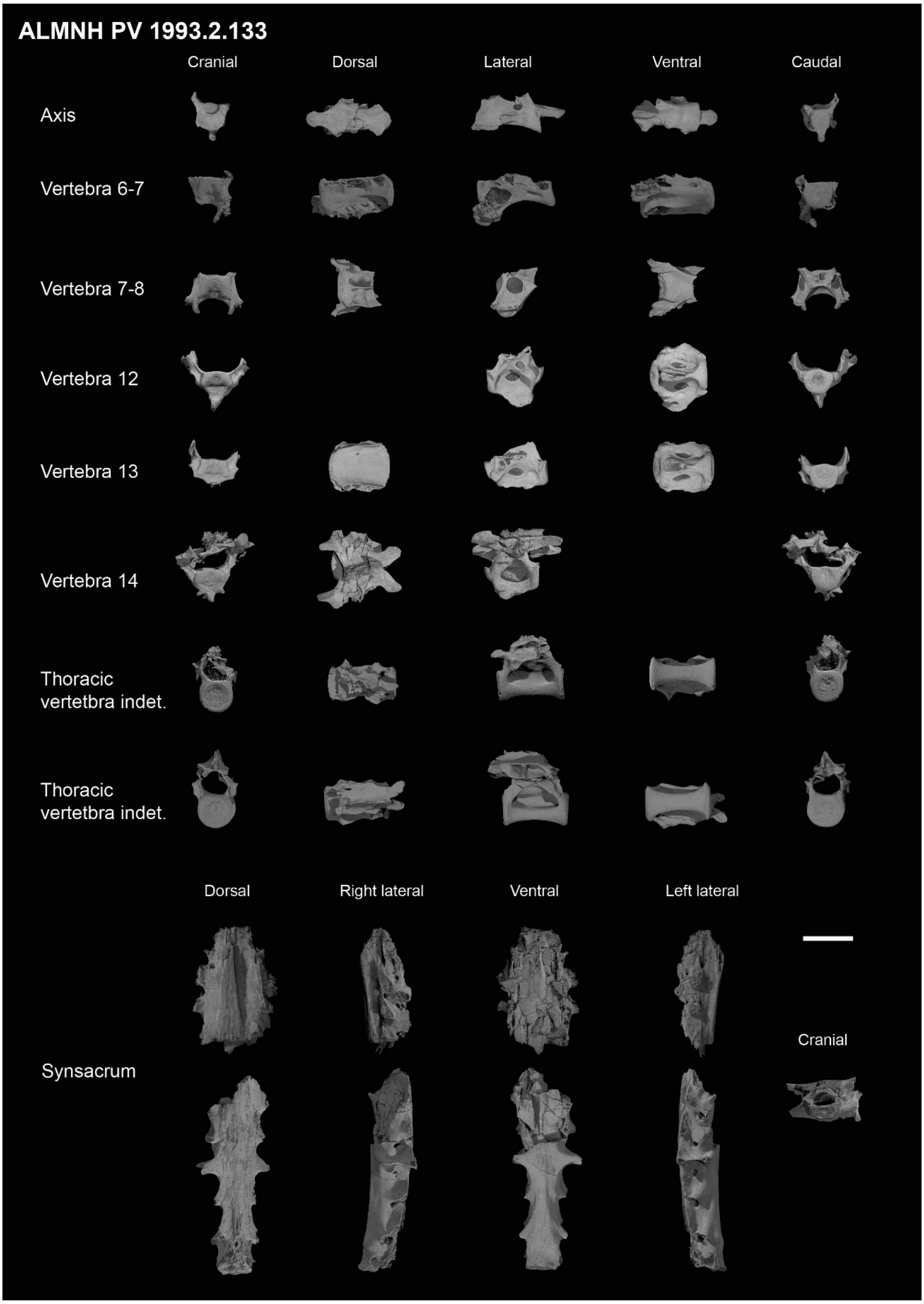


**Supplemental figure 1. Complete axial material from specimen ALMNH PV 1993.2.133.**

Scale bar equals 1 cm.

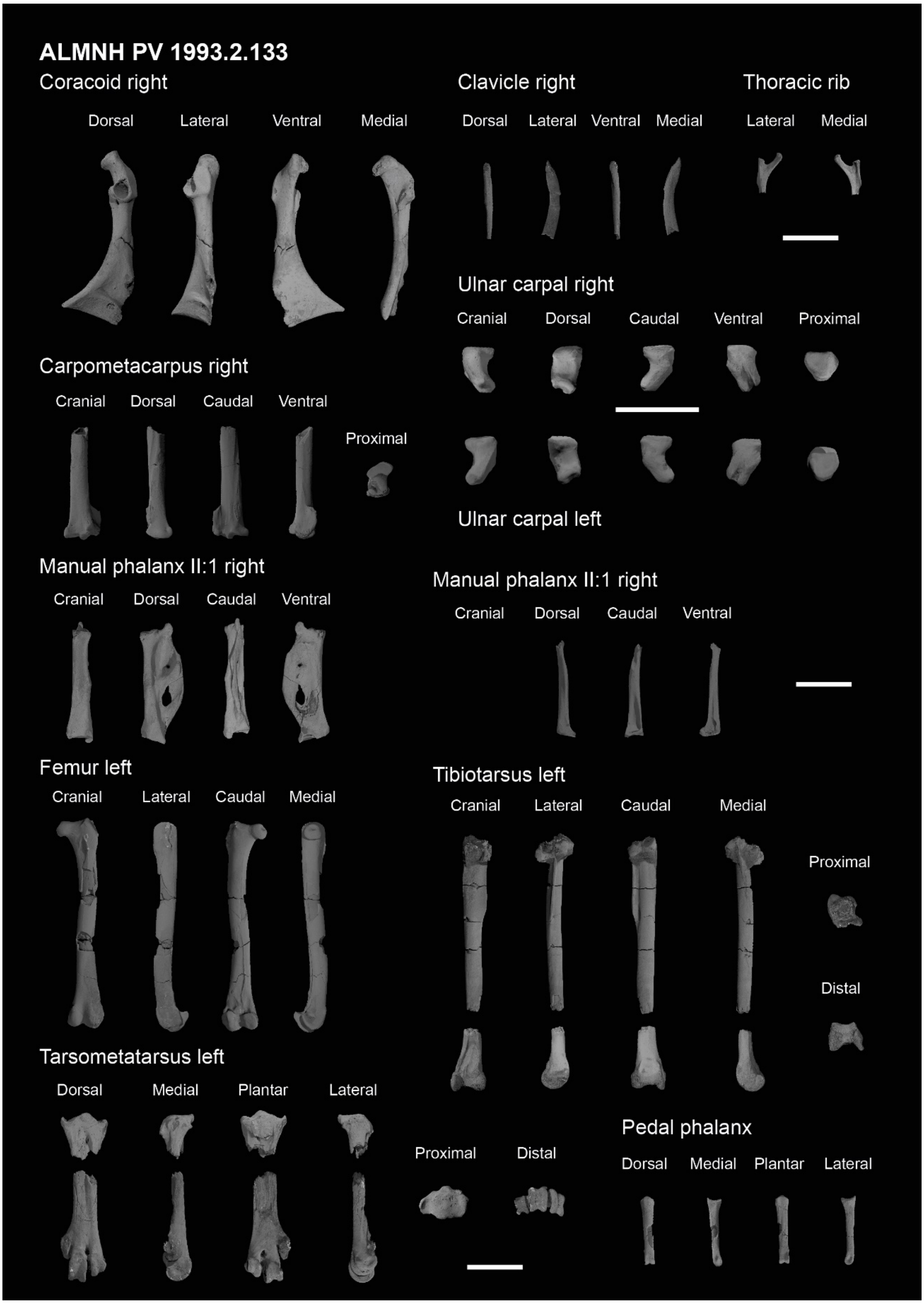


**Supplemental figure 2. Complete forelimb and hindlimb material from specimen ALMNH PV 1993.2.133.** Scale bars equals 1 cm.

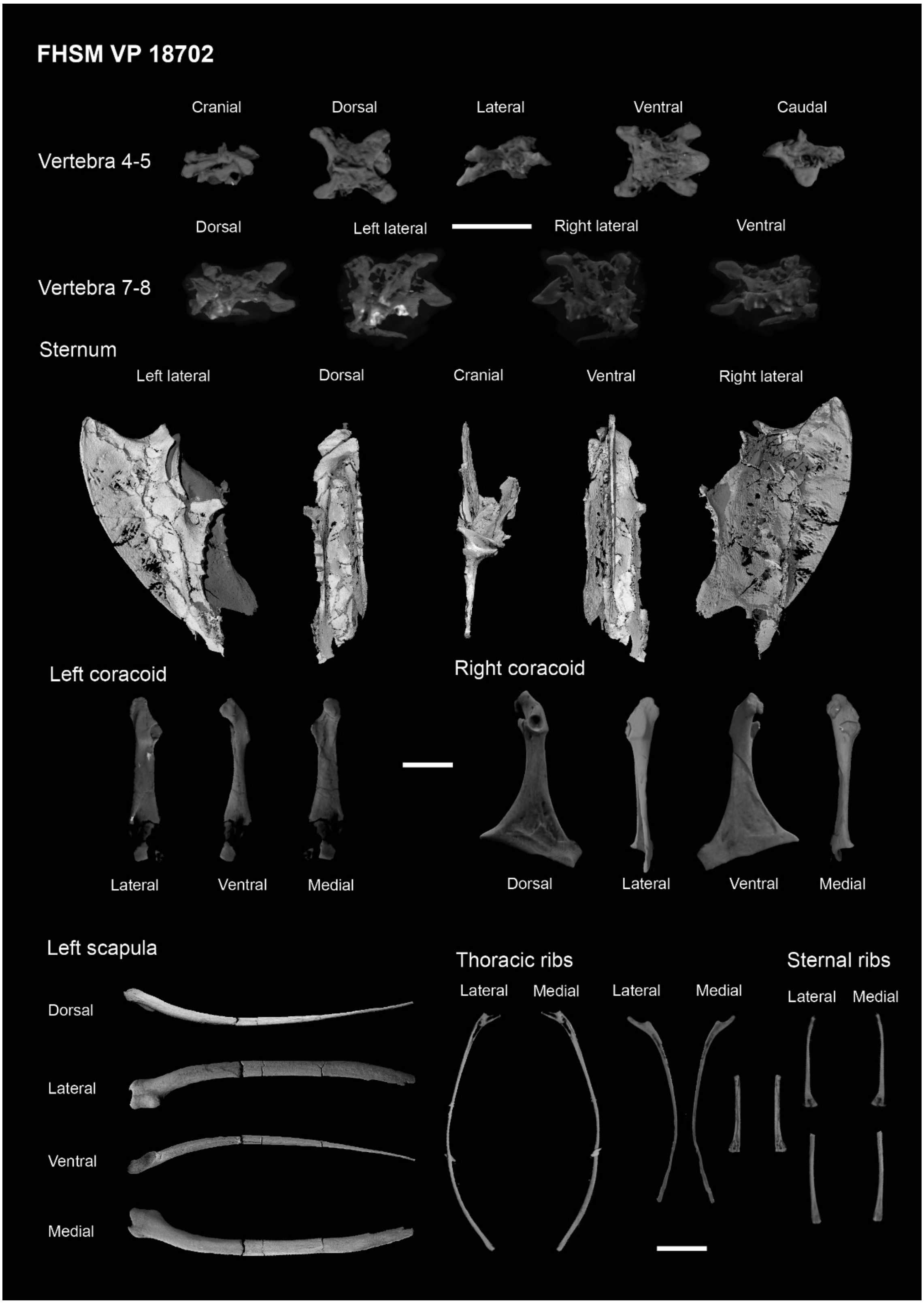


2273 **Supplemental figure 3. Complete cervical, pectoral and thoracic material from specimen**

2274 **FHSM VP 18702.** Scale bar equals 1 cm.

2275

2276

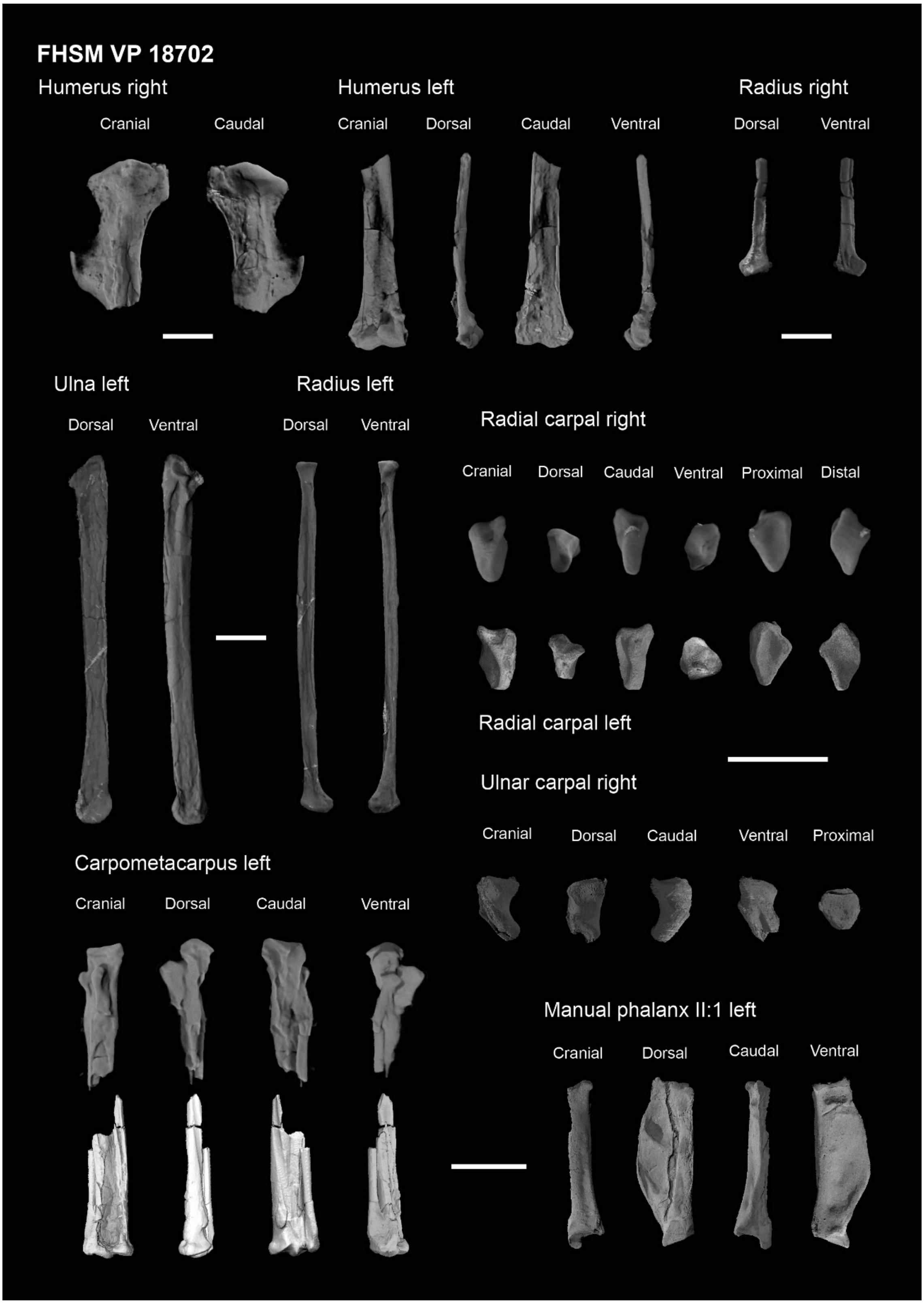


2278 **Supplemental figure 4. Complete forelimb material from specimen FHSM VP 18702.** Scale

2279 bars equals 1 cm.

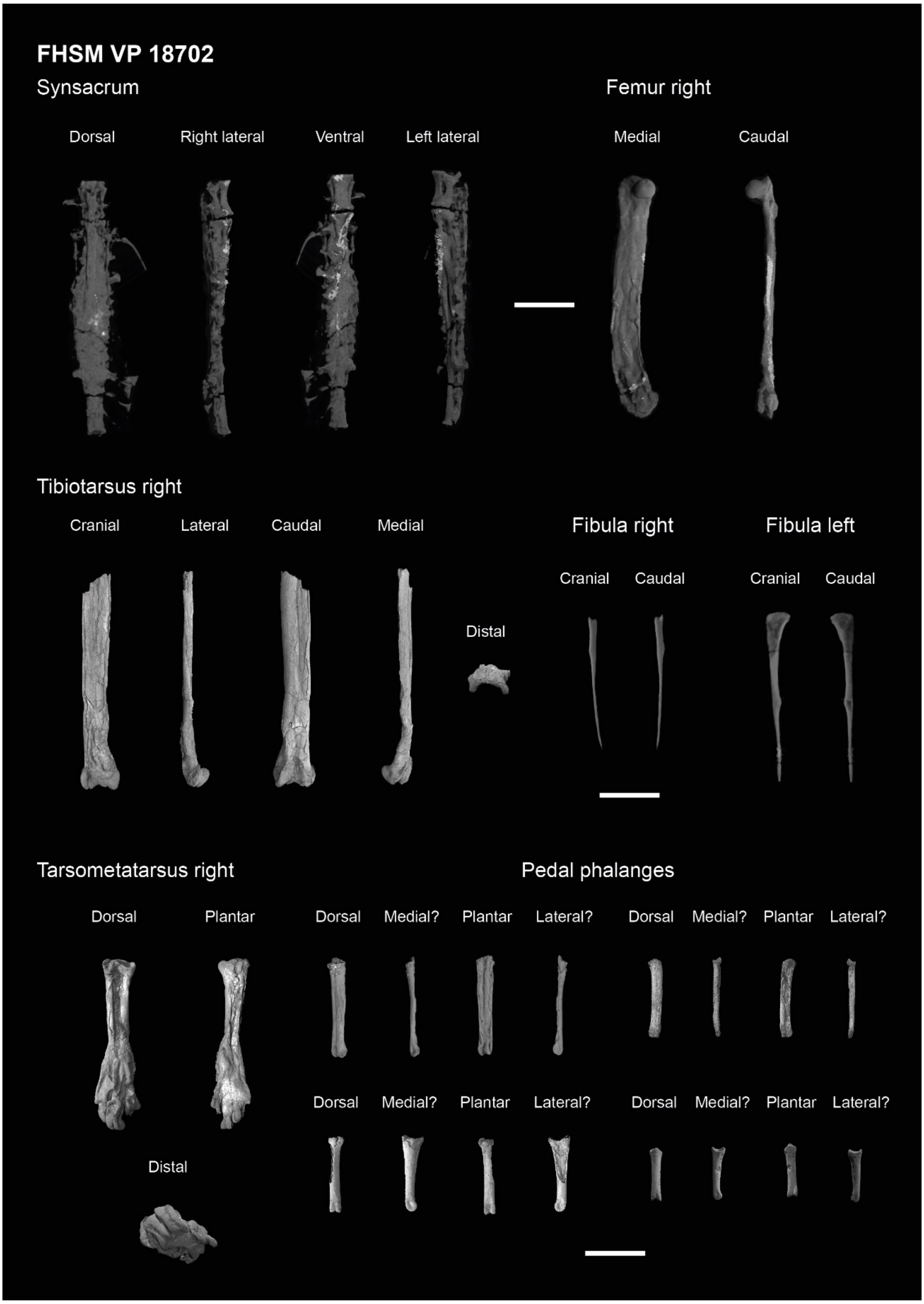


**Supplemental figure 5. Complete sacral and hindlimb material from specimen FHSM VP 18702.** Scale bars equals 1 cm.

2284

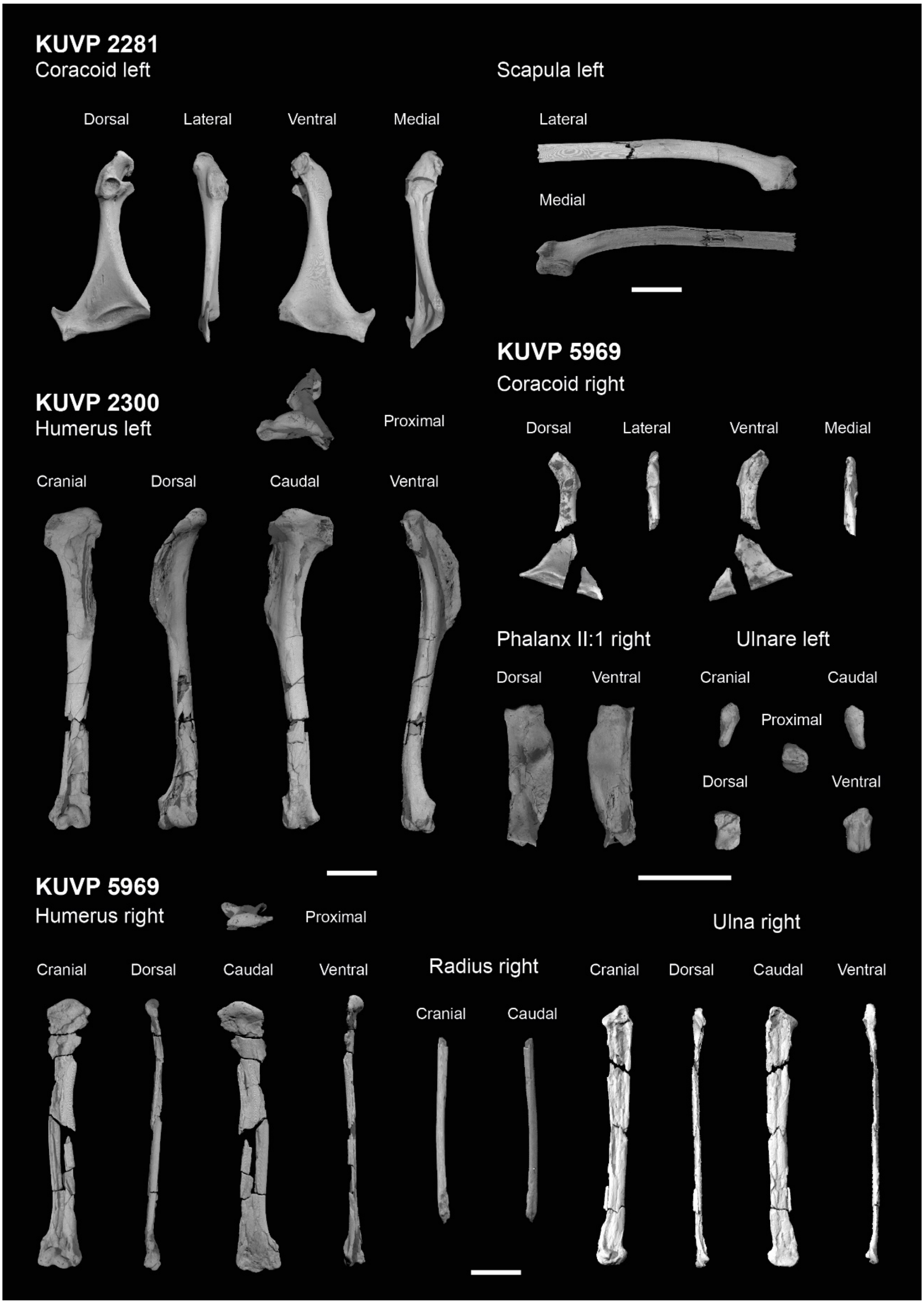


2286 **Supplemental figure 6. Complete pectoral and forelimb material from specimens KUVP**

2287 **2281, 2300 and 5969.** Scale bars equals 1 cm.

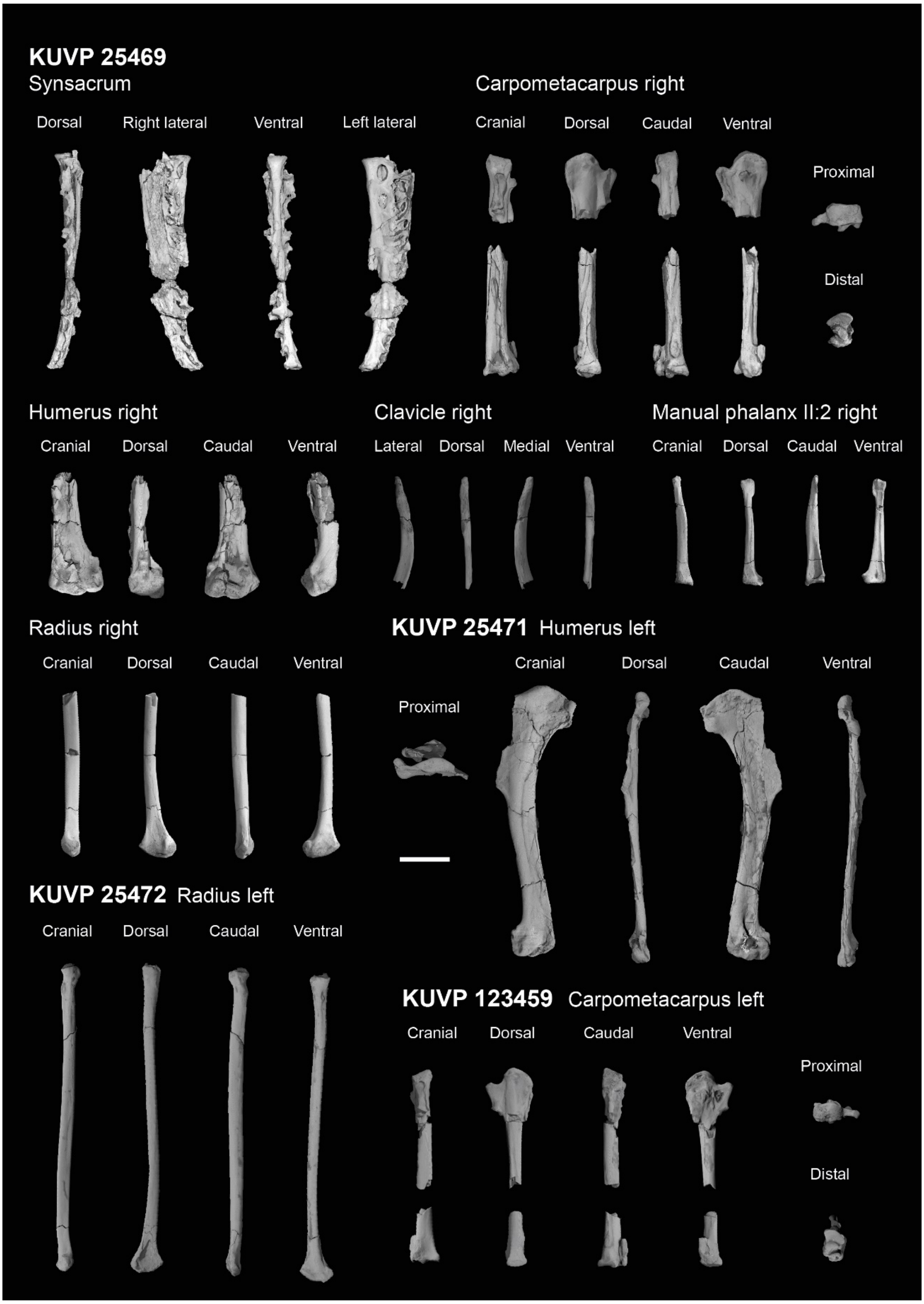


**Supplemental figure 7. Complete sacral, pectoral and forelimb material from specimens KUVP 25469, 25471, 25471 and 123459.** Scale bar equals 1 cm.

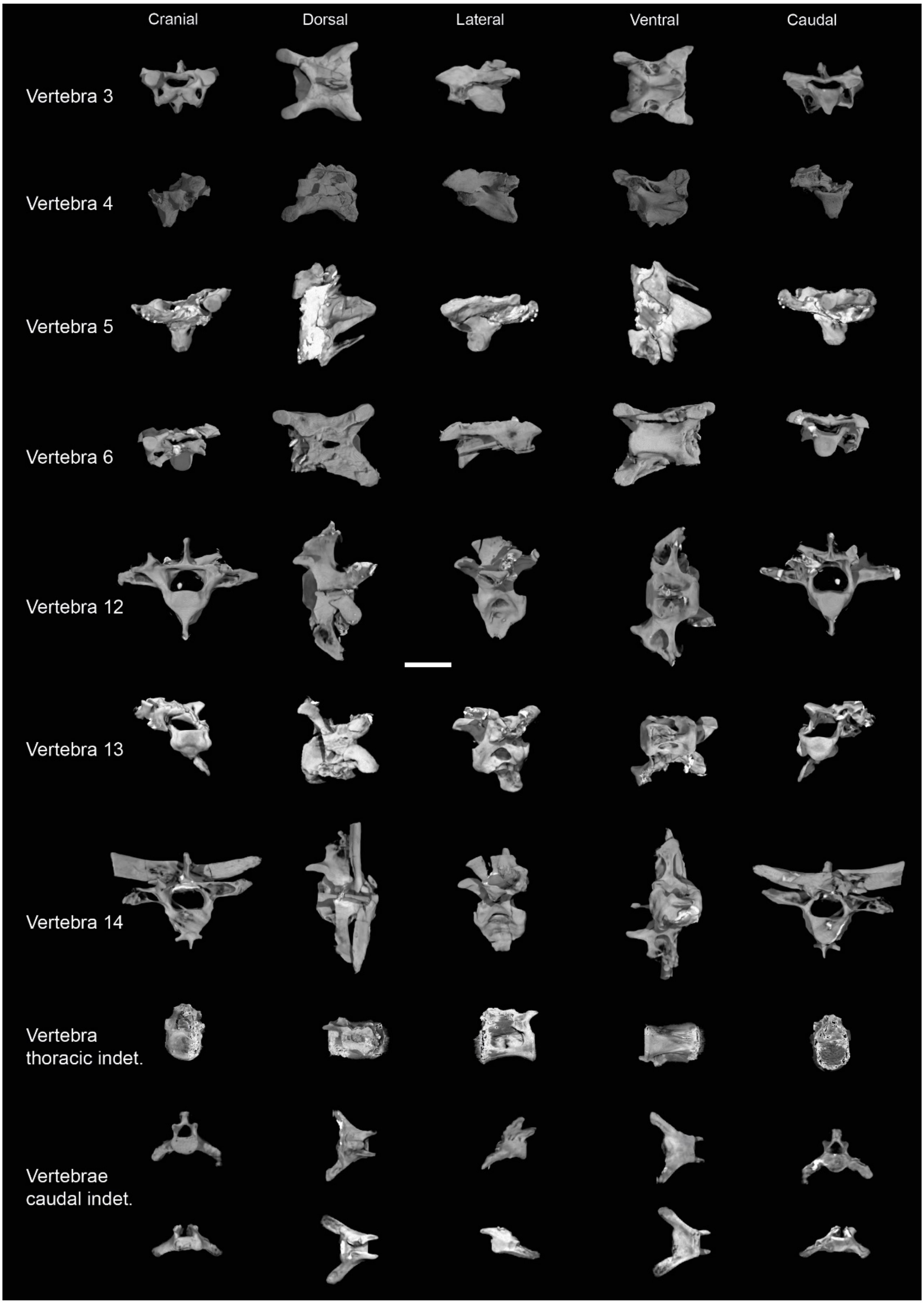


**Supplemental figure 8. Axial material from specimen KUVP 119673.** Scale bar equals 5 mm.

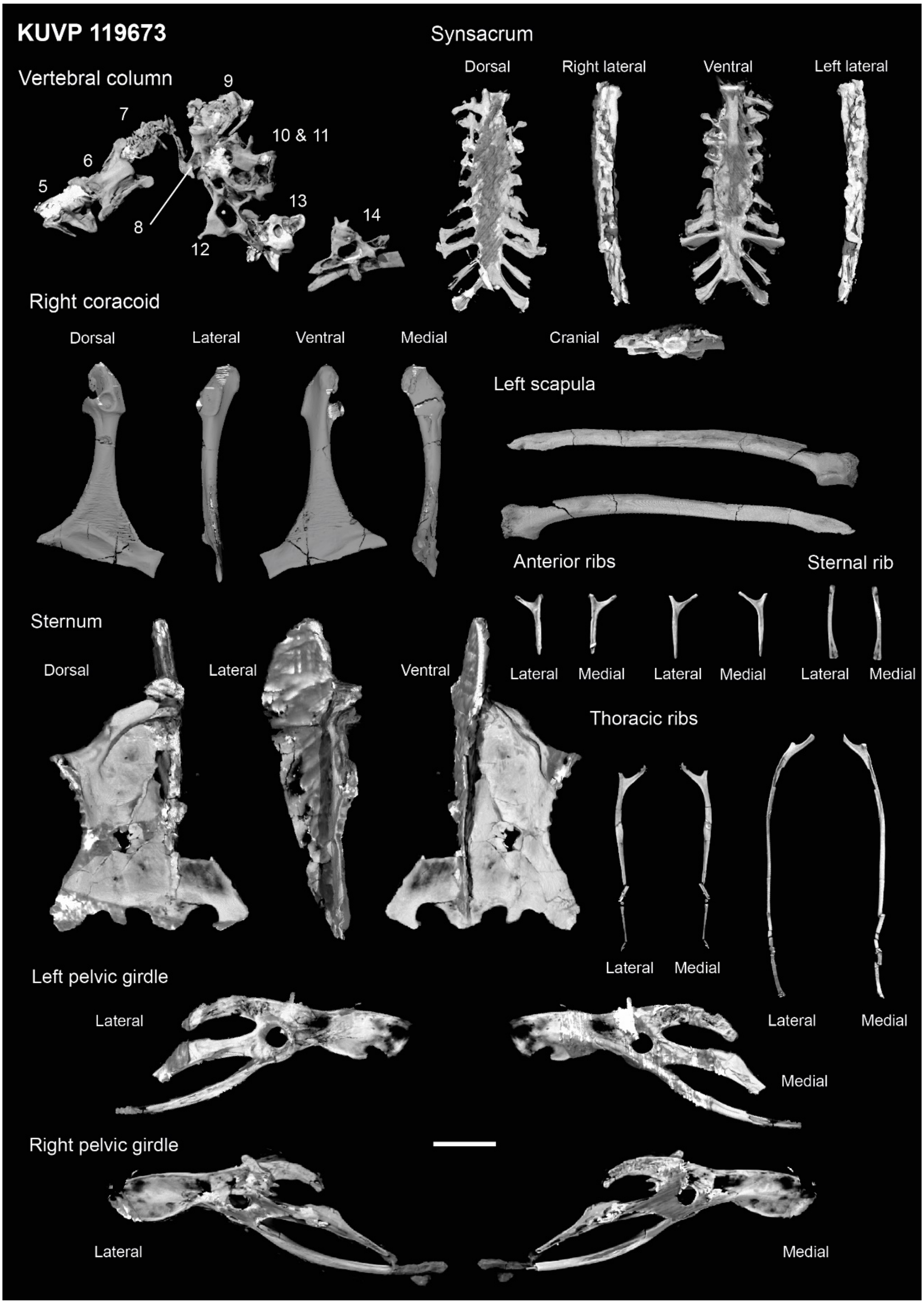


**Supplemental figure 9. Axial, pectoral, thoracic and pelvic material from specimen KUVP 119673.** Scale bar equals 1 cm.

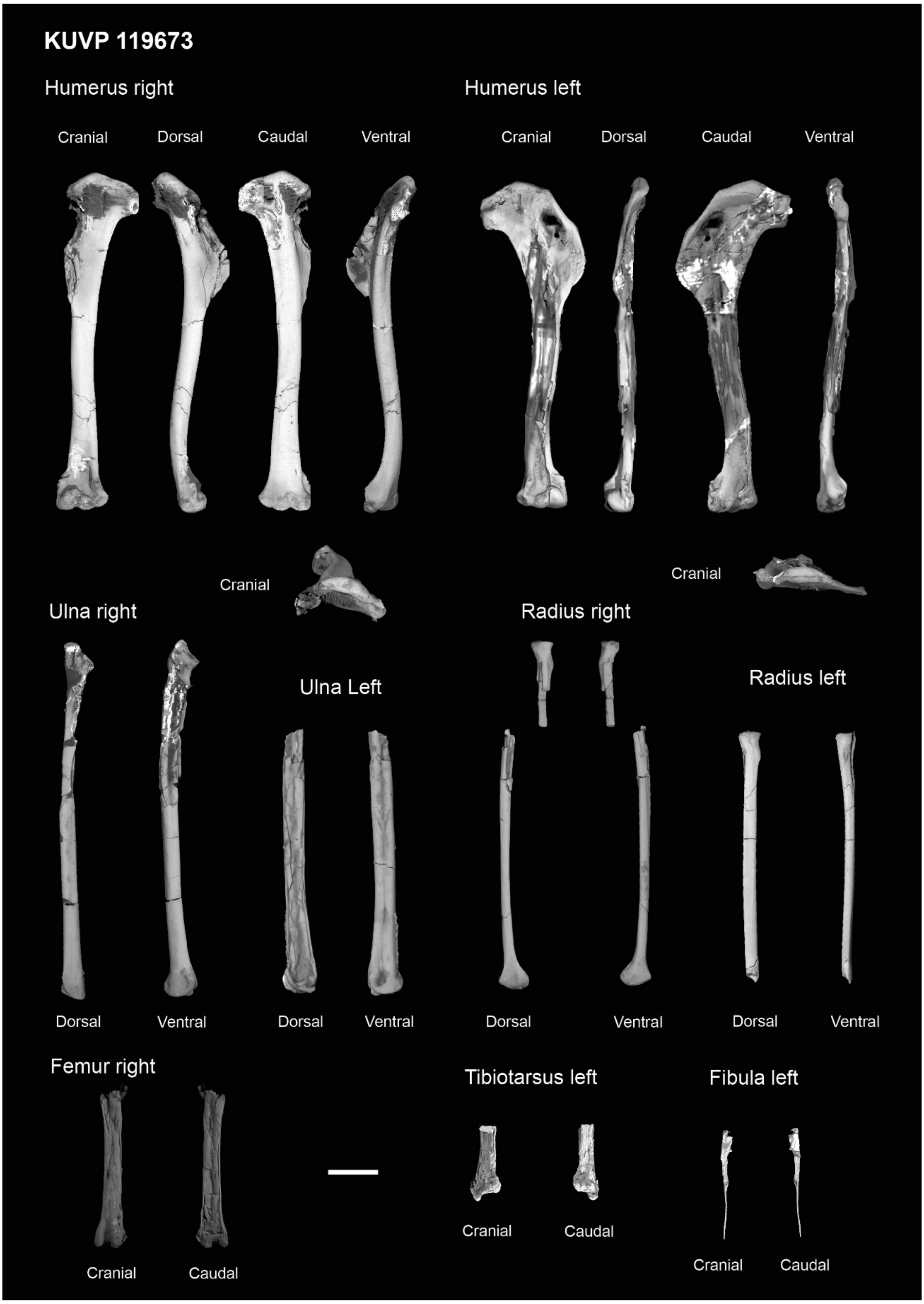


**Supplemental figure 10. Complete forelimb and hindlimb material from specimen KUVP 119673.** Scale bar equals 1 cm.

### SUPPLEMENTAL TREES

#### Supplemental Tree 1

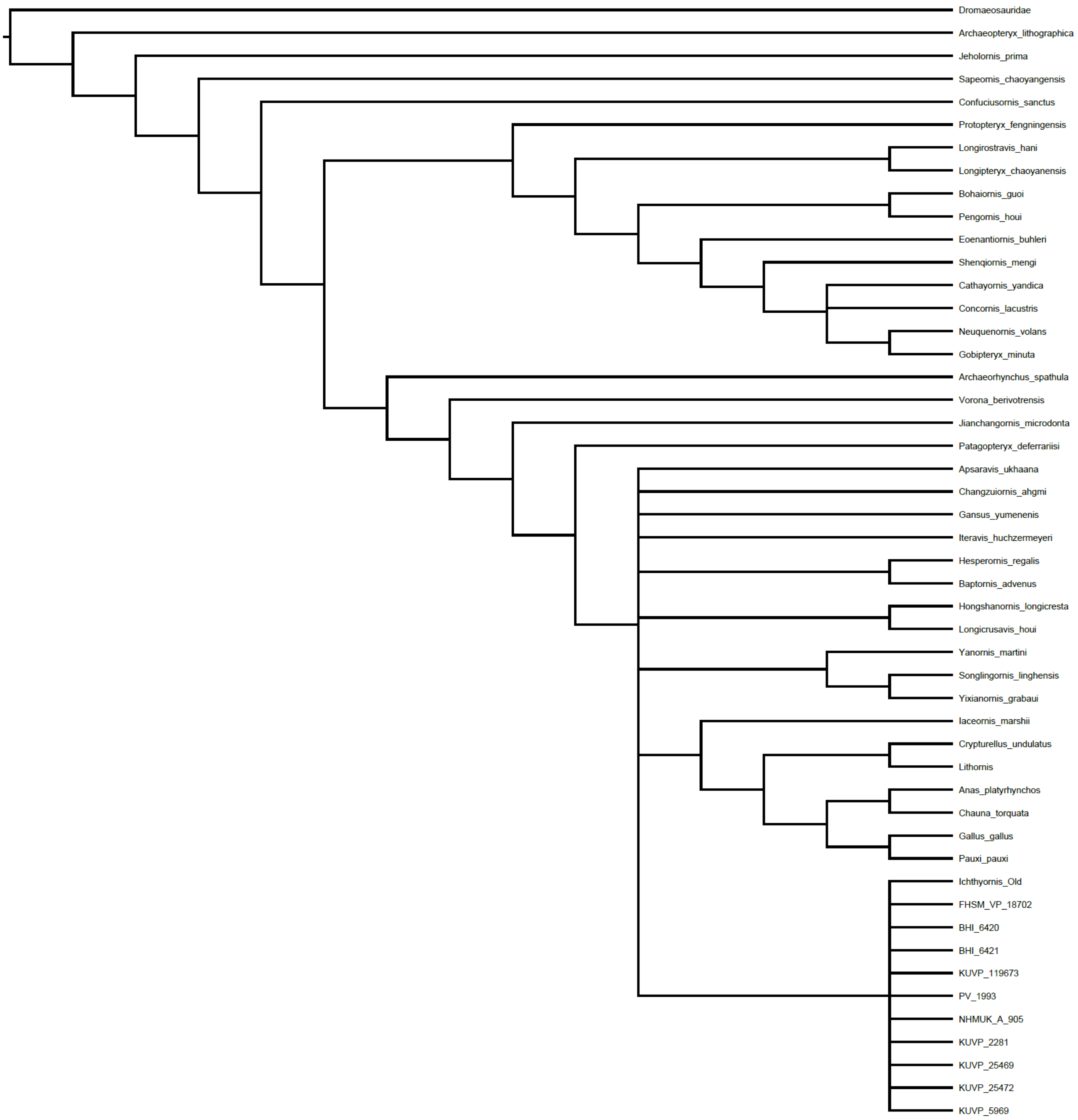

**Supplemental Tree 2**

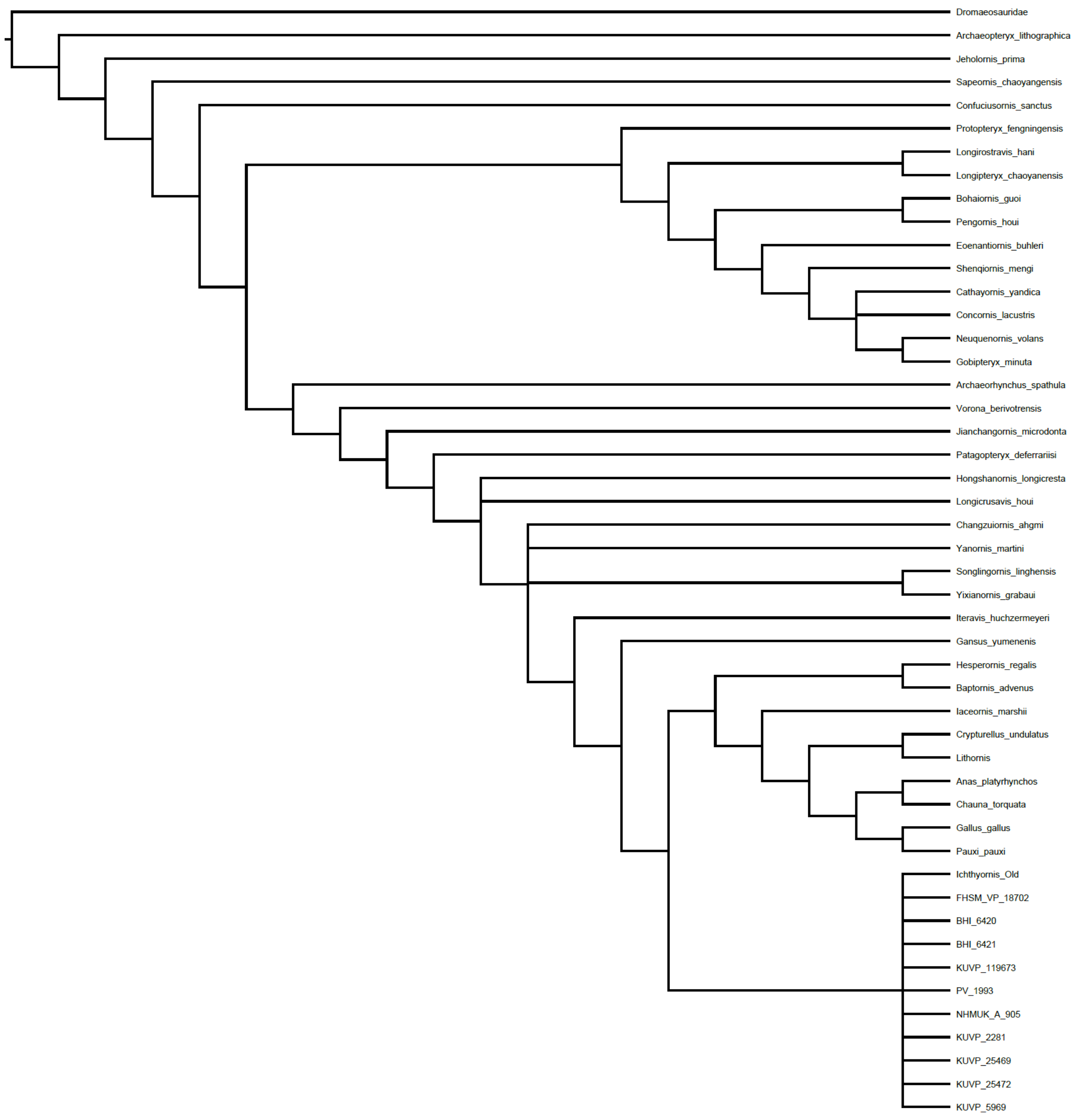

#### Supplemental Tree 3

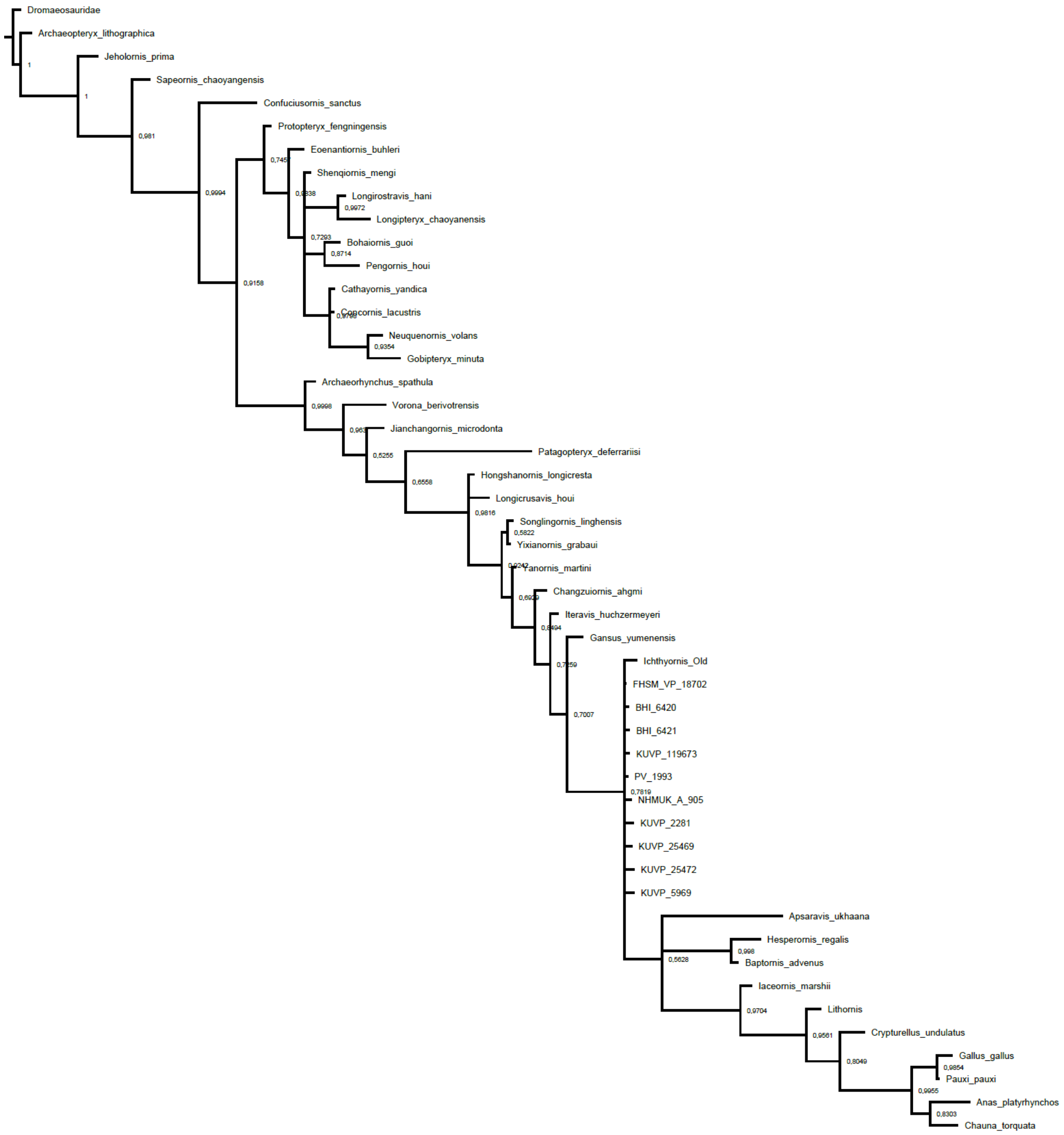

#### Supplemental Tree 4

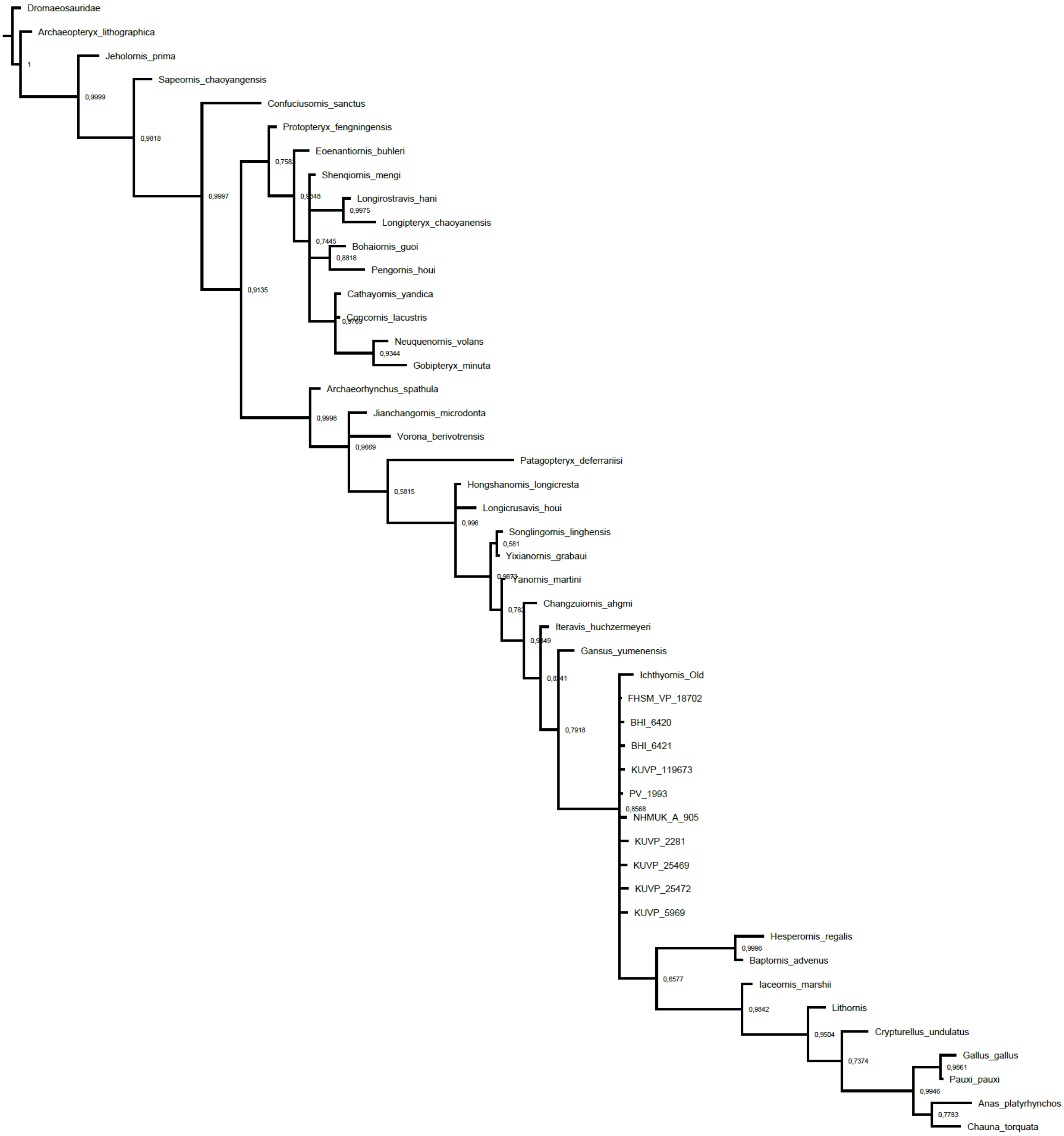

#### Supplemental Tree 5

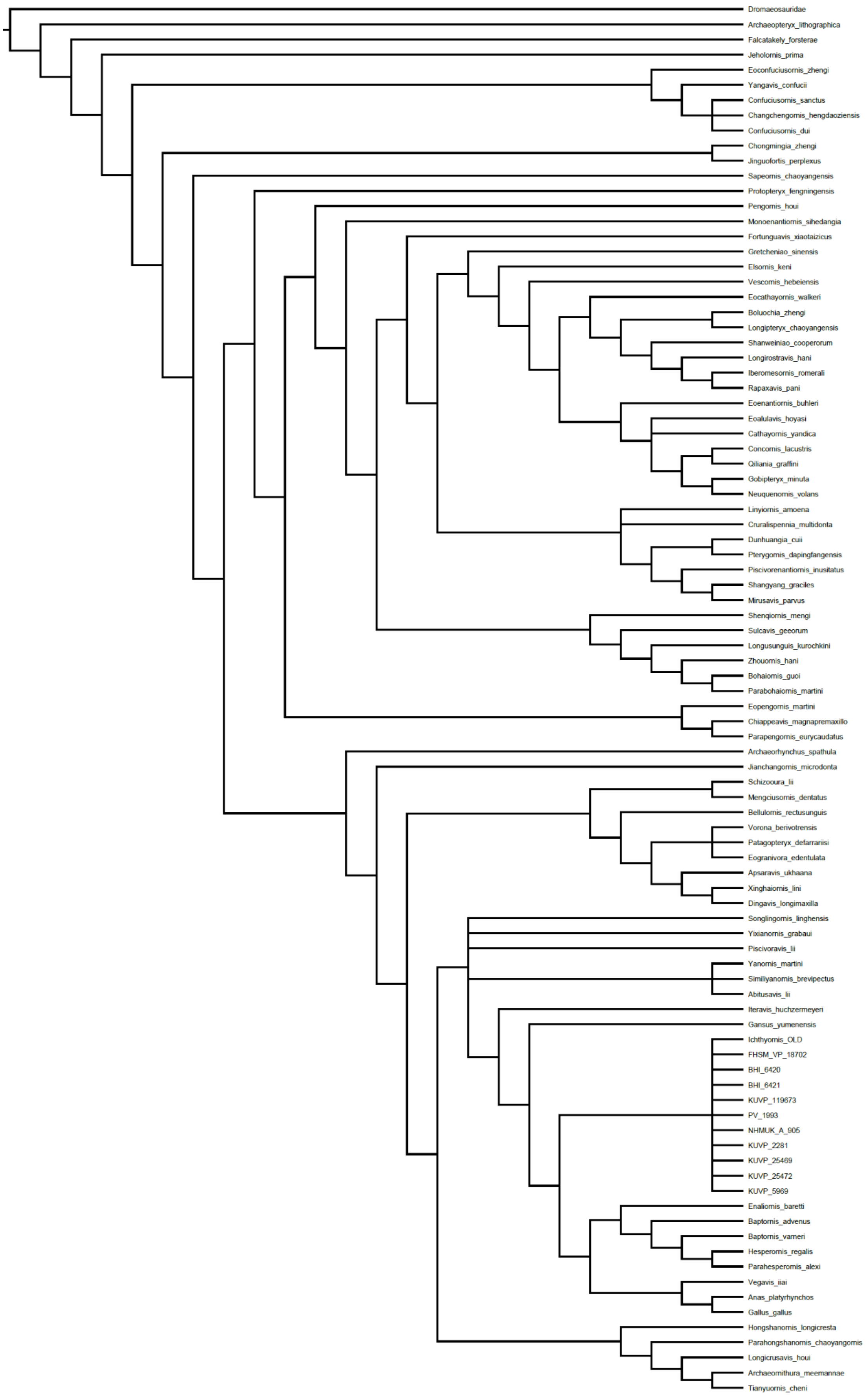

#### Supplemental Tree 6

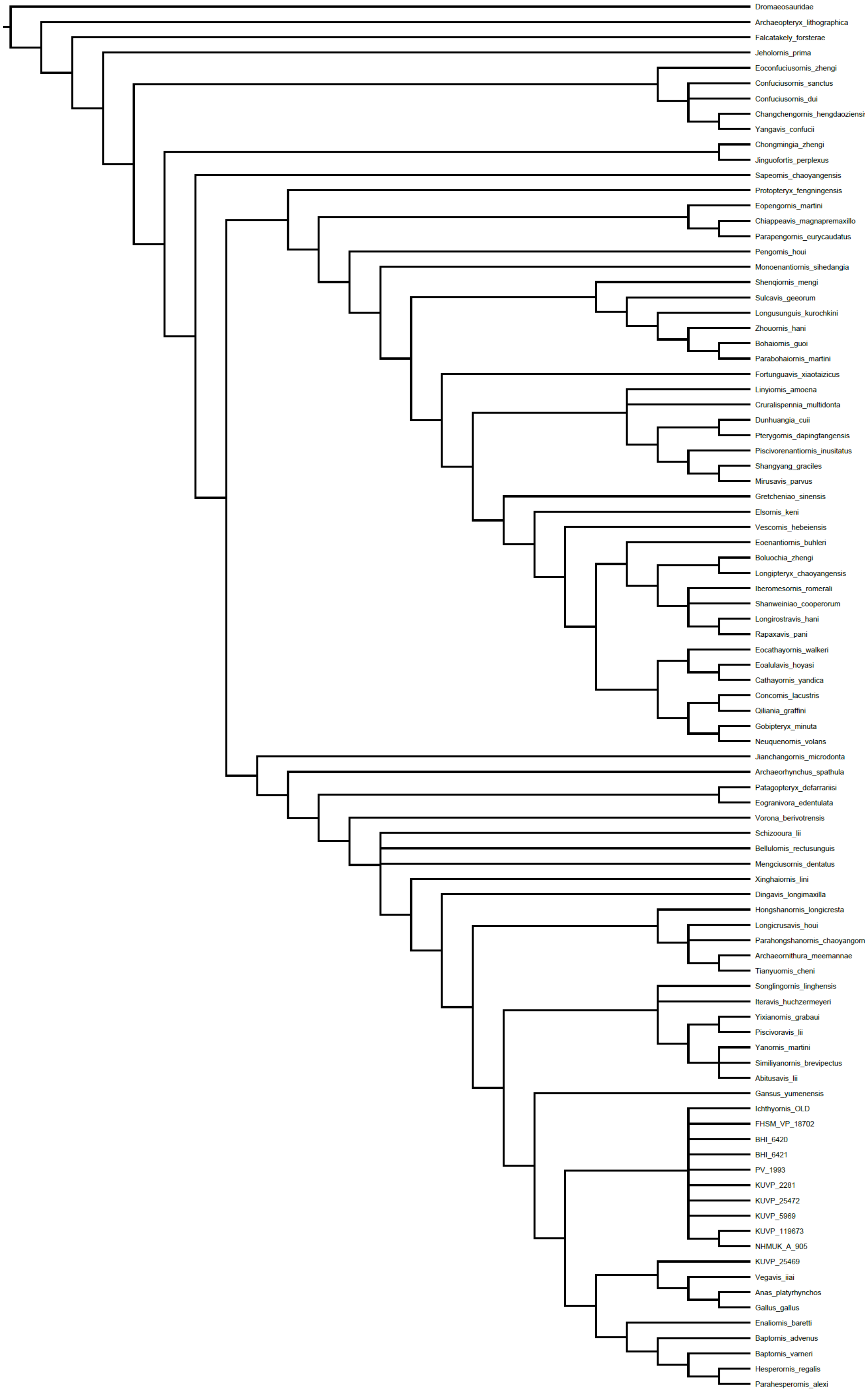

**Supplemental Tree 7**

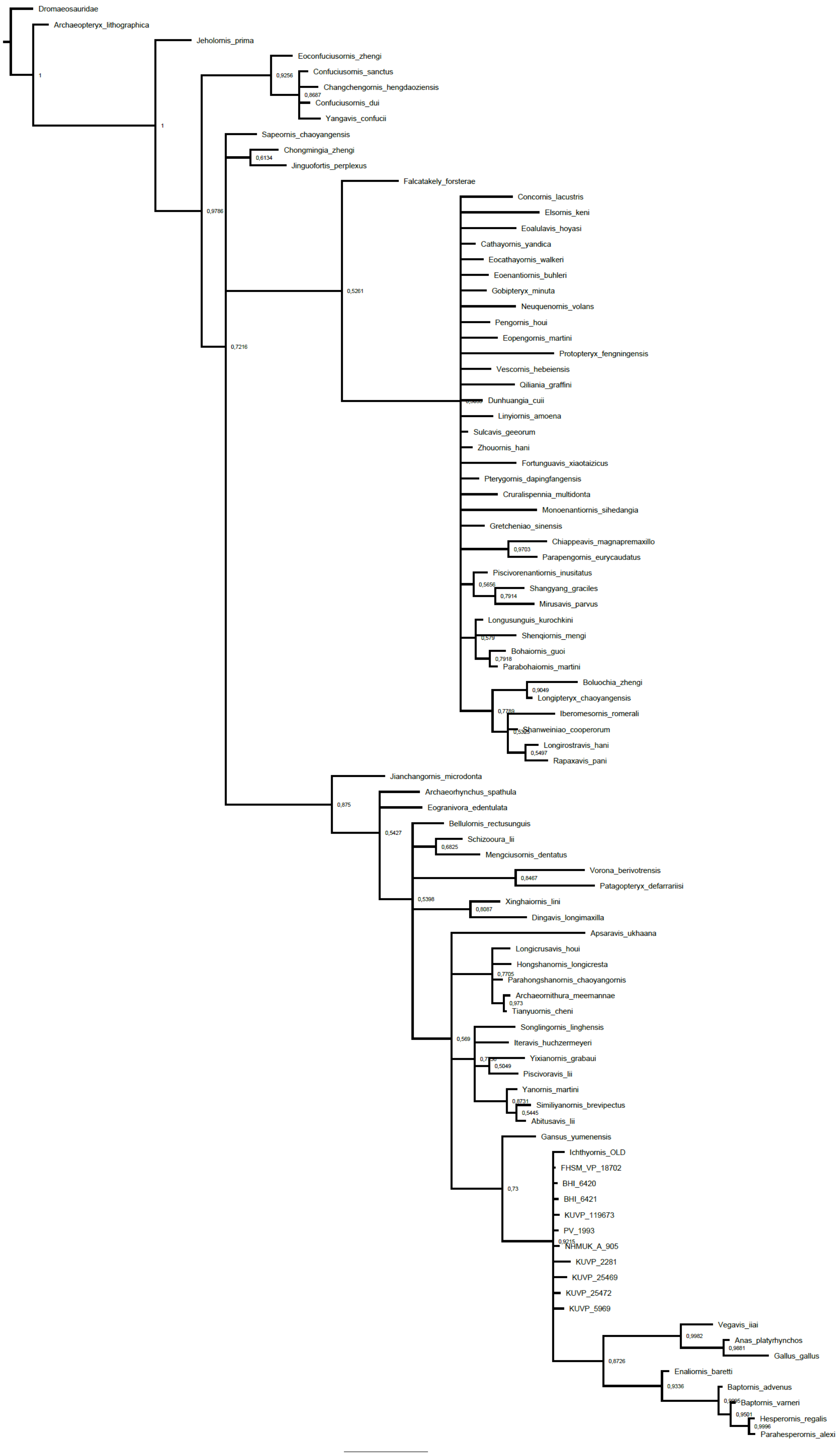

#### Supplemental Tree 8

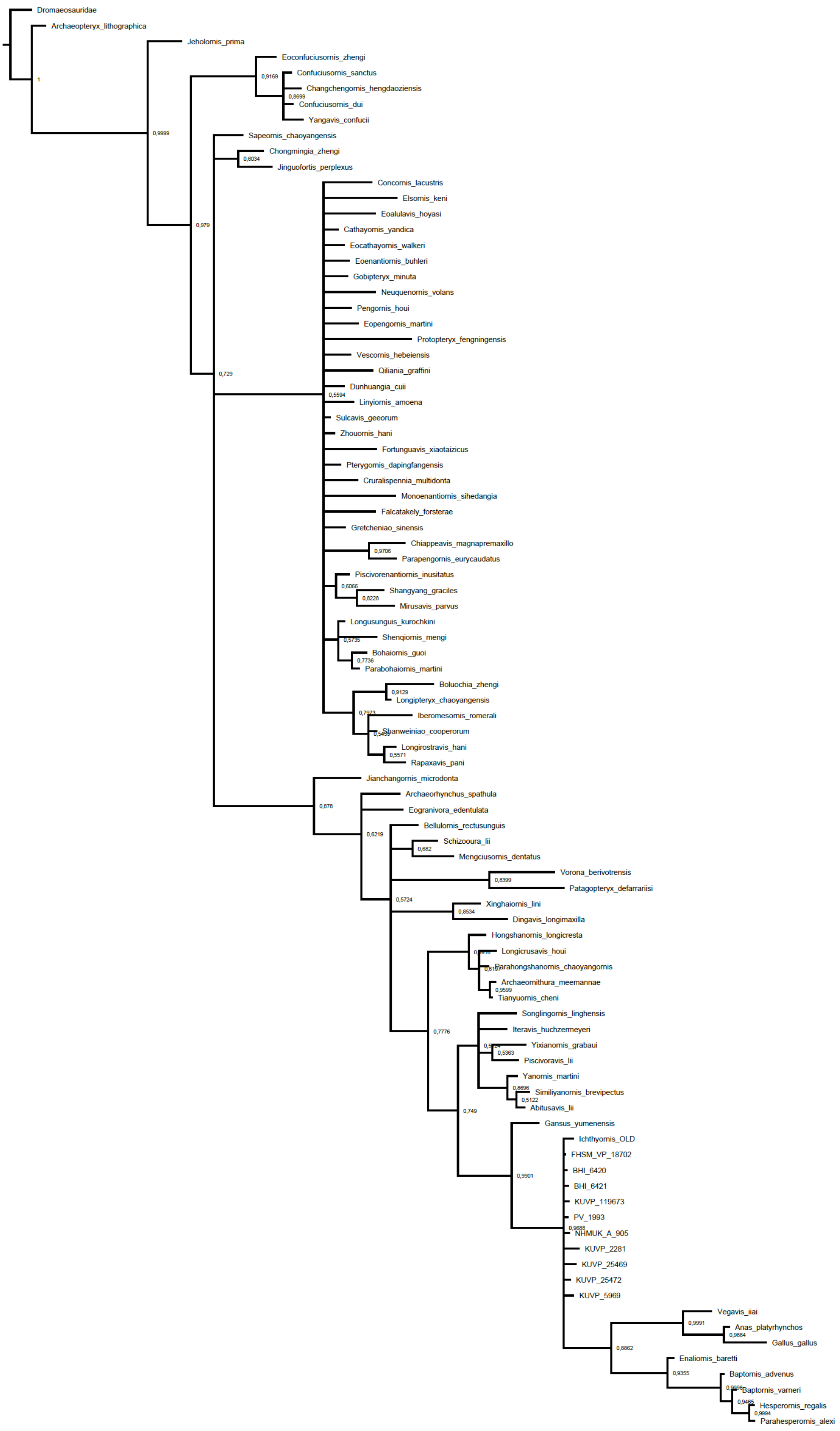

**Supplemental Tree 9**

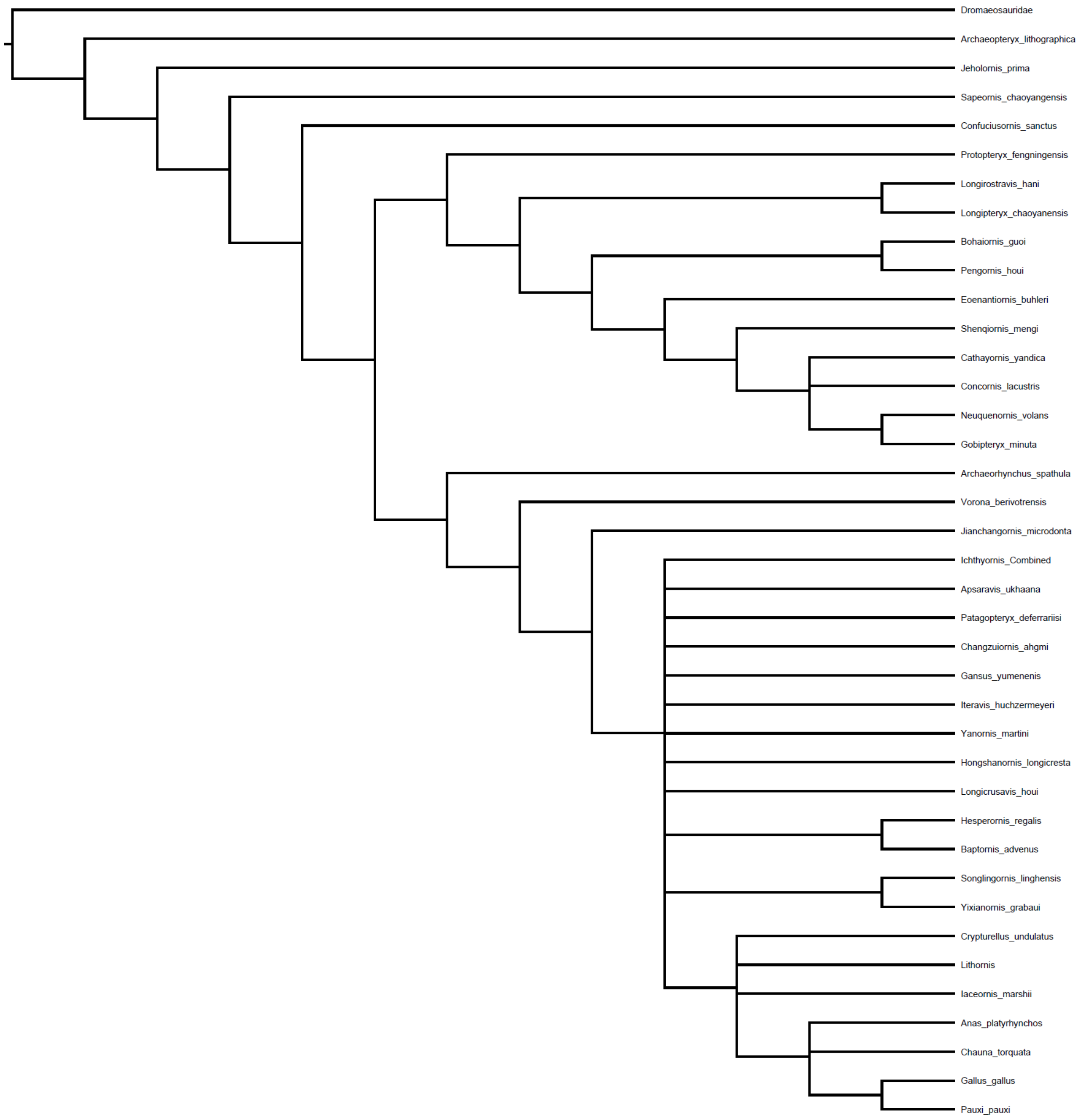

#### Supplemental Tree 10

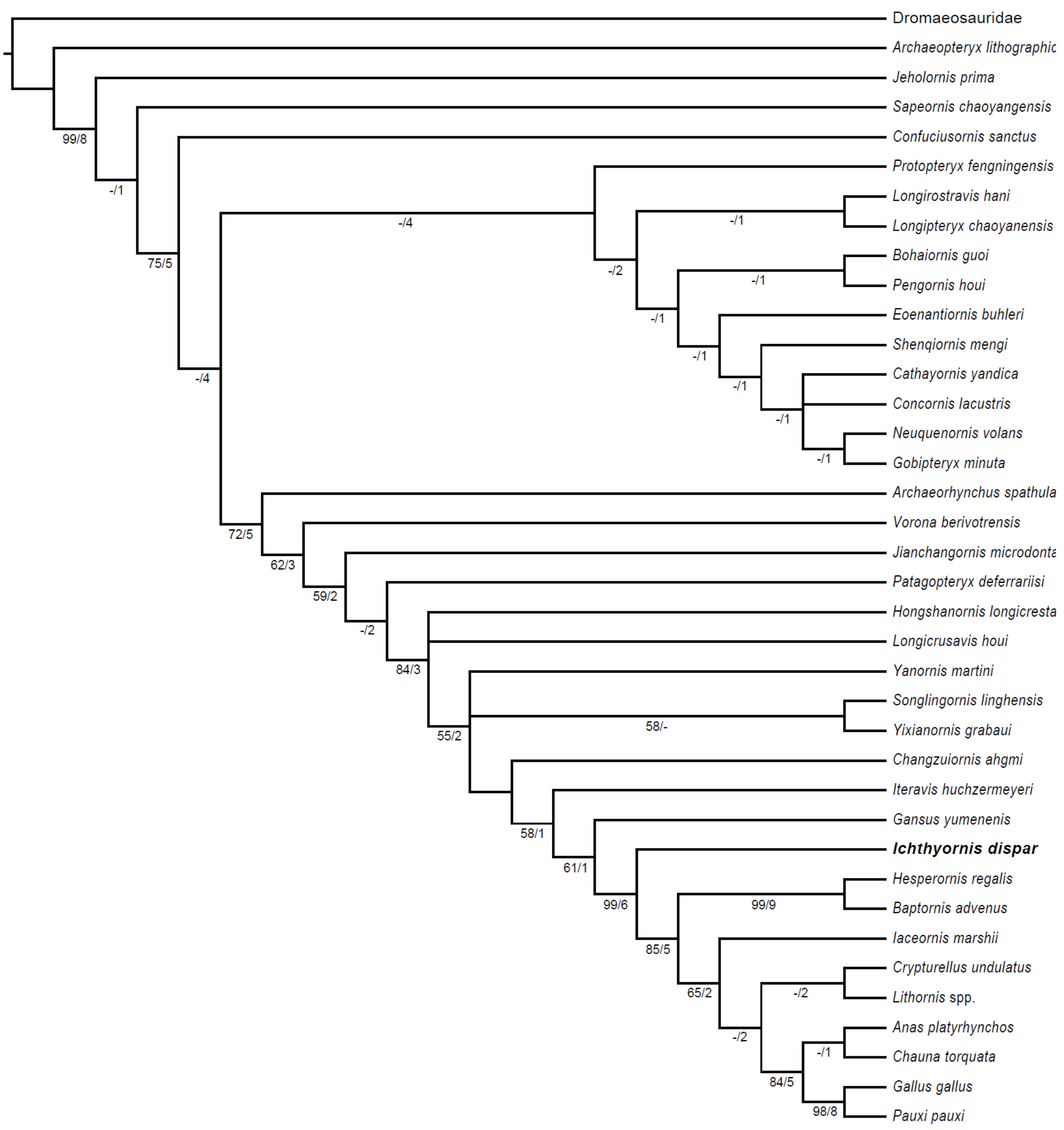

#### Supplemental Tree 11

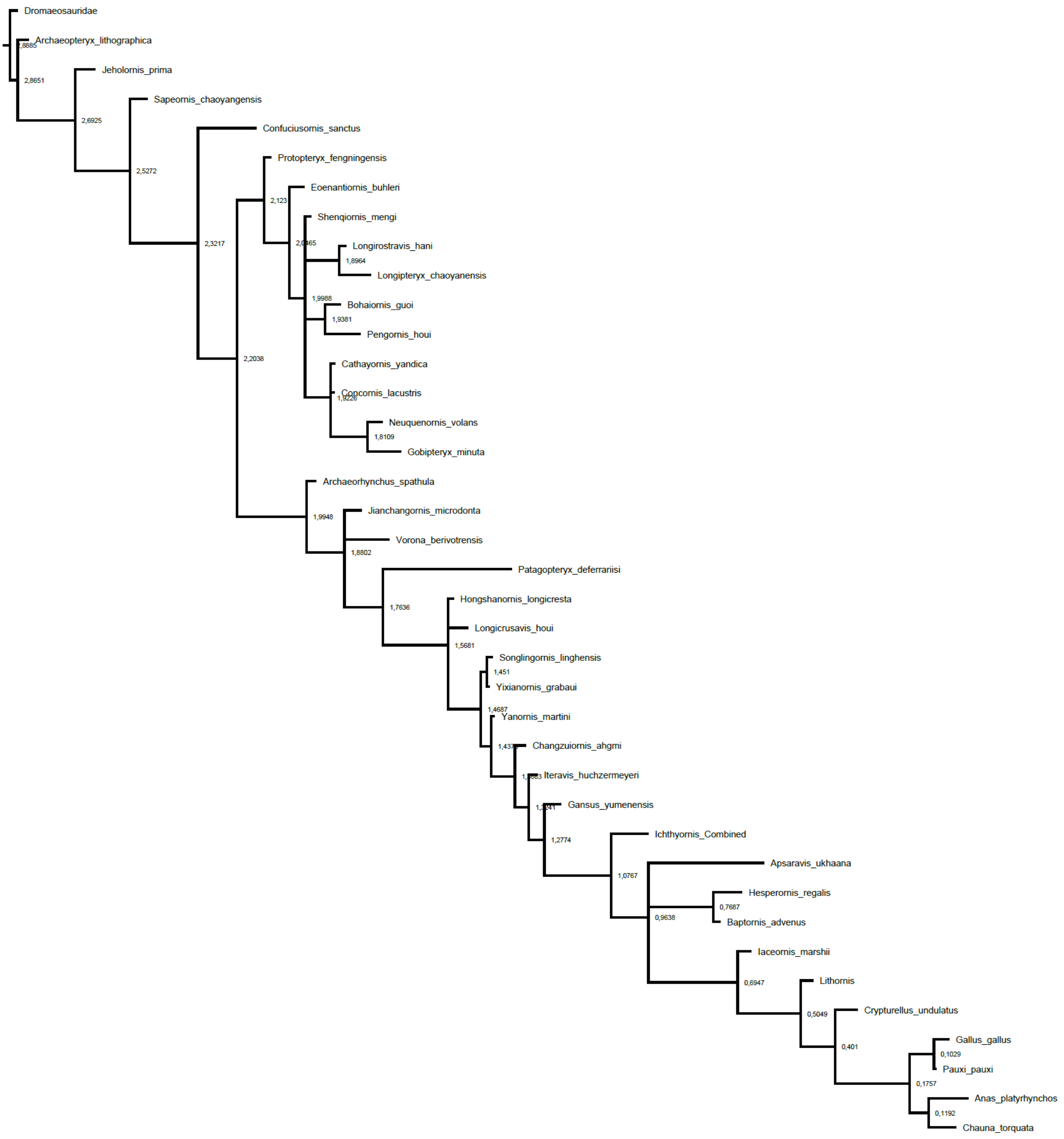

**Supplemental Tree 12**

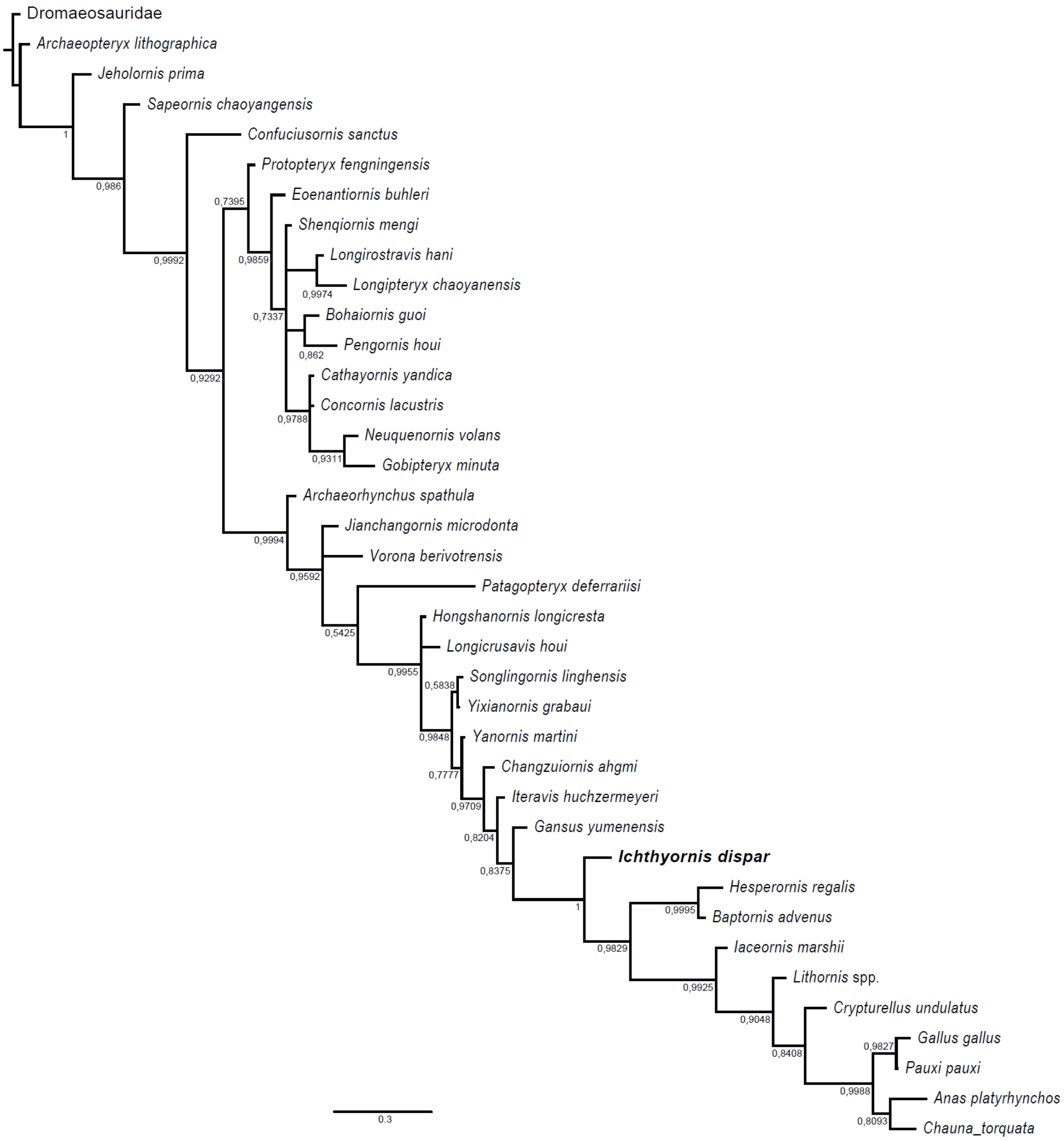

#### Supplemental Tree 14

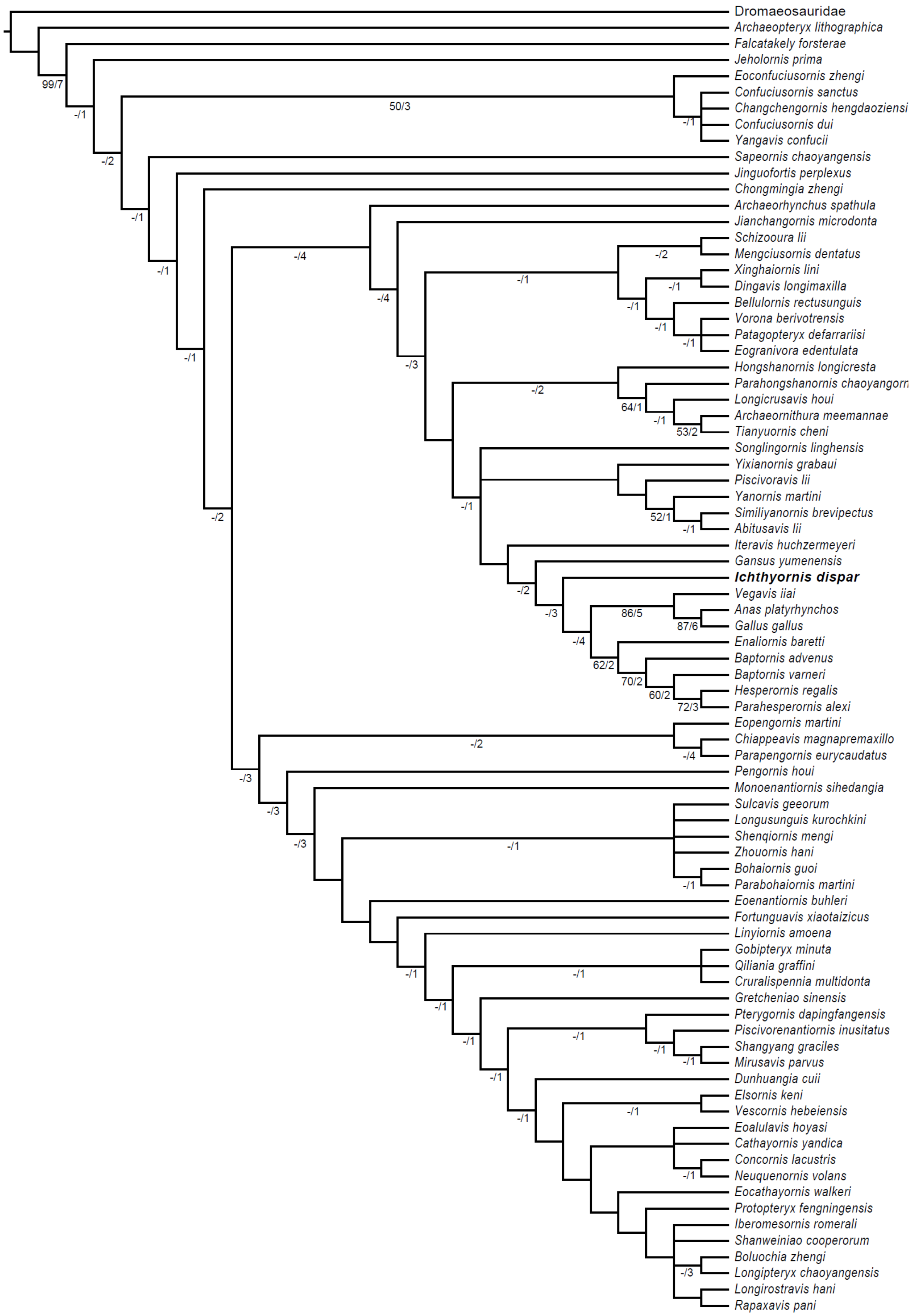

#### Supplemental Tree 15

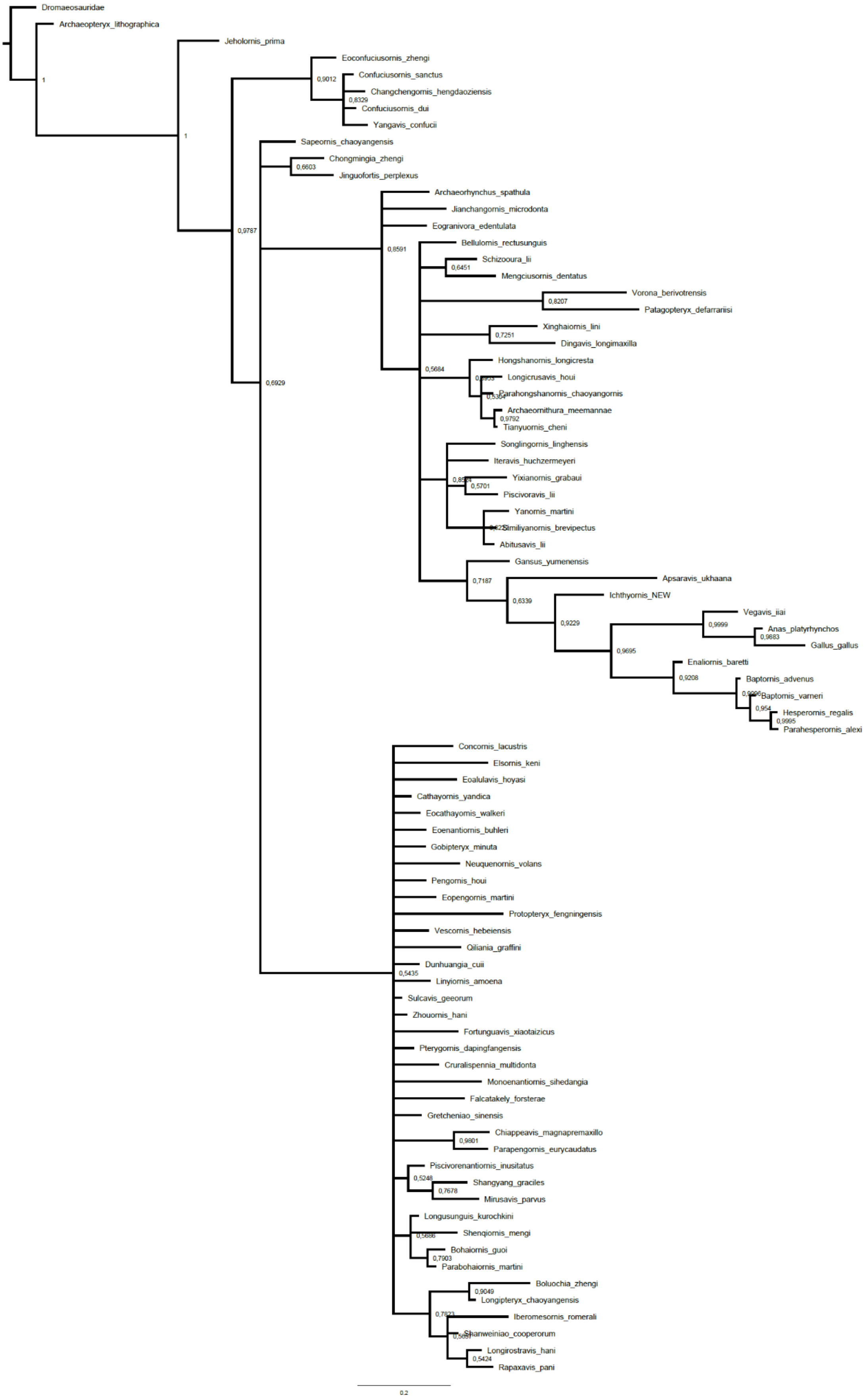

2377 **Supplemental Tree 16**

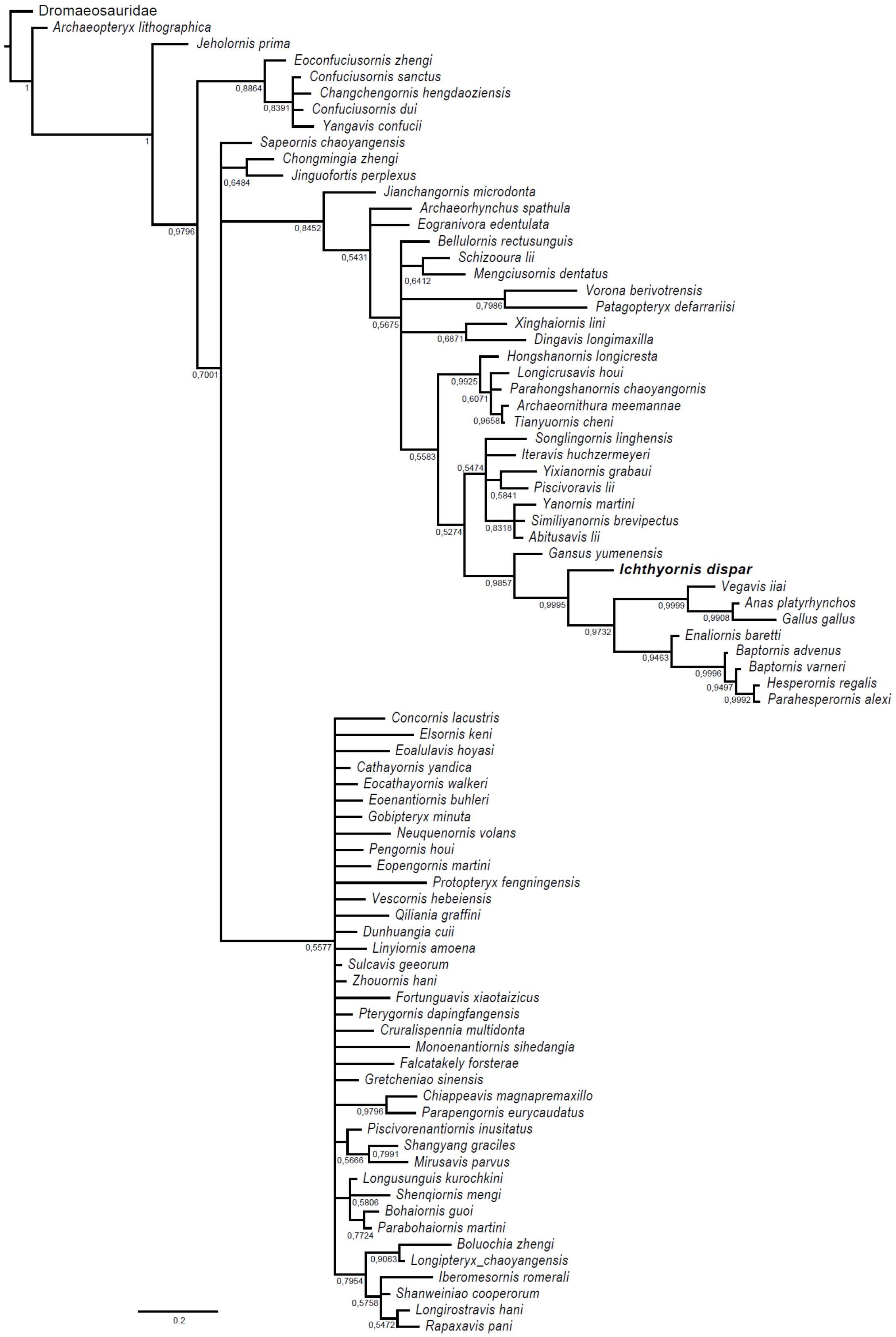

| Species | Specimen | Hum. Length | Ulna Length | Radius Length | CMC length | Phalanx II:1 | Phalanx II:2 | Manus lentgh | Femur length | Tibiotarsus length | TMT length | Max pedal phalanx length | Max pedal digit length | Brachial Index |
|---|---|---|---|---|---|---|---|---|---|---|---|---|---|---|
| *Anser albifrons* | UMZC 242.EA | 150.47 | 144.51 | 140.21 | 86.47 | 37.49 | 36.87 | - | 71.8 | 130.58 | 72.96 | 28.36 | 77.25 | 1.04 |
| *Anas platyrhynchos* | UMZC 224 | 97.83 | 83.2 | 76.2 | 57.7 | 22.48 | 19.1 | 98.2 | 55.72 | 96.72 | 50.85 | 24.03 | 55.36* | 1.18 |
| *Gallus gallus* | UMZC 402 | 70.5 | 69.74 | 63.95 | 34.94 | 14.34 | 14.18 | 62.31 | 82.29 | 115.12 | 75.49 | 17.12 | 42.84* | 1.01 |
| *Fulica atra* | UMZC 298.a | 82.24 | 72.37 | 67.73 | 43.07 | 16.13 | 15.64 | 74.48 | 60.09 | 110.28 | 63.51 | 29.12 | 89.1* | 1.14 |
| *Sterna sumatrana* | UMZC 269.BA | 44.62 | 51.7 | 48.54 | 28.6 | 16.61 | 15.79 | 55.2 | 21.61 | 37.54 | 16.88 | - | 20.31 | 0.86 |
| *Chroicocephalus novaehollandiae* | UMZC 274.c | 84.95 | 94.79 | 91.12 | 49.92 | 23.7 | 20.77 | 88.12 | 40.61 | 83.03 | 53.37 | 17.37 | 42.85 | 0.90 |
| *Puffinus lherminieri* | UMZC 284.AA | 64.51 | 65.17 | 60.37 | 33.83 | 17.55 | 19.11 | 67.84 | 23.4 | 60.58 | 36.65 | - | 42.21 | 0.99 |
| *Phaethon lepturus* | UMZC 261.CA | 80.27 | 87.44 | 82.41 | 42.07 | 24.27 | 21.23 | 76.09 | 30.34 | 43.82 | 27.59 | - | 37.08 | 0.92 |
| *Podiceps auritus* | UMCZ 206.B | 76.11 | 67.95 | 67.34 | 33.45 | 14.41 | 10.07 | - | 33.16 | 85.56 | 43.39 | 18.56 | - | 1.12 |
| *Podica senegalensis* | UMCZ 209.A | 56.64 | 43.17 | 38.96 | 37.58 | 13.25 | 10.97 | 59.81 | 49.57 | 80.09 | 45.12 | 20.47 | - | 1.31 |
| *Gavia arctica* | UMCZ 203 | - | 150.37 | 148.53 | 96.04 | 29.88 | 26.22 | - | 57.07 | - | 88.08 | 54.46 | - | - |
| *Fratercula artica* | UMCZ 191.B | 61.29 | 48.94 | 46.96 | 32.12 | 14.41 | 17.54 | - | 36.21 | 62.11 | 26.38 | 12.08 | - | 1.25 |
| *Oceanodroma leucorhoa* | UMCZ 287.D | 34.93 | 34.34 | 33.92 | 19.78 | 9.75 | 10.45 | - | 15.58 | 36.59 | 22.9 | 8.44 | 22.84 | 1.02 |
| *Rhyncops flavirostris* | UMCZ 268.a | 63.67 | 76.9 | 74.98 | 37.25 | 22.03 | 21.36 | - | 28.38 | 45.91 | 24.79 | 6.83 | - | 0.83 |
| *Larus fuscus* | UMCZ 274.1 | 118.66 | 132.34 | 128.36 | 68.14 | 32.28 | 28.66 | - | 51.45 | 101.67 | 57.51 | 18.98 | 48.44* | 0.90 |
| *Chlidonias niger* | UMCZ 271.B | 41.97 | 50.49 | 48.91 | 27.83 | 16.28 | 16.04 | - | 19.56 | 35.01 | 17.47 | 6.43 | 10.93 | 0.83 |
| *Alca torda* | UMCZ 187.AA | 58.8 | 46.3 | 44.11 | 32.25 | 16.32 | 15.74 | - | 35.04 | 57.28 | 25.51 | 12.96 | - | 1.27 |
| *Uria sp.* | UMCZ 192.A | 91.13 | 76.98 | 73.13 | 49.97 | 24.68 | 27.2 | - | 51.77 | 94.97 | 40.32 | 18.71 | 62.06 | 1.18 |
| *Charadrius rubricollis* | UMCZ 320.I | 39.69 | 41.36 | 34.47 | 23.84 | 11.85 | 9.02 | - | 27.18 | 47.1 | 27.51 | 7.64 | 20.84 | 0.96 |
| *Rallus striatus* | UMCZ 304.A | 49.16 | 44 | 41.18 | 28.44 | 9.13 | 10.29 | - | 49.88 | 69.18 | 41.68 | 14.17 | 39.51* | 1.12 |
| *Sterna hirundo* | Uncatalogued | 75.81 | 84.99 | 82.31 | 46.35 | 21.81 | 20.24 | - | 35.41 | 70.46 | 46.34 | 14.8 | - | 0.89 |

**Supplemental table 1. Forelimb and hindlimb measurements of comparative extant bird specimens.** Asterisks (*) denote measurements that might be unreliable due to breakage or surrounding soft tissue but are included for completeness purposes. All measurements are in mm. - = not measurable.
